# Endotherms trade body temperature regulation for the stress response

**DOI:** 10.1101/2023.01.09.523310

**Authors:** Joshua K.R. Tabh, Mariah Hartjes, Gary Burness

## Abstract

Responding to perceived threats is energetically expensive and can require animals to curtail somatic repair, immunity, and even reproduction to balance energy ledgers. Among birds and mammals, energetic demands of thermoregulation are often immense, yet whether homeostatic body temperatures are also compromised to aid the stress response is unknown. Using data sourced from over 60 years of literature and 24 endotherm species, we show that exposure to non-thermal challenges (e.g. human interaction, social threats) caused body temperatures to decrease in the cold and increase in the warmth, but particularly when species-specific costs of thermoregulation were high and surplus energy low. Biophysical models revealed that allowing body temperature to change in this way liberated up to 24% (mean = 5%) of resting energy expenditure for use toward coping. While useful to avoid energetic overload, such responses nevertheless heighten risks of cold- or heat-induced damage, particularly when coincident with cold- or heat-waves.

## Introduction

About 2000 years ago, the Greek physician Galen observed that the body temperatures of his patients tended to rise after psychologically distressing experiences [1]. Given the acute nature of these rises, Galen termed their occurrence “ephemeral fevers” [2] and speculated, like those after him (e.g. Ibn Sina) that they emerged from an abundance of humor [3,4]. “Why” (i.e., functionally) they might emerge at all, however, was apparently left unquestioned.

Numerous studies have now shown that Galen’s “ephemeral fever” is not just restricted to humans. Instead, many rodent, non-hominid primate, and even bird species appear to change their body temperatures after exposure to common disturbances, such as transportation or restraint [5–7]. Moreover, accumulating evidence suggests that the term “ephemeral fever” is likely a misnomer, with body temperatures of some individuals *declining* in response to some stressors rather than rising [8,9]. That these thermal responses to stressors exist across varied taxonomic groups and may differ under certain conditions strongly intimates a possible function, or, perhaps, a conserved and inherent nature (i.e., a constraint) behind their occurrence.

Recently, we proposed that changes in body temperature following stress exposure may be understood as consequences of an allocation trade-off between thermoregulation and the stress response ([10–11]; similar to ref. [12]). Under this hypothesis, fluctuations in body temperature that accompany the stress response represent the outcomes of reducing energy expenditure toward thermoregulation in favor of expenditure toward coping. Unlike previous hypotheses, this particular hypothesis provides an explanation for why body temperature might either rise *or* decline after a stress exposure, since relaxing thermoregulation should merely result in body temperature drifting toward ambient (particularly in the cold) or in the direction of accumulating metabolic heat (i.e. from activation of the stress response and reduced windows for heat loss; Fig. 1). Over the past few decades, a handful of studies have indeed supported a role of ambient temperature in shaping the direction and magnitude of stress-induced thermal responses [13–15]. Moreover, others extending over several decades have already shown that thermoregulatory investment can well be shifted to cope with other environmental challenges, such as water deprivation (most notably in dromedaries, *Camelus dromedarius,* ref [16]) and food limitation [17–18]. Together, these findings provide intriguing, albeit sometimes conflicting, support for our trade-off hypothesis. Nevertheless, whether an influence of ambient temperature holds across studies, species, and contexts (i.e. stressors), remains unknown, but would have importantly implications for understanding how, and at what cost, endotherms cope with non-thermal challenges.

**Figure 1.**
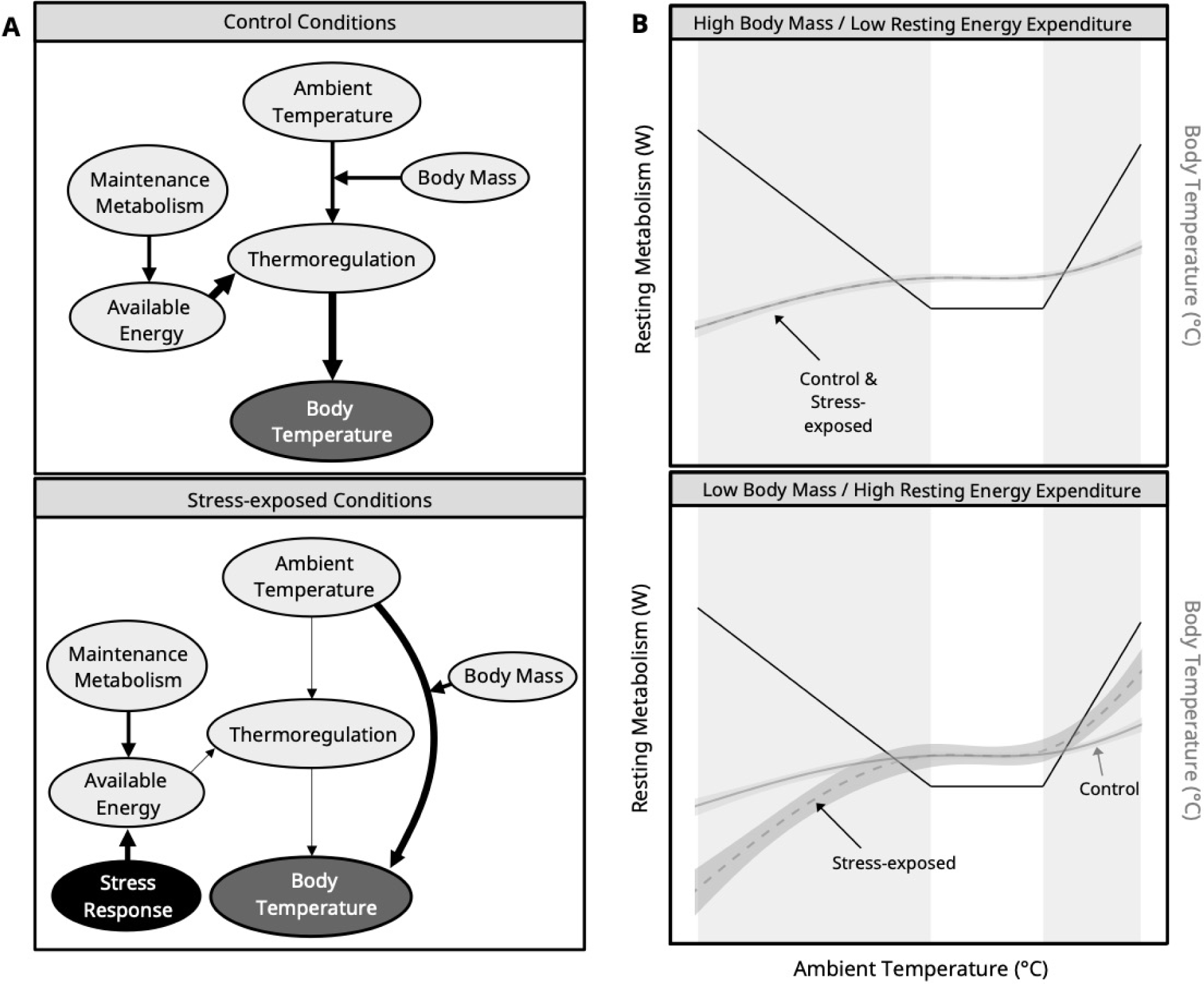
Trade-off model explaining the outcome of stress-induced changes in body temperature. **A |** “Available energy” indicates surplus energy available for allocation toward biological processes after accounting for maintenance metabolism (i.e. basal metabolism). Thickness of lines indicates the magnitude of influence that one variable exerts on another, in the direction of the arrow. **B |** Solid black lines indicate resting metabolism across ambient temperature for a hypothetical, terrestrial endotherm. Grey bands indicate ranges of ambient temperature where expenditure toward thermoregulation is expected. Solid grey lines indicate body temperature across ambient temperature for the same hypothetical endotherm during control (or unstressed) conditions and the dashed grey lines indicate body temperature across ambient temperature during stress-exposed conditions. Ribbons around grey lines are added to indicate that true body temperature responses to control and stress-exposed conditions will vary.

Here, we used data describing body temperature responses to stressors across 24 terrestrial endothermic species to test for an allocation trade-off between thermoregulation and the stress response (outlined in Fig. 1). Non-terrestrial endotherms were excluded from our study owing to a lack of available data, but implications with respect to these species are discussed. Under our trade-off hypothesis, we made three predictions. First, although body temperatures should typically increase after a stressor owing to accumulation of metabolic heat, these increases should be exacerbated in the warmth and forgone, or even reversed, in the cold. Second, the degree to which ambient temperature influences stress-induced thermal responses should depend on body mass, with smaller species/individuals becoming more heterothermic across observed temperature ranges, owing to higher surface area to volume ratios and greater susceptibilities to the cold (i.e. higher lower critical temperatures and lower thermal inertia). Third, how stress-induced thermal responses change across ambient temperature should depend on the relative amount of energy available for allocation toward both thermoregulation and the stress response (as per ref. [19]). More specifically, species with relatively high metabolic rates (thus indicating low energy availability; see ref. [20]) should be more likely to decrease their body temperature in the cold, and increase their body temperature in the warmth following a stressor than those with relatively low metabolic rates (indicating high energy availability).

Our findings indicate that body temperature regulations can be, and often is, compromised by non-thermal challenges among bird and mammal species. These findings raise important questions about risks posed by non-thermal stressors (e.g. human interaction) experienced during increasingly-common extreme climate events. Further still, we argue that they add to our growing understanding of how thermoregulatory strategies may be altered to avoid homeostatic disruptions or states of energetic overload in challenging environments (e.g. refs [16–18]).

## Methods

### Literature search

We searched primary literature for data describing the magnitude and context of stress-induced changes in body temperature across endotherms using Web of Science^TM^. To simplify interpretations of our findings, we considered only literature pertaining to terrestrial endotherms, excluding humans (owing to confounding effects of clothing on costs of thermoregulation).

Query terms for our search were: “stress”, “body temperature” and “NOT heat stress”. In total, 3022 studies were returned by our query. From those returned, we scanned abstracts for evidence that: (1) body temperature responses to stress exposure were indeed measured, and (2) the species in which body temperature responses were measured was endothermic, terrestrial, and non-human; those with abstracts lacking such evidence were removed from consideration. Next, because stress-induced changes in core body temperature rather than surface body temperature are: (1) arguably most relevant to whole animal energy expenditure and (2) subject to most investigation over the past century, studies describing stress-induced responses at surface body temperatures were also omitted. From all remaining studies (Supplemental Fig. 55), we scanned reference lists for relevant literature that had not been returned by Web of Science. Again, abstracts of these studies were reviewed for evidence that stress-induced changes in core body temperature were indeed measured and were retained or excluded accordingly (n = 31 studies added; Supplemental Fig. 55). Together, all searches and data collection were conducted between 28 April and 27 July, 2020, with publication dates in our data-set ranging from March, 1958 [21] to January, 2020 ([22]; mean publication year [rounded to nearest integer] ± s.e.m. = 2003 ± 0.879).

### Data extraction and inclusion criteria

From each study identified for possible inclusion (n = 136; Fig. 1), we harvested the following data:

1. The average stress-induced change in body temperature across a sample population (Δ°C; ± s.d. or s.e.m). In some studies, stress-induced thermal responses were monitored and reported across multiple study groups (i.e. per sex, air temperature treatment, discrete time period, etc.). In these cases, we chose to treat data from each group as an independent observation nested within a study. If multiple measurements of change were reported for a given study group, we collected only the value demonstrating the greatest increase or greatest decrease relative to baseline body temperature for that group.
2. The number of individuals measured.
3. The study species and family.
4. The age class of the sample population (factorial; adult, juvenile, or mixed, according to age of reproductive maturity). If age class was not reported, we assumed that the study population was comprised solely of adults.
5. Sex of individuals in sample population (factorial; female, male, or mixed). If sex was not reported, we assumed it to be mixed.
6. The season of study, categorized as spring, summer, fall, or winter. If the season of study was not provided, we set it as “not available (NA)”.
7. The average mass of individuals in the study population (grams, *±* s.d. or s.e.m; s.e.m. ≈ 0 [0.0001] if missing). If a range of masses was provided (i.e. in n = 220 observations across n = 84 studies), we calculated the theoretical median and used it in place of average mass, with the standard deviation equaling the mass range divided by 4. Where average mass of individuals was not reported, we extracted mean values for the species, sex, and age-class of interest (if provided; see categories 4-5 above) from primary literature (n = 285 observations across n = 43 species groupings and n = 117 studies). Categories (4) and (5) describe our approach when data on sex and age-class were missing.
8. Air temperature during study (in °C, ± s.d. or s.e.m). For field studies lacking measures of air temperature, we obtained average values from nearby weather-stations (< 50 km from the study location) across the reported duration of study (i.e. according to specified date and time ranges) where available (n = 13 observations from n = 3 studies; indicated in data files). Studies reporting stress-induced thermal responses in media other than air were discarded and those that did not report variation around air temperature means were assigned an s.e.m. of ≈ 0 (0.0001; n = 57 observations across n = 28 studies).
9. Latency between pre-stress exposure and post-stress exposure temperature measurements (seconds). Latency values greater than 8 hours (28800 seconds; > 3 s.d. above the mean [3544 seconds]) were excluded from our analyses (n = 4 observations).
10. Technique used to measure body temperature (intra-peritoneal telemetry or thermocouple [rectal/cloacal/throat]).
11. Whether pharmaceuticals other than saline were administered to study animals before or during stress exposure (binomial; true/false).

For most studies, we extracted these data from the article text or from article figures using the open-source software FIJI [23]. For all remaining studies, data were requested from article authors directly. Once data collection was complete, all studies lacking data from categories 1-4 and 8-10, and/or satisfying category 11 were discarded. Together, our final sample included 165 observations across 68 studies, 24 species, and 30 species strains (n = 12 avian; n = 18 mammalian; average number of observations/species ± s.d. = 6.98 ± 19.00; phylogeny provided in supplemental information). Final samples sizes and summaries of sample exclusions following the preferred reporting items for systematic reviews and meta-analyses (PRISMA) are outlined in Supplemental Fig. 55.

Types of stressors applied across our sample were varied and numerous, ranging from transportation (n = 4 studies), social defeat (n = 10), restraint (mechanically [n = 31 studies] or by handling [n = 43 studies]), exposure to a novel environment (n = 32 studies), and others.

#### Metabolic rate data

We were interested in testing whether a species’ relative energetic expenditure at rest (and thus, their remaining energy available for allocation) influences the magnitude of its stress-induced change in body temperature. To calculate relative energetic expenditure for species or species strains included in our data set (in publications that met our inclusion criteria: see “Data extraction and inclusion criteria” above), we again used Web of Science^TM^ to search for primary literature reporting either their basal or resting metabolic rates (“BMR” and “RMR” respectively). Measures of metabolic rate within each study were categorized as “basal” according to whether they were: (1) collected at thermoeneutrality, (2) collected post absorption, and (3) obtained during the circadian rest phase of the target species (i.e. criteria 1-3 from McKechnie and Wolf [24]). Measures that satisfied only criteria 1-2 were categorized as “resting” and all others were discarded. To enhance our search for metabolic data, we replicated our queries using Google Scholar^TM^.

Since both BMR and RMR likely vary across seasons, we sought to obtain metabolic data relevant to the season at which stress-induced changes in body temperature were observed (again, see “Data extraction and inclusion criteria” above). In most cases, however, the season in which metabolic expenditure and/or the stress-induced thermal response were measured were not reported (n = 66 studies of 68). Moreover, to our knowledge, some species lacked any primary data on basal or resting metabolism at all. Thus, where necessary, measures of BMR and/or RMR were obtained from available, but potentially mismatched, seasonal studies, from the most closely related species studied (n = 5 species; relatedness determined using a consensus phylogeny approach described below), or from order-specific allometric relationships (n = 2 species; 1 chiropteran species and 1 passerine species). Following collection, all metabolic data were converted to a common metric (watts, “W”) using conversion coefficients described in the appendix.

To account for effects of body mass on energy expenditure across species, we obtained ordinary residuals from a Bayesian linear mixed effects model (LMM) predicting natural log-transformed BMR or RMR as a function of natural log-transformed species mass (mean in g; prior = ***N***[0.67, 0.1]). Type of measurement was included in this model as a group-level intercept (i.e. “basal” or “resting”; group-level predictor; prior = Г[1, 1]; model coefficients: Intercept = −3.527 [−4.508, −1.873]; log mass = 0.668 [0.599, 0.734]; measurement type = 0.084 [−0.950, 0.869]) but phylogeny was not included to avoid redundancy in our subsequent analyses. Residuals from our model were then categorized as either positive or negative, with positive values for a given species indicating a relatively high energy expenditure at rest, and negative values indicating a relatively low energy expenditure at rest. Categorizing data in this way allowed us to broadly and conservatively evaluate how relative energy expenditure, and thus, energy availability, influenced stress-induced thermal responses while circumventing over-parameterization of our statistical model (described below).

### Effect size calculation

To ensure that our data were statistically comparable among studies and species, we converted our estimates of absolute changes in body temperature to relative changes in body temperature. This was achieved by calculating a response ratio as follows:

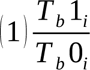

where T_b_1 represents an average minimal, or an average maximal body temperature of study population *i* that was observed after a stress exposure, and T_b_0 represents the body temperature of study population *i* before a stress exposure treatment. For studies that also recorded the body temperatures of control (or “unstressed”) individuals throughout experimentation, however, we used the following modified equations:

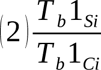

where T_b_1_S_ again represents an average minimal, or an average maximal stress-induced body temperature of the stress-exposed population *i*, and T_b_1_C_ represents the average body temperature of the control population at the equivalent time point.

To estimate uncertainty around our response ratio values, we calculated a pooled standard error using one of the following equation:

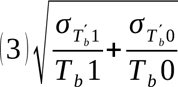

or:

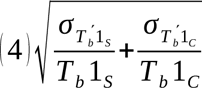

where equation (3) applies to studies without a control population, equation (4) applies to studies with a control population. In these equations, σ_x_ represents the standard error of a given subscripted variable *x*, and all other variables remain as previously described. In studies where a standard error around the stress-induced change in body temperature was reported, we assumed it to be a pooled estimate and used it in place of that derived from an equation above.

Finally, to both normalize our response ratio estimates and centre them on zero, we calculated their natural logarithms and used these transformed values as our final estimates of effect size (i.e. the log-response ratio, or “lRR”). Effect sizes for studies with and without control populations were visually compared to confirm that their distributions were similar.

#### Phylogeny

To estimate phylogenetic relationships among species included in our data-set, we extracted two sets of 1000 trimmed phylogenetic trees – one for mammals, and one for birds – from VertLife [25] and BirdTree [26] respectively. These tree-sets were drawn from pseudo-posterior distributions of relaxed-clock models to infer phylogeny from genetic and fossil data (number of genes: mammals = 31[25]; birds = 15 [26]; avian fossil backbone from ref. [27]). From each tree-set, we then estimated class-specific consensus trees using “TreeAnnotator” in Beast 2 [28], with a 25% burn-in, posterior probability limit of 1.0, and with node heights representing median values. Once our consensus trees were established, we used them to build variance-covariance matrices approximating the relative relatedness between species (via the R package “ape” [29]; matrices used in subsequent analyses). For simplicity, relatedness between all pairs of birds and mammals was assumed to be negligible and set to zero. Phylogeny estimation and consensus trees are detailed in the appendix (section 5).

### Statistical analyses

All of our statistical analyses were conducted in R version 4.03 [30] and figures were produced using the R package “ggplot2” [31]. For each model, we used Hamiltonian Monte Carlo (HMC) sampling in the R package “brms” [32] with 2500 burn-in iterations, 15000 sampling iterations, and a thinning factor of 10 across four HMC chains. Models were validated by visually diagnosing residual distributions and paired posterior coefficient estimates. Gelman-Rubin statistics 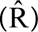 for each of our models fell between 0.95 and 1.05, suggesting that chain mixing was consistently adequate, and effective sample size to sample size ratios consistently exceeded 0.75, confirming minimal autocorrelation between HMC chain draws. Versions for all R packages used are reported in the appendix.

To confirm that our results were not biased by small sample sizes among our collated studies, we re-ran of our model using an adjusted log response ratio designed to correct for sample-size related error (“RR^Δ^”; [33]). Results from this model were then compared with our original model at 80% and 95% highest density credible intervals (HDIs). Outcomes of both models were highly similar (see supplemental information; Table S24), suggesting that sample variation in sample size among our collected studies was unlikely to influence our reported findings.

#### Modeling

We were interested in testing whether the direction and magnitude of stress-induced changes in body temperature depended on an individual’s thermoregulatory demand (described below) and relative energy available for investment in thermoregulation and the stress response (here, residual RMR after controlling from body mass, categorized as positive or negative). Clearly, thermoregulatory demand should vary according to air temperature, with the degree of variation being influenced by a species’ body size (here, proxied by body mass). Further, the relative energy available to a species is predicted to be inversely proportional to it’s relative expenditure (i.e. at rest; see [20]), barring other homeostatic compensations (refer to Discussion). Accordingly, we tested whether air temperature, body mass, relative energy expenditure, and the interactions between both air temperature and body mass, and air temperature and relative energy expenditure predicted the direction and magnitude of stress-induced changes in core body temperature. To achieve these ends, we constructed a Bayesian linear mixed effects model (LMM) with our log response ratio as the response variable, and air temperature (°C), the natural log of body mass (g), relative energy expenditure (factorial; positive or negative, indicating high or low expenditure for a given body mass respectively), the linear interaction between air temperature and body mass, and the linear interaction between air temperature and relative energy expenditure as population-level predictors. Given that both body mass and air temperature can vary considerably within studies, however, we modeled each parameter as latent, with their true values being unknown but falling within our observed values ± our observed error. Finally, both air temperature and the log of body mass were mean-centered to better facilitate interpretation of our model intercept.

To account for recovery or ceiling/floor effects in our analysis, we included the latency between baseline and stress-induced body temperature measurements as a population-level predictor in our LMM. Here, we assumed that the temporal effects of stress exposure on body temperature were non-linear, with body temperature rapidly changing, then plateauing or recovering across time [34–36]. For this reason, the latency between baseline and stress-exposed measurements was modeled as a second-order polynomial. Similar to air temperature and the log of body mass, latency between measurements was mean-centered to ease interpretation of our intercept. While the duration of a stressor may also influence the amount of energy allocated toward the stress response, estimates of this duration were correlated with our latency variable that was already included in our model (Bayesian LMM: Intercept = −349.46 [−688.85, 3.23]; effect of stressor duration [s; mean-centred] = 0.18 [0.09, 0.26]; study identity [group-level intercept] = 1533.48 [1231.85, 1852.27]; priors: Intercept = ***N***[0, 300]; stressor duration = ***N***(0, 10); study identity = exponential[0.01]; n = 149). Further, measuring stressor duration is often unclear (e.g. for stressors such as social defeat). As such, stressor duration was not included in our analysis. Finally, to correct for an influence of the method used to measure body temperature on stress-induced thermal responses, and to correct for the possibility of age effects, we included measurement technique (factorial; telemetry or thermocouple) and age of individuals in the study population (integer; juveniles only = −1, both juveniles and adults = 0, adults only = 1) in our model as population-level predictors. Species, study identity, and phylogeny (see section 6) were included as group-level predictors to control for non-independence between each factor.

For our LMM, we used weakly informative priors that assumed that log-responses ratios of ⪅ −1 and ⪆ 0.5 (e.g. representing a 24°C decrease and 10°C increase in body temperature respectively at baseline temperatures of 38°C) were unlikely. Specifically, priors for air temperature, log body mass, and the first-order effects of latency to body temperature measurement were set as ***N***(0, 0.1), while those for the interaction between log body mass and air temperature, and the interaction between relative energy expenditure and air temperature were set as ***N***(0, 0.05). Because we were less certain about the range of possible age-related and second-order time-related effects, priors for age class and the second-order effect of time were set as ***N***(0, 0.25). Next, because both relative energy expenditure and measurement methodology were factorial, priors for these parameters was set as skew-normal (***SN***) with ξ= 0, ω= 0.1, and α = −2.5. We chose to use a ***SN*** distribution to capture the increased likelihood of observing negative response ratios than positive response ratios (an inherent property of natural logarithm transformation). Importantly, however, a ***SN*** distribution was not assumed for priors of continuous predictors where absolute slope estimates were expected to be small. Lastly, priors for all group-level predictors, and σ were weak and flat (−∞ - ∞), or gamma-distributed (group-level predictors: Г[1.5, 1.0]; σ: Г[1.0, 1.0]).

Four observations (out of 165; ∼2.5% of our data) were removed from our analysis, owing to their abnormally large effect sizes relative to the mean (> 3.5 times the sample s.d.; absolute change in body temperature > 10°C). To account for the possibility that posterior distributions for our model coefficients were non-normal, we calculated modes for all coefficients in place of means, and highest posterior density credible intervals (80% and 95%) around these modes [37]). All calculated means were marginalized unless stated otherwise, and absolute changes in body temperature were calculated from log-response ratios and assumed a baseline body temperature of 38°C.

Where reported, positive log-response ratios represent a stress-induced increase in body temperature, while negative log-response ratios represent a stress-induced decrease.

#### Estimating change in energy expenditure attributed to the stress-induced thermal response

A critical assumption of our trade-off hypothesis is that, in response to a stressor, allowing body temperature to decrease in the cold or increase in the warmth liberates a meaningful amount of energy that can then be allocated toward coping with that stressor. With this assumption in mind, we sought to estimate the relative extent to which stress-induced changes in body temperature reduced expenditure toward thermoregulation among our sampled vertebrates. To do so, we first estimated resting metabolic rate of sampled animals at their known housing- and baseline body temperatures (henceforth “baseline RMR”) following established biophysical models [38], and by using the R package nicheMapR [39]. These estimates represented energy expenditure required to maintain thermal balance and assumed that: (1) baseline body temperature was held constant, (2) loss of body heat to the ambient environment occurred by radiation, forced convection, and free convection, but not conduction, and (3) the rate at which body heat was lost to the ambient environment was inversely related to the resistance of both the core body itself and the insulative layer of plumage or pelage [38]. For our estimates, we assumed that all animals were ellipsoids with volumes (m^3^) equal to their mass in g (body density = 1000 kg/m^3^; [39]), and with semi-major axes being 1.2 and 1.3 times longer than semi-minor axes for birds and mammals respectively. Further, we also assumed that: (1) pelage or plumage depth of sampled animals was equal the logarithm of their body mass (g) in cm (e.g. pelage depth = 1 and 2 cm for a 10 and 100 g mammal respectively), (2) thermal conductivity of an animal’s pelage or plumage was 0.0272 W/m-°C [38], and (3) wind speed during sampling was negligible (0.1 m/s). Following these estimates, we then re-estimated resting metabolic rates across the same ambient temperatures, but with body temperatures representing those measured after stress exposure (henceforth, “stress-induced RMR”), rather than at baseline. The difference between stress-induced and baseline RMR values then represented the total energy saved or lost by adjusting body temperature in the manner observed. These change values are expected to be conservative, owing to the exclusion of temperature effects of metabolic processes themselves (i.e. through q_10_ effects; [40]).

## Results

### Stressors trigger heterothermy and reduce costs of thermoregulation, particularly in small endotherms

Exposure to stressors coincided with, on average, an increase in body temperature of 0.64°C (indicated by our model intercept; β ≈ 1.67⨯10^-2^ [80%: 4.41⨯10^-3^, 2.73⨯10^-2^; 95%: −1.78⨯10^-3^, 3.50⨯10^-2^]). Both the magnitude and direction of these stress-induced responses were clearly influenced by ambient temperature, as predicted by our trade-off hypothesis (β ≈ 7.70⨯10^-4^ [80%: 3.10⨯10^-4^, 1.23⨯10^-3^; 95%: 4.00⨯10^-5^, 1.44⨯10^-3^]; Fig. 2). At low ambient temperatures, where costs of warming are highest, body temperatures was more likely to decrease after a stressor than at high ambient temperatures. By contrast, at high ambient temperatures, where costs of warming are lowest and those of cooling highest, body temperatures was more likely to increase after a stressor, and to a greater degree, than at low ambient temperatures (Fig. 2).

**Figure 2.**
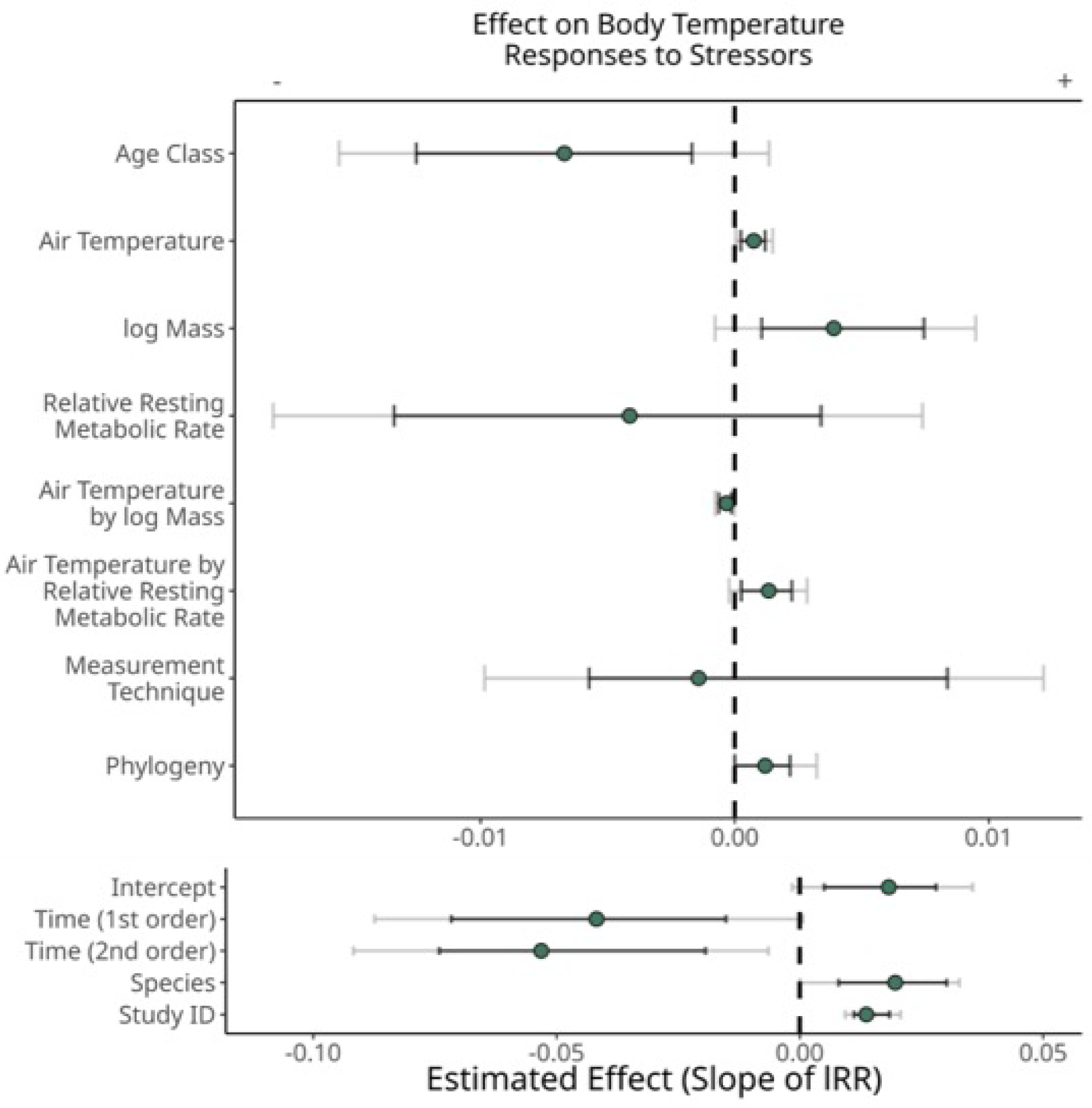
Factors influencing stress-induced changes in core body temperature among terrestrial endotherms. Dots represent the estimated effect of a given parameter on the magnitude and direction of stress-induced changes in body temperature (measured by a log response ratio, “lRR”). “Time” indicates latency to a maximum or minimum body temperature measurement after stressor onset; “relative resting metabolic rate” indicates the residuals of log-transformed resting metabolic rate (W) regressed against log-transformed body mass (g), then categorized as positive or negative. Negative slopes represent a tendency to reduce core body temperature, while positive slopes represent a tendency to increase core body temperature. Black grey represent 80% highest posterior density intervals credible intervals (HPDIs) and black whiskers represent 95% HPDIs.

An influence of ambient temperature on stress-induced responses was not ubiquitous, and clearly depended on an individual’s body mass (interaction between air temperature and body mass: β = −4.00⨯10^-4^ [80%: −6.40⨯10^-4^, −1.50⨯10^-4^; 95%: −7.80⨯10^-4^, −1.00⨯10^-5^]; Fig. 3a). Given that body mass is capable of dictating both costs of thermoregulation and susceptibilities to heat and cold among animals (i.e. the breadth of their thermoneutral zones), this interactive effect between mass and ambient temperature was expected under our trade-off hypothesis. Among individuals with low body masses (< 222 g, our sample mean), exposure to a stressor tended to induce a state of heterothermy, with body temperatures decreasing after a stressor in cool temperatures and increasing after a stressor in the warmth (Fig. 3a). For example, when ambient temperatures were below our sample mean (20.1°C), body temperatures of these individuals fell by an average of 0.54°C after a stress exposure, but when ambient temperatures were above our sample mean, body temperatures of the same individuals rose by an average of 0.79°C (Fig. 3b). Thermal responses of individuals with relatively high body masses (> 222 g), however, were less nuanced. Specifically, body temperatures of these individuals tended to increase after a stressor regardless of the current ambient temperature (Fig. 3; mean change in body temperature = +2.02°C at below-average air temperatures, and +1.18°C at above-average air temperatures). Regardless of body mass and ambient temperature, however, biophysical models predicted that how body temperatures responded to stressors across these animals reduced energetic costs of maintenance by on average of 4.8%, and up to 24% of resting metabolic rate (Fig. 3c).

**Figure 3.**
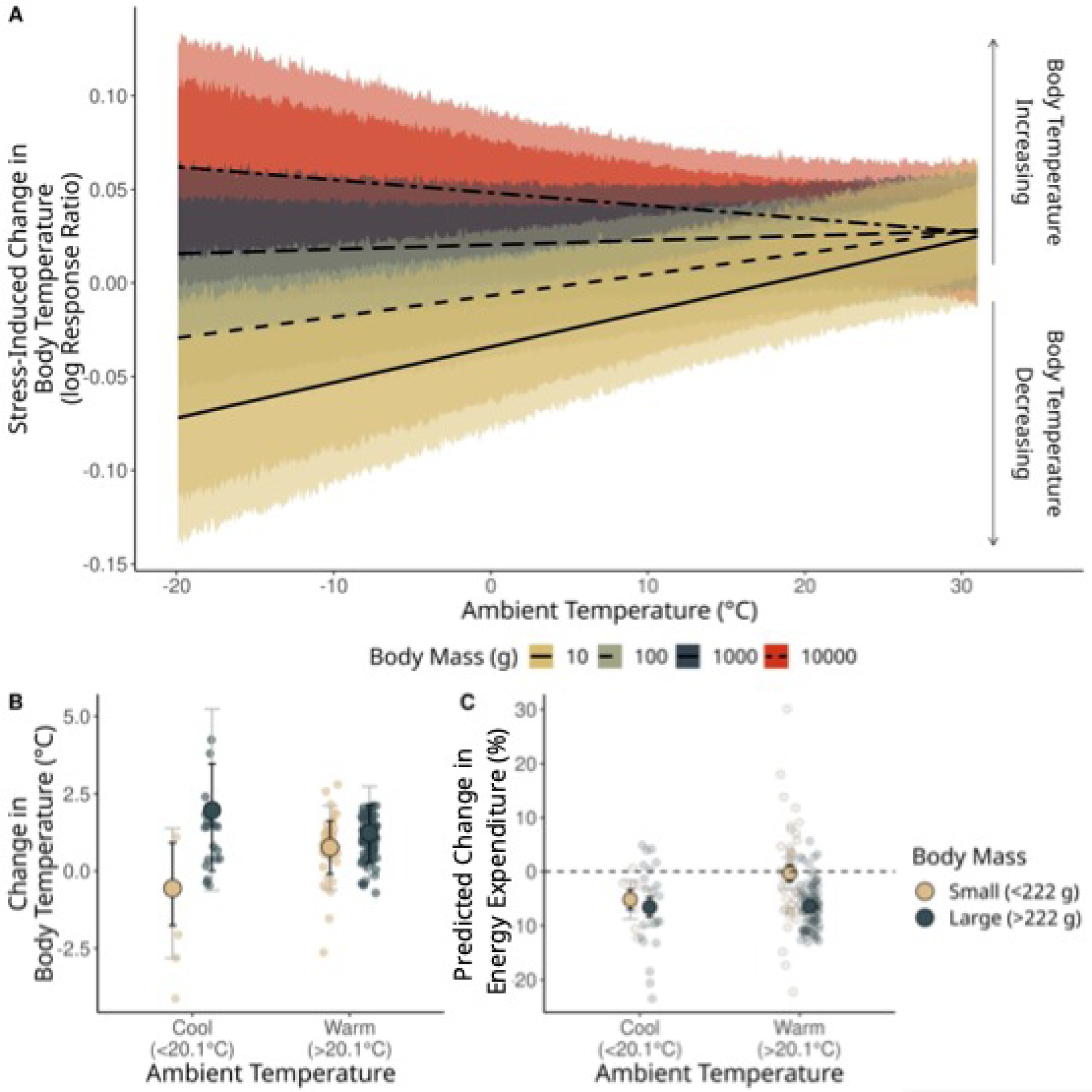
Effect of air temperature and body mass on the magnitude and direction of stress-induced changed in core temperature. **A.** Negative log response ratios represent stress-induced declines in body temperature, while positive log response ratios represent stress-induced increases in body temperature. Trend lines represent predicted relationships for an animal of a given body mass (g), while holding all other model covariates (shown in Fig. 2) at their averages. Dark ribbons represent 80% highest posterior density intervals (HPDIs) and pale ribbons represent 95% HPDIs. **B.** Large dots indicate average changes in body temperature for species above and below mean body mass for our sample (∼222g; beige and green respectively), and at ambient temperatures above and below the average across studies (20.1°C). Mean changes in body temperature are calculated from log-response ratios and assume a baseline body temperature of 38°C. Small dots represent raw data values and error bars indicate 80% and 95% highest posterior density intervals (HPDIs; black and grey respectively). **C.** Large dots display mean changes in resting energy expenditure attributed to stress-induced thermal responses, as predicted by biophysical models. Again, small dots represent raw data values and error bars indicate 80% and 95% highest posterior density intervals (HPDIs; black and grey respectively).

Independent of air temperature, larger animals were marginally more likely to increase their body temperature after a stress exposure, and by a greater degree, than smaller animals (main effect of body mass: β = 4.25⨯10^-3^ [80%: 1.26⨯10^-3^, 7.68⨯10^-3^; 95%: −1.28⨯10^-3^, 9.17⨯10^-3^]; Fig. 2).

### Limited surplus energy drives heterothermic responses to stressors

Body temperatures tended to increase after a stress exposure at moderate-to-high ambient temperatures (indicated by a positive log response ratio at ambient temperatures > approximately 10°C), regardless of whether a species expended relatively low, or relatively high amounts of energy at rest (i.e. for their given mass; Fig. 4). The magnitude by which these increases occurred also did not depend upon a species’ relative energy expenditure at rest (main effect of relative resting metabolic rate: β ≈ −3.98⨯10^-3^ [80%: −1.34⨯10^-2^, 3.43⨯10^-3^; 95%: −1.76⨯10^-2^, 8.47⨯10^-3^]; Fig. 2). In the cold (ambient temperatures < approximately 10°C), however, species expending relatively high amounts of energy at rest (and thus, having less energy available for allocation elsewhere) tended to decrease their body temperatures, or allow them to fall, relative to those expending comparably little energy at rest (interaction between air temperature and relative energy expenditure: β = 1.38⨯10^-3^ [80%: 2.60⨯10^-4^, 2.23⨯10^-3^; 95%: −2.70⨯10^-4^, 2.79⨯10^-3^]; Fig. 4a). This effect was predicted by our trade-off hypothesis. Since baseline body temperatures did not differ between species with relatively high, and relative low energy expenditure, this interactive effect with ambient temperature could not be explained by body temperatures being near maximum or minimum potential values alone (e.g. “ceiling” or “floor” effects; mean baseline body temperature ± s.d.: low energy expenditure = 37.5°C ± 1.70°C; high energy expenditure = 38.3°C ± 1.75°C).

**Figure 4.**
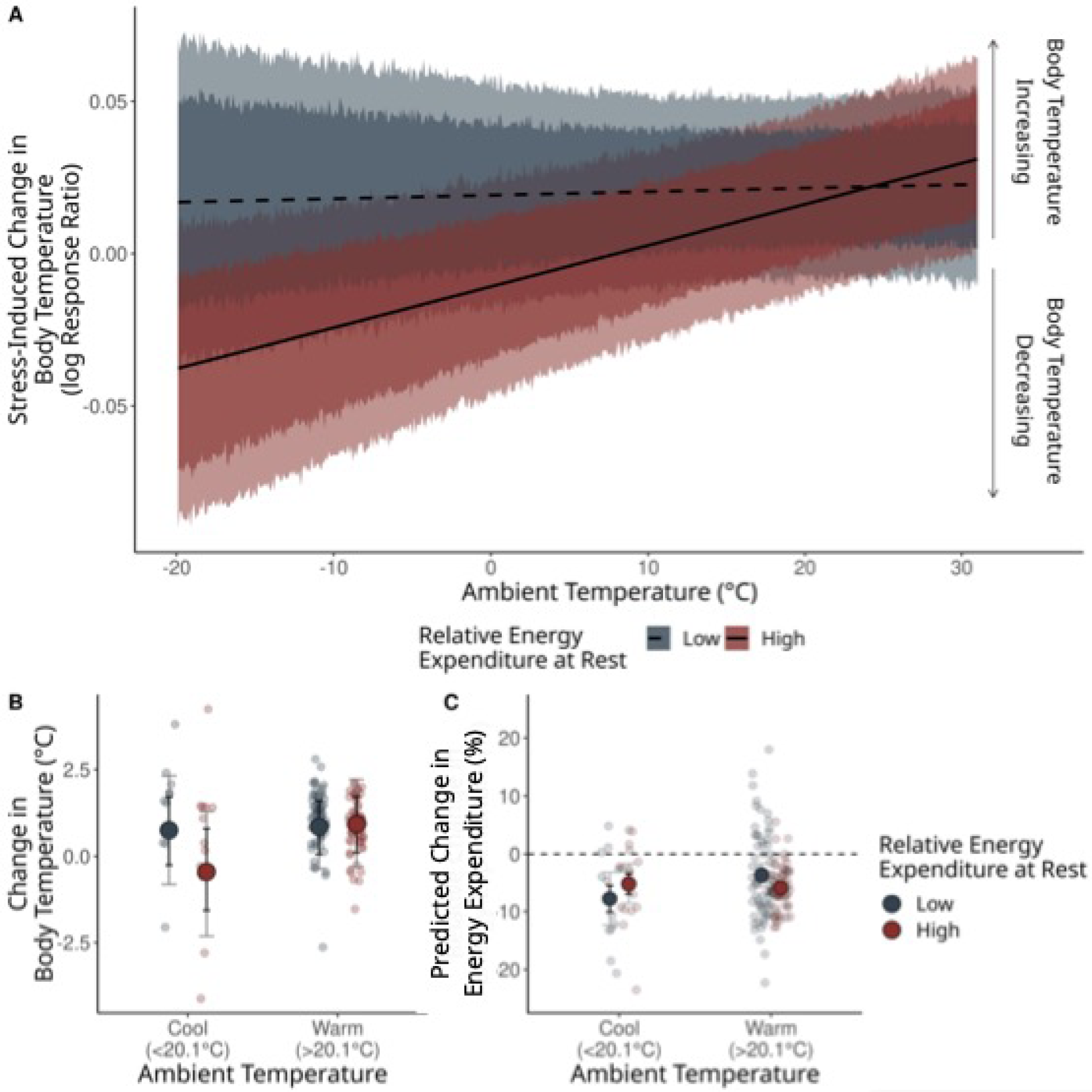
Effect of relative energy expenditure at rest on the magnitude and direction of stress-induced changed in core temperature. Relative energy expenditure at rest represents the residuals extracted from a linear mixed effects model regressing log-transformed resting metabolic rate (W) against log-transformed body mass (g), then categorized as “low” (i.e. negative residuals) or “high” (i.e. positive residuals). **A.** Negative log response ratios represent stress-induced declines in body temperature, while positive log response ratios represent stress-induced increases in body temperature. Dark ribbons represent 80% highest posterior density intervals (HPDIs) and pale ribbons represent 95% HPDIs. Trend lines and credible intervals are marginalized across all other model predictors. **B.** Large dots represent average changes in body temperature for species with high or low relative energy expenditure (red and navy respectively), and at ambient temperatures above and below the average across studies (20.1°C). Mean changes in body temperature were calculated from log-response ratios and assume a baseline body temperature of 38°C. Small dots represent raw data values and whiskers indicate 80% and 95% highest posterior density intervals (HPDIs) around means (black and grey respectively). **C.** Large dots display mean changes in resting energy expenditure attributed to stress-induced thermal responses, as predicted by biophysical models. Again, small dots represent raw data values and error bars indicate 80% and 95% highest posterior density intervals (HPDIs; black and grey respectively).

Across all studies where ambient temperatures were below our sample average (20.1°C), body temperatures of those with relatively high resting energy expenditure decreased by an average of 0.37°C in response to a stressor (Fig. 4b). Among those with relatively low resting energy expenditure, however, body temperatures increase by an average of 0.74°C across the same ambient temperature range (Fig. 4b). In both cases, biophysical models indicate that changing body temperature in these ways contributed to a 5% reduction (mean = 5.46%) in maintenance expenditure (Fig. 4c).

### Time, age class, and species identity also influence the effect of stress exposures on thermal physiology

Beyond air temperature, body mass, and relative energy expenditure, study methodology had mixed effects on the outcomes of stress-induced thermal responses. Alone, latency between baseline and stress-induced body temperature measurements clearly influenced the observed magnitude of stress-induced thermal responses (first order: β = −3.61⨯10^-2^ [80%: −7.14⨯10^-2^, −1.37⨯10^-2^; 95%: −8.53⨯10^-2^, 2.50⨯10^-3^]; second order: β = −5.04⨯10^-2^ [80%: −7.50⨯10^-2^, −1.96⨯10^-2^; 95%: −8.99⨯10^-2^, −3.34⨯10^-3^]; Figs. 2 & Supplemental Fig 56). Across all studies assessed here, and when holding all other parameters constant, body temperatures tended to increase immediately after stress exposure, and then decrease after an average of approximately 35 minutes (2100 seconds; Supplemental Fig. 56). Whether body temperature was measured using implanted telemeters or thermocouples, however, did not appear to influence how body temperatures changed in response to stressors (β = 4.80⨯10^-4^ [80%: −5.45⨯10^-3^, 8.66⨯10^-3^; 95%: −9.32⨯10^-3^, 1.24⨯10^-2^]; Fig. 2).

Although not predicted here, body temperatures were more likely to decrease after a stress exposure among study populations comprised of solely adults than among those comprised of solely juveniles (β = −6.97⨯10^-3^ [80%: −1.25⨯10^-2^, −1.37⨯10^-3^; 95%: −1.62⨯10^-3^, 1.39⨯10^-3^]; Fig. 2). How body temperature responded to stress exposures was also weakly genetically linked, with species identity and phylogeny explaining a small degree of variation in our data (group-level intercept for species: µ = 1.49⨯10^-2^ [80%: 7.55⨯10^-3^, 2.94⨯10^-2^; 95%: 4.00⨯10^-4^, 3.22⨯10^-2^]; group-level intercept for phylogeny: µ = 1.50⨯10^-3^[80%: 0, 2.23⨯10^-3^; 95%: 0, 3.25⨯10^-3^]; Fig. 2).

## Discussion

### Costs of thermoregulation dictate body temperature under stress

Changes in body temperature that accompany the stress response are often understood as byproducts of the physiological stress response and serve no function in and of themselves [41–43]. Our results do not support this understanding. Instead, we provide evidence that stress-induced changes in body temperature result from attempts to balance energetic expenditure between body temperature regulation and the stress response. More specifically, stress-induced thermal responses represent allocation trade-offs between thermoregulation and stress physiology [10,11].

Supporting our trade-off hypothesis, our findings indicate that, among terrestrial endotherms, how body temperature changes after a stress exposure may depends on how much energy a species expends on body temperature regulation (as influenced by their body mass, and thus, their surface area to volume ratio). In cool environments, for example, species predicted to have relatively high expenditure toward warming (i.e. those with low body mass, and thus, high surface area to volume ratios) tended to decrease their body temperatures, or allow them to fall, following a stress exposure, while those with relatively low expenditure toward warming, or even slight expenditure toward cooling (i.e. those with large body masses, and thus, low surface area to volume ratios) did not (Fig. 3). Under a trade-off hypothesis, we predicted this to be the case because expenditure toward the stress response should only necessitate reductions in expenditure toward thermoregulation when costs of warming or cooling are already high. By contrast, in the absence of a trade-off, expenditure toward thermoregulation should be irrelevant to the direction or magnitude of stress-induced thermal responses.

An interactive effect of ambient temperature and body mass on the outcome of stress-induced thermal responses is not new to physiological theory. In birds, Nord and Folkow [44] highlighted that hypothermic responses to stress exposure seem more likely to occur at low ambient temperatures, but specifically among species with small body size (e.g. Parids; [12,45]). Oka [46] alluded to a similar trend in small mammals (e.g. rats and domestic rabbits; [13,14,47]). For Nord and Folkow [44], this interaction between body mass and ambient temperature was interpreted as a byproduct of experimental methodology, with commonly-used restraint stressors causing greater disruptions to the retention of body heat in small birds relative to large birds (i.e. by compression of plumage). Although this interpretation may be valid in some contexts, it alone does not seem sufficient to explain our general findings. Numerous studies included in our analysis (n = 28), employed stressors that did not involve direct contact or handling of experimental animals at all (e.g. social isolation, exposure to a dominant individual, introduction to a novel environment etc). Yet despite inclusion of these studies in our analysis, we still detected a strong interactive effect of body mass and ambient temperature on the outcome of stress-induced thermal responses (Fig. 3). Thus, interpreting these responses as passive, unregulated consequences of experimental design is likely premature.

Under a trade-off hypothesis, we speculated that body mass influenced stress-induced thermal responses through its effects on heat loss and, subsequently, energetic demands for heat production (discussed above). Nevertheless, it remains possible that how body mass influences thermal responses to stressors occurs though other avenues, for example, by influencing thermal inertia. Arguably the most direct way to test our speculation would be to compare how body temperature responds to stressors at ambient temperatures below, within, and above the zone of thermoneutrality within a given species, where relative expenditure toward thermoregulation differs predictably. To the best of our knowledge, very few studies have sought to conduct this type of comparison and outcomes of those that have are equivocal [13,14,48].

### Energy availability influences whether stressors induce heteothermy

Above and beyond body mass, our results indicate that relative energy expenditure at rest influences how body temperature responds to stressors in different thermal environment. Specifically, we found that species with relatively high rates of energy expenditure at rest ‒ whose energy available to allocate to both the stress response and thermoregulation should be limited [20](but see below) ‒ tended to lower their body temperatures after stressors in the cold, but raise them in the warmth (Fig. 4). By contrast, species with relatively low rates of energy expenditure at rest ‒ whose energy available for allocation toward the stress response and thermoregulation should abound ‒ tended to raise their body temperatures after stressors regardless of the thermal environment (Fig. 4). According to our trade-off hypothesis, we predicted this combined effect of energy availability and the thermal environment on stress-induced thermal responses, since adjustments of expenditure toward body temperature regulation should predominantly occur when energy is limited [19].

For this study, we estimated that species with relatively high resting metabolism (RMR) for their given mass would have less energy available for allocation than others (as per [20]). While useful for broad inference, we recognize that this method for estimating energy availability is coarse and ignores: (1) individual-level variation, and (2) the possibility of high RMR species having other mechanisms in place to free, or acquire (but at possible risk of predation) energy for allocation in the face of a challenge. Nevertheless, studies directly or indirectly manipulating energy availability via food restriction also support a trade-off hypothesis. In adult rats, for example, food restriction reversed thermal responses to stressors, with fasted individuals decreasing their body temperatures in response to restraint, and fed individuals increasing them in equivalent, sub-thermoneutral conditions [49] (see ref [50] for range of thermoneutrality). Further, in birds, socially-mediated food restriction (i.e. by social subordination) was shown to stimulate stress-induced hypothermia in the cold and hyperthermia in the warmth (core body temperature proxied by eye region temperature; [11]; similar findings in ref [51]). These findings indicate that how body temperature changes after a stressor probably does depend on an individual’s relative energy balance, and may well represent active calibration of expenditure toward warming or cooling. Further studies employing experimental food would, however, be valuable to further clarify the role of energy balance in dictated stress-induced thermal responses.

Not surprisingly, our results reiterate a key effect of time on the magnitude and direction of stress-induced thermal responses. When controlling for all other parameters, body temperature was more likely to increase shortly after stress exposure, then decrease to baseline temperatures, or below, shortly thereafter (Fig. 2; Supplemental Fig. 56). These trends strongly align with those reported for individual species [34–36] and probably represent the timeline of physiologically responding to, and recovering from, a stressor. Perhaps more interestingly, our results also indicated that age likely influences how body temperature responds to a stressor. When averaged across species and thermal environments, we found that younger individuals were more likely to increase their body temperatures after a stressor, and to a greater extent, than older individuals (refer to negative slope of age class in Fig. 2). Given that juveniles are expected to allocate high amounts of energy toward growth [52,53], this result seems surprising. Although a similar effect of age has been reported in a handful of other studies [49,54–56], why these differences between juveniles and adults occur is not known. One possible explanation is that juveniles are comparatively less averse to wear-and-tear associated with homeostatic overload, once expenditure toward growth, thermoregulation, and the stress response are combined (i.e., a “better safe than sorry” strategy; 56,57). Alternatively, the capacity to change body temperature after a stressor may simply differ between age groups owing to age-related changes to nervous system function (reviewed in ref. [46]). In any case, understanding whether these age-related effects have meaningful implications on energy balance is yet necessary to determine whether they support or refute our trade-off hypothesis.

### Putative mechanisms

Contextualizing stress-induced changes in body temperature as allocation trade-offs (i.e. between thermoregulation and the stress response) may suggest that body temperature set-point is actively changed when a non-thermal stressor is perceived. Thermoregulatory control centers should therefore be involved in mediating these stress-induced responses. So far, studies using pharmacological agents to manipulate neural function in these control centers suggest that they probably do regulate body temperature responses to stressors [57–60], at least indirectly. Additionally, their capacity to shape stress-induced thermal responses is becoming increasingly clear [61,62]. In an effort to synthesize these findings, Angilletta et al [63] recently proposed that inhibitory pathways linking the limbic system with thermoregulatory nuclei of the hypothalamus may be activated during the stress response, thus temporarily knocking down select thermoregulatory processes. According to their model, activation of, and investment in, the acute stress response is therefore actively prioritized over thermoregulation (representing an “active” allocation trade-off). Efforts to map inhibitory pathways between limbic structures and thermoregulatory control centers are limited but generally confirmatory [64,65], thus providing some early mechanistic support for our trade-off hypothesis.

While changes to body temperature that accompany the stress response may well represent “active” trade-offs, we still cannot rule out the possibility that they merely represent passive allocation constraints. Namely, declines in body temperature in the cold and increases in body temperature in the warmth might simply indicate a failing to allocate energy toward responding to the thermal environment experienced during a stress exposure. In this context, thermal responses to stressors may not constitute functional responses to balance energetic expenditure between thermoregulation and the stress response, but rather, may indicate an inability to regulate body temperature at all under distress (see, for example, ref. [56]). Whether these stress-induced changes in body temperature do indeed represent regulated trade-offs or passive constraints is an important question that will directly influence whether these responses may respond to selection or are simply fixed consequences of how physiological responses to stressors shape body temperature.

### Ectotherms and non-terrestrial endotherms

Descriptions of how body temperatures change after stressors among ectotherms and non-terrestrial endotherms are sparse. We were therefore unable to formally evaluate whether evidence for an allocation trade-off between thermoregulation and the stress response extends to these species’ groupings. However, indirect evidence for such a trade-off in ectotherms is widely supportive. Some species of lizards, grasshoppers, and even larval newts, for example, reportedly abandon basking and/or heat-seeking behavior in the presence of predator cues (reviewed in ref. [63]). In the Yarrow’s spiny lizard (*Sceloporus jarrovii*), this change in behavior results in body temperature both declining toward ambient and away from preference [66]. While not strictly evidencing an energy allocation trade-off, these reports do highlight that responding to stressors can, and often does, take precedence over body temperature regulation in ectotherms with clear consequences for the latter. Among non-terrestrial endotherms, thermoregulation may also be relaxed, or even forgone, in the presence of stressors. In southern bluefin tuna (*Thunnus maccoyii*), for example, body temperatures can rise dramatically in response to suspected predators [67], which, in severe cases, is thought to contribute to physiological damage ([68]; a phenomenon known as “burnt meat”). We therefore speculate that our support for a trade-off between coping was a stressor and regulating body temperature probably extends beyond the species grouping analyzed here.

## Conclusions

Our results provide strong evidence that changes in body temperature accompanying the stress response represent allocation trade-offs between the stress-response and thermoregulation. Across species, we highlight that by allowing body temperature to drift toward ambient, or in the direction of accumulating metabolic heat, these responses contribute to reductions in the energetic costs of thermoregulation, particularly when such are already high, or when energy available for allocation to both the stress response and body temperature regulation are low. These findings, therefore, highlight a new model for understanding both how and “why” stress-induced changes in body temperature might occur (outlined in Fig. 1). In doing so, they provide novel, cross-species evidence that the acute stress response may directly influence body temperature regulation in endotherms. Given the clear implication of these findings for a warming world, further experimental studies testing our trade-off hypothesis within species should be a priority in coming years.

## Acknowledgements

We thank Ms. Elin Persson and Maria Correia, Drs. Arne Hegemann, Caroline Isaksson, Andreas Nord, and Jim Schafer for their perspectives on our study. We are particularly grateful for analytical suggestions provided by Dr. Mari Fjelldal. Funding for this research was provided by a Natural Sciences and Engineering Research Council grant to GB. JT and GB conceived the study, MH and JT collected the data, JT conducted statistical analyses and wrote the first draft of the manuscript with GB contributing to manuscript revisions. The authors have no conflicts of interest to declare. All data needed to evaluate this study’s findings are given in the Supplementary Materials.

## Notes

### Competing Interest Statement

The authors have declared no competing interest.

### Summary of Updates

(1) Brevity. In this revised draft, we have refined our text to more directly circle around our key research questions and findings. Doing so has required us to shift some previously main figures to supplemental information. (2) Clarity regarding predictions of our trade-off hypothesis. These predictions, particularly with regard to specific environmental contexts, are now more explicit and accompanied by a new prediction figure (Fig. 1b). (3) Integration with broader literature. In our revised introduction, we now contextualise our findings with the broader literature around endotherm thermoregulation in challenging environments.

