## Supplemental Information for "Endotherms trade body temperature regulation for the stress response"

This document displays and describes all data compilation, data organisation, and data analysis that is required to reproduce the results reported in "Endotherms trade body temperature regulation for the stress response". All code below has been executed in R version 4.2.1 (R Core Team, 2022) on a x86_64-pc-linux-gnu platform. Annotations (preceded by the pound symbol, "#") are provided throughout the document to explain the purpose of core operations, or to guide the interpretation of data tables and plots.

All data files imported in the code below will be available at a publicly-available GitHub™ repository upon publication. We encourage readers to download these data and follow our document by executing the code in their own R terminal. Before doing so, however, readers should acknowledge that our latent-variable models employ Hamiltonian Monte Carlo (HMC) sampling with relatively small step sizes, high numbers of iterations, and often weakly informative priors. These models can therefore be both computationally demanding and time consuming. Those with limited computational power, then, may wish to reduce the number of Markov Chain Monte Carlo (MCMC) chains or iterations in our models before executing them. Doing so should reduce computational demands, but, as a forewarning, it may also hinder chain convergence and/or shift posterior densities. We therefore advise caution when interpreting the outcomes of such adjusted models.

Those familiar with the R package used to execute Bayesian models in Stan ('brms'; Bürkner, 2018) will see that we have chosen to save the outputs of our models in .Rds files. These .Rds files have been provided with the manuscript submission, and readers may bypass model re-execution by downloading and calling them in the "file" argument of a given brms function. If the objective of the reader is to replicate our results, however, we recommend re-constructing and running our models on one's own.

Finally, below, we provide the version numbers of each package used in our analysis. To ensure that the outcomes of all code processes below are similar to those shown, we advise that readers specify the provided version numbers during package installation.

---

#### 1 | Importing and compiling data

We begin by importing all necessary packages and defining additional, supportive functions.

```
# Note that packages are loaded in bulk using
# 'easypackages'.
library("devtools")
# install_github('jacintak/colRoz')
# install_github('karthik/wesanderson')

library("easypackages")
library("tidyverse")
libraries("ape", "bayesplot", "bayestestR", "brms", "coda", "colRoz",
    "cmdstanr", "DiagrammeRsvg", "elementalist", "gghalves",
    "ggplotify", "ggtext", "ggtree", "grid", "gridExtra", "kableExtra",
    "NicheMapR", "performance", "PRISMAstatement", "rstan", "rstanarm",
    "rsvg", "showtext", "sn", "tidybayes", "wesanderson")
```

```r
# Viewing package numbers.

pV <- function(x) {
    y <- packageVersion(x)
    y <- as.character(y)
    return(y)
}

caption = "List of R package versions used in the subsequent analyses."

sapply(c("ape", "bayesplot", "bayestestR", "brms", "coda", "colRoz",
    "cmdstanr", "easypackages", "gghalves", "ggplotify", "ggtext",
    "ggtree", "grid", "gridExtra", "kableExtra", "performance",
    "PRISMAstatement", "rstan", "rstanarm", "showtext", "sn",
    "tidybayes", "tidyverse", "wesanderson"), pV, simplify = FALSE) %>%
    enframe(., name = "Package", value = "Version") %>%
    as.data.frame(.) %>%
    kbl(., longtable = T, booktabs = T, caption = caption) %>%
    kable_styling(latex_options = "striped")
```

Table 1: List of R package versions used in the subsequent analyses.

| Package | Version |
|---|---|
| ape | 5.7.1 |
| bayesplot | 1.10.0 |
| bayestestR | 0.13.1 |
| brms | 2.19.0 |
| coda | 0.19.4 |
| colRoz | 0.2.2 |
| cmdstanr | 0.5.3 |
| easypackages | 0.1.0 |
| gghalves | 0.1.4 |
| ggplotify | 0.1.0 |
| ggtext | 0.1.2 |
| ggtree | 3.6.2 |
| grid | 4.2.3 |
| gridExtra | 2.3 |
| kableExtra | 1.3.4 |
| performance | 0.10.3 |
| PRISMAstatement | 1.1.1 |
| rstan | 2.26.22 |
| rstanarm | 2.21.4 |
| showtext | 0.9.6 |
| sn | 2.1.1 |
| tidybayes | 3.0.4 |
| tidyverse | 2.0.0 |
| wesanderson | 0.3.6.9000 |

```r
# Some plots below use the freely-available Google Font
# 'Noto Sans'. The below line imports that font.

font_add_google(name = "Noto Sans", family = "Noto Sans")

# Here, the colour scheme for diagnostic plots produced by
# the 'bayesplot' package is set.

color_scheme_set("pink")
nice_pink = "#b97c9b"

# The below function, reported by 'ellbur' at
# https://strugglingthroughproblems.wordpress.com/author/ellbur/page/3/,
# allows users to assign multiple objects to different
# variable names at once.
```

```
{
    "%=%" <- function(l, r, ...) UseMethod("%=%")

    "%=%.lbunch" <- function(l, r, ..., List = NA) {
        Envir <- as.environment(-1)

        if (!is.na(List)) {
            l <- List[[1]]
            r <- List[[2]]
        }

        if (length(r) > length(l)) {
            warning("RHS has more args than LHS. Only first",
                length(l), "used.")
        }

        if (length(l) > length(r)) {
            warning("LHS has more args than RHS. RHS will be repeated.")
            r <- extendToMatch(r, l)
        }

        for (II in 1:length(l)) {
            do.call("<-", list(l[[II]], r[[II]]), envir = Envir)
        }
    }

    extendToMatch <- function(source, destin) {
        s <- length(source)
        d <- length(destin)

        if (d == 1 && s > 1 && !is.null(as.numeric(destin))) {
            d <- destin
        }

        dif <- d - s
        if (dif > 0) {
            source <- rep(source, ceiling(d/s))[1:d]
        }
        return(source)
    }

    g <- function(...) {
        List <- as.list(substitute(list(...)))[-1L]
        class(List) <- "lbunch"
        return(List)
    }
}

# Adding functions that estimate 95% and 50% credible
# intervals from quantiles

{
    LCL_Fun <- function(x) {
        quantile(x, 0.025, type = 8)
    }
    UCL_Fun <- function(x) {
        quantile(x, 0.975, type = 8)
    }
    LCL_Fun_50 <- function(x) {
        quantile(x, 0.25, type = 8)
    }
    UCL_Fun_50 <- function(x) {
        quantile(x, 0.75, type = 8)
    }
}

# Adding a condensed function to call median Bayesian or
```

```
# Loo R2 values as defined by Gelman et al (2018; main text
# and appendix).

bR2 <- function(x, resp = NA, digits = 2, loo = FALSE) {
    require(brms)
    if (!is.na(resp)) {
        if (loo == TRUE) {
            return(round(median(loo_R2(x, resp = resp)), digits))
        } else {
            return(round(median(bayes_R2(x, resp = resp)), digits))
        }
    } else {
        if (loo == TRUE) {
            return(round(loo_R2(bayes_R2(x)), digits))
        } else {
            return(round(median(bayes_R2(x)), digits))
        }
    }
}

# A function to calculate the mode of a vector.

md <- function(x) {
    all_values <- unique(x)
    all_values[which.max(tabulate(match(x, all_values)))]
}

# A function to simplify the output of bayestestR's hdi
# function.

simple_hdi <- function(x, rnd = 3, cis = c(50, 95), sci_note = FALSE) {
    if (class(x)[1] != "brmsfit") {
        return("x must be a brmsfit object.")
    }
    if (length(cis) != 2) {
        return("cis must be a vector of integers with length 2")
    }

    HDI_low <- bayestestR::hdi(x, effects = "all", ci = min(cis)/100)
    HDI_high <- bayestestR::hdi(x, effects = "all", ci = max(cis)/100)

    if (sci_note == FALSE) {
        Results <- data.frame(Parameter = HDI_low$Parameter,
            `1` = round(HDI_low$CI_low, rnd), `2` = round(HDI_low$CI_high,
                rnd), `3` = round(HDI_high$CI_low, rnd), `4` = round(HDI_high$CI_high,
                rnd))
        colnames(Results)[c(2:5)] <- c(paste0("Low_HDI_", min(cis)),
            paste0("High_HDI_", min(cis)), paste0("Low_HDI_",
                max(cis)), paste0("High_HDI_", max(cis)))
    } else if (sci_note == TRUE) {
        Results <- data.frame(Parameter = HDI_low$Parameter,
            `1` = format(round(HDI_low$CI_low, rnd), scientific = TRUE),
            `2` = format(round(HDI_low$CI_high, rnd), scientific = TRUE),
            `3` = format(round(HDI_high$CI_low, rnd), scientific = TRUE),
            `4` = format(round(HDI_high$CI_high, rnd), scientific = TRUE))
        colnames(Results)[c(2:5)] <- c(paste0("Low_HDI_", min(cis)),
            paste0("High_HDI_", min(cis)), paste0("Low_HDI_",
                max(cis)), paste0("High_HDI_", max(cis)))
    }

    return(Results)
}

# Four functions to print clean MCMC density, MCMC pairs,
# MCMC intervals, and MCMC autocorrelations plots that are
# rendered by bayesplot and rstan.
```

```
clean_dens <- function(x, prs = NA, names = NA) {
    require("bayesplot")

    if (class(x)[1] != "brmsfit") {
        return("x must be a brmsfit object.")
    }

    if (is.na(prs[1])) {
        return(mcmc_dens(x))
    }

    if (!is.na(prs[1]) & is.na(names[1])) {
        return(mcmc_dens(x, pars = prs))
    }

    if (length(prs) != length(names)) {
        return("Length of pars and names must be equal.")
    }

    Dens_Panel <- mcmc_dens(x, pars = prs)
    Dens_Panel$data$Parameter <- as.character(Dens_Panel$data$Parameter)

    for (i in 1:length(prs)) {
        Dens_Panel$data$Parameter[c(which(Dens_Panel$data$Parameter ==
            prs[i]))] <- names[i]
    }

    if (length(prs) > 1) {
        Dens_Panel$facet$params$ncol = 2
    }

    return(Dens_Panel)
}

clean_pairs <- function(x, prs = NA, names = NA, np = NA) {
    require("bayesplot")
    require("ggplotify")
    require("grid")

    if (class(x)[1] != "brmsfit") {
        return("x must be a brmsfit object.")
    }

    if (is.na(prs[1])) {
        if (is.na(np)[1] == TRUE) {
            return(mcmc_pairs(x))
        } else {
            return(mcmc_pairs(x, np = np))
        }
    }

    if (!is.na(prs[1]) & is.na(names[1])) {
        if (is.na(np)[1] == TRUE) {
            return(mcmc_pairs(x, pars = prs))
        } else {
            return(mcmc_pairs(x, pars = prs, np = np))
        }
    }

    if (length(prs) != length(names)) {
        return("Length of pars and names must be equal.")
    }

    if (is.na(np)[1] == TRUE) {
        Pairs_Panel <- mcmc_pairs(x, pars = prs, off_diag_args = list(size = 1.5))
    } else {
        Pairs_Panel <- mcmc_pairs(x, pars = prs, off_diag_args = list(size = 1.5),
```

```
            np = np)
    }

    Diagonal <- matrix(nrow = length(prs), ncol = length(prs))
    diag(Diagonal) <- 1
    To_Grab <- c(which(Diagonal == 1))

    for (i in 1:length(To_Grab)) {
        Pairs_Panel[["grobs"]][[To_Grab[i]]][["grobs"]][[15]] <- nullGrob()
        New_Grob <- ggplotGrob(as.ggplot(Pairs_Panel[["grobs"]][[To_Grab[i]]]) +
            labs(subtitle = names[i]) + theme(plot.title = element_text(hjust = 0.5)))
        Pairs_Panel[["grobs"]][[To_Grab[i]]] <- New_Grob
    }
    return(Pairs_Panel)
}

clean_int <- function(x, prs = NA, names = NA) {
    require("bayesplot")

    if (class(x)[1] != "brmsfit") {
        return("x must be a brmsfit object.")
    }

    if (is.na(prs[1])) {
        return(mcmc_intervals(x))
    }

    if (!is.na(prs[1]) & is.na(names[1])) {
        return(mcmc_intervals(x, pars = prs))
    }

    if (length(prs) != length(names)) {
        return("Length of pars and names must be equal.")
    }

    Int_plot <- mcmc_intervals(x, pars = prs)

    for (i in 1:length(prs)) {
        Int_plot <- Int_plot + scale_y_discrete(labels = c(names))
    }

    return(Int_plot)
}

clean_ac <- function(x, prs = NA, names = NA) {
    require("rstan")

    if (class(x)[1] != "brmsfit") {
        return("x must be a brmsfit object.")
    }

    if (is.na(prs[1])) {
        return(stan_ac(x$fit))
    }

    if (!is.na(prs[1]) & is.na(names[1])) {
        return(stan_ac(x$fit, pars = prs))
    }

    if (length(prs) != length(names)) {
        return("Length of pars and names must be equal.")
    }

    Base_plot <- stan_ac(x$fit, pars = prs, fill = nice_pink)
    Base_plot$data$parameters <- as.character(Base_plot$data$parameters)

    for (i in 1:length(prs)) {
```

```
        Base_plot$data$parameters[c(which(Base_plot$data$parameters ==
            prs[i]))] <- names[i]
    }
    Base_plot$data$parameters <- as.factor(Base_plot$data$parameters)

    return(Base_plot)
}
```

Next, we begin importing, compiling, and summarising data pertaining to stress-induced changes in body temperature across species. Collection of these data is described in the main text of our manuscript. In this study, we were interested in testing whether changes in body temperature that accompany a stress exposure represent a trade-off between *core* body temperature regulation and the stress response. For this reason, thermal responses to stress exposure that were measured at the body surface and not the body core were removed from our data-set.

```
Data <- read.csv("/Users/joshuatabh/Documents/researchProjects/trent/sihMetaregression/data/final/stressData.csv")

# Reviewing species names and correcting where necessary.

caption = "List of species incuded in dataset by species names."
Data %>%
    group_by(Species.Name) %>%
    summarise(Count = n()) %>%
    as.data.frame(.) %>%
    rename(`Latin Name` = Species.Name) %>%
    kbl(., longtable = T, booktabs = T, caption = caption) %>%
    kable_styling(latex_options = "striped")
```

Table 2: List of species incuded in dataset by species names.

| Latin Name | Count |
|---|---|
| Aepyceros melampus | 3 |
| Amazona aestiva | 1 |
| Amazona ventralis | 2 |
| Anas platyrhynchos | 1 |
| Bombina bombina | 1 |
| Bos taurus | 11 |
| Bufo marinus | 1 |
| Callithrix penicillata | 6 |
| Callopistes maculatus | 1 |
| Canis lupus familiaris | 8 |
| Capra aegagrus hircus | 2 |
| Capreolus capreolus | 2 |
| Cavia porcellus | 6 |
| Ceratotherium simum | 1 |
| Columba livia | 10 |
| Cyanistes caeruleus | 1 |
| Danio rerio | 2 |
| Dipodomys merriami | 4 |
| Dromaius novaehollandiae | 2 |
| Equus caballus | 7 |
| Equus ferus caballus | 1 |
| Felis catus | 4 |
| Gallus domesticus | 1 |
| Gallus gallus domesticus | 24 |
| Glyptemys insculpta | 1 |
| Hirundo rustica | 2 |
| Lagopus muta hyperborea | 16 |
| Lasionycteris noctivagans | 3 |
| Macaca mulatta | 1 |
| Melopsittacus undulatus | 3 |

| | |
|---|---|
| Mesocricetus auratus | 2 |
| Mus musculus | 84 |
| Mustela vison | 4 |
| Oryctolagus cuniculus | 3 |
| Oryctolagus cuniculus domesticus | 9 |
| Ovis aries | 9 |
| Ovis canadensis | 1 |
| Panthera tigris | 1 |
| Parus major | 5 |
| Poecile atricapillus | 2 |
| Rangifer tarandus platyrhynchus | 2 |
| Rattus norvegicus | 49 |
| Rattus norvegicus domestica | 99 |
| Saxicola rubetra | 1 |
| Somateria mollissima | 2 |
| Spermophilus beecheyi | 8 |
| Strix varia | 1 |
| Struthio camelus | 1 |
| Sus scrofa domesticus | 3 |
| Sylvia borin | 1 |
| Tamias striatus | 1 |
| Tupaia belangeri | 3 |
| Vulpes vulpes | 3 |

```
# Subspecies names will be used temporarily here.

Data <- Data %>%
    mutate(Species.Name = ifelse(Species.Name == "Equus ferus caballus",
        "Equus caballus", ifelse(Species.Name == "Gallus domesticus",
            "Gallus gallus domesticus", ifelse(Species.Name ==
                "Rattus norvegicus " | Species.Name == "Rattus norvegicus",
                "Rattus norvegicus domestica", ifelse(Species.Name ==
                  "Oryctolagus cuniculus", "Oryctolagus cuniculus domesticus",
                  Species.Name)))))

# Reviewing species common names to double-check.

caption = "List of species incuded in dataset by common names."
Data %>%
    group_by(Common.Name) %>%
    summarise(Count = n()) %>%
    as.data.frame(.) %>%
    rename(`Common Name` = Common.Name) %>%
    kbl(., longtable = T, booktabs = T, caption = caption) %>%
    kable_styling(latex_options = "striped")
```

Table 3: List of species incuded in dataset by common names.

| Common Name | Count |
|---|---|
| American Mink | 4 |
| Ames Dwarf Mouse | 1 |
| Ames Mouse | 1 |
| Angus Cattle | 3 |
| Barn Swallow | 2 |
| Barred Owls | 1 |
| Bengal Tiger | 1 |
| Bighorn Sheep | 1 |
| Black-Tufted Marmosets | 6 |
| Black-capped Chickadee | 2 |
| Blue Tit | 1 |
| Blue-Fronted Parrot | 1 |
| Brazilian Mangalarga Machador Horse | 1 |
| Brown Swiss Cows | 2 |

| | |
|---|---|
| Budgerigars | 3 |
| California Ground Squirrel | 8 |
| Cane Toad | 1 |
| Common Eider | 2 |
| Common Ostrich | 1 |
| Domestic Albino Rabbit | 8 |
| Domestic Cat | 4 |
| Domestic Chicken | 25 |
| Domestic Dog | 8 |
| Domestic Goat | 2 |
| Domestic Horse | 7 |
| Domestic Pig | 3 |
| Domestic Rabbit | 4 |
| Domestic Sheep | 6 |
| Eastern Chipmunk | 1 |
| Emu | 2 |
| Fire-bellied Toad | 1 |
| Fischer Rat | 17 |
| Garden Warblers | 1 |
| Golden Hamster | 2 |
| Great Tit | 5 |
| Guinea Pig | 6 |
| Hispaniolan Amazon Parrot | 2 |
| Holstein Cattle | 3 |
| Holstein Friesian Cow | 3 |
| House Mouse | 82 |
| Impala | 3 |
| Kangaroo Rat | 4 |
| Lizard | 1 |
| Long-Evans Rat | 1 |
| Pekin Duck | 1 |
| Rat | 3 |
| Rat (unknown strain) | 2 |
| Rhesus Macaque | 1 |
| Rock Pigeon | 10 |
| Roe Deer | 2 |
| Romney Sheep | 3 |
| Silver Fox | 3 |
| Silver-haired Bat | 3 |
| Sprague Dawley Rat | 75 |
| Svalbard Reindeer | 2 |
| Svalbard Rock Ptarmigan | 16 |
| Treeshrew | 3 |
| Whinchats | 1 |
| White Rhino | 1 |
| Wild Type Groningen Rat | 4 |
| Wistar Rat | 46 |
| Wood Turtle | 1 |
| Zebrafish | 2 |

```
# Unknown or unorthodox rat strains are renamed
# generically.

Data <- Data %>%
    mutate(Common.Name = ifelse(Common.Name == "Wild Type Groningen Rat",
        "Rat", ifelse(Common.Name == "Rat (unknown strain)",
            "Rat", Common.Name)))

# Reviewing corrections.

caption = "List of species incuded in dataset by common names, after name corrections."
Data %>%
```

```
    group_by(Species.Name, Common.Name) %>%
    summarise(Count = n()) %>%
    as.data.frame(.) %>%
    rename(`Latin Name` = Species.Name, `Common Name` = Common.Name) %>%
    kbl(., longtable = T, booktabs = T, caption = caption) %>%
    kable_styling(latex_options = "striped")
```

Table 4: List of species incuded in dataset by common names, after name corrections.

| Latin Name | Common Name | Count |
|---|---|---|
| Aepyceros melampus | Impala | 3 |
| Amazona aestiva | Blue-Fronted Parrot | 1 |
| Amazona ventralis | Hispaniolan Amazon Parrot | 2 |
| Anas platyrhynchos | Pekin Duck | 1 |
| Bombina bombina | Fire-bellied Toad | 1 |
| Bos taurus | Angus Cattle | 3 |
| Bos taurus | Brown Swiss Cows | 2 |
| Bos taurus | Holstein Cattle | 3 |
| Bos taurus | Holstein Friesian Cow | 3 |
| Bufo marinus | Cane Toad | 1 |
| Callithrix penicillata | Black-Tufted Marmosets | 6 |
| Callopistes maculatus | Lizard | 1 |
| Canis lupus familiaris | Domestic Dog | 8 |
| Capra aegagrus hircus | Domestic Goat | 2 |
| Capreolus capreolus | Roe Deer | 2 |
| Cavia porcellus | Guinea Pig | 6 |
| Ceratotherium simum | White Rhino | 1 |
| Columba livia | Rock Pigeon | 10 |
| Cyanistes caeruleus | Blue Tit | 1 |
| Danio rerio | Zebrafish | 2 |
| Dipodomys merriami | Kangaroo Rat | 4 |
| Dromaius novaehollandiae | Emu | 2 |
| Equus caballus | Brazilian Mangalarga Machador Horse | 1 |
| Equus caballus | Domestic Horse | 7 |
| Felis catus | Domestic Cat | 4 |
| Gallus gallus domesticus | Domestic Chicken | 25 |
| Glyptemys insculpta | Wood Turtle | 1 |
| Hirundo rustica | Barn Swallow | 2 |
| Lagopus muta hyperborea | Svalbard Rock Ptarmigan | 16 |
| Lasionycteris noctivagans | Silver-haired Bat | 3 |
| Macaca mulatta | Rhesus Macaque | 1 |
| Melopsittacus undulatus | Budgerigars | 3 |
| Mesocricetus auratus | Golden Hamster | 2 |
| Mus musculus | Ames Dwarf Mouse | 1 |
| Mus musculus | Ames Mouse | 1 |
| Mus musculus | House Mouse | 82 |
| Mustela vison | American Mink | 4 |
| Oryctolagus cuniculus domesticus | Domestic Albino Rabbit | 8 |
| Oryctolagus cuniculus domesticus | Domestic Rabbit | 4 |
| Ovis aries | Domestic Sheep | 6 |
| Ovis aries | Romney Sheep | 3 |
| Ovis canadensis | Bighorn Sheep | 1 |
| Panthera tigris | Bengal Tiger | 1 |
| Parus major | Great Tit | 5 |
| Poecile atricapillus | Black-capped Chickadee | 2 |
| Rangifer tarandus platyrhynchus | Svalbard Reindeer | 2 |
| Rattus norvegicus domestica | Fischer Rat | 17 |
| Rattus norvegicus domestica | Long-Evans Rat | 1 |
| Rattus norvegicus domestica | Rat | 9 |
| Rattus norvegicus domestica | Sprague Dawley Rat | 75 |
| Rattus norvegicus domestica | Wistar Rat | 46 |

| | | |
|---|---|---|
| Saxicola rubetra | Whinchats | 1 |
| Somateria mollissima | Common Eider | 2 |
| Spermophilus beecheyi | California Ground Squirrel | 8 |
| Strix varia | Barred Owls | 1 |
| Struthio camelus | Common Ostrich | 1 |
| Sus scrofa domesticus | Domestic Pig | 3 |
| Sylvia borin | Garden Warblers | 1 |
| Tamias striatus | Eastern Chipmunk | 1 |
| Tupaia belangeri | Treeshrew | 3 |
| Vulpes vulpes | Silver Fox | 3 |

```
# Good. Checking and correcting orders.

caption = "List of orders represented in dataset."
Data %>%
    group_by(Order) %>%
    summarise(Count = n()) %>%
    kbl(., longtable = T, booktabs = T, caption = caption) %>%
    kable_styling(latex_options = "striped")
```

Table 5: List of orders represented in dataset.

| Order | Count |
|---|---|
| Anseriformes | 3 |
| Anura | 2 |
| Artiodactyla | 25 |
| Carnivora | 20 |
| Casuariiformes | 2 |
| Chiroptera | 3 |
| Columbidae | 10 |
| Cypriniformes | 2 |
| Galliformes | 41 |
| Lagomorpha | 11 |
| Passeriformes | 9 |
| Passerine | 3 |
| Perissodactyla | 8 |
| Primates | 7 |
| Psittaciformes | 6 |
| Rodentia | 254 |
| Scandentia | 3 |
| Squamata | 1 |
| Strigiformes | 1 |
| Struthioniformes | 1 |
| Testudines | 1 |
| Therapsid | 9 |

```
Data <- Data %>%
    mutate(Order = ifelse(Order == "Passerine", "Passeriformes",
        ifelse(Order == "Even-toed ungulates", "Artiodactyla",
            ifelse(Order == "Odd-toed ungulates", "Perissodactyla",
                ifelse(Order == "Columbidae", "Columbiformes",
                  Order)))))

# Filtering out data that pertain to responses at the
# surface of the body, and not at the body core.

Data <- Data %>%
    filter(Surface.Core == "Core")

# Checking mean publication year and sem in raw data.
```

```
Data %>%
    group_by(StudyID) %>%
    summarise(Year = mean(Year.Study)) %>%
    ungroup() %>%
    summarise(`Mean Publication Year` = round(mean(Year), digits = 1),
        SEM = sd(Year)/sqrt(n()))
```

```
## # A tibble: 1 x 2
##   `Mean Publication Year`   SEM
##                     <dbl> <dbl>
## 1                   2003. 0.879
```

Because body mass is likely to affect the magnitude of stress-induced changes in body temperature (owing to an influence on thermal inertia), we extracted estimates of average body mass (in grams, $\pm$ s.d.) from study populations addressed in our meta-regression. Often, however, a range of body masses was reported rather than a population mean. In these cases, we assumed the population mean to be the hypothetical median of the mass range, and the standard deviation around the mean to be the reported range divided by 4 (see Brase and Brase, 2017). Where standard errors were reported in place of standard deviations, we back-converted them to standard deviation values using the reported sample sizes. Finally, if neither mean body mass, nor a body mass range were reported for a given sample population, we assumed mean body mass to be the average mass for the species (and/or strain) in question, as reported from other primary sources. Standard errors around these means were then derived from the mean cofficient of variation (i.e. standard deviation divided by the mean) for our sample population. If the study population lacking a measure of mass was comprised of juvenile (defined as pre-reproductive age for a given species), we identified and assumed their mass to be the average for their species, strain, and weekly age, as reported from other sources.

```
# Beginning by ensuring that no characters beyond hyphens
# are included in the mass range column.

Data$Mass <- Data$Mean.Mass.g
Data$Mass.Range.g[grepl(pattern = "g", Data$Mass.Range.g)]
```

```
## [1] "8-12 g"
```

```
Data$Mass.Range.g <- mapply(gsub, pattern = "g", replacement = "",
    Data$Mass.Range.g)

Data$Mass.Range.g <- mapply(gsub, pattern = " ", replacement = "",
    Data$Mass.Range.g)

Data$Mass <- as.character(Data$Mass)

# Grabbing mass ranges and calculating hypothetical median.

mean_est <- function(x) {
    if (!is.numeric(x[1]) & !is.numeric(x[2])) {
        x <- as.numeric(x)
    }
    return(mean(x))
}

To_Split <- grepl(pattern = "[[:digit:]]-", Data$Mass.Range.g) &
    is.na(Data$Mean.Mass.g)

Data$Mass[To_Split] <- as.character(unlist(lapply(str_split(Data$Mass.Range.g[To_Split],
    "-"), mean_est)))

Data$Mass <- as.numeric(Data$Mass)
Data$Mass_SD <- as.character(Data$Mass.sd)

# Estimating hypothetical standard deviation of mass ranges

sd_est <- function(x) {
```

```
    if (!is.numeric(x[1]) & !is.numeric(x[2])) {
        x <- as.numeric(x)
    }
    return((x[2] - x[1])/4)
}

To_Split <- grepl(pattern = "[[:digit:]]-", Data$Mass.Range.g) &
    is.na(Data$Mass.sd)

Data$Mass_SD[To_Split] <- as.character(unlist(lapply(str_split(Data$Mass.Range.g[To_Split],
    "-"), sd_est)))

Data$Mass_SD <- as.numeric(Data$Mass_SD)

# Checking which observations lack a mass measurements

caption = "List of observations where body mass measurements are lacking."

Data %>%
    filter(is.na(Mass)) %>%
    select(c(ObservationID, StudyID, Common.Name, Species.Name)) %>%
    arrange(Common.Name, Species.Name, StudyID) %>%
    as.data.frame(.) %>%
    rename(`Common Name` = Common.Name, `Latin Name` = Species.Name,
        `Study ID` = StudyID, `Observation ID` = ObservationID) %>%
    kbl(., longtable = T, booktabs = T, caption = caption) %>%
    kable_styling(latex_options = "striped")
```

Table 6: List of observations where body mass measurements are lacking.

| Observation ID | Study ID | Common Name | Latin Name |
|---|---|---|---|
| BV | Q | American Mink | Mustela vison |
| BW | Q | American Mink | Mustela vison |
| BX | Q | American Mink | Mustela vison |
| BY | Q | American Mink | Mustela vison |
| HX | W2 | Barn Swallow | Hirundo rustica |
| KB | V3 | Bengal Tiger | Panthera tigris |
| NZ | W5 | Bighorn Sheep | Ovis canadensis |
| MX | C5 | Brazilian Mangalarga Machador Horse | Equus caballus |
| AZ1 | M | California Ground Squirrel | Spermophilus beecheyi |
| AZ2 | M | California Ground Squirrel | Spermophilus beecheyi |
| AJ | E | Common Eider | Somateria mollissima |
| HA | R2 | Domestic Cat | Felis catus |
| HB | R2 | Domestic Cat | Felis catus |
| HC | R2 | Domestic Cat | Felis catus |
| HD | R2 | Domestic Cat | Felis catus |
| MG | R4 | Domestic Chicken | Gallus gallus domesticus |
| OT | C6 | Domestic Dog | Canis lupus familiaris |
| OU | C6 | Domestic Dog | Canis lupus familiaris |
| NP | P5 | Domestic Dog | Canis lupus familiaris |
| NH | K5 | Domestic Horse | Equus caballus |
| GA | D2 | Domestic Pig | Sus scrofa domesticus |
| HV | W2 | Garden Warblers | Sylvia borin |
| NF | J5 | Guinea Pig | Cavia porcellus |
| NG | J5 | Guinea Pig | Cavia porcellus |
| NK | M5 | Holstein Cattle | Bos taurus |
| KQ | C4 | House Mouse | Mus musculus |
| LG | G4 | House Mouse | Mus musculus |
| LH | G4 | House Mouse | Mus musculus |
| LI | G4 | House Mouse | Mus musculus |
| GG | H2 | House Mouse | Mus musculus |
| JB | J3 | House Mouse | Mus musculus |
| JC | J3 | House Mouse | Mus musculus |
| GO | L2 | House Mouse | Mus musculus |

| | | | |
|---|---|---|---|
| GP | L2 | House Mouse | Mus musculus |
| GP1 | L2 | House Mouse | Mus musculus |
| GP2 | L2 | House Mouse | Mus musculus |
| GP3 | L2 | House Mouse | Mus musculus |
| GP4 | L2 | House Mouse | Mus musculus |
| NN | O5 | House Mouse | Mus musculus |
| NO | O5 | House Mouse | Mus musculus |
| BE | O | Impala | Aepyceros melampus |
| BE1 | O | Impala | Aepyceros melampus |
| BE2 | O | Impala | Aepyceros melampus |
| IT | G3 | Rat | Rattus norvegicus domestica |
| IU | G3 | Rat | Rattus norvegicus domestica |
| IV | G3 | Rat | Rattus norvegicus domestica |
| IW | G3 | Rat | Rattus norvegicus domestica |
| ED | O1 | Rat | Rattus norvegicus domestica |
| ME | Q4 | Roe Deer | Capreolus capreolus |
| MF | Q4 | Roe Deer | Capreolus capreolus |
| EI | Q1 | Silver Fox | Vulpes vulpes |
| EJ | Q1 | Silver Fox | Vulpes vulpes |
| MM | W4 | Sprague Dawley Rat | Rattus norvegicus domestica |
| MQ | Y4 | Sprague Dawley Rat | Rattus norvegicus domestica |
| MR | Y4 | Sprague Dawley Rat | Rattus norvegicus domestica |
| MR2 | Y4 | Sprague Dawley Rat | Rattus norvegicus domestica |
| MR3 | Y4 | Sprague Dawley Rat | Rattus norvegicus domestica |
| JM1 | N3 | Svalbard Reindeer | Rangifer tarandus platyrhynchus |
| JM2 | N3 | Svalbard Reindeer | Rangifer tarandus platyrhynchus |
| KA | U3 | Treeshrew | Tupaia belangeri |
| HW | W2 | Whinchats | Saxicola rubetra |
| IQ | E3 | Wistar Rat | Rattus norvegicus domestica |
| IR | E3 | Wistar Rat | Rattus norvegicus domestica |
| GE | G2 | Wistar Rat | Rattus norvegicus domestica |
| GF | G2 | Wistar Rat | Rattus norvegicus domestica |
| JD | J3 | Wistar Rat | Rattus norvegicus domestica |
| LN | J4 | Wistar Rat | Rattus norvegicus domestica |
| LO | J4 | Wistar Rat | Rattus norvegicus domestica |
| JJ | L3 | Wistar Rat | Rattus norvegicus domestica |
| JK | L3 | Wistar Rat | Rattus norvegicus domestica |
| EB | M1 | Wistar Rat | Rattus norvegicus domestica |
| JS | Q3 | Wistar Rat | Rattus norvegicus domestica |
| JT | Q3 | Wistar Rat | Rattus norvegicus domestica |
| CB | T | Wistar Rat | Rattus norvegicus domestica |
| KL | Z3 | Zebrafish | Danio rerio |

```
# Now reading in mass from external (additional) sources,
# and adjusting species names to match those provided in
# the global data frame.

Mass <- read.csv("/Users/joshuatabh/Documents/researchProjects/trent/sihMetaregression/data/final/bodyMassData.csv") %>%
    mutate(Species = ifelse(Species == "Rattus norvegicus", "Rattus norvegicus domestica",
        Species)) %>%
    select(StudyID, Species.Name = Species, Mass, Mass_SD = Mass.sd,
        Age) %>%
    ungroup()

# Pulling out rows that lack mass...

Patch <- Data %>%
    mutate(RowID = c(1:nrow(.))) %>%
    filter(is.na(Mass)) %>%
    select(-Mass, -Mass_SD) %>%
    ungroup()

# and binding them by species and age after confirming that
```

```
# all available combinations are met.

caption1 <- "List of species missing mass data"
caption2 <- paste0("List of species from which mass data has ",
    "been sourced elsewhere")

Patch %>%
    group_by(Species.Name, Age) %>%
    count() %>%
    select(-n) %>%
    rename(Species = Species.Name) %>%
    mutate(Species = sub("^(\\S+) (\\S+) ", "\\1 \\2\n", Species)) %>%
    arrange(Species) %>%
    kbl(., longtable = T, booktabs = T, caption = caption1) %>%
    kable_styling(latex_options = "striped")
```

Table 7: List of species missing mass data

| Species | Age |
|---|---|
| Aepyceros melampus | Adult |
| Bos taurus | Adult |
| Canis lupus familiaris | Adult |
| Capreolus capreolus | Adult |
| Cavia porcellus | Juvenile |
| Danio rerio | Adult |
| Equus caballus | Adult |
| Felis catus | Adult |
| Gallus gallus domesticus | Juvenile |
| Hirundo rustica | Adult |
| Mus musculus | Adult |
| Mustela vison | Adult |
| Ovis canadensis | Adult |
| Panthera tigris | Adult |
| Rangifer tarandus platyrhynchus | Adult |
| Rattus norvegicus domestica | Adult |
| Rattus norvegicus domestica | Juvenile |
| Rattus norvegicus domestica | Juvenile_14 |
| Rattus norvegicus domestica | Juvenile_7 |
| Saxicola rubetra | Adult |
| Somateria mollissima | Adult |
| Spermophilus beecheyi | Adult |
| Sus scrofa domesticus | Juvenile |
| Sylvia borin | Adult |
| Tupaia belangeri | Adult |
| Vulpes vulpes | Adult |

```
Mass %>%
    group_by(Species.Name, Age) %>%
    count() %>%
    select(-n) %>%
    rename(Species = Species.Name) %>%
    mutate(Species = sub("^(\\S+) (\\S+) ", "\\1 \\2\n", Species)) %>%
    arrange(Species) %>%
    kbl(., longtable = T, booktabs = T, caption = caption2) %>%
    kable_styling(latex_options = "striped")
```

Table 8: List of species from which mass data has been sourced elsewhere

| Species | Age |
|---|---|
| Aepyceros melampus | Adult |
| Bos taurus | Adult |
| Canis lupus familiaris | Adult |

| | |
|---|---|
| Capreolus capreolus | Adult |
| Cavia porcellus | Adult |
| Cavia porcellus | Juvenile |
| Ceratotherium simum | Adult |
| Cyanistes caeruleus | Adult |
| Equus caballus | Adult |
| Felis catus | Adult |
| Gallus gallus domesticus | Adult |
| Hirundo rustica | Adult |
| Macaca mulatta | Adult |
| Melopsittacus undulatus | Adult |
| Mus musculus | Adult |
| Mustela vison | Adult |
| Oryctolagus cuniculus domesticus | Adult |
| Oryctolagus cuniculus domesticus | Juvenile |
| Ovis canadensis | Adult |
| Panthera tigris | Adult |
| Rangifer tarandus platyrhynchus | Adult |
| Rattus norvegicus domestica | Adult |
| Rattus norvegicus domestica | Juvenile |
| Rattus norvegicus domestica | Juvenile_14 |
| Rattus norvegicus domestica | Juvenile_7 |
| Saxicola rubetra | Adult |
| Somateria mollissima | Adult |
| Spermophilus beecheyi | Adult |
| Sus scrofa domesticus | Adult |
| Sylvia borin | Adult |
| Tupaia belangeri | Adult |
| Vulpes vulpes | Adult |

```
rm(caption1, caption2)

# Good - joining data.

Patch <- left_join(Patch, Mass, by = c("StudyID", "Species.Name",
    "Age")) %>%
    select(c(RowID, colnames(Data)))

Gap <- Data %>%
    mutate(RowID = c(1:nrow(.))) %>%
    filter(!is.na(Mass)) %>%
    select(c(RowID, colnames(Data)))

# Checking bind, returning data object name to its original
# value, and removing objects that are no longer needed.

if ("FALSE" %in% c(rbind(Patch, Gap) %>%
    arrange(RowID) %>%
    pull(RowID) == c(1:nrow(Data)))) {
    print("Bind failed")
} else {
    print("Bind correct")
}
```

```
## [1] "Bind correct"
```

```
Data <- rbind(Patch, Gap) %>%
    arrange(RowID)

rm(RowID, Gap, Mass)

# Calculating the mean coefficient of variation around mass
# then using this CV to estimate missing standard
# deviations
```

```
CV = mean(with(Data, Mass_SD/Mass), na.rm = T)
Data = Data %>%
    mutate(Mass_SD = ifelse(is.na(Mass_SD), Mass * CV, Mass_SD))

# Lastly, mass and it's standard deviation are transformed
# on a log scale. Standard deviation for samples with no
# known error are assigned a negligible standard deviation
# value (0.0001).

Data$logMass <- log(Data$Mass)
Data$logMass_SD <- log(Data$Mass + Data$Mass_SD) - log(Data$Mass)
```

Several studies have reported that ambient temperature influences the magnitude of stress-induced changes in core body temperature and surface body temperature (see Wilber and Robinson, 1958; Yokoi, 1966; Briese, 1991; Drugan et al, 2005; Lewden et al, 2017; Muise et al, 2018; Nord and Folkow, 2019; Robertson et al, 2020a; but see: Briese and Cabanac, 1991; Long et al, 1990). We were interested in testing this relationship across studies and species. To do so, we sought to extract the average ambient temperature (°C ± s.d.) from each study included in our meta-regression. However, some studies reported a range of ambient temperatures observed during data collection. Similar to our body mass approximations, we assumed mean ambient temperature to be the hypothetical median of the temperature range in these cases, and the associated standard deviation to be the range divided by four. If ambient temperature was not reported, we sought to obtain measures of ambient temperature by contacting authors or by extracting hourly values from a weather station no greater than 50 km from the location of study. Similar to our body mass measurements, if no error around mean ambient temperature was provided, we assumed it to be the average standard deviation observed across all studies. Here, we did not estimate error from our sample mean coefficient of variation because both fluctuations in ambient temperature and error in ambient temperature measurement was unlikely to vary across mean ambient temperatures.

```
# Similar to mass, where ambient temperature during stress
# exposure is not reported, we assume it to be the same as
# that reported for general animal housing or observation.

Data <- Data %>%
    mutate(Ambient.Temp.Stressor = ifelse(Ambient.Temp.Stressor ==
        "", NA, Ambient.Temp.Stressor)) %>%
    mutate(Ambient.Temp.Stressor = ifelse(is.na(Ambient.Temp.Stressor) &
        !is.na(Ambient.Temp.No.Stressor), Ambient.Temp.No.Stressor,
        Ambient.Temp.Stressor)) %>%
    as.data.frame()

# Ambient temperature ranges are split and used to estimate
# a hypothetical median.

Range_Split <- function(x) {
    X_chop <- str_split(x, "-")
    correct <- function(x) {
        x <- lapply(x, as.numeric)
        x <- lapply(x, mean)
        return(x)
    }
    y <- correct(X_chop)
    return(unlist(y))
}

To_Split <- grepl(pattern = "[[:digit:]]-", Data$Ambient.Temp.Stressor)
Data$Ambient.Temp.Stressor[To_Split] <- as.character(Range_Split(Data$Ambient.Temp.Stressor[To_Split]))

To_Split <- grepl(pattern = "[[:digit:]]-", Data$Ambient.Temp.No.Stressor)
Data$Ambient.Temp.No.Stressor[To_Split] <- as.character(Range_Split(Data$Ambient.Temp.No.Stressor[To_Split]))

Data <- Data %>%
    mutate(Ambient.Temp.Stressor = as.numeric(Ambient.Temp.Stressor),
```

```
        Ambient.Temp.No.Stressor = as.numeric(Ambient.Temp.No.Stressor))

Data <- Data %>%
    mutate(Ambient.Temp.No.Stressor.sd = ifelse(is.na(Ambient.Temp.No.Stressor.sd),
        mean(Ambient.Temp.No.Stressor.sd, na.rm = T), Ambient.Temp.No.Stressor.sd),
        Ambient.Temp.Stressor.sd = ifelse(is.na(Ambient.Temp.Stressor.sd),
            mean(Ambient.Temp.Stressor.sd, na.rm = T), Ambient.Temp.Stressor.sd))
```

#### 2 | Filtering data according to criteria for inclusion

Although we modeled body mass and ambient temperature as latent variables in our later analyses (described in sections 6-8 below), we chose not to interpolate missing values of each variable. This decision was made for two reasons: (1) we did not have reliable species or context specific predictors of each variable, and (2) we wanted our estimates of correlations between our response variable (the magnitude of stress-induced changes in body temperature) and both body mass and ambient temperature to be conservative and conditional upon true values of each variable (plus some known, or approximated, error). Thus, all studies where body mass or ambient temperature could not be estimated were excluded from further analyses.

Beyond studies missing measures of body mass or ambient temperature, we also chose to exclude studies from our analysis that: (1) did not measure the magnitude of stress-induced changes in body temperature, (2) measured stress-induced changes in body temperature 8 hours or longer after stressor onset (thus limiting our data to "acute" responses), (3) did not report an experimental or observational sample size (a critical value for effect size calculation) (4) treated study individuals with pharmaceutical agents during experimentation, (5) were exposed to a stressor in any medium other than air (in which disruptions to body temperature may already occur), (6) used transgenic lines of animals, thus producing results that could not be generalised to wild populations, or (7) used an ectothermic species as a model, for which the energetic interpretation of a change in body temperature may differ from that for endothermic species. All studies falling within these seven exclusion criteria are removed below either directly, or indirectly by having been "flagged" during the data collection process (i.e. given an "X" in the data column "Flag").

```
# First, counting studies that lacked measures of body mass
# or ambient temperature and removing them.

nrow(subset(Data, is.na(logMass)))
```

```
## [1] 6
```

```
Data <- subset(Data, !is.na(logMass))

nrow(subset(Data, is.na(Ambient.Temp.Stressor)))
```

```
## [1] 36
```

```
Data <- subset(Data, !is.na(Ambient.Temp.Stressor))

# Second, counting and excluding studies that did not
# report a sample size

nrow(subset(Data, is.na(Exp.Sample.Size)))
```

```
## [1] 7
```

```
Data <- subset(Data, !is.na(Exp.Sample.Size))

# Third, counting and removing studies that occurred in
# water.

nrow(subset(Data, Stress.Medium == "Water"))
```

```
## [1] 13
```

```
Data <- subset(Data, Stress.Medium == "Air")

# Fourth, counting and excluded data from non-endotherms
```

```
nrow(subset(Data, !(Class %in% c("Aves", "Mammalia"))))
```

```
## [1] 5
```

```
Data <- subset(Data, Class %in% c("Aves", "Mammalia"))

# Fifth, calculating mean time to measurement, then
# counting and excluding data pertaining to measurements >
# 8 hours after the onset of a stress exposure

Data %>%
    mutate(Time = as.numeric(Time.to.Peak.s.)) %>%
    summarise(Mean = mean(Time, na.rm = T), SD = sd(Time, na.rm = T))
```

```
##       Mean       SD
## 1 3517.553 7859.244
```

```
Data <- Data %>%
    mutate(Time.to.Peak.s. = as.numeric(Time.to.Peak.s.))
nrow(subset(Data, Time.to.Peak.s. >= 3600 * 8))
```

```
## [1] 4
```

```
Data <- subset(Data, Time.to.Peak.s. < 3600 * 8)

# Fifth, counting remaining flagged studies (that fall
# under exclusion criteria 1, 3, and 6) and removing them.

nrow(subset(Data, Flag == "X"))
```

```
## [1] 4
```

```
Data <- subset(Data, Flag != "X")
```

#### 3 | Estimating relative resting metabolic rate across species

Theory predicts that trade-offs between biological processes should only occur when energy or resources are limited (Stearns, 1992). One way that this limitation may occur is by expending relatively high amounts of energy or resources on maintenance, thus leaving little remaining for allocation elsewhere (Careau, 2017). This potential avenue toward a trade-off is particularly convenient to study, given that rates of expenditure towards maintenance are already known for many species (i.e. "resting metabolic rate", or "RMR"). In our study, we tested whether these readily-available measures of expenditure could explain whether and how the body temperature of a given species sample changed after exposure to a stressor. Under our trade-off hypothesis, those with a relatively high expenditure toward maintenance (or relatively high RMR) should be more likely to decrease their body temperatures at, or below, thermoneutrality than those with a relatively low expenditure toward maintenance.

To first ensure that our measure of RMR was indeed relative and not largely explained by a species body mass, we regressed log-transformed measures of metabolic rate (here, in Watts, or "W") against log-transformed body mass (g) and extracted mean ordinary residuals (henceforth "residuals") from our posterior distribution. Importantly, because measures of metabolic rate in our data set represented both basal or resting values (which are likely to differ), we included metabolic rate metric (binomial, "resting" or "basal") as a group-level intercept in our regression. Metabolic rate metric was not included as a population-level predictor because we were not interested in estimating its effect on cross-species metabolic rate *per se*, and to ease interpretation of our residuals.

Before conducting our regression, we imported, compiled, and summarised data pertaining to species-specific basal or resting metabolic rate. Some studies included in our meta-regression report stress-induced changes in body temperature for various specified seasons, among which, energetic expenditure toward maintenance is likely to vary. Where possible, we therefore sought to identify basal or resting metabolic rate for each season at which stress-induced changes in body temperature were reported for a given species. In some case, however, such season-specific data was unavailable in primary literature. Moreover, to our knowledge, some species lacked any primary data on basal or resting metabolism regardless of season. Thus, where necessary, metabolic data from the nearest available season and/or species (using consensus phylogeny described in section 6 below) were paired with season-specific measures of stress-induced changes in body temperature.

```r
# Loading in data pertaining to species-specific energy
# consumption. Note that column names are corrected to ease
# data-frame bind later on.

MO2 <- read.csv("/Users/joshuatabh/Documents/researchProjects/trent/sihMetaregression/data/final/metabolicData.csv") %>%
    rename(Species.Name = Latin.Name, Season = MR.Season) %>%
    select(-X)

# Now, season-specific changes in body temperature are
# separated out so that we may pair these values with
# season-specific energy expenditure.

Seasonal_Data <- subset(Data, Common.Name %in% c("Black-capped Chickadee",
    "California Ground Squirrel", "Svalbard Rock Ptarmigan"))

# Checking whether other studies included these species
# exist without season reported.

nrow(Seasonal_Data[c(which(is.na(Seasonal_Data$Season))), ]) >
    0  # Yes.
```

```
## [1] TRUE
```

```r
Seasonal_Data_Subset <- subset(Seasonal_Data, !is.na(Season))

# Switching spring season for Svalbard Rock Ptarmigan to
# summer (because only data for summer and winter are
# available).

Seasonal_Data_Subset$Season[c(which(Seasonal_Data_Subset$Common.Name ==
    "Svalbard Rock Ptarmigan" & Seasonal_Data_Subset$Season ==
    "Spring"))] <- "Summer"

MO2_Data_Subset <- subset(MO2, Common.Name %in% c("Black-capped Chickadee",
    "California Ground Squirrel", "Svalbard Rock Ptarmigan"))

# Binding together energy expenditure and delta body
# temperature data for seasonal measures.

Subset_Bound <- left_join(Seasonal_Data_Subset, MO2_Data_Subset,
    by = c("Species.Name", "Common.Name", "Season"))

# Appending California Ground Squirrel study with season
# undefined (for which energy expenditure is averaged
# across season, and the thermoneutral zone is assumed,
# conservatively, to be the widest possible).

CGS <- subset(MO2_Data_Subset, Common.Name == "California Ground Squirrel") %>%
    group_by(Common.Name, Species.Name) %>%
    summarise(LCT = min(LCT), UCT = max(UCT), BMR.RMR = BMR.RMR[1],
        MR.W = mean(MR.W)) %>%
    mutate(Conversion.Coefficient = paste(subset(MO2_Data_Subset,
        Common.Name == "California Ground
                    Squirrel")$Conversion.Coefficient[1:2],
        collapse = "; ")) %>%
    mutate(Conversion.Coefficient.Source = paste(subset(MO2_Data_Subset,
        Common.Name == "California Ground
                    Squirrel")$Conversion.Coefficient.Source[1:2],
        collapse = "; ")) %>%
    mutate(MR.Protocol = "Respirometry") %>%
    mutate(Season = "All") %>%
    mutate(Notes = paste(subset(MO2_Data_Subset, Common.Name ==
        "California Ground Squirrel")$Notes[1], subset(MO2_Data_Subset,
        Common.Name == "California Ground Squirrel")$Notes[2],
        sep = ";")) %>%
```

```r
    mutate(Source = paste(subset(MO2_Data_Subset, Common.Name ==
        "California Ground Squirrel")$Source[1], subset(MO2_Data_Subset,
        Common.Name == "California Ground Squirrel")$Source[2],
        sep = ";"))

# Checking that no changes in column names have occurred
# during this process.

length(grep("FALSE", colnames(CGS) == colnames(MO2_Data_Subset))) >
    0
```

```
## [1] FALSE
```

```r
# Good. Now integrating this data from the California
# Ground Squirrel.

Seasonal_Data_Subset <- subset(Seasonal_Data, is.na(Season))
Second_Subset_Bound <- left_join(Seasonal_Data_Subset, CGS, by = c("Species.Name",
    "Common.Name")) %>%
    rename(Season = Season.x) %>%
    select(-Season.y)

Season_Specific <- rbind(Subset_Bound, Second_Subset_Bound)

# Cleaning column names for simplicity.

Season_Specific <- Season_Specific %>%
    rename(Tb_Notes = Notes.x, Tb_Source = Source.x, Metabolic_Notes = Notes.y,
        Metabolic_Source = Source.y, Tb_Season = Season) %>%
    mutate(MR_Season = Tb_Season)

# Now binding non-seasonal data pertaining to
# stress-induced changes in body temperature and energy
# expenditure.

Nonseasonal_Data <- subset(Data, !(Common.Name %in% c("Black-capped Chickadee",
    "California Ground Squirrel", "Svalbard Rock Ptarmigan")))

# Column names must be adjusted to match the new names
# defined above.

Nonseasonal_Joined <- left_join(Nonseasonal_Data, MO2, by = c("Species.Name",
    "Common.Name")) %>%
    rename(Tb_Notes = Notes.x, Tb_Source = Source.x, Metabolic_Notes = Notes.y,
        Metabolic_Source = Source.y, Tb_Season = Season.x, MR_Season = Season.y)

# Here, we make sure that column names are matched between
# seasonal and non-seasonal data sets.

Nonseasonal_Joined <- Nonseasonal_Joined %>%
    select(colnames(Season_Specific))

# And now, we bind seasonal and non-seasonal data together,
# check that no data duplications have occurred, then
# rename our data objected back to 'Data'. All old data
# objects are also dropped below.

All_Data <- rbind(Season_Specific, Nonseasonal_Joined)
nrow(All_Data) == nrow(Data)
```

```
## [1] TRUE
```

```r
Data <- All_Data

rm(MO2, Seasonal_Data, Seasonal_Data_Subset, Subset_Bound, CGS,
    Second_Subset_Bound, Season_Specific, Nonseasonal_Data, Nonseasonal_Joined,
    All_Data)
```

```r
# Proceeding by plotting log-transformed metabolic rate
# measures against log-transformed body mass.

pal <- colRoz_pal(name = "c.azureus", n = 20, type = "continuous")

Data %>%
    group_by(Class, Order, Species.Name) %>%
    summarise(logMass = log(mean(Mass, na.rm = T)), logMR = log(mean(MR.W,
        na.rm = T))) %>%
    ggplot(aes(x = logMass, y = logMR)) + geom_point(size = 2,
    pch = 21, colour = "black", alpha = 0.7, aes(fill = Order)) +
    geom_smooth(method = "lm", colour = "black", size = 1, se = FALSE) +
    theme_classic() + scale_colour_gradientn(colours = pal) +
    xlab("log Body Mass (g)") + ylab("log Metabolic Rate (W)")
```

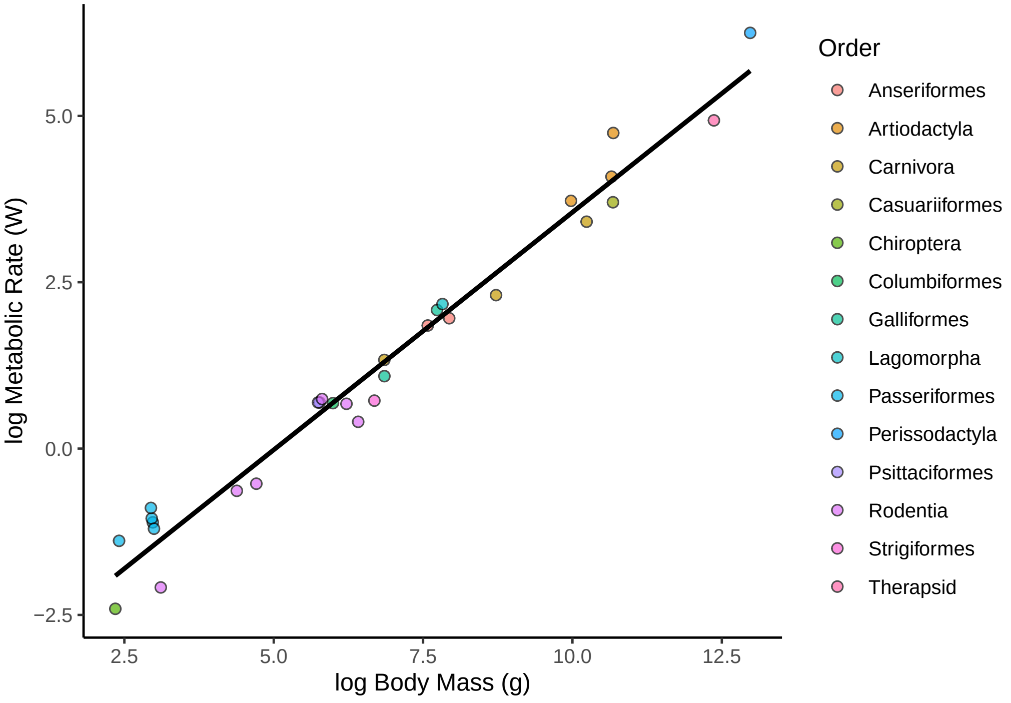


Figure 1: Natural log-transformed resting metabolic rate as a function of average body mass among terrestrial endothermic species.

```r
# And checking raw differences between basal and resting
# metabolic rate measurements.

Data %>%
    group_by(Class, Order, Species.Name, Common.Name) %>%
    summarise(logMR = log(mean(MR.W, na.rm = T)), BMR.RMR = BMR.RMR[1]) %>%
    mutate(BMR.RMR = ifelse(BMR.RMR == "BMR", "Basal", "Resting")) %>%
    ggplot(aes(x = BMR.RMR, y = logMR, fill = BMR.RMR)) + geom_boxplot(width = 0.2) +
    ggdist::stat_halfeye(slab_colour = "black", adjust = 0.7,
        width = 0.3, .width = 0, justification = -0.8, point_colour = NA) +
    geom_point(size = 1, alpha = 0.3, position = position_jitter(seed = 1,
        width = 0.1)) + coord_cartesian(xlim = c(1.2, NA), clip = "off") +
    theme_classic() + scale_fill_manual(values = c(nice_pink,
    "slateblue")) + theme(legend.position = "none") + xlab("Measurement Type") +
    ylab("log Metabolic Rate (W)")
```

```r
# Clearly higher metabolic rate when classified as resting.

# Now creating data to be used in regression.
```

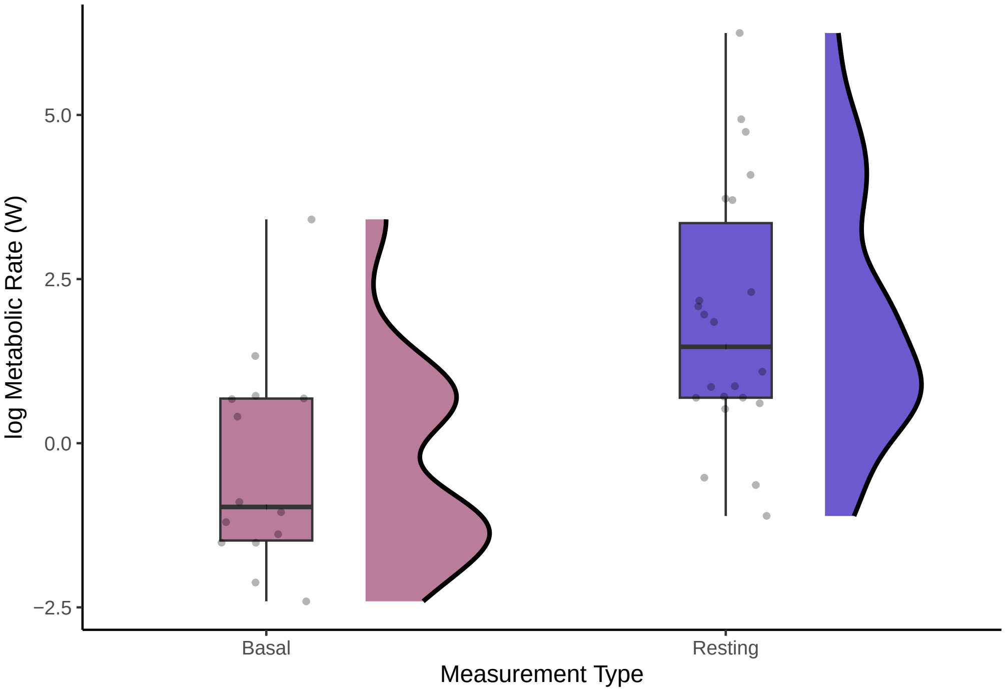


Figure 2: Distribution of natural log-transformed metabolic rate measurements according to whether such were classified as resting or basal.

```
ResData <- Data %>%
    group_by(Class, Order, Species.Name, Common.Name) %>%
    # filter(Age == 'Adult') %>%
summarise(logMass = log(mean(Mass, na.rm = T)), logMR = log(mean(MR.W,
    na.rm = T)), BMR.RMR = BMR.RMR[1]) %>%
    filter(!is.na(logMR)) %>%
    filter(!is.na(logMass))

# Defining an informative prior for our model with a slope
# between log-mass and log-metabolic rate of 0.67. The
# influence of metabolic rate metric on metabolic rate
# measurements is assumed weak and vague, with a gamma
# distribution of alpha = 1 and beta = 1.

prior_Mass <- c(set_prior("normal(0.67, 0.1)", class = "b"),
    set_prior("gamma(1, 1)", class = "sd"), set_prior("gamma(1, 1)",
        class = "sigma")  #
)

# Running model and saving output.

MR_Mod_Output = "/Users/joshuatabh/Documents/researchProjects/trent/sihMetaregression/models/metabolismByMass.Rds"

MR_Mod <- brm(logMR ~ logMass + (1 | BMR.RMR), data = ResData,
    cores = 1, chains = 4, seed = 100, family = "gaussian", iter = 50000,
    warmup = 5000, thin = 10, control = list(adapt_delta = 0.99,
        max_treedepth = 16), prior = prior_Mass, file = MR_Mod_Output)

# Checking the Bayesian (conditional) R2 of our model.

r2_bayes(MR_Mod)
```

```
## # Bayesian R2 with Compatibility Interval
##
```

```
##   Conditional R2: 0.972 (95% CI [0.966, 0.974])
##      Marginal R2: 0.970 (95% CI [0.961, 0.972])
```

```
# Marginal R2 = 0.970; quite high. Checking for
# autocorrelation in our posteriors, and assessing chain
# convergence with a Gelman-Rubin (Rhat) statistic.
```

```
clean_ac(MR_Mod, prs = c("b_Intercept", "b_logMass", "sd_BMR.RMR__Intercept",
    "sigma"), names = c("Intercept", "log Mass (g)", "MR Intercept",
    "Sigma"))
```

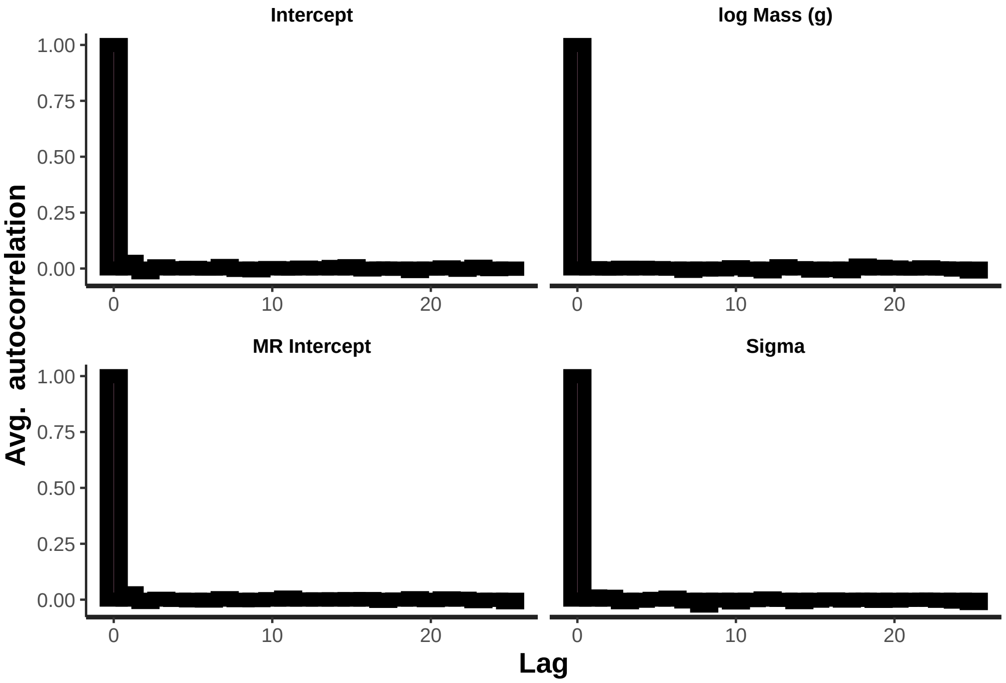


Figure 3: Lag plots displaying degree of sequential autocorrelation between chain draws, per parameter, for a Bayesian, linear model predicting log resting metabolic rate as a function of log body mass.

```
grid.arrange(mcmc_neff(neff_ratio(MR_Mod), size = 2) + theme(legend.position = "none"),
    mcmc_rhat(rhat(MR_Mod)) + theme(legend.position = "none"),
    nrow = 1)
```

```
# No evident autocorrelation, but ratio of effective sample
# sizes to sample sizes is quite poor in some instances.
# Rhats values are very close to 1, however, suggesting
# nice chain convergence/mixing.
```

```
# Now viewing MCMC draws by parameter, as intervals
# (quantile) and pairs plots.

clean_int(MR_Mod, prs = c("b_Intercept", "b_logMass", "sd_BMR.RMR__Intercept",
    "sigma"), names = c("Intercept", "log Mass", "MR Intercept",
    "Sigma"))
```

```
clean_pairs(MR_Mod, prs = c("b_logMass", "sd_BMR.RMR__Intercept",
    "sigma"), names = c("log Mass", "MR Intercept", "Sigma"))
```

```
# Some noise around random intercept, but this is expected,
# particularly given our weak prior and differences in
# study techniques.

# Checking the distribution of our mean, ordinary
# residuals.
```

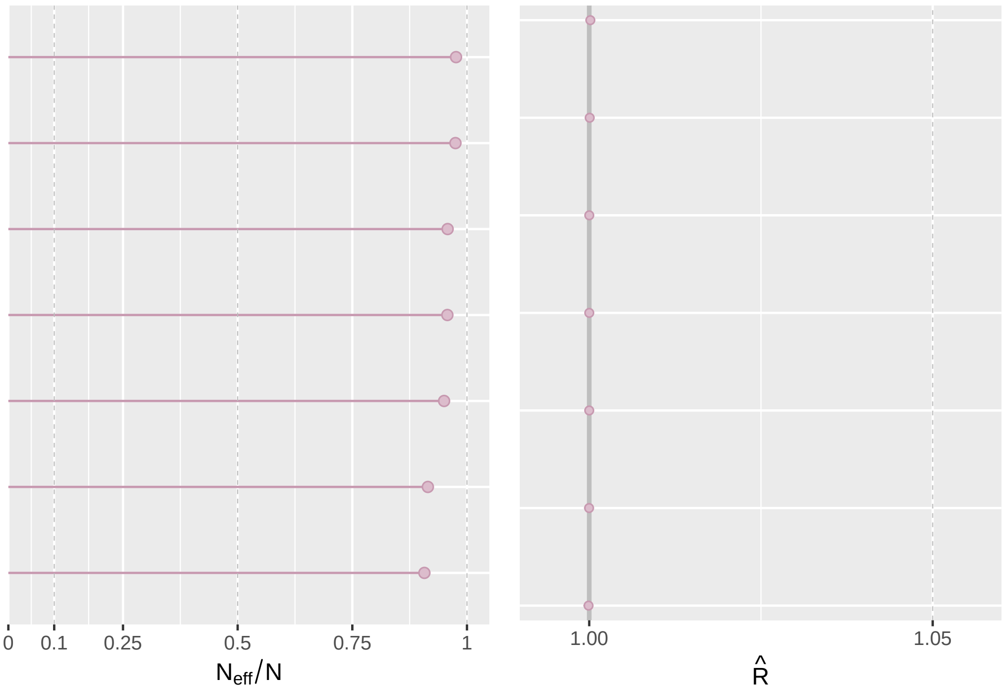


Figure 4: Ratio of effective sample sizes to sample sizes (left panel) and Gelman-Rubin (Rhat) statistics (right panel), per parameter, for a Bayesian, linear model predicting log resting metabolic rate as a function of log body mass.

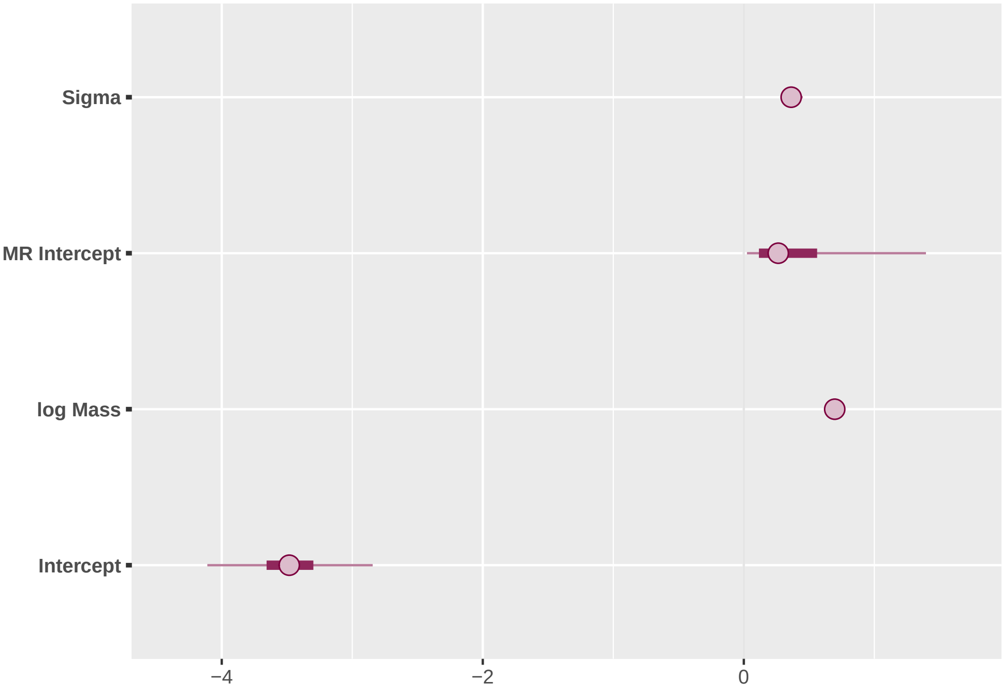


Figure 5: Select parameters and their 90 percent posterior intervals from a Bayesian, linear model predicting log resting metabolic rate as a function of log body mass.

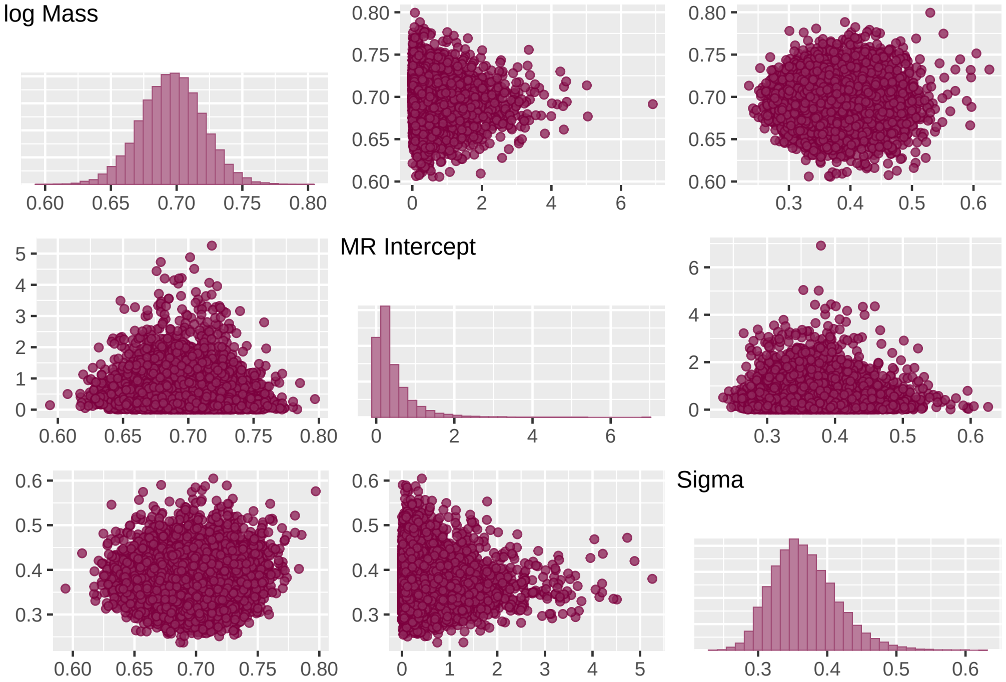


Figure 6: Posterior draws across model parameters from a Bayesian, linear model predicting log resting metabolic rate as a function of log body mass.

```
MR_Mod$data %>%
    dplyr::select(logMR, logMass, BMR.RMR) %>%
    na.omit(.) %>%
    add_residual_draws(MR_Mod) %>%
    group_by(logMR, logMass, BMR.RMR) %>%
    summarise(Res = mean(.residual)) %>%
    ggplot(aes(x = 1:nrow(.), y = Res)) + geom_point() + geom_smooth(colour = "black") +
    xlab("Sample Number") + ylab("Mean Ordinary Residuals") +
    theme_classic()
```

```
# No clear patterning. Now assessing residual values by
# fitted values

MR_Mod$data %>%
    dplyr::select(logMR, logMass, BMR.RMR) %>%
    na.omit(.) %>%
    add_residual_draws(MR_Mod) %>%
    group_by(logMR, logMass, BMR.RMR) %>%
    summarise(Res = mean(.residual)) %>%
    ungroup() %>%
    mutate(Fit = colMeans(posterior_predict(MR_Mod))) %>%
    ggplot(aes(x = Fit, y = Res)) + geom_point() + geom_smooth(colour = "black") +
    xlab("Mean Fitted Values") + ylab("Mean Ordinary Residuals") +
    theme_classic()
```

```
# Also good. Plotting the posterior distributions, printing
# out mode coefficients, and calculating 50% and 95%
# highest posterior density intervals.
```

```
clean_dens(MR_Mod, prs = c("b_Intercept", "b_logMass", "sd_BMR.RMR__Intercept"),
    names = c("Intercept", "log Mass (g)", "MR Intercept (s.d.)"))
```

```
MR_Mod %>%
    spread_draws(b_Intercept, b_logMass, sd_BMR.RMR__Intercept,
        `r_BMR.RMR[BMR,Intercept]`, `r_BMR.RMR[RMR,Intercept]`) %>%
    select(-c(.chain, .iteration, .draw), Intercept = b_Intercept,
```

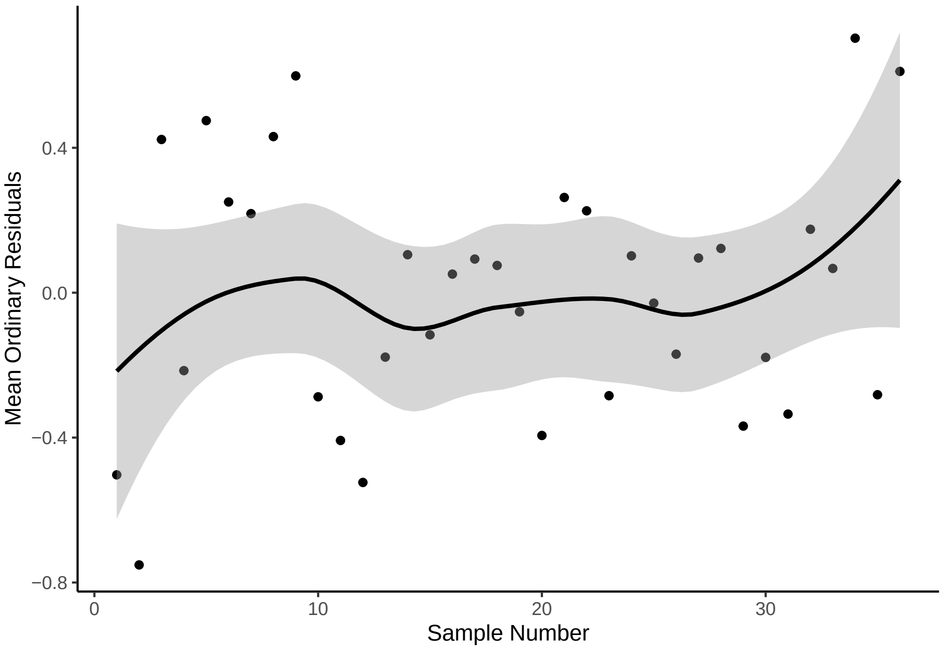


Figure 7: Mean ordinary residuals according to data-set sample number from a Bayesian, linear model predicting log resting metabolic rate as a function of log body mass.

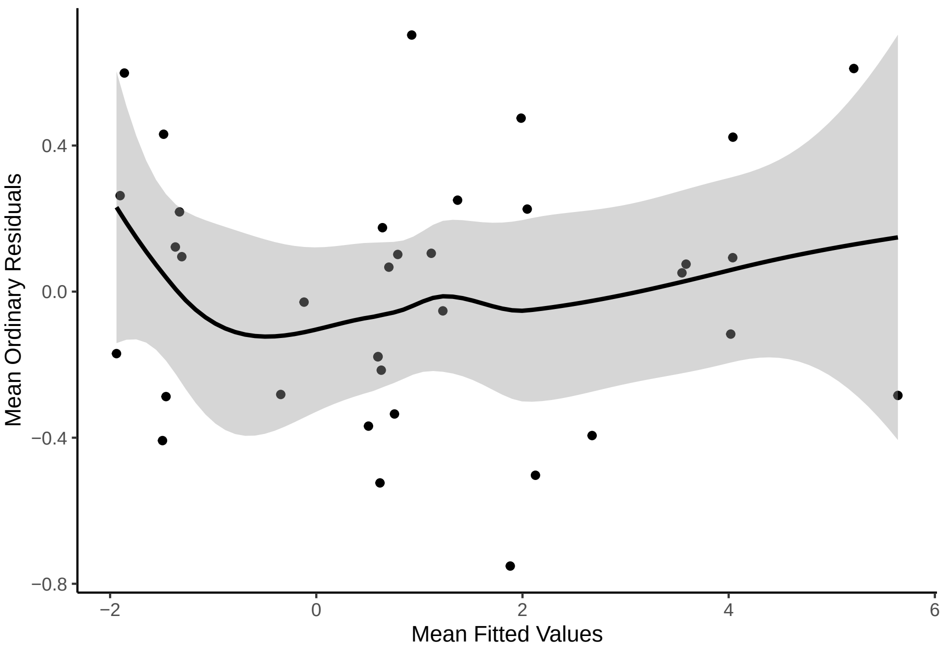


Figure 8: Mean ordinary residuals according to fitted values from a Bayesian, linear model predicting log resting metabolic rate as a function of log body mass.

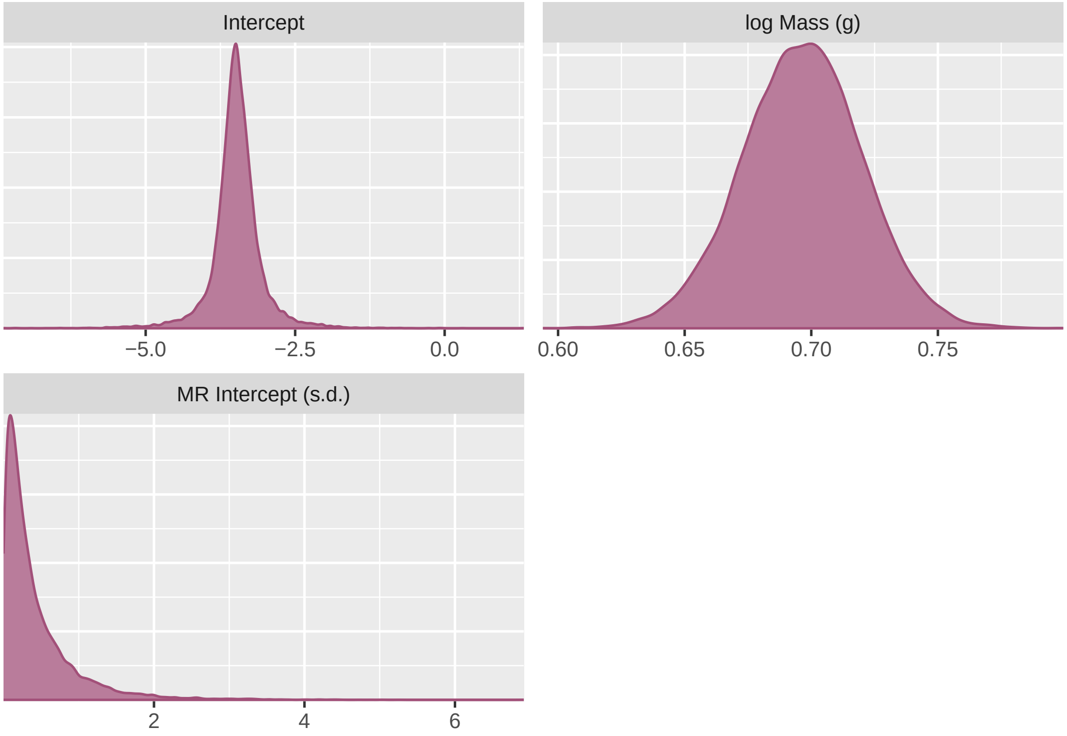


Figure 9: Densities of posterior draws, per model parameter, from a Bayesian, linear model predicting log resting metabolic rate as a function of log body mass.

```
        `log Mass` = b_logMass, `MR Intercept (s.d.)` = sd_BMR.RMR__Intercept,
        `BMR Intercept` = `r_BMR.RMR[BMR,Intercept]`, `RMR Intercept` = `r_BMR.RMR[RMR,Intercept]`) %>%
    mutate_all(., .funs = round, 5) %>%
    summarise_all(., .funs = md) %>%
    pivot_longer(everything(), names_to = "Parameter", values_to = "Mode") %>%
    cbind(., simple_hdi(MR_Mod, cis = c(80, 95), rnd = 5) %>%
        select(-Parameter)) %>%
    mutate(`80% HDI` = paste0("(", paste(Low_HDI_80, High_HDI_80,
        sep = ", "), ")"), `95% HDI` = paste0("(", paste(Low_HDI_95,
        High_HDI_95, sep = ", "), ")")) %>%
    select(-c(Low_HDI_80, High_HDI_80, Low_HDI_95, High_HDI_95)) %>%
    kbl(., longtable = T, booktabs = T) %>%
    kable_styling(latex_options = "striped")
```

| Parameter | Mode | 80% HDI | 95% HDI |
|---|---|---|---|
| Intercept | -3.41956 | (-3.87324, -3.06732) | (-4.35678, -2.54882) |
| log Mass | 0.70269 | (0.66711, 0.72711) | (0.64858, 0.74231) |
| MR Intercept (s.d.) | 0.05786 | (-0.44083, 0.2689) | (-0.97693, 0.76541) |
| BMR Intercept | 0.00088 | (-0.25155, 0.45238) | (-0.74672, 1.00493) |
| RMR Intercept | -0.00074 | (2e-05, 0.66655) | (2e-05, 1.39636) |

```
# And now extracting mean residuals and standard deviations
# for later analysis.

ResData <- ResData %>%
    ungroup() %>%
    mutate(ResMR = residuals(MR_Mod, type = "ordinary")[, 1],
        ResMR_SE = residuals(MR_Mod, type = "ordinary")[, 2]) %>%
    mutate(ResMR_Bin = ifelse(ResMR < 0, "Low", "High"))

# Binding these residuals into our main data object.

ResBind <- left_join(Data, ResData %>%
    select(-c(logMass, logMR)), by = c("Class", "Order", "Species.Name",
    "Common.Name"))
```



```
# Note that log-transformed metabolic rate estimates were
# derived only from adults. Given demands for growth, we
# cannot assume that these metabolic rate estimates apply
```

#### 4 | Calculating log-transformed response ratios

Average body temperature is known to vary among endothermic species (discussed in Angilletta et al, 2010), and among endothermic conspecifics (e.g. Bozinovic, 2007; Møller, 2010; Szafranska et al, 2020). Such variation in average body temperature is likely to influence the proportional value of a given, discrete change in body temperature per species and per individual (i.e. in °C). Thus, to relativise our measures of stress-induced changes in body temperature across sample populations, we converted our measures of absolute change in body temperature to a relative change in body temperature (or, a "response ratio") defined as:

$$\frac{T_b 1_i}{T_b 0_i}$$

where $T_b1$ represents an average minimal, or an average maximal body temperature of study population *i* that was observed after the onset of a stress exposure treatment, and $T_b0$ represents the body temperature of study population *i prior* to a stress exposure treatment. For studies that also recorded the body temperatures of control (or "unstressed") individuals throughout experimentation, the above equation was replaced with:

$$\frac{T_b 1_{Si}}{T_b 1_{Ci}}$$

where $T_b1_S$ again represents an average minimal, or an average maximal body temperature of the stress exposed population *i*, observed after the onset of a stress exposure treatment, and $T_b1_C$ represents the average body temperature of the control population at the equivalent time point.

To estimate uncertainty around our response ratio values, we calculated a pooled standard error using one of the following equation:

$$(1) \quad \sqrt{\frac{\sigma_{\bar{T_b}1}}{T_b 1} + \frac{\sigma_{\bar{T_b}0}}{T_b 0}}$$

or

$$(2) \quad \sqrt{\frac{\sigma_{\bar{T_b}1_S}}{T_b 1_S} + \frac{\sigma_{\bar{T_b}1_C}}{T_b 1_C}}$$

where equation (1) applies to studies without a control population, equation (2) applies to studies with a control population, $\bar{\sigma}$ represents the standard error of a given subscripted variables, and all other variables remain as previously described. In studies where a standard error around a stress-induced change in body temperature was reported, we assumed it to be a pooled estimate and used it in place of any estimate derived from an equation above.

Many studies included in our original data-set reported a stress-induced change in body temperature (i.e. Δ °C) but did not report initial or final body temperature measurements (here, minimal or maximal, as described above). Still others did not report a measure of uncertainty around inital and/or final body temperature measurements. For these studies, we were therefore unable to estimate a response ratio, or a standard error around a response ratio value. Thus, all studies lacking measures or initial body temperature, final body temperature, or the uncertainty around each were removed from further study.

Finally, to both normalise our response ratio estimates and centre them on zero, we derived their natural logarithm and used these transformed values as our final estimates of effect size. Effect sizes for studies with and without control populations were visually compared to assess bias according to experimental approach.

```
# Beginning by splitting data according to whether or not a
# control population was observed.

g(Control_Data, Paired_Data) %=% list(subset(Data, Unmanipulated.Control.Y.N. ==
    "Y"), subset(Data, Unmanipulated.Control.Y.N. != "Y"))

# Proceeding to calculating effect sizes (here, a log
# response ratio). For now, dropping studies that do not
# have both an initial and peak Tb measurement. Also note
# that bias correction is not used.

Paired_Data <- Paired_Data %>%
    mutate(lRR = ifelse(!is.na(Initial.Tb) & !is.na(Peak.Trough.Tb),
        log(Peak.Trough.Tb/Initial.Tb), NA), lRV = ifelse(!is.na(Initial.Tb.SE) &
        !is.na(Peak.Tb.SE), sqrt(((Peak.Tb.SE^2)/(Peak.Trough.Tb^2)) +
        ((Initial.Tb.SE^2)/(Initial.Tb^2))), NA))

Control_Data <- Control_Data %>%
    mutate(lRR = ifelse(!is.na(Control.Tb.at.Stress.Peak) & !is.na(Control.Tb.at.Stress.Peak.SE) &
        !is.na(Peak.Trough.Tb) & !is.na(Peak.Tb.SE), log(Peak.Trough.Tb/Control.Tb.at.Stress.Peak),
        NA), lRV = ifelse(!is.na(Control.Tb.at.Stress.Peak) &
        !is.na(Control.Tb.at.Stress.Peak.SE) & !is.na(Peak.Trough.Tb) &
        !is.na(Peak.Tb.SE), sqrt(((Peak.Tb.SE^2)/(Peak.Trough.Tb^2)) +
        ((Control.Tb.at.Stress.Peak.SE^2)/(Control.Tb.at.Stress.Peak^2))),
        NA), )

Data_Bound <- rbind(Paired_Data, Control_Data)

# Comparing distributions of effect sizes and errors
# calculated from each measurement approach.
```

```
pal <- colRoz_pal(name = "c.azureus", n = 2, type = "discrete")

grid.arrange(rbind(Paired_Data %>%
    mutate(Type = "Paired"), Control_Data %>%
    mutate(Type = "Unpaired")) %>%
    ggplot(aes(x = lRR, fill = Type)) + geom_density(adjust = 3,
    colour = "black", alpha = 0.5) + theme_classic() + xlab("log Response Ratio") +
    scale_fill_manual(values = pal), rbind(Paired_Data %>%
    mutate(Type = "Paired"), Control_Data %>%
    mutate(Type = "Unpaired")) %>%
    ggplot(aes(x = lRV, fill = Type)) + geom_density(adjust = 3,
    colour = "black", alpha = 0.5) + theme_classic() + xlab("SE log Response Ratio") +
    scale_fill_manual(values = pal), nrow = 1)
```

```
# No evidence of bias. Counting samples with and without
# effect sizes.

Data_Bound %>%
    mutate(Group = ifelse(is.na(lRR), "No", "Yes")) %>%
    group_by(Group) %>%
    summarise(Count = n())
```

```
## # A tibble: 2 x 2
##   Group Count
##   <chr> <int>
## 1 No       30
## 2 Yes     199
```

```
# A substantial number of samples lost (29). Checking
# whether studies labelled as having controls lack control
# data, but contain paired, non-controlled data.

nrow(Data_Bound %>%
    mutate(Group = ifelse(is.na(lRR), "No", "Yes")) %>%
    filter(Group == "No" & Unmanipulated.Control.Y.N. == "Y") %>%
    filter(!is.na(Initial.Tb) & !is.na(Peak.Trough.Tb) & !is.na(Initial.Tb.SE) &
        !is.na(Peak.Tb.SE)) %>%
    filter(is.na(Control.Tb.at.Stress.Peak) | is.na(Control.Tb.at.Stress.Peak.SE)))
```

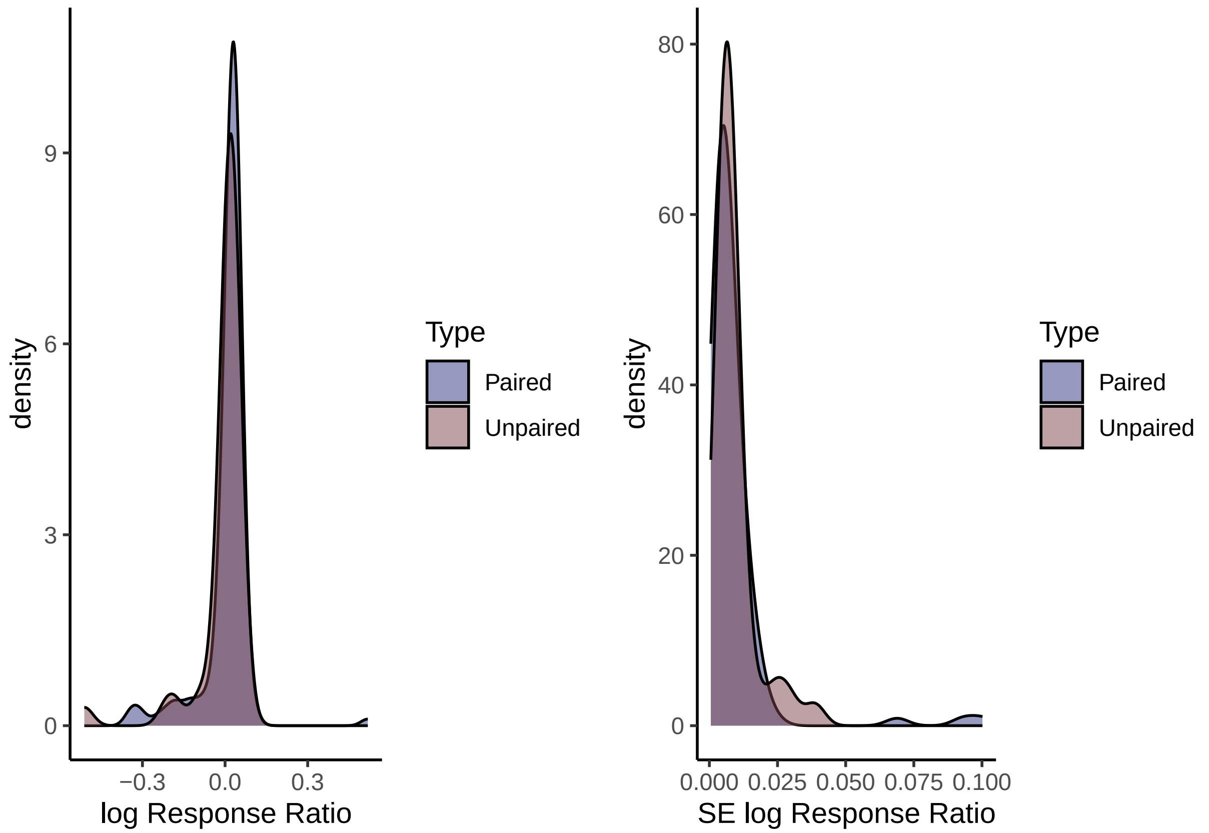


Figure 10: Comparison of effect sizes (log response ratio, or 'lRR'; left panel) and standard errors (SEs) around effect sizes as calculated from paired studies and unpaired studies. Paired studies represented those where concurrent treatment and control groups were used, whereas unpaired studies represent those lacking control groups (i.e. where comparions were made within individuals and with respect to baseline measurements).

```
## [1] 0
```

```r
# Checking adjustment

nrow(Data_Bound %>%
    mutate(Group = ifelse(is.na(lRR), "No", "Yes")) %>%
    filter(Group == "No" & Unmanipulated.Control.Y.N. == "Y") %>%
    filter(!is.na(Initial.Tb) & !is.na(Peak.Trough.Tb) & !is.na(Initial.Tb.SE) &
        !is.na(Peak.Tb.SE)) %>%
    filter(is.na(Control.Tb.at.Stress.Peak) | is.na(Control.Tb.at.Stress.Peak.SE))) ==
    0
```

```
## [1] TRUE
```

```r
# Now grabbing studies with a pooled standard error already
# estimated for a change in body temperature.

Observations <- Data_Bound %>%
    mutate(Group = ifelse(!is.na(lRR), "No", "Yes")) %>%
    filter(Group == "No") %>%
    filter(!is.na(Delta.Tb) & !is.na(Delta.Tb.SE) & !is.na(Initial.Tb)) %>%
    pull(ObservationID)

Adjust <- subset(Data_Bound, ObservationID %in% Observations)
Base <- subset(Data_Bound, !(ObservationID %in% Observations))

Adjust <- Adjust %>%
    mutate(lRR = log((Initial.Tb + Delta.Tb)/Initial.Tb), lRV = Delta.Tb.SE)

Data_Second_Bind <- rbind(Adjust, Base)
nrow(Data_Second_Bind) == nrow(Data_Bound)
```

```
## [1] TRUE
```

```r
# Viewing species missing response ratios or error data

Data_Second_Bind %>%
    filter(is.na(lRR) | is.na(lRV)) %>%
    group_by(Species.Name, Common.Name) %>%
    summarise()
```

```
## # A tibble: 13 x 2
## # Groups:   Species.Name [11]
##    Species.Name                     Common.Name
##    <chr>                            <chr>
##  1 Amazona aestiva                  Blue-Fronted Parrot
##  2 Amazona ventralis                Hispaniolan Amazon Parrot
##  3 Canis lupus familiaris           Domestic Dog
##  4 Capreolus capreolus              Roe Deer
##  5 Cavia porcellus                  Guinea Pig
##  6 Dromaius novaehollandiae         Emu
##  7 Mus musculus                     House Mouse
##  8 Oryctolagus cuniculus domesticus Domestic Albino Rabbit
##  9 Rattus norvegicus domestica      Fischer Rat
## 10 Rattus norvegicus domestica      Sprague Dawley Rat
## 11 Rattus norvegicus domestica      Wistar Rat
## 12 Spermophilus beecheyi            California Ground Squirrel
## 13 Vulpes vulpes                    Silver Fox
```

```r
nrow(Data_Second_Bind %>%
    filter(is.na(lRR) | is.na(lRV)))  # 59 data points
```

```
## [1] 59
```

```r
# Lastly, removing studies without a measure of effect size
# or effect size error.

Data = Data_Second_Bind %>%
    filter(!is.na(lRR) & !is.na(lRV))

rm(Removals, Base, Adjust)
```

Now that our measures of effect size have been calculated and all data that do not meet our inclusion criteria have been discarded, we can proceed to summarising sample sizes for our study.

```
# First, counting total number of observations.

Data %>%
    group_by(Surface.Core) %>%
    count()
```

```
## # A tibble: 1 x 2
## # Groups:   Surface.Core [1]
##   Surface.Core     n
##   <chr>        <int>
## 1 Core           170
```

```
# Number of studies included.

Data %>%
    group_by(StudyID) %>%
    summarise() %>%
    ungroup() %>%
    summarise(Count = nrow(.)) %>%
    pull(Count)
```

```
## [1] 70
```

```
# Number of species included.

Data %>%
    group_by(Species.Name) %>%
    summarise() %>%
    ungroup() %>%
    summarise(Count = nrow(.)) %>%
    pull(Count)
```

```
## [1] 25
```

```
# Average samples per species

mean(Data %>%
    group_by(Species.Name) %>%
    summarise(Count = n()) %>%
    pull(Count))
```

```
## [1] 6.8
```

```
sd(Data %>%
    group_by(Species.Name) %>%
    summarise(Count = n()) %>%
    pull(Count))
```

```
## [1] 18.62794
```

```
# Number of orders included.

Data %>%
    group_by(Order) %>%
    summarise() %>%
    ungroup() %>%
    summarise(Count = nrow(.)) %>%
    pull(Count)
```

```
## [1] 12
```

```
# Breakdown of species and classes

caption = "Number of species per class represented in final data-set."
Data %>%
    group_by(Class, Species.Name) %>%
    summarise() %>%
    ungroup() %>%
```

```
    group_by(Class) %>%
    summarise(Count = n()) %>%
    kbl(., longtable = T, booktabs = T, caption = caption) %>%
    kable_styling(latex_options = "striped")
```

Table 10: Number of species per class represented in final data-set.

| Class | Count |
|---|---|
| Aves | 12 |
| Mammalia | 13 |

#### 5 | Estimating phylogenetic relationships among study species

Because the magnitude and direction of stress-induced changes in body temperature may be correlated among closely related species, we next sought to estimate, and control for, the relative phylogenetic distance among all species included in our study. To do so, we extracted two sets of 1000 trimmed phylogenetic trees - one pertaining to mammals, and one pertaining to birds - from VertLife$^{TM}$ (Upham et al, 2019) and BirdTree$^{TM}$ (Jetz et al, 2012) respectively. These tree-sets were drawn from pseudo-posterior distributions of models that used relaxed-clock methods to infer phylogeny from genetic and fossil data (mammals: $n_{Genes} = 31$ ; birds: $n_{Genes} = 15$; Jetz et al, 2012; Upham et al, 2019; avian fossil backbone: Ericson et al, 2006). From each tree-set, we then estimated class-specific concensus trees using 'TreeAnnotator' in Beast2$^{TM}$ (Bouckaert et al, 2014), with a 25% burn-in, posterior probability limit of 1.0, and with node heights representing median values. Next, consensus trees were used to build variance-covariance matrices that approximated the relative relatedness between species (here, using the R package "ape"; Paradis and Schliep, 2019). These variance-covariance matrices were later used to correct for phylogenetic relationships between species in sections 7-9.

```
# Saving species list to build phylogeny.

Species_List <- data.frame(Class = Data$Class, Species = Data$Species.Name) %>%
    distinct() %>%
    mutate(Species = word(Species, 1, 2)) %>%
    arrange(Class, Species)

write.csv(Species_List, "/Users/joshuatabh/Documents/researchProjects/trent/sihMetaregression/data/speciesList.csv",
    row.names = F)

# Reading in consensus trees derived from TREEANNOTATOR
# (provided with BEAST2).

Mammals_DNA <- read.nexus("/Users/joshuatabh/Documents/researchProjects/trent/sihMetaregression/trees/Beast_Trees/mammals.nex")
Birds_DNA <- read.nexus("/Users/joshuatabh/Documents/researchProjects/trent/sihMetaregression/trees/Beast_Trees/birds.nex")

# Quickly plotting trees, then building covariance matrices
# from trees for modelling.

MammalTree <- Mammals_DNA
MammalTree$tip.label <- gsub("_", " ", MammalTree$tip.label)

max(MammalTree$edge.length) + MammalTree$edge.length[7]
```

```
## [1] 84.15982
```

```
edge <- data.frame(MammalTree$edge, edge_num = 1:nrow(MammalTree$edge))
colnames(edge) <- c("parent", "node", "edge_num")

# Summing lengths of edges 1 and 17 will provide total
# time-frame.

MammalTree$edge.length[1] + MammalTree$edge.length[17]
```

```
## [1] 32.23683
```

```r
# Rounding to 81 for the purposes of a plot.

MammalTreePlot <- ggtree(MammalTree) + geom_nodepoint(pch = 21,
    alpha = 0.6, size = 4, colour = "black", fill = "lightseagreen") +
    geom_tiplab(size = 4) + theme_tree2() + scale_x_continuous(limits = c(-10,
    105), breaks = c(1, 21, 41, 61, 81), labels = c(80, 60, 40,
    20, 0)) + xlab("Millions of Years") + theme(text = element_text(family = "Noto Sans"),
    axis.title = element_text(size = 14))

print(MammalTreePlot)
```

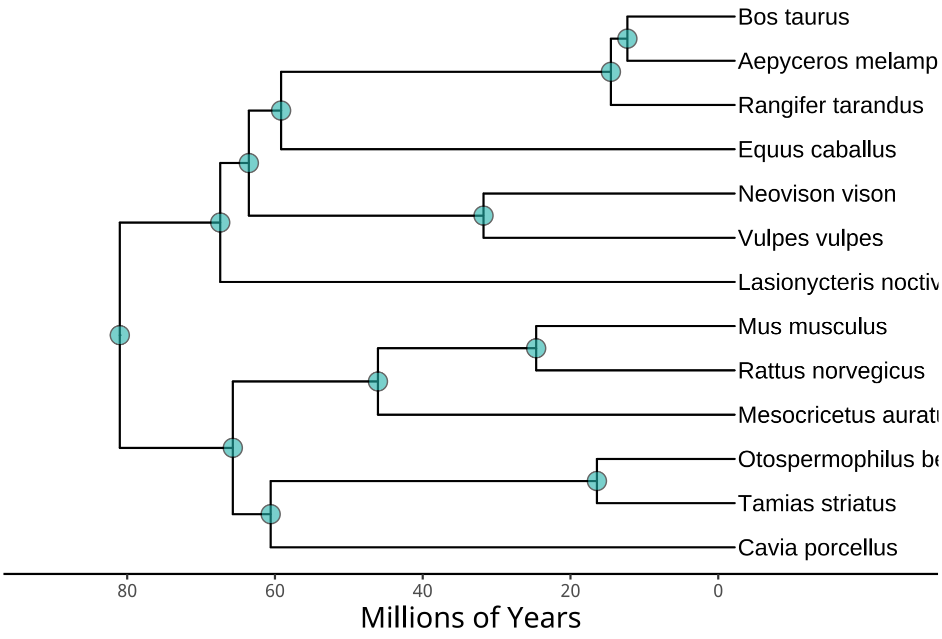


Figure 11: Consensus phylogenetic tree displaying relatedness of mammalian species included in our data-set, and estimates of time-lags between species divergences. Consensus trees were estimated using genetic data from 31 loci, as derived from VertTree (Upham et al, 2019). Details regarding tree construction are provided in the text above.

```r
ggsave("/Users/joshuatabh/Documents/researchProjects/trent/sihMetaregression/figures/mammalTree.jpg",
    MammalTreePlot, dpi = 800, width = 11, height = 7.5)
```

```r
BirdTree <- Birds_DNA
BirdTree$tip.label <- gsub("_", " ", BirdTree$tip.label)

max(BirdTree$edge.length)  # Extracting maximum edge length to determine range of x axis.
```

```
## [1] 87.21574
```

```r
BirdTreePlot <- ggtree(BirdTree) + geom_nodepoint(pch = 21, alpha = 0.5,
    size = 3, colour = "black", fill = "slateblue") + geom_tiplab(size = 3.5) +
    theme_tree2() + scale_x_continuous(limits = c(-20, 140),
    breaks = c(-15, 10, 35, 60, 85, 110), labels = c(125, 100,
        75, 50, 25, 0), ) + xlab("Millions of Years") + theme(text = element_text(family = "Noto Sans"),
    axis.title = element_text(size = 14))

print(BirdTreePlot)
```

```r
ggsave("/Users/joshuatabh/Documents/researchProjects/trent/sihMetaregression/figures/birdTree.jpg",
    BirdTreePlot, dpi = 800, width = 10.5, height = 7.5)

# Now building covariance matrix for model.

Mammals_VCV <- vcv.phylo(Mammals_DNA)
```

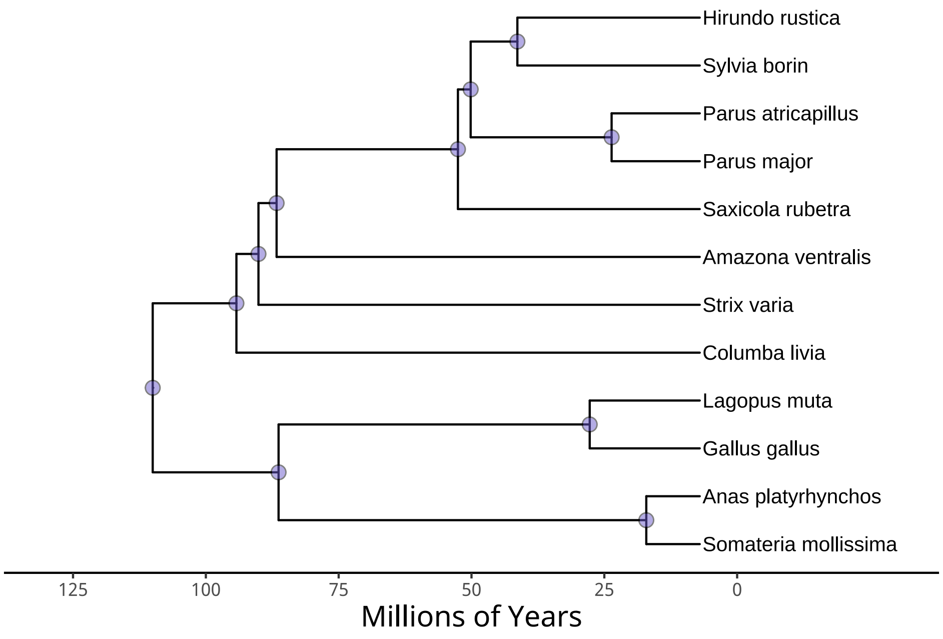


Figure 12: Consensus phylogenetic tree displaying relatedness of avian species included in our data-set, and estimates of time-lags between species divergences. Here, consensus trees were estimated using genetic data from 15 loci, as derived from BirdTree (Jetz et al, 2012). Again, details regarding tree construction are provided in the text above.

```
Birds_VCV <- vcv.phylo(Birds_DNA)

Bird_Tack <- data.frame(matrix(0, nrow = nrow(Mammals_VCV), ncol = ncol(Birds_VCV)))
colnames(Bird_Tack) <- colnames(Birds_VCV)
rownames(Bird_Tack) <- rownames(Mammals_VCV)

Mammal_Tack <- data.frame(matrix(0, nrow = nrow(Birds_VCV), ncol = ncol(Mammals_VCV)))
colnames(Mammal_Tack) <- colnames(Mammals_VCV)
rownames(Mammal_Tack) <- rownames(Birds_VCV)

Mammals_VCV <- cbind(Mammals_VCV, Bird_Tack)
Birds_VCV <- cbind(Mammal_Tack, Birds_VCV)

All_VCV <- rbind(Mammals_VCV, Birds_VCV)

# Correcting nomenclature.

colnames(All_VCV)[c(which(colnames(All_VCV) == "Parus_atricapillus"))] <- "Poecile_atricapillus"
rownames(All_VCV)[c(which(rownames(All_VCV) == "Parus_atricapillus"))] <- "Poecile_atricapillus"

# Lastly, adding phylogeny column in main data.

Data <- Data %>%
    mutate(Phylo = word(Species.Name, 1, 2)) %>%
    mutate(Phylo = gsub("[[:space:]]", "_", Phylo)) %>%
    mutate(Phylo = ifelse(Phylo == "Spermophilus_beecheyi", "Otospermophilus_beecheyi",
        ifelse(Phylo == "Mustela_vison", "Neovison_vison", Phylo)))

rm(Bird_Tack, Birds_DNA, Birds_VCV, BirdTree, BirdTreePlot, edge,
    Mammal_Tack, Mammals_DNA, Mammals_VCV, MammalTree, MammalTreePlot,
    TestView)
```

#### 6 | Modeling core body temperature responses to stress exposure

We were interested in testing whether the magnitude and direction of stress-induced changes in core body temperature are influenced by a species' thermoregulatory expenditure (that is, their energetic expenditure *toward* thermoregulation) and/or their relative energy available to expend toward body temperature maintenance and the stress response at all. Clearly, expenditure toward thermoregulation should vary across ambient temperature, with the degree of variation being determined by a species' body size, or mass. More specifically, a relatively small species is expected to display a more rapid increase in thermoregulatory expenditure across declining ambient temperatures than a relatively large species. Moreover, energy available to expend toward thermoregulation and the stress response should depend upon relative energy use at rest (see section 3). Accordingly, we tested whether ambient temperature, body mass, relative energy expenditure, and the interactions between body mass and ambient temperature, and relative energy expenditure and ambient temperature were correlated with the magnitude and direction of stress-induced changes in core body temperature (here, measured by a log-response ratio; refer to section 5). This was achieved by including each parameter as a population-level predictor in the below analysis. Importantly, however, the methodology of studies included in our final data set often varied in ways that could bias estimates of stress-induced changes in body temperature. For example, several studies monitored changes in body temperatures by intra-abdominal or intra-caecal telemetry, while others employed thermocouples inserted into the rectum, cloaca, or throat. Because measurement by each method is likely to be influenced by local vascular responses to a stressor, we might expect regular and systematic differences to occur between them. To control for these difference in methodology, we broadly categorised studies by body temperature measurement technique (binomial; internal telemetry or thermocouple) and included this category in our below analysis as a population-level predictor. Finally, because we could not discredit the possibility that estimates of stress-induced change in body temperature are explained by the time at which body temperature was measured (see Jerem et al, 2019), the time-length between baseline body temperature measurements and maximal or minimal body temperature measurements was also included as a population-level predictor in our analysis. Here, this time measurement was assumed modeled as a second-order polynomial to account for known non-linearity in stress-induced body temperature responses (again, see Jerem et al, 2019). As group-level predictors, we included species, study identity, and phylogeny (see section 6) to control for non-independence between species, observations reported in the same study, and between closely related species.

Individuals within a study population often vary considerably in size. Moreover, the ambient temperature at which a study population is observed is also rarely constant. These two sources of variation, if sufficiently large, may lead to incorrect predictions about the relationships between mass or ambient temperature and the magnitude and direction of stress-induced changes in body temperature. To capture and adjust for this uncertainty, we modeled body mass and ambient temperature as latent variables, with their true values being unknown, but falling within by our observed values ± our observed error.

For this analysis, we used weakly informative priors that assumed log-responses ratios $\lessapprox$ -1 (representing a 24°C decline for an animal with a baseline core body temperature of 38°C) and $\gtrapprox$ 0.5 (representing a 10°C increase for the same animal described above) to be unlikely. Specifically, priors for air temperature, body mass, and the first-order effects of latency to body temperature measurement were set as normally distributed with a mean of 0 and a standard deviation of 0.1, while those for the interaction between body mass and air temperature, and the interaction between relative energy expenditure and air temperature were set as normally distributed with a mean of 0 and a standard deviation of 0.05. Because we were less certain about the range of possible age-related and second-order time-related effects, priors for age class and the second-order effect of time were set as normally distributed with a mean of zero and a larger standard deviation of 0.25. Next, because both relative energy expenditure and measurement methodology were factorial, priors for these parameters was set as skew-normal with $\xi$ equaling 0, $\omega$ equaling 0.1, and $\alpha$ equaling -2.5. Here, a skew-normal distribution was used to capture the increased likelihood of observing negative response ratio values than positive response ratio values (an inherent property of natural logarithm transformation). Importantly, however, a skew-normal distribution was not assumed for priors of continuous predictors where absolute slope estimates were expected to be small. Finally, priors for measurement technique, all group-level predictors, and $\sigma$ were weak with flat (across $\mathbb{R}$), and gamma distributions (group-level predictors: $\alpha = 1.5$, $\beta = 1.0$; $\sigma$: $\alpha = 1$, $\beta = 1$), and our model was run across 4 HMC chains with 15,000 iterations, burn-ins of

2,500, and thinning values of 10.

```
# Renaming data to be more explicit.

Core_Data <- Data

# Note that some species will only have data for surface body temperature responses and not core
# body temperature responses. This means that we need to correct our phylogenetic
# variance-covariance matrix to only include species for which core body temperature responses
# are observed.

Core_VCV <- All_VCV[(colnames(All_VCV) %in% unique(Core_Data$Phylo)),
                    (colnames(All_VCV) %in% unique(Core_Data$Phylo))]

# Checking inclusion.

colnames(Core_VCV) %in% unique(Core_Data$Phylo)
```

```
##  [1] TRUE TRUE TRUE TRUE TRUE TRUE TRUE TRUE TRUE TRUE TRUE TRUE TRUE TRUE TRUE
## [16] TRUE TRUE TRUE TRUE TRUE TRUE TRUE TRUE TRUE TRUE
```

```
dimnames(Core_VCV)[[1]][!(dimnames(Core_VCV)[[1]] %in% unique(Core_Data$Phylo))]
```

```
## character(0)
```

```
# Good. Next, constructing Cleveland Dot-Plot to assess outliers.

ggplot(Core_Data, aes(x = 1:nrow(Core_Data), y = lRR)) +
  geom_point(size = 2, colour = "black", fill = "lightseagreen", alpha = 0.8) +
  annotate("rect",
    xmin = 1, xmax = nrow(Core_Data),
    ymin = mean(Core_Data$lRR) - 3.5 * sd(Core_Data$lRR),
    ymax = mean(Core_Data$lRR) + 3.5 * sd(Core_Data$lRR),
    colour = "black", fill = "grey70", alpha = 0.4
  ) + xlab("Sample Number") + ylab("Log Response Ratio") +
  annotate("text", x = 10, y = mean(Core_Data$lRR) - 3.5 * sd(Core_Data$lRR) - 0.05,
           label = paste0("LL: ", round(mean(Core_Data$lRR) - 3.5 *
                                          sd(Core_Data$lRR), digits = 3))) +
  annotate("text", x = 10, y = mean(Core_Data$lRR) + 3.5 * sd(Core_Data$lRR) + 0.05,
           label = paste0("UL: ", round(mean(Core_Data$lRR) + 3.5 *
                                          sd(Core_Data$lRR), digits = 3))) +
  theme_classic()
```

```
# At least two clear outliers. Viewing and removing.

caption = "Observations with log response ratio (lRR) values falling outside of the mean +/- 3.5 times the standard deviation."

Core_Data[c(which(Core_Data$lRR < -0.33 | Core_Data$lRR > 0.33)), ] %>%
  select("Study ID" = StudyID, "Observation ID" = ObservationID,
         "Latin Name" = Species.Name,
         Mass, "Change in Body Temperature" = Delta.Tb) %>%
  kbl(., longtable = T, booktabs = T, caption = caption) %>%
  kable_styling(latex_options = "striped")
```

Table 11: Observations with log response ratio (lRR) values falling outside of the mean +/- 3.5 times the standard deviation.

| | Study ID | Observation ID | Latin Name | Mass | Change in Body Temperature |
|---|---|---|---|---|---|
| 11 | U | CE | Lasionycteris noctivagans | 9.47 | 10.80000 |
| 18 | A2 | FN | Rattus norvegicus domestica | 242.50 | -10.72740 |
| 22 | A2 | FR | Rattus norvegicus domestica | 304.00 | -11.05358 |
| 128 | K | AV | Cavia porcellus | 700.00 | -18.72300 |

```
# Four apparent.

Core_Data_Short <- Core_Data[-c(which(Core_Data$lRR < -0.33 | Core_Data$lRR > 0.33)), ]
```

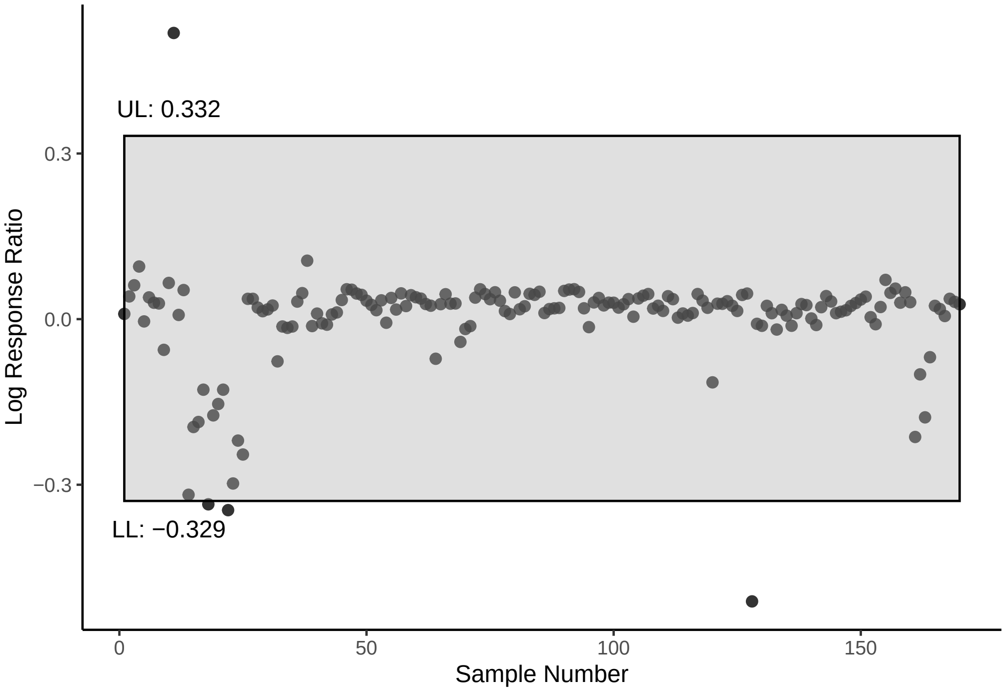


Figure 13: Raw response ratio values (log response ratios, or 'lRR') by data-set sample number. The light grey box represents that span of lRR values covered by the mean plus/minus 3.5 times the standard devation. Values falling outside of this box may be considered potential outliers.

```
Core_VCV_Short <- All_VCV[(colnames(All_VCV) %in% unique(Core_Data_Short$Phylo)),
                          (colnames(All_VCV) %in% unique(Core_Data_Short$Phylo))]

# Lastly, categorising measurement techniques and binning residual metabolic rate measurements as negative or positive.

Core_Data_Short$Technique <- "Probe"
Core_Data_Short$Technique[c(grep("Telemetry", Core_Data_Short$Measurement.Technique))] <- "Telemetry"
Core_Data_Short$Technique[c(grep("Throat.Probe", Core_Data_Short$Measurement.Technique))] <- "Probe"

Core_Data_Short = Core_Data_Short %>%
  mutate(Technique = factor(Technique))

# Mean-centreing all variables to ease interpretation of model coefficients

Core_Data_MR = Core_Data_Short %>%
  mutate("Centred_AT" = Ambient.Temp.Stressor - mean(Ambient.Temp.Stressor, na.rm = T),
         "Centred_logMass" = logMass - mean(logMass, na.rm = T),
         "Centred_Time" = Time.to.Peak.s. - mean(Time.to.Peak.s., na.rm = T)) %>%
  mutate("Age_Class" = ifelse(Age == "Juvenile", -1,
                              ifelse(Age == "Mixed", 0,
                                     ifelse(Age == "Adult", 1, NA)))
         ) %>%
  mutate("ResMR_Bin" = ifelse(ResMR_Bin == "Low", -0.5,
                              ifelse(ResMR_Bin == "High", 0.5, NA)
                              )
         ) %>%
  mutate("Technique" = ifelse(Technique == "Probe", -0.5,
                              ifelse(Technique == "Telemetry", 0.5, NA)
                              )
         ) %>%
  filter(!is.na(Age_Class) & !is.na(ResMR_Bin)) %>%
  mutate(Age_Class = as.integer(Age_Class))
```

```
Core_VCV_MR <- All_VCV[(colnames(All_VCV) %in% unique(Core_Data_MR$Phylo)),
                          (colnames(All_VCV) %in% unique(Core_Data_MR$Phylo))]

# Setting up prior distributions and running model

prior_Core <- c(
  set_prior("normal(0, 0.5)", class = "Intercept", resp = "lRR"),
  set_prior("normal(0, 0.1)", class = "b", coef = "miCentred_AT", resp = "lRR"),
  set_prior("normal(0, 0.1)", class = "b", coef = "miCentred_logMass", resp = "lRR"),
  set_prior("normal(0, 0.05)", class = "b", coef = "miCentred_AT:ResMR_Bin", resp = "lRR"),
  set_prior("normal(0, 0.1)", class = "b", coef = "Age_Class", resp = "lRR"),
  set_prior("skew_normal(0, 0.1, -2.1)", class = "b", coef = "Technique", resp = "lRR"),
  set_prior("skew_normal(0, 0.1, -2.1)", class = "b", coef = "ResMR_Bin", resp = "lRR"),
  set_prior("normal(0, 0.05)", class = "b", coef = "polyCentred_Time21", resp = "lRR"),
  set_prior("normal(0, 0.1)", class = "b", coef = "polyCentred_Time22", resp = "lRR"),
  set_prior("exponential(7.5)", class = "sd", resp = "lRR"),
  set_prior("gamma(1,2)", class = "sigma", resp = "lRR")
  )

modelSource <- paste0(
  "/Users/joshuatabh/Documents/researchProjects/trent/sihMetaregression/models/",
  "modelCore.Rds"
)

Core_Model <- brm(bf(Centred_logMass | mi(logMass_SD) ~ 0,
                     family = "gaussian") +
                    bf(Centred_AT | mi(Ambient.Temp.Stressor.sd) ~ 0,
                      family = "gaussian") +
                    bf(lRR | se(lRV, sigma = TRUE) ~ mi(Centred_AT)*mi(Centred_logMass) +
                           ResMR_Bin + ResMR_Bin:mi(Centred_AT) + Age_Class + Technique +
                           poly(Centred_Time, 2) + (1 | StudyID) + (1 | Species.Name) +
                         (1 | gr(Phylo, cov = A)), family = "gaussian") +
                    set_rescor(FALSE),
data = Core_Data_MR,
data2 = list(A = Core_VCV_MR),
cores = 4, chains = 4,
seed = 100, #refresh = 0,
iter = 50000, warmup = 25000, thin = 10,
prior = prior_Core,
save_pars = save_pars(latent = TRUE),
file = modelSource
)

rm(modelSource)
```

A handful of divergent transitions, but less than 5% of samples (after thinning). Nevertheless, we checked for any sampling biases or pathologies by visually assessing the distributions of our HMC draws across pairs of predictors. Furthermore, we visually checked for sequential non-independence in our HMC samples (i.e autocorrelation), and by assessing the ratio of our parameter-specific effective samples sizes to true sample size ratios. To validate convergence of our 4 HMC chains, we then calculated a Gelman-Rubin (Rhat) statistic, where 1 represents complete chain convergence. Finally, we then proceeded to check whether our model met basic assumptions (e.g. normality and homogeneity of error, linearity, etc.) by visually diagnosing residual distributions.

```
clean_pairs(Core_Model, prs = c("b_lRR_Intercept", "b_lRR_Technique",
    "b_lRR_ResMR_Bin", "b_lRR_Age_Class", "bsp_lRR_miCentred_AT",
    "bsp_lRR_miCentred_logMass"), names = c("Intercept", "Technique",
    "Residual MR", "Age Class", "Amb Temp", "Mass"))
```

```
clean_pairs(Core_Model, prs = c("bsp_lRR_miCentred_AT:miCentred_logMass",
    "bsp_lRR_miCentred_AT:ResMR_Bin", "sigma_lRR"), names = c("log Mass^2 by °C",
    "Residual MR by °C", "Sigma"))
```

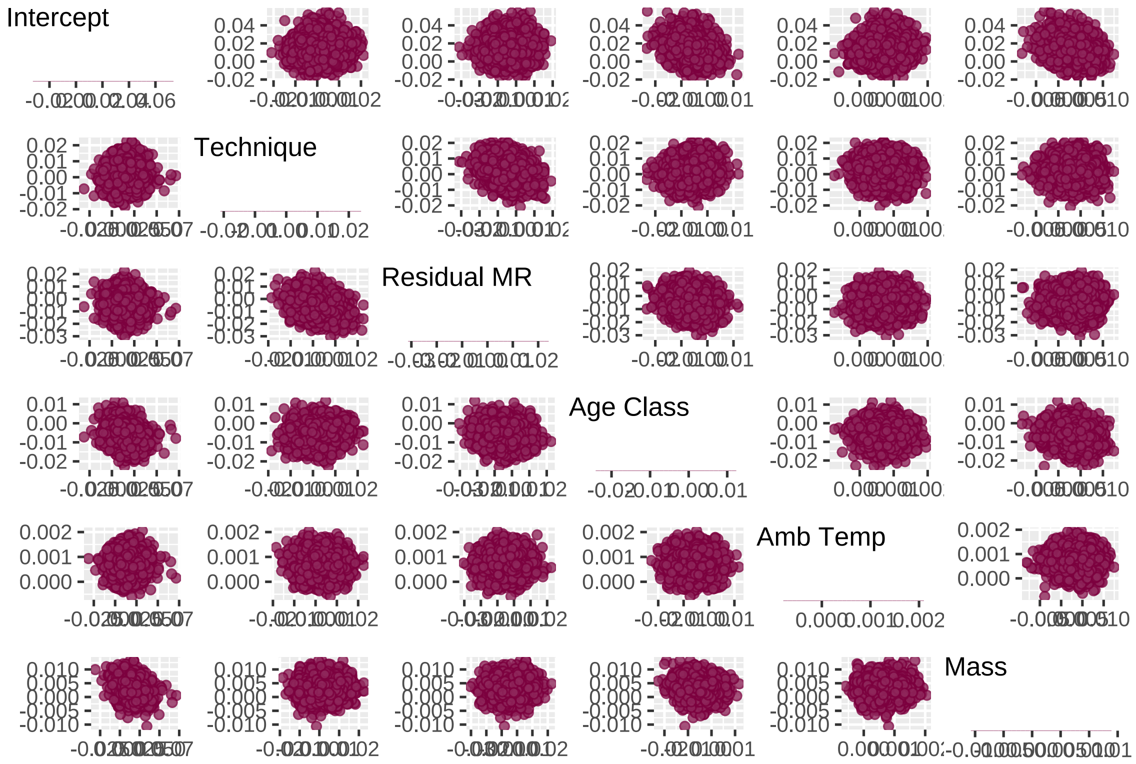


Figure 14: Paired, posterior coefficient estimates from a Bayesian, linear model predicting the magnitude and direction of stress-induced changes in body temperature (measured as log response ratios). Note that only select parameter pairs are displayed here, with the remainder presented in the plot immediately below.

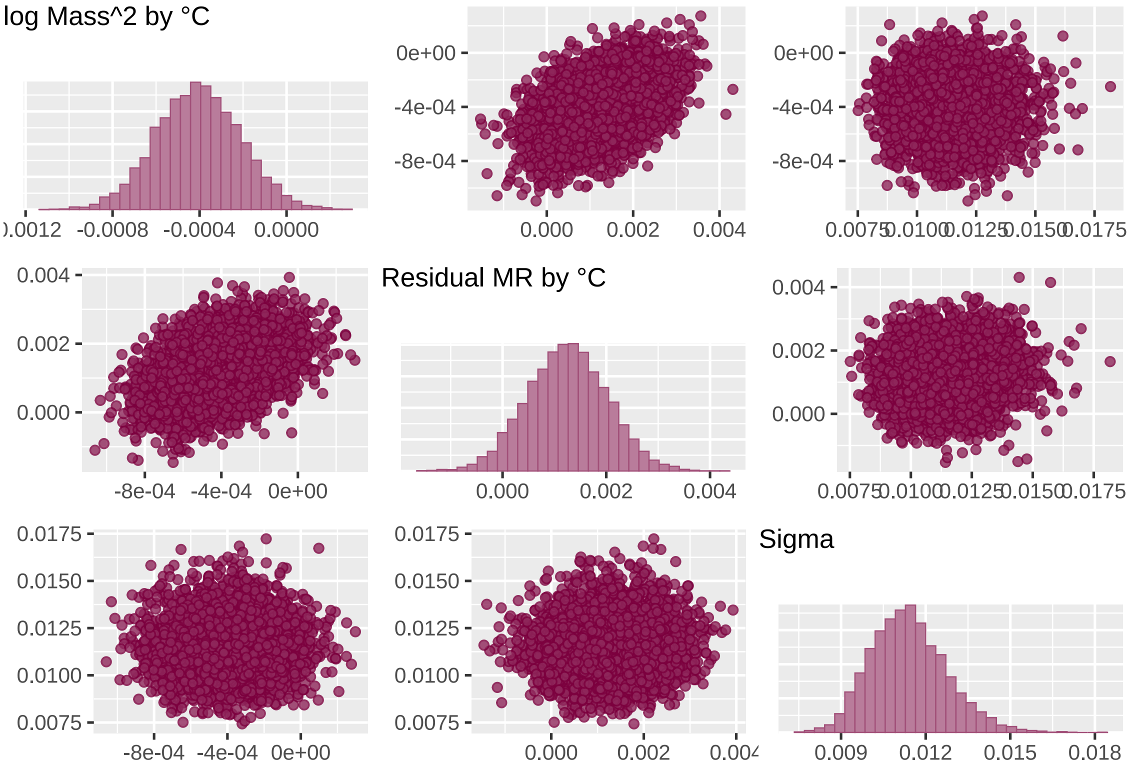


Figure 15: Paired, posterior coefficient estimates from a Bayesian, linear model predicting the magnitude and direction of stress-induced changes in body temperature (measured as log response ratios). Parameter pairs not displayed in the above plot are displayed here.

```
# Great. Mild correlations between posteriors where
# correlations are expected (that is, given the inclusion
# of interaction terms). Otherwise, all appears well
# dispersed, except for a potentially curious correlation
# between time and age class. Assessing this in raw data.

Core_Model$data %>%
    left_join(Core_Model$data %>%
        group_by(Age_Class) %>%
        count() %>%
        mutate(Label = paste0(Age_Class, "\n(n = ", n, ")")),
        by = "Age_Class") %>%
    ggplot(aes(x = Label, y = Centred_Time, fill = Label)) +
    geom_point(size = 2, pch = 21, colour = "black", alpha = 0.5,
        position = position_jitter(width = 0.25)) + stat_summary(geom = "errorbar",
    fun.data = "mean_se", width = 0.2, colour = "black") + stat_summary(geom = "point",
    fun = "mean", size = 4, pch = 21, colour = "black") + theme_classic() +
    scale_fill_manual(values = c(nice_pink, "black", "grey80")) +
    theme(legend.position = "none", axis.title = element_text(size = 12,
        family = "Noto Sans"), axis.text.x = element_text(size = 10)) +
    xlab("Age Class") + ylab("Time To Peak/Trough Tb\n(S; Mean-Centred)")
```

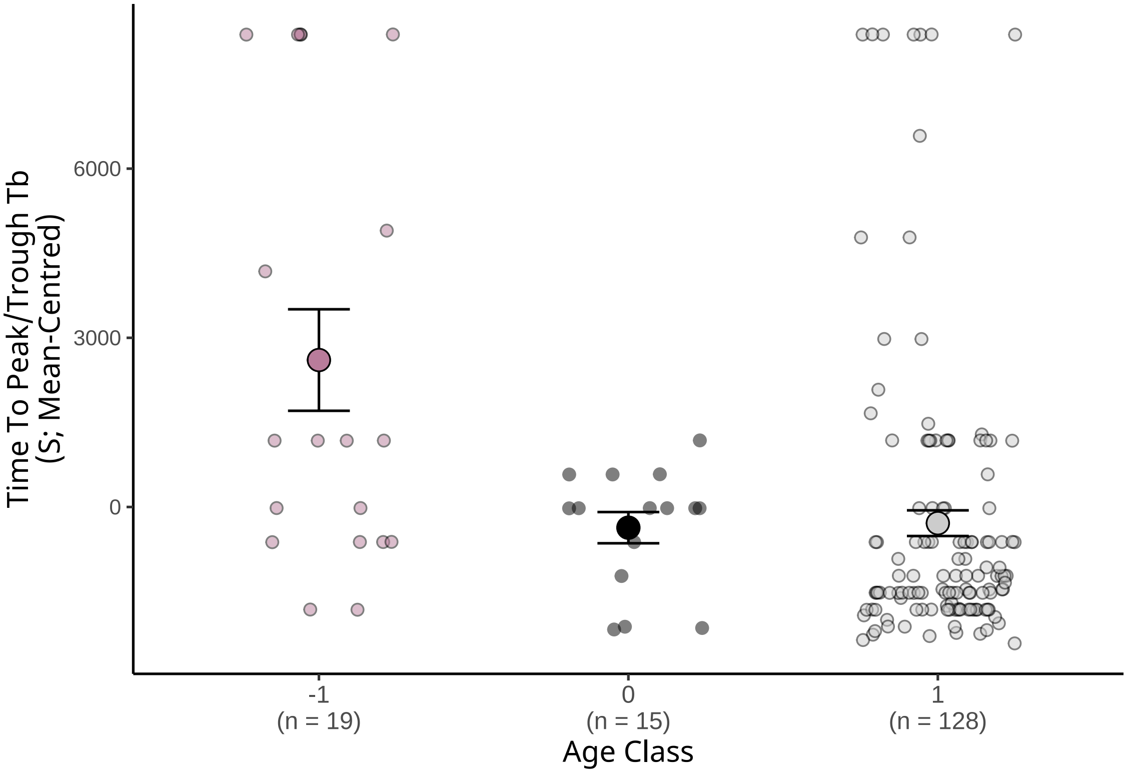


Figure 16: Effect of age class (adult, juvenile, or both) on the latency from stress exposure to a maximum or minimum change in body temperature across bird and mammal species. Each small dot represents a single obervation for a single species (obtained from one study). Large dots represent means for each age class grouping and errorbars represent standard errors around means.

```
# Potentially some distinctions here, however, they appear
# to be a product of differences in sample sizes

clean_ac(Core_Model, prs = c("b_lRR_Intercept", "b_lRR_Technique",
    "b_lRR_ResMR_Bin", "b_lRR_Age_Class", "bsp_lRR_miCentred_AT",
    "bsp_lRR_miCentred_logMass"), names = c("Intercept", "Technique",
    "Residual MR", "Age Class", "Amb Temp", "Mass"))
```

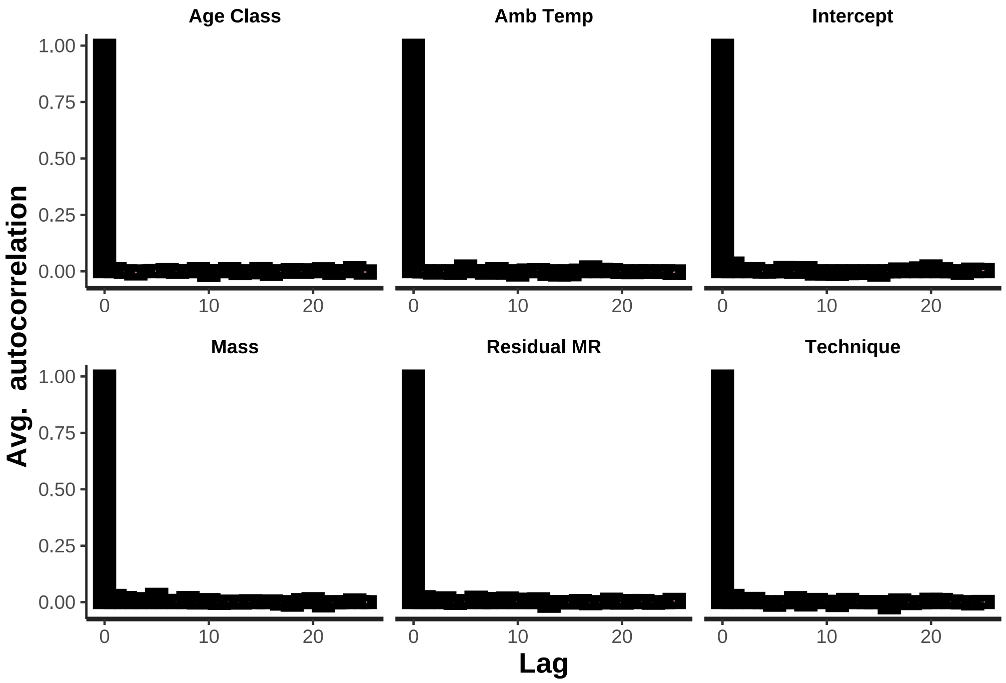


Figure 17: Degree of sequential autocorrelation between chain draws, per parameter, for a Bayesian, linear mixed effects model predicting stress-induced changes in body temperature (natural log-transformed response ratio). Autocorrelation for select predictors is displayed, with that for further predictors displayed in the plot below.

```
clean_ac(Core_Model, prs = c("bsp_lRR_miCentred_AT:miCentred_logMass",
    "bsp_lRR_miCentred_AT:ResMR_Bin", "sigma_lRR"), names = c("log Mass^2 by °C",
    "Residual MR by °C", "Sigma"))
```

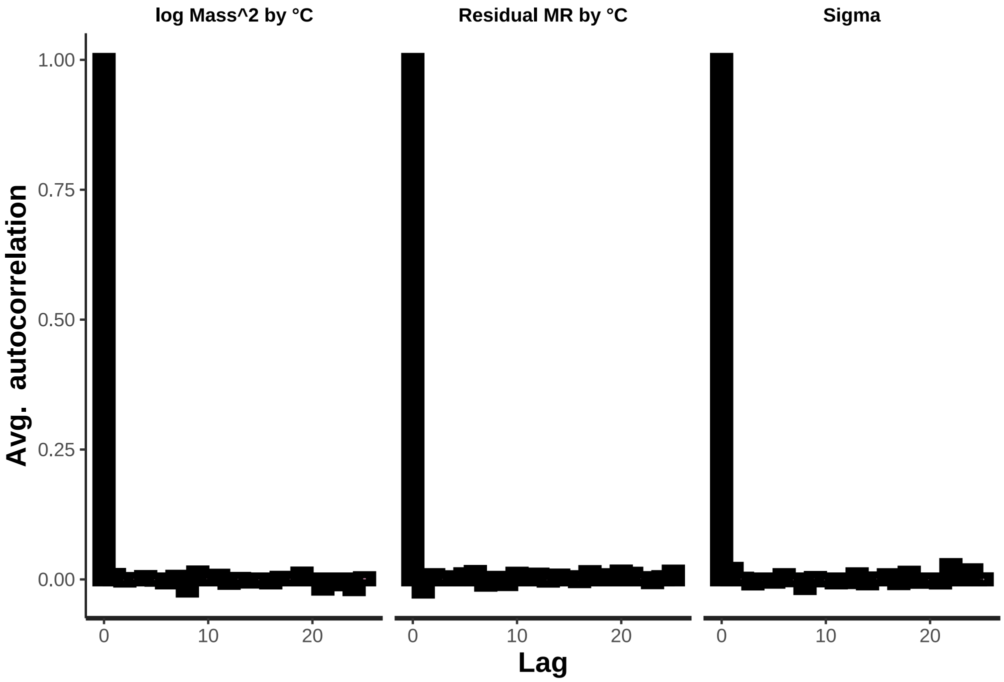


Figure 18: Degree of sequential autocorrelation between chain draws, per parameter, for a Bayesian, linear mixed effects model predicting stress-induced changes in body temperature (natural log-transformed response ratio, lRR'). Autocorrelation for select predictors is displayed; further predictors are displayed in the plot above.

```
mcmc_neff(neff_ratio(Core_Model), size = 2)
```

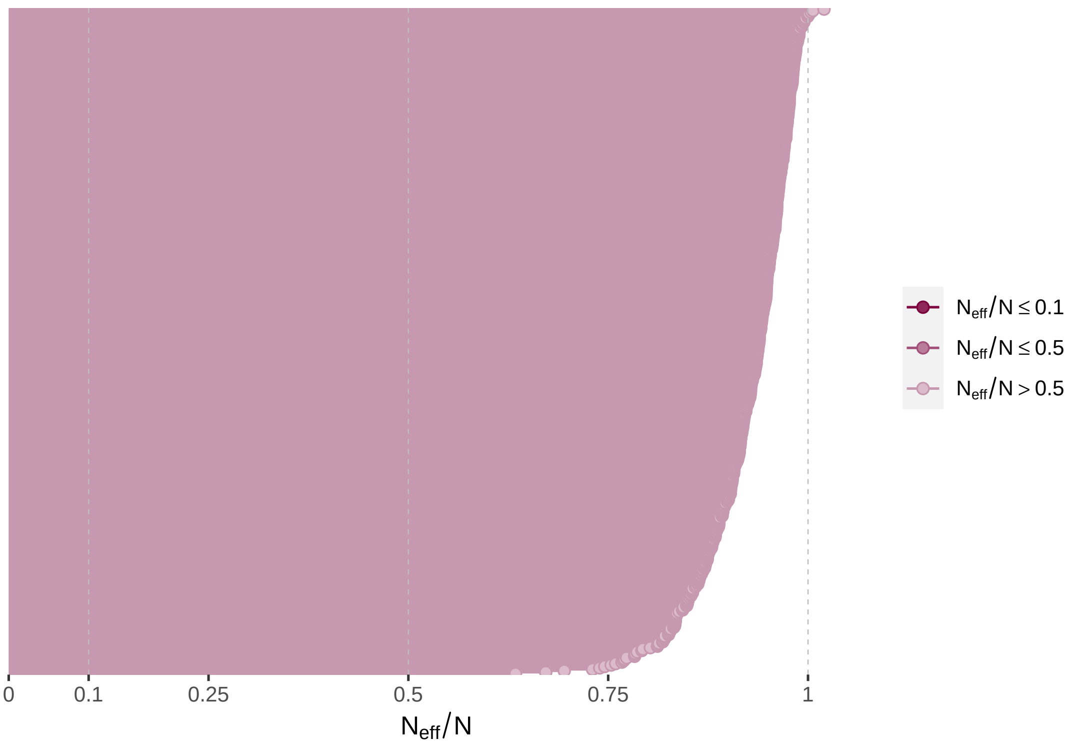


Figure 19: Ratio of effective sample sizes to sample sizes for a Bayesian, linear mixed effects model predicting stress-induced changes in body temperature (natural log-transformed response ratio, lRR').

```
# No apparent autocorrelation and ratio of effective sample
# sizes to sample sizes appear acceptable. Checking Rhat
# values.

mcmc_rhat(rhat(Core_Model))
```

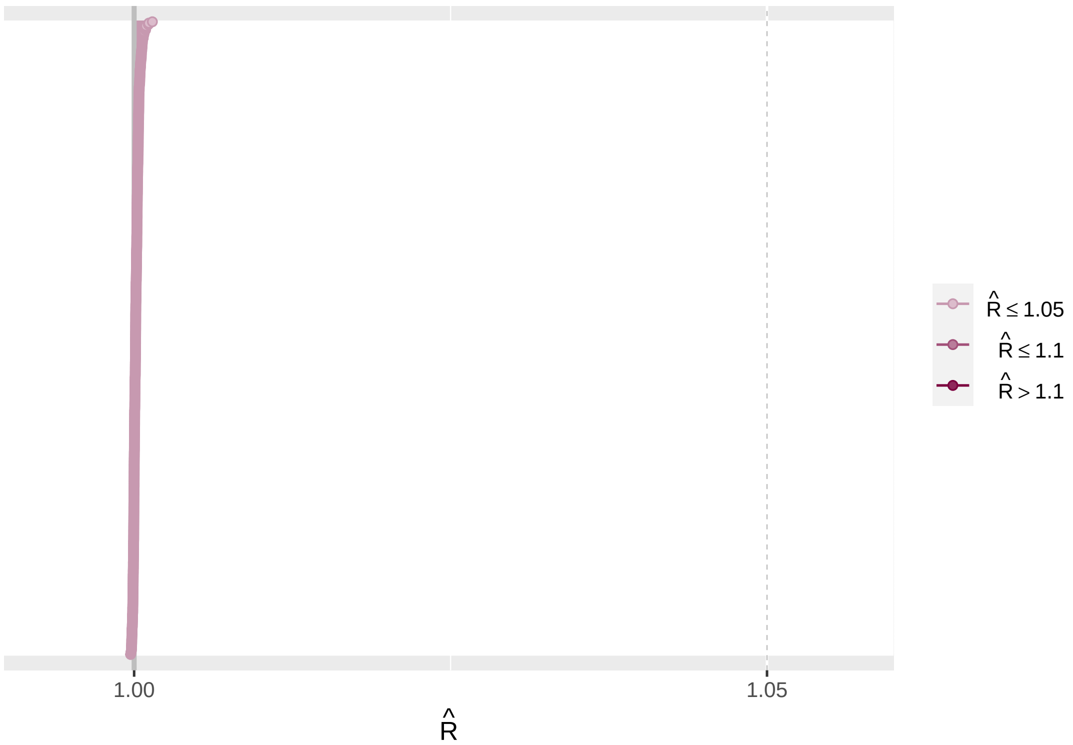


Figure 20: Gelman-Rubin statistics per model parameter for a Bayesian, linear mixed effects model predicting stress-induced changes in body temperature (natural log-transformed response ratio, lRR').

```
# Rhat values quite tight to 1, suggesting nice chain
# mixing. Checking residuals next.
```

```
Core_Model$data %>%
    mutate(Res = residuals(Core_Model, type = "ordinary", method = "posterior_predict")[,
        "Estimate", "lRR"]) %>%
    ggplot(aes(x = 1:nrow(.), y = Res)) + geom_point() + theme_classic() +
    xlab("Sample Number") + ylab("Ordinary Residuals")
```

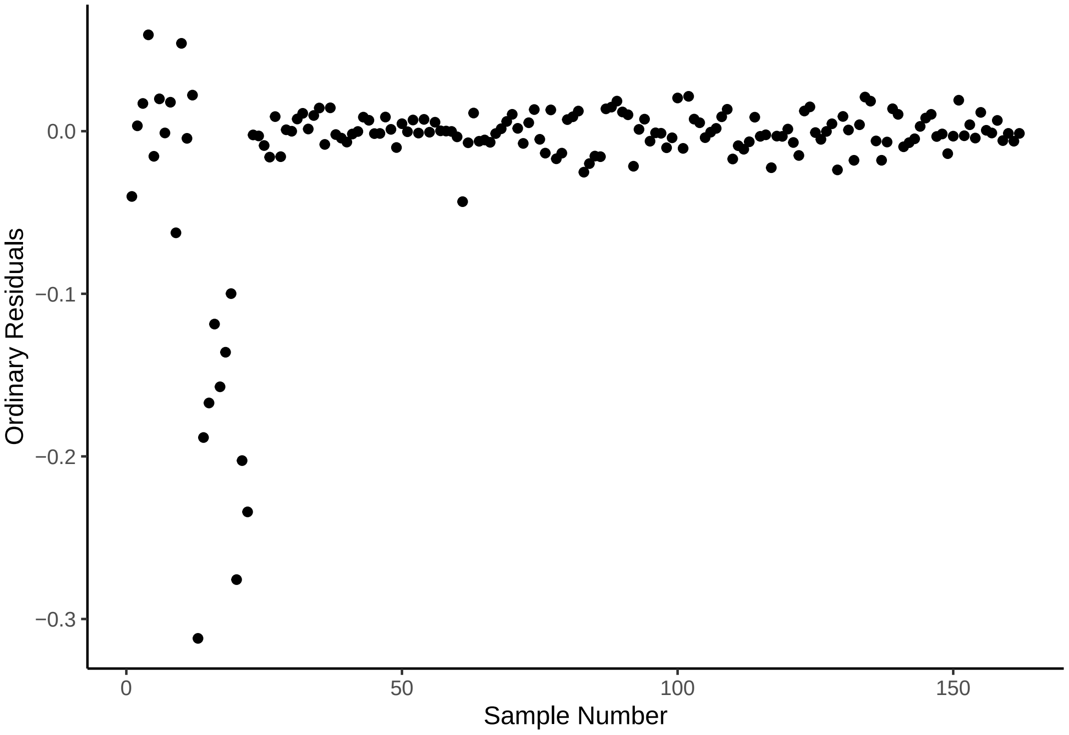


Figure 21: Ordinary, untransformed residuals for a Bayesian, linear mixed effects model predicting stress-induced changes in body temperature (natural log-transformed response ratio, lRR'), diplayed per data-frame row. Each dot represents the residual value for a single observation of a stress-induced change in body temperature for a single species.

```
# Some extreme values appear to be pulling downward.
# Assessing which.

caption = "Data-points where ordinary residuals fall below -0.05."
Core_Model$data %>%
    mutate(Res = residuals(Core_Model, type = "ordinary", method = "posterior_predict")[,
        "Estimate", "lRR"]) %>%
    filter(Res < -0.05) %>%
    select(`Study ID` = StudyID, `Latin Name` = Species.Name,
        `Ambient Temperature` = Centred_AT, `log Mass` = Centred_logMass,
        `log Response Ratio` = lRR, `Ordinary Residuals` = Res) %>%
    kbl(., longtable = T, booktabs = T, caption = caption) %>%
    kable_styling(latex_options = "striped")
```

Table 12: Data-points where ordinary residuals fall below -0.05.

| Study ID | Latin Name | Ambient Temperature | log Mass | log Response Ratio | Ordinary Residuals |
|---|---|---|---|---|---|
| U | Lasionycteris noctivagans | -3.038916 | -3.0603060 | -0.0556798 | -0.0547019 |
| A2 | Rattus norvegicus domestica | -8.538916 | -0.1387556 | -0.3181413 | -0.3070007 |
| A2 | Rattus norvegicus domestica | -8.538916 | -0.1387556 | -0.1954068 | -0.1886799 |
| A2 | Rattus norvegicus domestica | -8.538916 | -0.1387556 | -0.1863296 | -0.1837381 |
| A2 | Rattus norvegicus domestica | -8.538916 | -0.1387556 | -0.1278334 | -0.1091115 |
| A2 | Rattus norvegicus domestica | -8.538916 | 0.0869686 | -0.1743534 | -0.1499253 |
| A2 | Rattus norvegicus domestica | -8.538916 | 0.0869686 | -0.1537341 | -0.1476879 |
| A2 | Rattus norvegicus domestica | -8.538916 | 0.0869686 | -0.1278334 | -0.1064464 |
| A2 | Rattus norvegicus domestica | -8.538916 | 0.3129946 | -0.2977324 | -0.2792959 |
| A2 | Rattus norvegicus domestica | -8.538916 | 0.3129946 | -0.2200234 | -0.1942354 |
| A2 | Rattus norvegicus domestica | -8.538916 | 0.3129946 | -0.2452614 | -0.2198452 |

```
# Nearly all points from the same study. Checking where
# these values fall with respect to the mean and standard
# deviation.

Core_Model$data %>%
    mutate(Res = residuals(Core_Model, type = "ordinary", method = "posterior_predict")[,
        "Estimate", "lRR"]) %>%
    summarise(`Mean - 3.5*SD` = mean(Res) - 3.5 * sd(Res), `Mean + 3.5*SD` = mean(Res) +
        3.5 * sd(Res))
```

```
##   Mean - 3.5*SD Mean + 3.5*SD
## 1    -0.1887087     0.1653175
```

```
# Several values fall outside of bounds. Removing this
# study and re-running model.
```

Clearly, the log response ratio (representing the magnitude of a stress-induced change in body temperature) of some observations laid outside of our general error and may be biasing our model. To ensure that these samples did not bias our results, we removed them from our data-set and reconstructed our model. All model assessments were then repeated as described above.

```
rm(Core_Model)

Core_Data_MR_Shortened = Core_Data_MR  %>%
  filter(StudyID != "A2")

modelSource <- paste0(
 "/Users/joshuatabh/Documents/researchProjects/trent/sihMetaregression/models/",
 "modelCoreCorrected.Rds"
)

Core_Model_New <- brm(bf(Centred_logMass | mi(logMass_SD) ~ 0,
                     family = "gaussian") +
```

```
                    bf(Centred_AT | mi(Ambient.Temp.Stressor.sd) ~ 0,
                       family = "gaussian") +
                    bf(lRR | se(lRV, sigma = TRUE) ~ mi(Centred_AT)*mi(Centred_logMass) +
                           ResMR_Bin + ResMR_Bin:mi(Centred_AT) + Age_Class + Technique +
                           poly(Centred_Time, 2) + (1 | StudyID) + (1 | Species.Name) +
                         (1 | gr(Phylo, cov = A)), family = "gaussian") +
                    set_rescor(FALSE),
data = Core_Data_MR_Shortened,
data2 = list(A = Core_VCV_MR),
cores = 4, chains = 4,
seed = 100, #refresh = 0,
iter = 50000, warmup = 25000, thin = 10,
prior = prior_Core,
save_pars = save_pars(latent = TRUE),
file = modelSource
)
```

```
# Quickly looking at posterior estimates of log response
# ratio.

pp_check(Core_Model_New, resp = "lRR", ndraws = 500) + ylab("Density") +
    xlab("log Response Ratio")
```

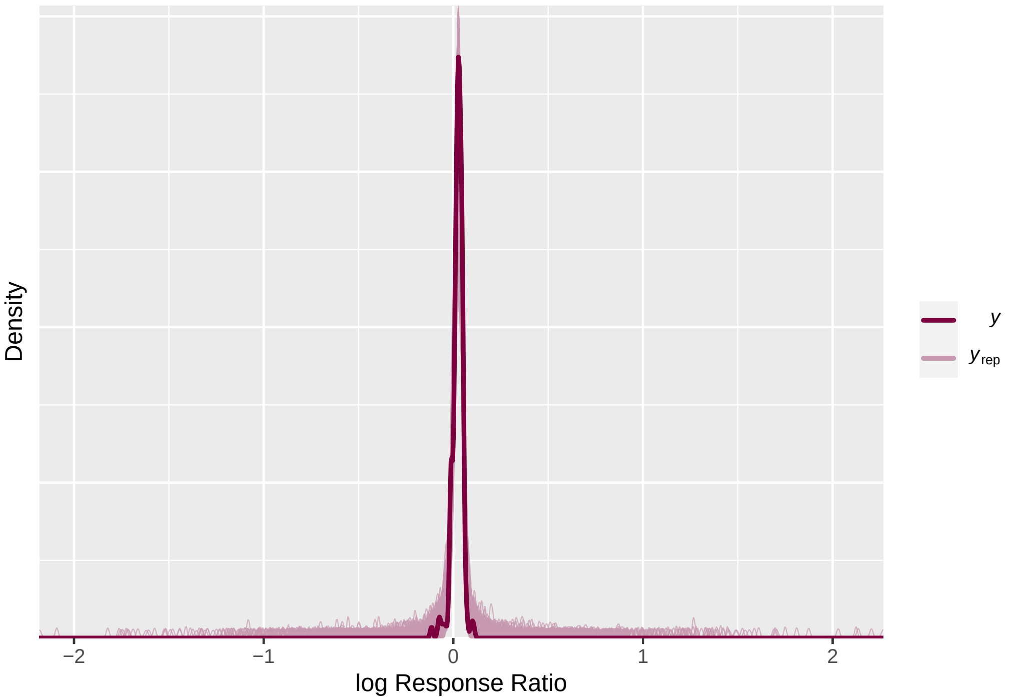


Figure 22: Density of stress-induced change in body temperature values (natural log-transformed response ratios) observed in raw data (maroon) and predicted by a Bayesian linear mixed effects model (light pink).

```
# Okay, but slightly tight near 0. Comparing posterior
# predictions across classes, residual metabolic rate bins,
# and across ambient temperature and mass.

yrep <- posterior_predict(Core_Model_New, ndraws = 100)

y <- Core_Model_New$data %>%
    pull(lRR)

g(Class_Group, Temp_Group, Mass_Group, ResMR_Bin) %=% (List <- Core_Data_MR_Shortened %>%
    mutate(Temp_Group = ifelse(Ambient.Temp.Stressor < 15, "Cold",
        ifelse(Ambient.Temp.Stressor >= 15 & Ambient.Temp.Stressor <
```

```
            25, "Warm", "Hot")), Mass_Group = ifelse(Mass < 100,
        "Small", ifelse(Mass >= 100 & Mass < 2000, "Mid", "Large"))) %>%
    select(Class, Temp_Group, Mass_Group, ResMR_Bin) %>%
    as.list(.))

g(pp1, pp2, pp3, pp4, pp5, pp6, pp7) %=% list(ppc_violin_grouped(y,
    yrep[, , "lRR"], group = Class_Group, probs = c(0.05, 0.95),
    alpha = 0.05, y_draw = "points"), ppc_violin_grouped(y, yrep[,
    , "lRR"], group = Temp_Group, probs = c(0.05, 0.95), alpha = 0.05,
    y_draw = "points"), ppc_violin_grouped(y, yrep[, , "lRR"],
    group = Mass_Group, probs = c(0.05, 0.95), alpha = 0.05,
    y_draw = "points"), ppc_violin_grouped(y, yrep[, , "lRR"],
    group = ResMR_Bin, probs = c(0.05, 0.95), alpha = 0.05, y_draw = "points"),
    ppc_scatter_avg_grouped(y, yrep[, , "lRR"], group = Class_Group) +
        geom_smooth(method = "lm", colour = "black"), ppc_scatter_avg_grouped(y,
        yrep[, , "lRR"], group = Temp_Group) + geom_smooth(method = "lm",
        colour = "black"), ppc_scatter_avg_grouped(y, yrep[,
        , "lRR"], group = Mass_Group) + geom_smooth(method = "lm",
        colour = "black"))
```

```
grid.arrange(pp1, pp2, pp3, pp4, nrow = 2)
```

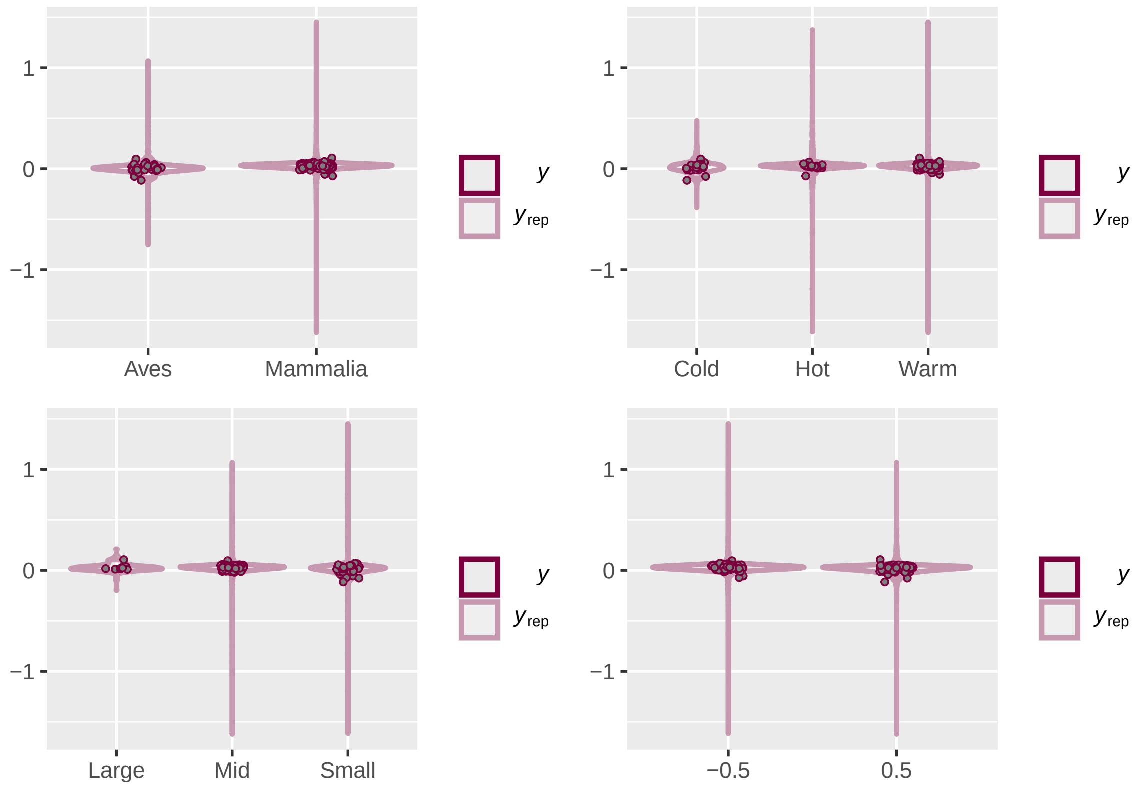


Figure 23: Distribution of stress-induced change in body temperature measurements (natural log response ratios) across classes (top left), temperature classes (top right; cold = less than 15°C, warm = >15°C and <25°C, hot = >25°C), body masses (bottom left; small = < 100 g, mid = >100 g and < 2000 g, large = >2000 g), and relative metabolic expenditures (bottom right; low = -0.5, high = 0.5). Dots represent individual change measurements, per study and species, and violins indicate ranges in predicted change values (y limits) and their distributions (width of violin), as derived from a Bayesian linear mixed effects model.

```
grid.arrange(pp5, pp6, pp7, nrow = 3)
```

```
# Little concern across groupings. Removing unnecessary
# objects and checking model chains.
```

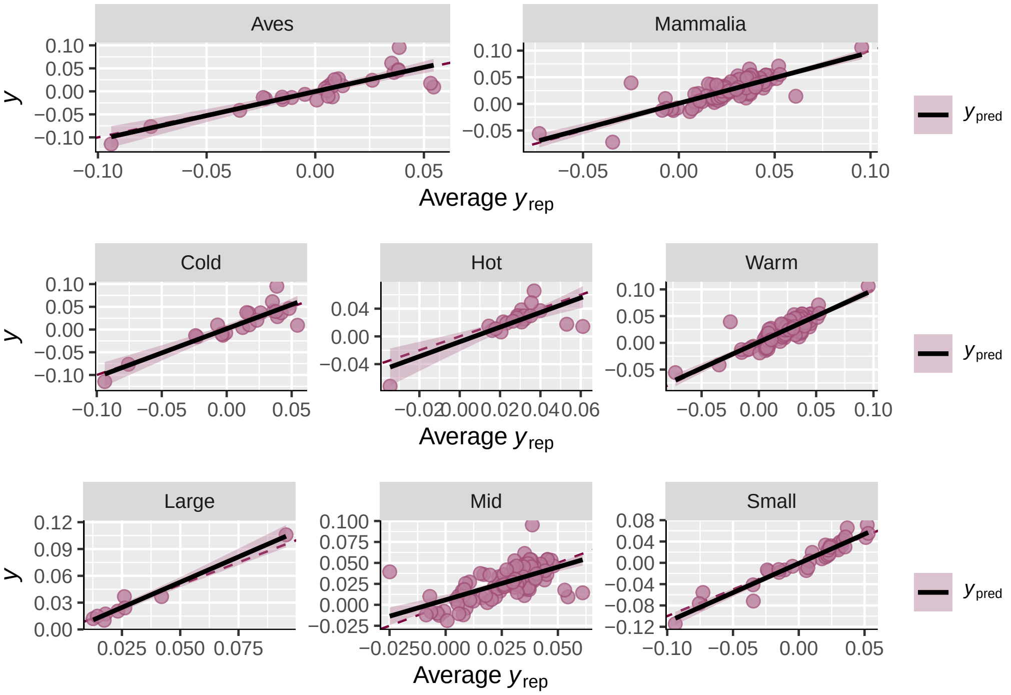


Figure 24: Spread of stress-induced change in body temperature measurements (natural log response ratios) across classes (top two panels), temperature classes (middle three panels; cold = <15°C, warm = >15°C and <25°C, hot = >25°C) and body masses (lower three panels; small = < 100 g, mid = >100 g and < 2000 g, large = >2000 g). Dots represent individual change measurements, per study and species (y-axis), as correlated with their their predicted change values (x-axis), via a Bayesian linear mixed effects models. Lines represent lines of best fit, as estimated for a simple linear correlation, and grey bands indicate 95 percent credible intervales around these trend lines.

```
rm(Class_Group, Temp_Group, Mass_Group, ResMR_Bin, pp1, pp2,
    pp3, pp4, pp5, pp6, pp7)
```

```
lp <- log_posterior(Core_Model_New)
nuts <- nuts_params(Core_Model_New)

MC_Div <- mcmc_nuts_divergence(nuts, lp, chain = 4)[1][["grobs"]][[1]]
Ylab <- ggplotGrob(ggplot() + ylab("Log Posterior Density"))[["grobs"]][[13]]
MC_Div[["grobs"]][[13]] <- Ylab
MC_Div <- as.ggplot(MC_Div)

print(MC_Div)
```

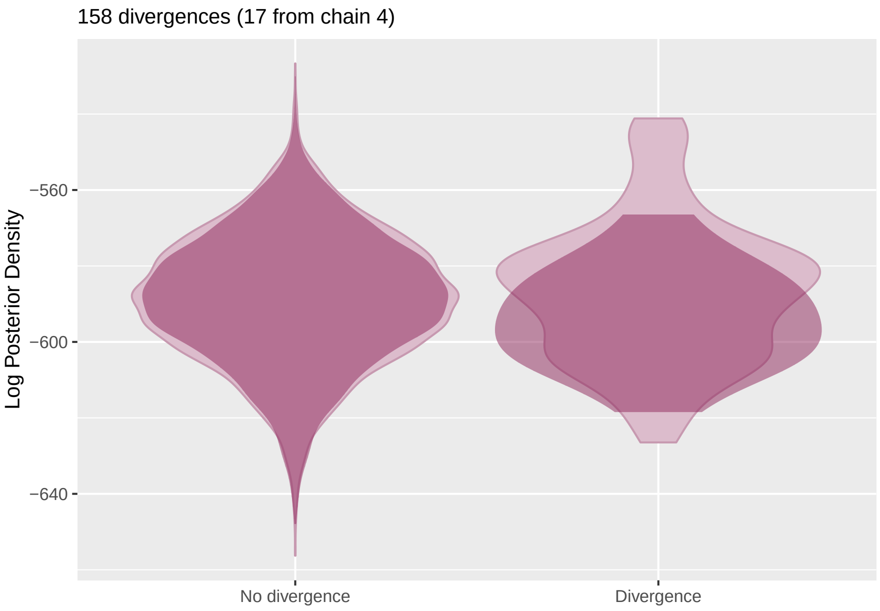


Figure 25: Natural log posterior densities for chain draws with and without divergences, as derived from a Bayesian linear mixed effects model predicted stress-induced changes in body temperature (natural log-transformed response ratio).

```
# No divergences or obvious constrictions in the log
# posterior density. Assessing autocorrelation in chains.
rm(lp, nuts, MC_Div, Ylab)
```

```
clean_ac(Core_Model_New, prs = c("bsp_lRR_miCentred_AT", "bsp_lRR_miCentred_AT:miCentred_logMass",
    "bsp_lRR_miCentred_AT:ResMR_Bin", "bsp_lRR_miCentred_logMass",
    "sigma_lRR"), names = c("Air Temp", "log Mass by °C", "Residual MR by °C",
    "log Mass", "Sigma"))
```

```
mcmc_neff(neff_ratio(Core_Model_New), size = 2)
```

```
modelSource <- paste0(
 "/Users/joshuatabh/Documents/researchProjects/trent/sihMetaregression/models/",
 "modelCoreCorrectedLong.Rds"
)

# Some possible autocorrelation remaining among a few parameters (presumably phylogeny and species-level intercepts again).
# Running model with more iterations, a longer warm-up duration, and a widened thinning process.

Core_Model_New <- brm(bf(Centred_logMass | mi(logMass_SD) ~ 0,
                     family = "gaussian") +
                    bf(Centred_AT | mi(Ambient.Temp.Stressor.sd) ~ 0,
                       family = "gaussian") +
                    bf(lRR | se(lRV, sigma = TRUE) ~ mi(Centred_AT)*mi(Centred_logMass) +
                           ResMR_Bin + ResMR_Bin:mi(Centred_AT) + Age_Class + Technique +
```

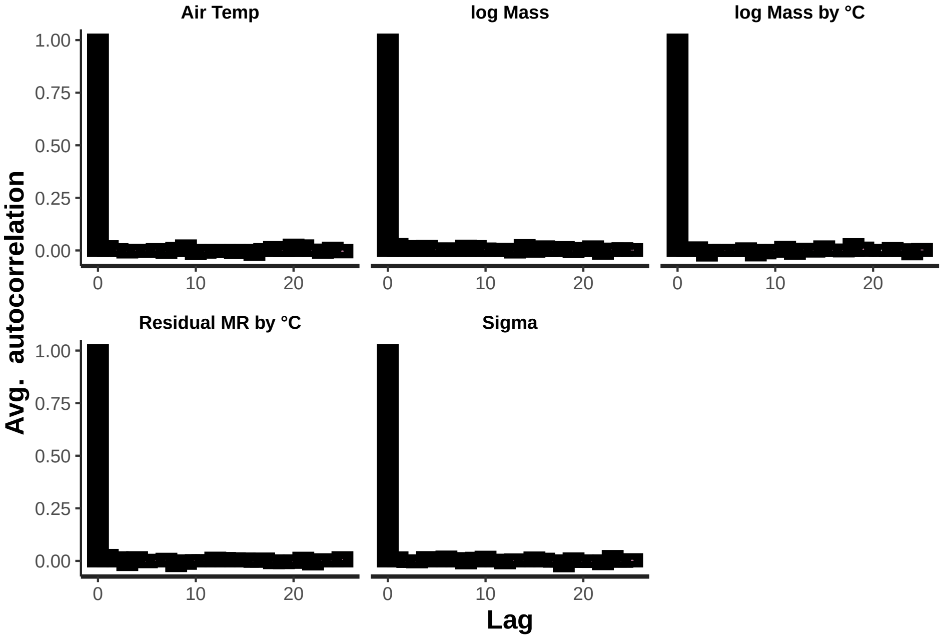


Figure 26: Degree of sequential autocorrelation between chain draws, per parameter, for a revised Bayesian, linear mixed effects model predicting stress-induced changes in body temperature (natural log-transformed response ratio, lRR'). Autocorrelation for select predictors is displayed.

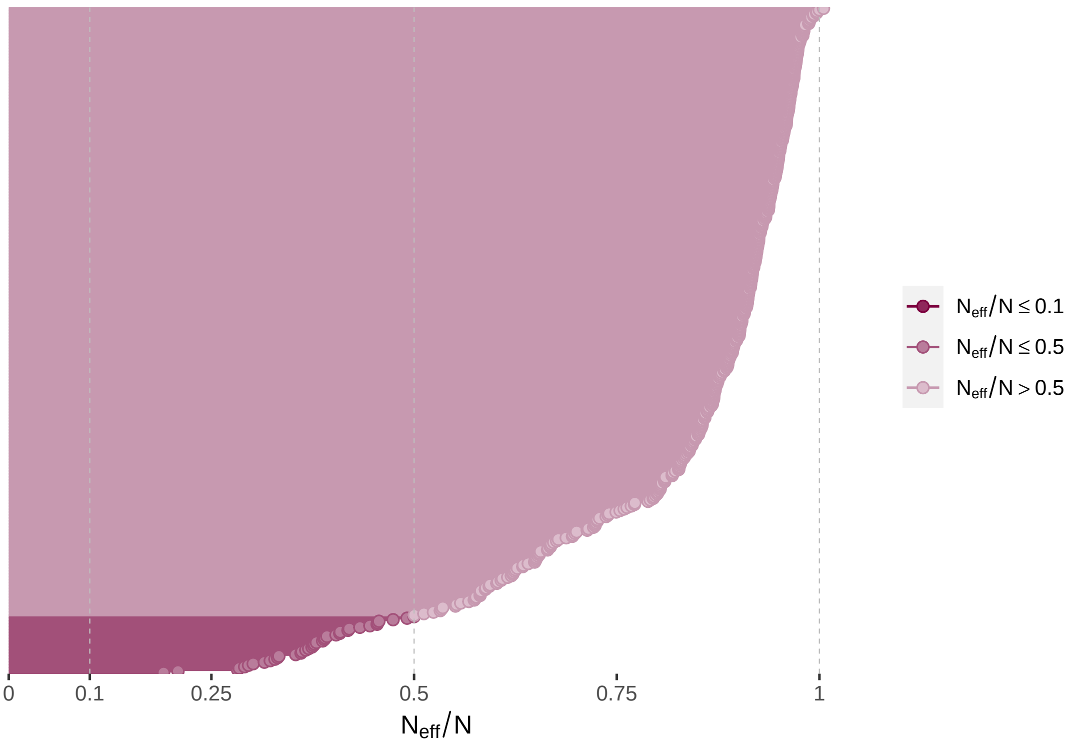


Figure 27: Ratio of effective sample sizes to sample sizes for a revised Bayesian, linear mixed effects model predicting stress-induced changes in body temperature (natural log-transformed response ratio, lRR').

```
                           poly(Centred_Time, 2) + (1 | StudyID) + (1 | Species.Name) +
                         (1 | gr(Phylo, cov = A)), family = "gaussian") +
                    set_rescor(FALSE),
data = Core_Data_MR_Shortened,
data2 = list(A = Core_VCV_MR),
cores = 4, chains = 4,
seed = 100, #refresh = 0,
iter = 100000, warmup = 50000, thin = 25,
prior = prior_Core,
save_pars = save_pars(latent = TRUE),
file = modelSource
)
```

```
mcmc_neff(neff_ratio(Core_Model_New), size = 2)
```

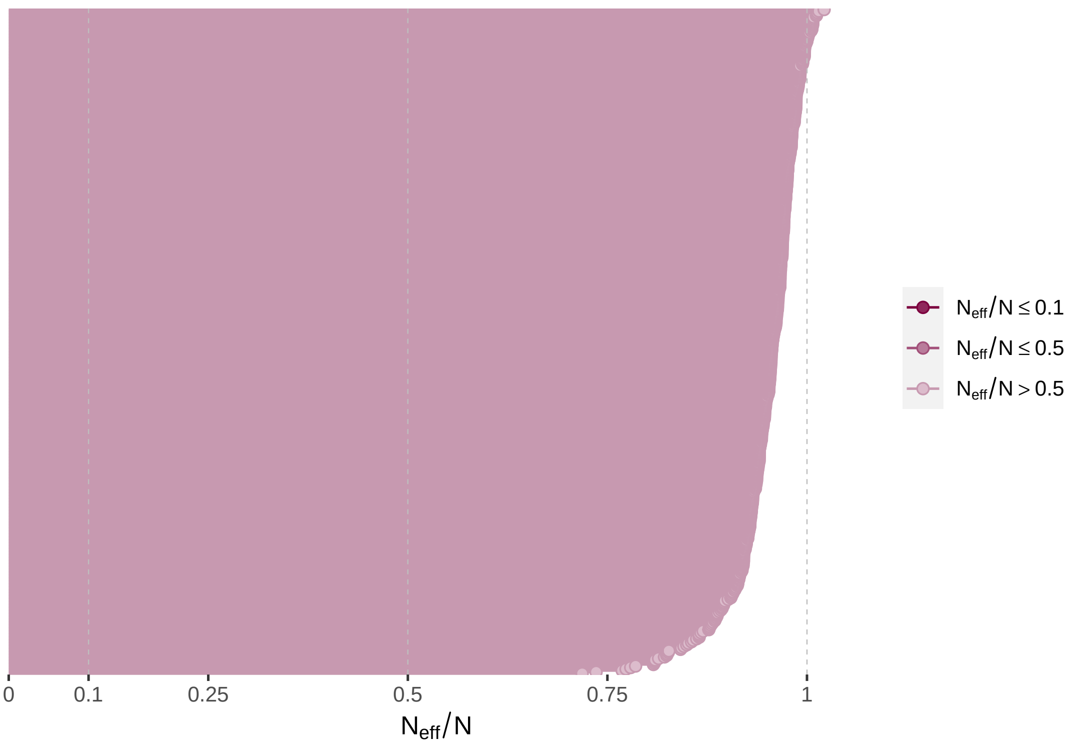


Figure 28: Updated ratio of effective sample sizes to sample sizes for the Bayesian, linear mixed effects model addressed in Fig. 27.

```
# Good. Proceeding to Rhats.
```

```
mcmc_rhat(rhat(Core_Model_New))
```

```
# Rhats tight to 1. Checking HMC intervals

clean_int(Core_Model_New, prs = c("b_lRR_Intercept", "b_lRR_Technique",
    "b_lRR_polyCentred_Time21", "b_lRR_polyCentred_Time22", "bsp_lRR_miCentred_AT",
    "bsp_lRR_miCentred_AT:miCentred_logMass", "bsp_lRR_miCentred_logMass",
    "b_lRR_ResMR_Bin", "bsp_lRR_miCentred_AT:ResMR_Bin", "b_lRR_Age_Class",
    "sigma_lRR"), names = c("Intercept", "Technique", "Time_1",
    "Time_2", "Air Temperature", "Air Temp by log Mass", "log Mass",
    "Residual RMR", "Air Temp by Residual MR", "Age Class", "Sigma"))
```

```
# Some noise around time. Pairing chain draws across key
# predictors.

clean_pairs(Core_Model_New, prs = c("bsp_lRR_miCentred_AT", "b_lRR_ResMR_Bin",
    "bsp_lRR_miCentred_logMass", "bsp_lRR_miCentred_AT:miCentred_logMass",
```

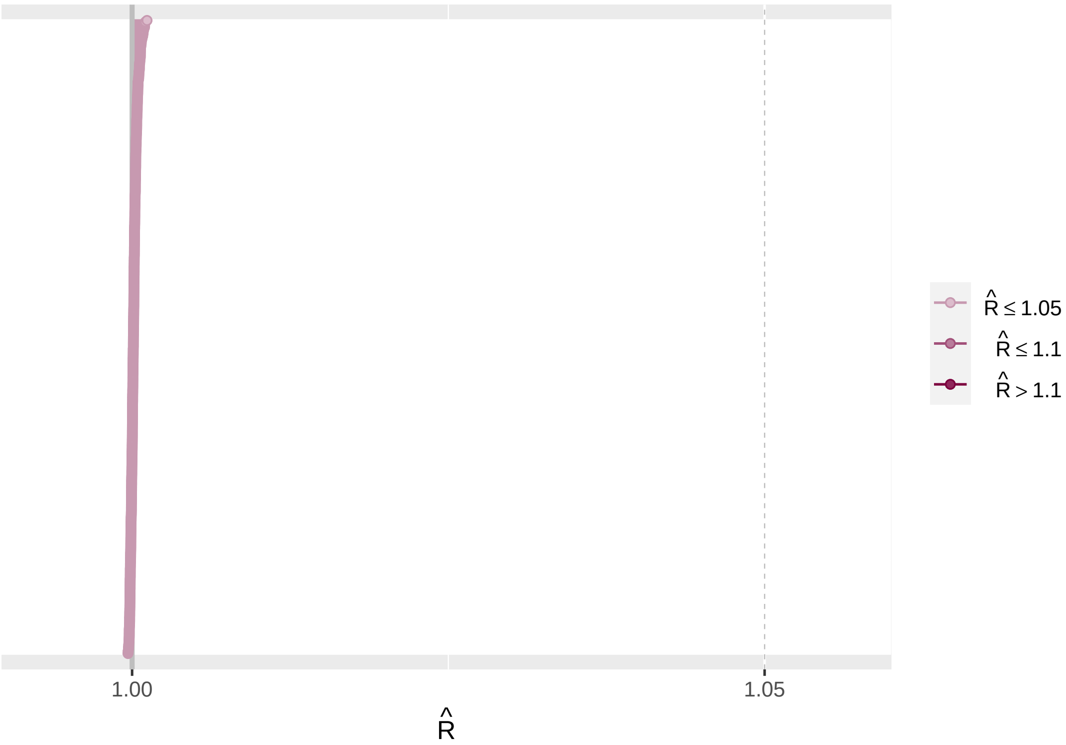


Figure 29: Gelman-Rubin statistics per model parameter for a revised Bayesian, linear mixed effects model predicting stress-induced changes in body temperature (natural log-transformed response ratio, lRR').

```
    "bsp_lRR_miCentred_AT:ResMR_Bin", "sigma_lRR"), names = c("Air Temp",
    "Residual RMR", "log Mass", "Air Temp by Mass", "Air Temp by Residual RMR",
    "Sigma"))
```

```
# No concerning patterns. However, double-checking that
# there is no serendipitous collinearity between mass and
# ambient temperature or residual resting metabolic rate
# and air temperature to begin with.
```

```
ggplot(Core_Data_MR_Shortened, aes(x = Ambient.Temp.Stressor,
    y = logMass)) + geom_errorbarh(aes(y = logMass, xmin = Ambient.Temp.Stressor -
    Ambient.Temp.Stressor.sd, xmax = Ambient.Temp.Stressor +
    Ambient.Temp.Stressor.sd), height = 0.4) + geom_errorbar(aes(x = Ambient.Temp.Stressor,
    ymin = logMass - logMass_SD, ymax = logMass + logMass_SD),
    width = 1) + geom_point(size = 3, pch = 21, alpha = 0.5,
    colour = "black", fill = nice_pink) + stat_ellipse(fill = "grey70",
    colour = "black", alpha = 0.5, geom = "polygon") + theme_classic() +
    xlab("Ambient Temperature (°C)") + ylab("Mass (g)")
```

```
ggplot(Core_Data_MR_Shortened, aes(x = Ambient.Temp.Stressor,
    y = ResMR)) + geom_errorbarh(aes(y = ResMR, xmin = Ambient.Temp.Stressor -
    Ambient.Temp.Stressor.sd, xmax = Ambient.Temp.Stressor +
    Ambient.Temp.Stressor.sd), height = 0.4) + geom_point(size = 3,
    pch = 21, alpha = 0.5, colour = "black", fill = nice_pink) +
    stat_ellipse(fill = "grey70", colour = "black", alpha = 0.5,
        geom = "polygon") + theme_classic() + xlab("Ambient Temperature (°C)") +
    ylab("Residual RMR (W)")
```

```
# Very little concern.
```

```
# Checking model residuals.

grid.arrange(Core_Model_New$data %>%
```

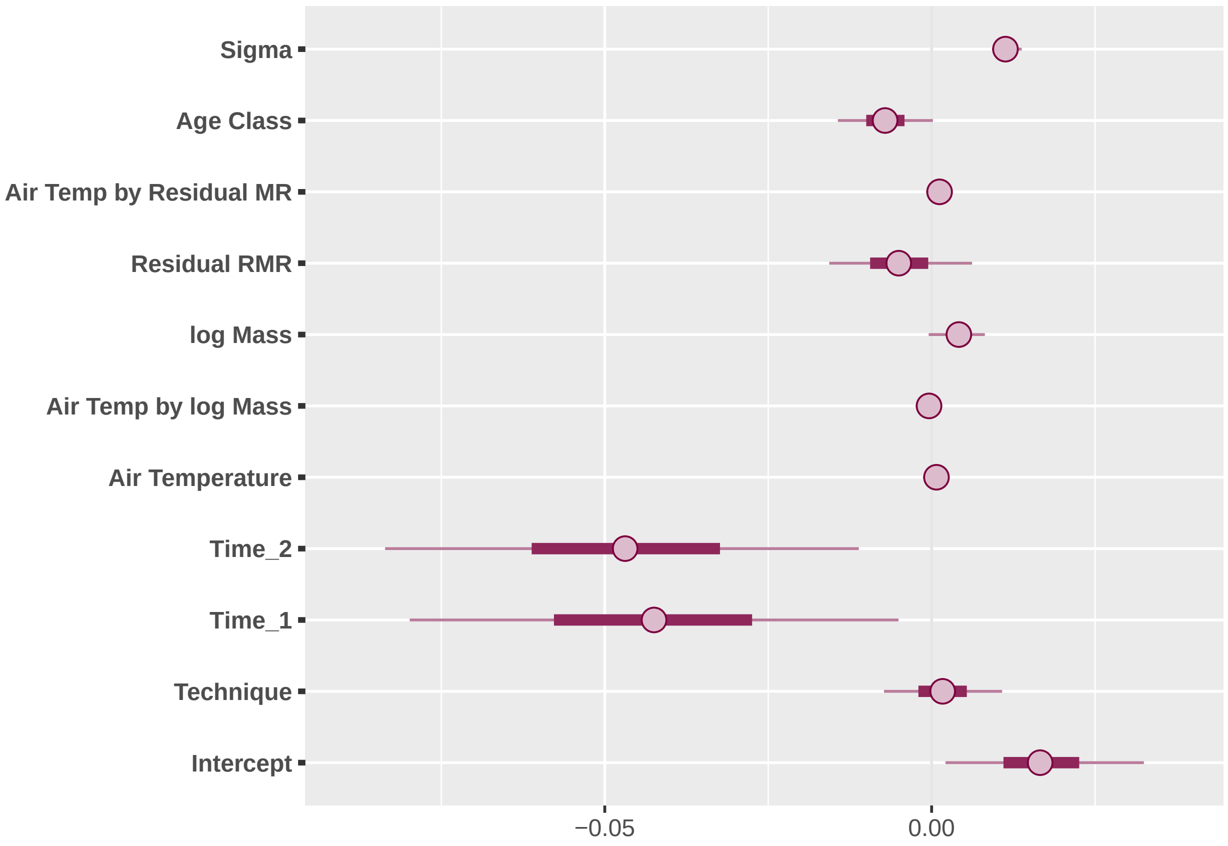


Figure 30: Mean coefficient estimates and 90 percent posterior intervals around such for select predictes from a revised Bayesian, linear mixed effects model predicting stress-induced changes in body temperature (natural log-transformed response ratios).

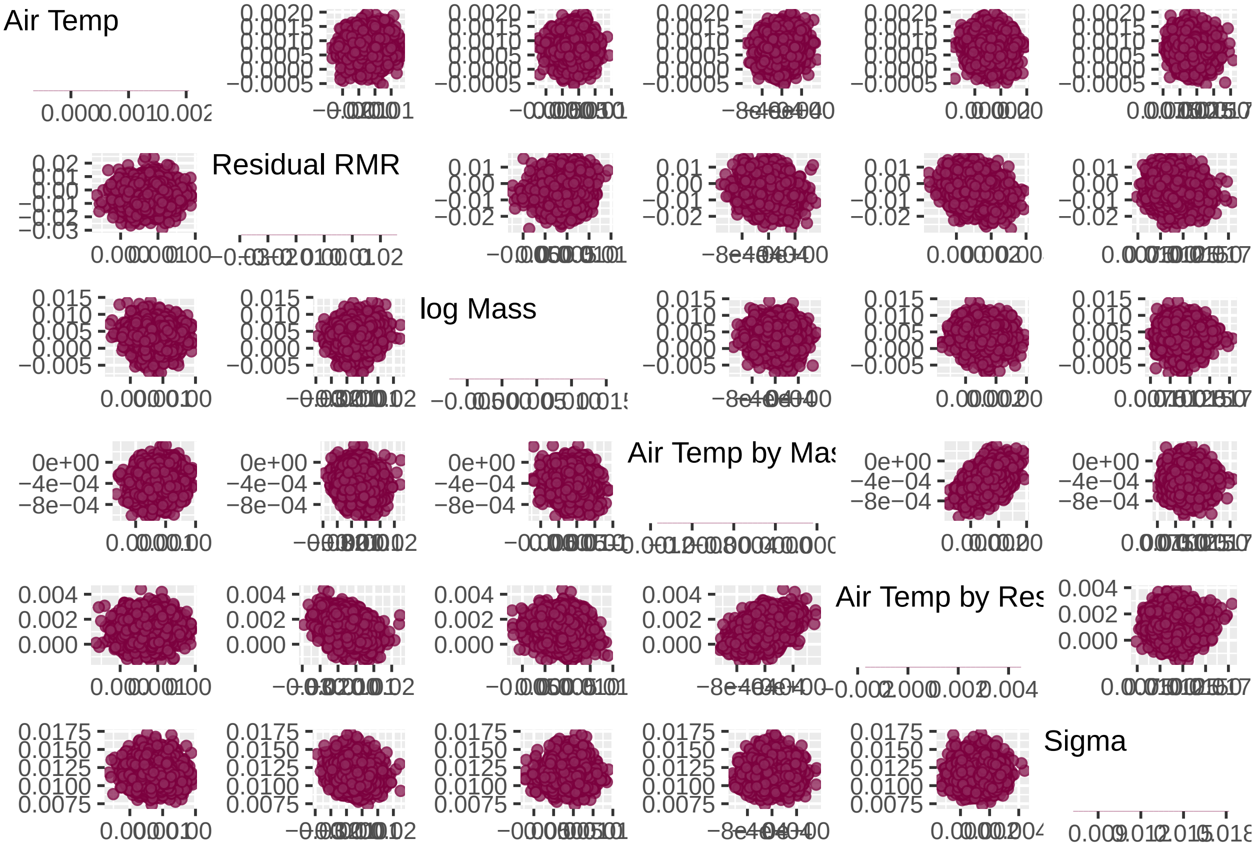


Figure 31: Paired, posterior coefficient estimates from a Bayesian, linear model predicting the magnitude and direction of stress-induced changes in body temperature (measured as natural log transformed response ratios). Select Parameter pairs for are displayed.

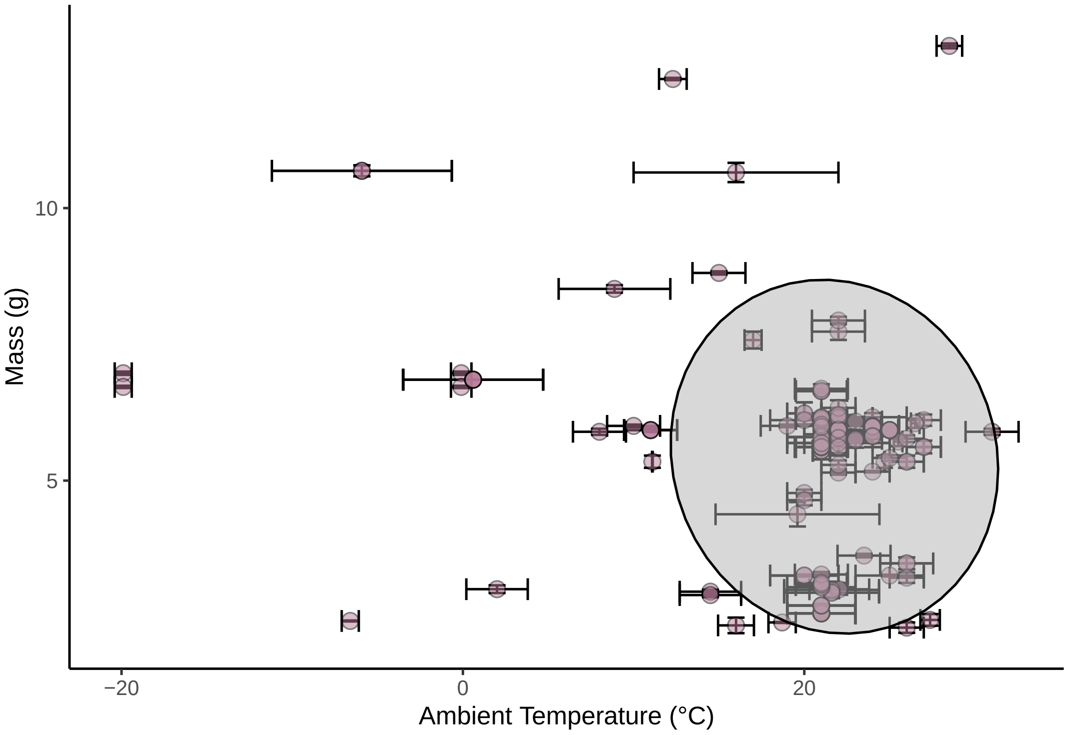


Figure 32: Mean body mass of sample, terrestrial endotherm populations (per study) as displayed against the ambient temperature at which stress-induced changes in body temperature (log response ratios) were measured. Dots respresent mean ambient temperature and body mass values per sample population and errorbars represent standard deviations around ambient temperature values.

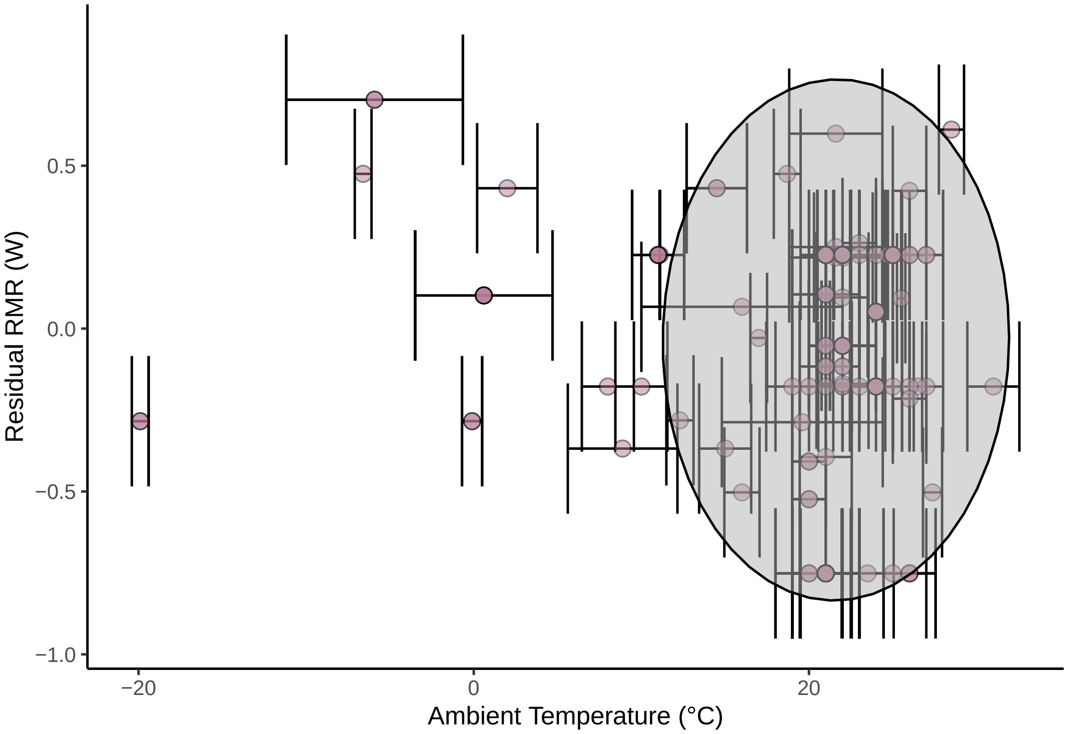


Figure 33: Relative metabolic rate (residual resting metabolic rate) of sample, terrestrial endotherm populations (per study) as displayed against the ambient temperature at which stress-induced changes in body temperature (log response ratios) were measured. Again, dots respresent mean ambient temperature and body mass values per sample population and errorbars represent standard deviations around ambient temperature values.

```
    mutate(Res = residuals(Core_Model_New, type = "ordinary",
        method = "posterior_predict")[, "Estimate", "lRR"]) %>%
    ggplot(aes(x = 1:nrow(.), y = Res)) + geom_point(size = 3,
    colour = nice_pink) + theme_classic() + xlab("Sample Number") +
    ylab("Ordinary Residuals"), Core_Model_New$data %>%
    mutate(Res = residuals(Core_Model_New, type = "ordinary",
        method = "posterior_predict")[, "Estimate", "lRR"]) %>%
    ggplot(aes(x = Res)) + geom_density(colour = "black", fill = nice_pink,
    adjust = 1.5) + theme_classic() + xlab("Ordinary Residuals") +
    ylab("Density"), nrow = 2)
```

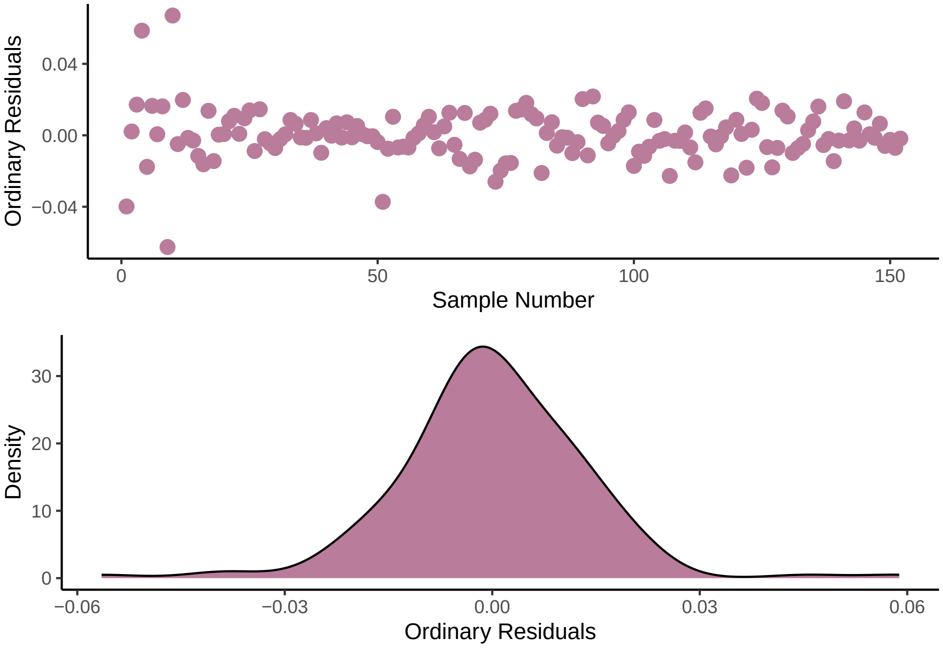


Figure 34: Ordinary, untransformed residuals for a revised Bayesian, linear mixed effects model predicting stress-induced changes in body temperature (natural log-transformed response ratio, lRR'), diplayed per dataframe row (top panel). Each dot represents the residual value for a single observation of a stress-induced change in body temperature for a single species. Density of ordinary residual values (bottom panel).

```
# Looks much cleaner. One data points straying slightly
# high, but not egregious.  Checking qqplot with labelled
# points
```

```
qq_base <- Core_Model_New$data %>%
    mutate(Res = residuals(Core_Model_New, type = "ordinary",
        method = "posterior_predict")[, "Estimate", "lRR"]) %>%
    na.omit(.) %>%
    ggplot(aes(sample = Res)) + stat_qq() + stat_qq_line()

qq_base_dat <- ggplot_build(qq_base)$data[[1]]
qq_base_dat$OBID <- Core_Data_MR_Shortened$ObservationID[order(Core_Data_MR_Shortened$ObservationID)]

qq_base_dat %>%
    ggplot(aes(x = theoretical, y = sample, label = OBID)) +
    geom_text() + geom_smooth(method = "lm", linetype = "dashed",
    colour = "darkred") + theme_classic() + ylim(c(-0.05, 0.07))
```

```
# Viewing those that diverge

Core_Data_MR_Shortened %>%
    mutate(Res = residuals(Core_Model_New, type = "ordinary")[,
        "Estimate", "lRR"]) %>%
    filter(ObservationID %in% c("AD", "PI")) %>%
```

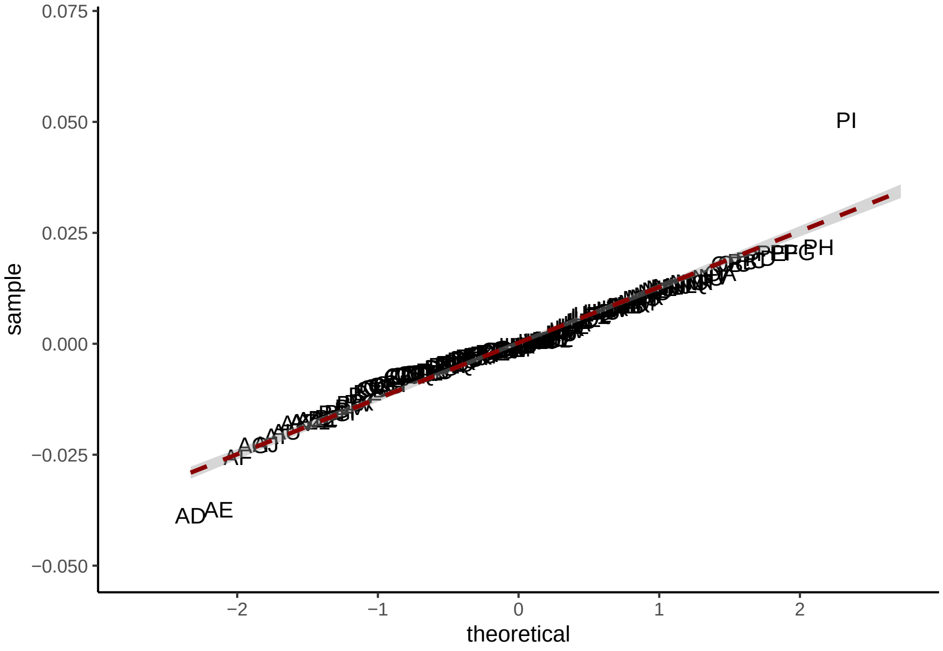


Figure 35: Quantile-quantile ('qq') plot of, untransformed residuals for a revised Bayesian, linear mixed effects model predicting stress-induced changes in body temperature (natural log-transformed response ratio, lRR').

```
select(Observation = ObservationID, `Study ID` = StudyID,
    `Latin Name` = Species.Name, `°C` = Ambient.Temp.Stressor,
    `log Mass` = logMass, `log Response Ratio` = lRR, Residuals = Res) %>%
kbl(., longtable = T, booktabs = T) %>%
kable_styling(latex_options = "striped")
```

| Observation | Study ID | Latin Name | °C | log Mass | log Response Ratio | Residuals |
|---|---|---|---|---|---|---|
| AD | B | Parus major | 14.5 | 2.900322 | -0.0133578 | 0.0108180 |
| PI | H6 | Rattus norvegicus domestica | 22.0 | 5.936216 | 0.0463007 | 0.0150745 |

```
# Nothing of obvious concern in these studies. Proceeding
# to check residuals against predictors and fitted values

Core_Model_New$data %>%
    mutate(Res = residuals(Core_Model_New, type = "ordinary",
        method = "predict")[, "Estimate", "lRR"], Res_SE = residuals(Core_Model_New,
        type = "ordinary", method = "predict")[, "Est.Error",
        "lRR"], Fit = fitted(Core_Model_New)[, "Estimate", "lRR"],
        Fit_SE = fitted(Core_Model_New)[, "Est.Error", "lRR"]) %>%
    ggplot(aes(x = Fit, y = Res)) + geom_errorbarh(aes(y = Res,
    xmin = Fit - Fit_SE, xmax = Fit + Fit_SE), height = 0.005,
    colour = "black", alpha = 0.5) + geom_errorbar(aes(x = Fit,
    ymin = Res - Res_SE, ymax = Res + Res_SE), width = 0.005,
    colour = "black", alpha = 0.5) + geom_point(size = 3, pch = 21,
    colour = "black", fill = nice_pink, alpha = 0.5) + stat_ellipse(geom = "polygon",
    fill = "gray70", alpha = 0.7, colour = "black") + theme_classic() +
    xlab("Ordinary Residuals") + ylab("Fitted Values")
```

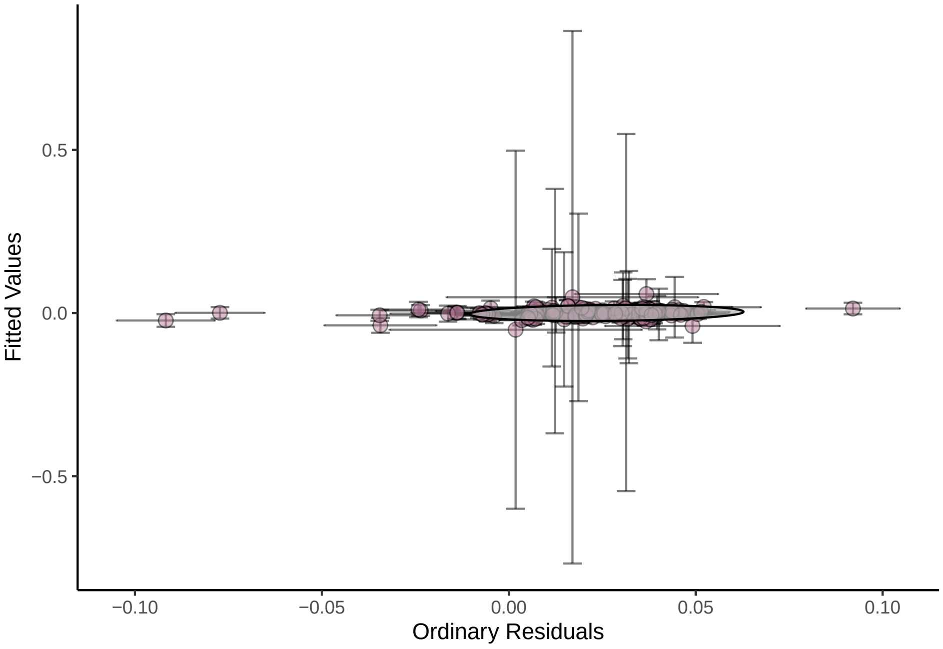


Figure 36: Fitted values against ordinary residual values for a revised Bayesian, linear mixed effects model predicting stress-induced changes in body temperature (natural log-transformed response ratio, lRR'). Dots represent mean fitted values across posterior draws while errorbars represent +/- standard error values around these means.

```
# Looks well dispersed. Plotting residuals across
# predictors to check for heteroskedasticity.
```

```
{
    p1 <- Core_Model_New$data %>%
        mutate(Res = residuals(Core_Model_New, type = "ordinary",
            method = "predict")[, "Estimate", "lRR"]) %>%
        mutate(Technique = ifelse(Technique == -0.5, "Probe",
            "Telemetry")) %>%
        ggplot(aes(x = Technique, y = Res, fill = Technique)) +
        geom_boxplot(width = 0.2) + ggdist::stat_halfeye(slab_colour = "black",
        adjust = 0.7, width = 0.3, .width = 0, justification = -0.8,
        point_colour = NA) + geom_point(size = 1, alpha = 0.3,
        position = position_jitter(seed = 1, width = 0.1)) +
        coord_cartesian(xlim = c(1.2, NA), clip = "off") + theme_classic() +
        scale_fill_manual(values = c(nice_pink, "slateblue"),
            name = "Measurement Technique") + theme(legend.position = "none") +
```

```
        xlab("Measurement Technique") + ylab("Ordinary Residuals")

    p2 <- Core_Model_New$data %>%
        mutate(Res = residuals(Core_Model_New, type = "ordinary",
            method = "predict")[, "Estimate", "lRR"], Res_SE = residuals(Core_Model_New,
            type = "ordinary", method = "predict")[, "Est.Error",
            "lRR"]) %>%
        ggplot(aes(x = Centred_AT, y = Res)) + geom_errorbar(aes(x = Centred_AT,
        ymin = Res - Res_SE, ymax = Res + Res_SE), width = 0.8,
        alpha = 0.5) + geom_point(size = 3, pch = 21, colour = "black",
        fill = nice_pink, alpha = 0.5) + geom_smooth(method = "loess",
        colour = "black") + theme_classic() + xlab("Mean-Centred Ambient Temperature (°C)") +
        ylab("Ordinary Residuals")

    p3 <- Core_Model_New$data %>%
        mutate(Res = residuals(Core_Model_New, type = "ordinary",
            method = "predict")[, "Estimate", "lRR"]) %>%
        mutate(ResMR_Bin = ifelse(ResMR_Bin == -0.5, "Low", "High")) %>%
        ggplot(aes(x = ResMR_Bin, y = Res, fill = ResMR_Bin)) +
        geom_boxplot(width = 0.2) + ggdist::stat_halfeye(slab_colour = "black",
        adjust = 0.7, width = 0.3, .width = 0, justification = -0.8,
        point_colour = NA) + geom_point(size = 1, alpha = 0.3,
        position = position_jitter(seed = 1, width = 0.1)) +
        coord_cartesian(xlim = c(1.2, NA), clip = "off") + theme_classic() +
        scale_fill_manual(values = c(nice_pink, "slateblue"),
            name = "Measurement Technique") + theme(legend.position = "none") +
        xlab("Residual Resting Metabolic Rate (W)") + ylab("Ordinary Residuals")

    p4 <- Core_Model_New$data %>%
        mutate(Res = residuals(Core_Model_New, type = "ordinary",
            method = "predict")[, "Estimate", "lRR"], Res_SE = residuals(Core_Model_New,
            type = "ordinary", method = "predict")[, "Est.Error",
            "lRR"]) %>%
        mutate(ResMR_Bin = ifelse(ResMR_Bin == -0.5, "Low", "High")) %>%
        ggplot(aes(x = Centred_AT, y = Res, fill = ResMR_Bin,
            )) + geom_errorbar(aes(x = Centred_AT, ymin = Res -
        Res_SE, ymax = Res + Res_SE), width = 0.5, alpha = 0.5) +
        geom_point(size = 3, pch = 21, colour = "black", alpha = 0.5) +
        geom_smooth(method = "loess", colour = "black") + scale_fill_manual(values = c(nice_pink,
        "slateblue"), name = "Relative Resting\nMetabolic Rate") +
        theme_classic() + xlab("Centred Air Temperature (°C)") +
        ylab("Ordinary Residuals")

    p5 <- grid.arrange(Core_Model_New$data %>%
        mutate(Res = residuals(Core_Model_New, type = "ordinary",
            method = "predict")[, "Estimate", "lRR"], Res_SE = residuals(Core_Model_New,
            type = "ordinary", method = "predict")[, "Est.Error",
            "lRR"]) %>%
        ggplot(aes(x = Centred_logMass, y = Res, fill = Centred_AT,
            )) + geom_errorbar(aes(x = Centred_logMass, ymin = Res -
        Res_SE, ymax = Res + Res_SE), width = 0.5, alpha = 0.5) +
        geom_point(size = 3, pch = 21, colour = "black", alpha = 0.5) +
        geom_smooth(method = "loess", colour = "black") + scale_fill_gradient2(low = "slateblue",
        mid = "navajowhite", high = nice_pink) + theme_classic() +
        theme(legend.position = "none") + xlab("Centred log Mass (g)") +
        ylab("Ordinary Residuals"), Core_Model_New$data %>%
        mutate(Res = residuals(Core_Model_New, type = "ordinary",
            method = "predict")[, "Estimate", "lRR"], Res_SE = residuals(Core_Model_New,
            type = "ordinary", method = "predict")[, "Est.Error",
            "lRR"]) %>%
        ggplot(aes(x = Centred_logMass, y = Res, fill = Centred_AT,
            )) + geom_point(size = 3, pch = 21, colour = "black",
        alpha = 0.5) + stat_ellipse(geom = "polygon", fill = "gray70",
        alpha = 0.3, colour = "black") + scale_fill_gradient2(low = "slateblue",
        mid = "navajowhite", high = nice_pink, name = "Centred Ambient\nTemperature (°C)") +
        theme_classic() + xlab("Centred log Mass (g)") + ylab("Ordinary Residuals"),
        nrow = 1)
```

```
    p6 <- Core_Model_New$data %>%
        mutate(Res = residuals(Core_Model_New, type = "ordinary",
            method = "predict")[, "Estimate", "lRR"], Res_SE = residuals(Core_Model_New,
            type = "ordinary", method = "predict")[, "Est.Error",
            "lRR"]) %>%
        ggplot(aes(x = Centred_Time, y = Res)) + geom_errorbar(aes(x = Centred_Time,
        ymin = Res - Res_SE, ymax = Res + Res_SE), width = 0.5,
        alpha = 0.5) + geom_point(size = 3, pch = 21, colour = "black",
        fill = nice_pink) + geom_smooth(method = "loess", colour = "black") +
        theme_classic() + xlab("Time to Peak or Trough\nBody Temperature (s)") +
        ylab("Ordinary Residuals")

    p7 <- Core_Model_New$data %>%
        mutate(Res = residuals(Core_Model_New, type = "ordinary",
            method = "predict")[, "Estimate", "lRR"]) %>%
        ggplot(aes(x = Species.Name, y = Res, fill = Species.Name)) +
        geom_boxplot() + scale_fill_viridis_d(option = "B") +
        theme_classic() + theme(legend.position = "none", axis.text.x = element_text(angle = 90)) +
        xlab("Species") + ylab("Ordinary Residuals")
}
```

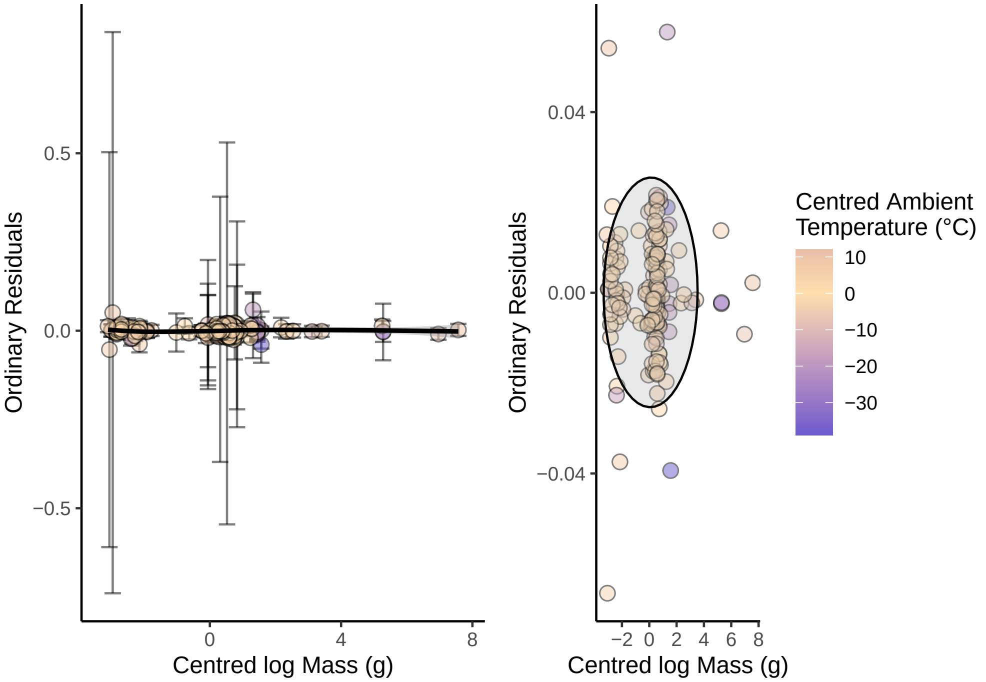


Figure 37: Distribution of ordinary residuals across body temperature measurements techniques. Ordinary residual values are means across posterior draws and are derived from a revised Bayesian, linear mixed effects model predicting stress-induced changes in body temperature (natural log-transformed response ratio, lRR’). Boxplots represent median values (central bars) with inter-quartiles ranges (box limits) and ranges exluding outliers (whiskers). Outliers represent values beyond the median +/- 1.5 x the interquartile range. Black dots represent individual residual values per smaple population.

```
print(p1)
```

```
print(p2)
```

```
print(p3)
```

```
print(p4)
```

```
plot(p5)
```

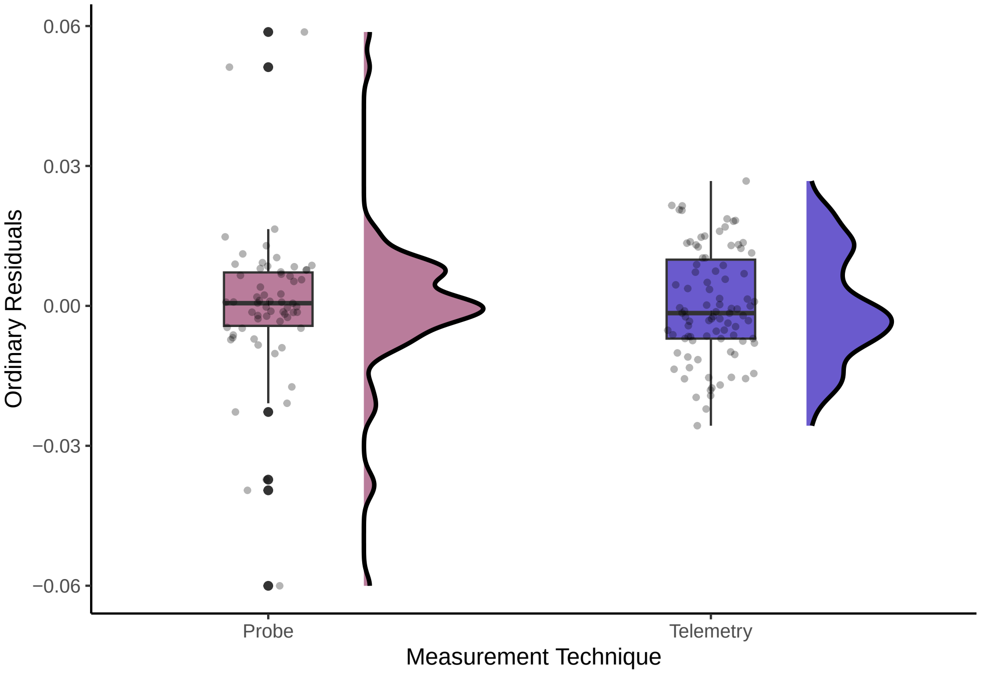


Figure 38: Distribution of ordinary residuals across body temperature measurements techniques. Ordinary residual values are means across posterior draws and are derived from a revised Bayesian, linear mixed effects model predicting stress-induced changes in body temperature (natural log-transformed response ratio, lRR'). Boxplots represent median values (central bars) with inter-quartiles ranges (box limits) and ranges exluding outliers (whiskers). Outliers represent values beyond the median +/- 1.5 x the interquartile range. Black dots represent individual residual values per smaple population.

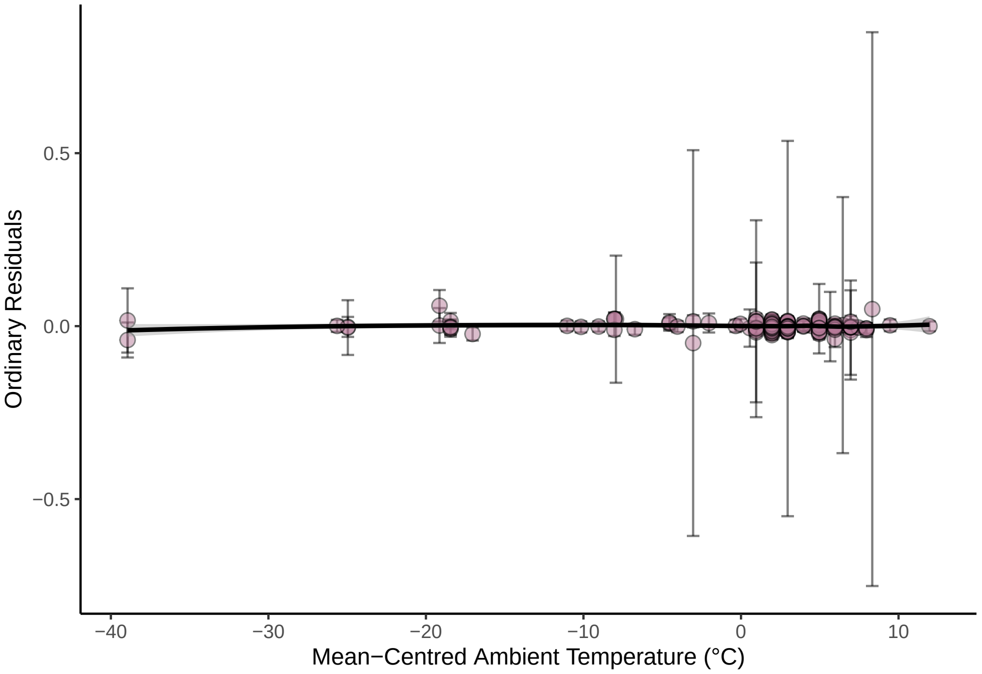


Figure 39: Ordinary residuals across mean-centred ambient temperature measurements (°C) during stress-induced change in body temperature measurements. Ordinary residual values are means across posterior draws and are derived from a revised Bayesian, linear mixed effects model predicting stress-induced changes in body temperature (natural log-transformed response ratio, lRR'). Errorbars represent standard errors aroung mean residual values.

```
print(p6)
```

```
# Perhaps some heteroskedasticity by measurement technique,
# but not of particular concern.
```

```
print(p7)
```

```
rm(p1, p2, p3, p4, p5, p6, p7)
```

Next, we visually assess both the conditional and marginal fits of our model by regressing fitted values (obtained with and without group-level effects respectively) against observed values of our log response ratio.

```
# Visualising model fit.

Core_Model_New$data %>%
    mutate(Fit = fitted(Core_Model_New)[, "Estimate", "lRR"],
        Fit_SE = fitted(Core_Model_New)[, "Est.Error", "lRR"]) %>%
    ggplot(aes(x = Fit, y = lRR)) + geom_errorbarh(aes(y = lRR,
    xmin = Fit - Fit_SE, xmax = Fit + Fit_SE), height = 0.01,
    alpha = 0.5) + geom_point(size = 3, pch = 21, fill = nice_pink,
    alpha = 0.5) + geom_smooth(method = "lm", colour = "black",
    linetype = "dashed", se = FALSE) + theme_classic() + xlab("Fitted Value") +
    ylab("Log Response Ratio")
```

```
# Good. Plotting marginal fit.

Core_Model_New$data %>%
    mutate(Fit = fitted(Core_Model_New, re_formula = NA)[, "Estimate",
        "lRR"], Fit_SE = fitted(Core_Model_New, re_formula = NA)[,
```

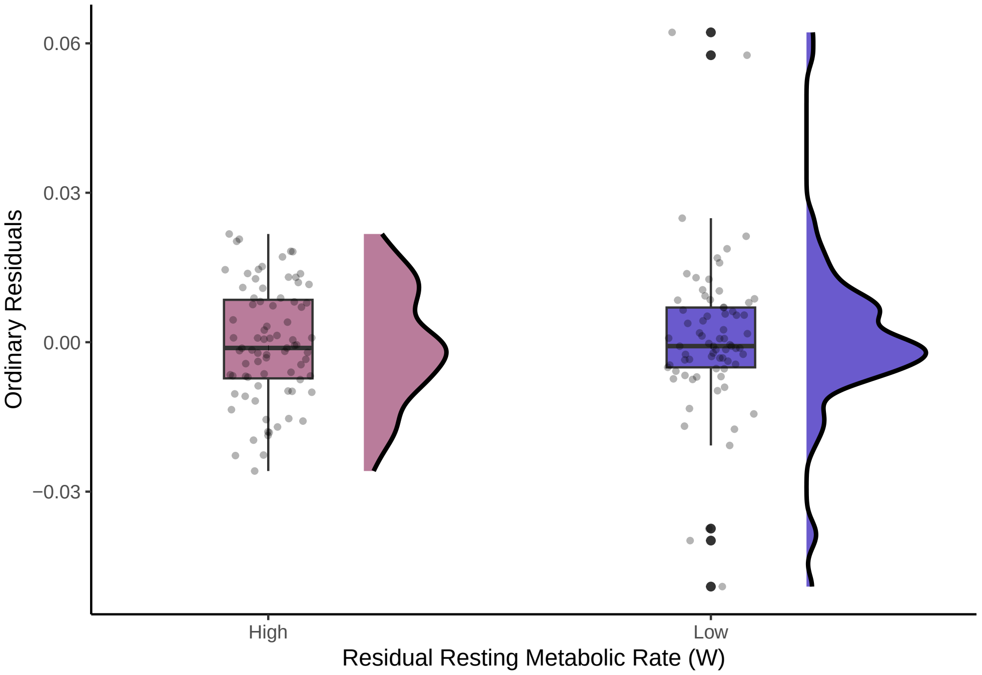


Figure 40: Distribution of ordinary residuals according to relative (residual) resting metabolic expenditure. Ordinary residual values are means across posterior draws and are derived from a revised Bayesian, linear mixed effects model predicting stress-induced changes in body temperature (natural log-transformed response ratio, lRR'). Boxplots represent median values (central bars) with inter-quartiles ranges (box limits) and ranges exluding outliers (whiskers). Outliers represent values beyond the median +/- 1.5 x the interquartile range. Black dots represent individual residual values per smaple population.

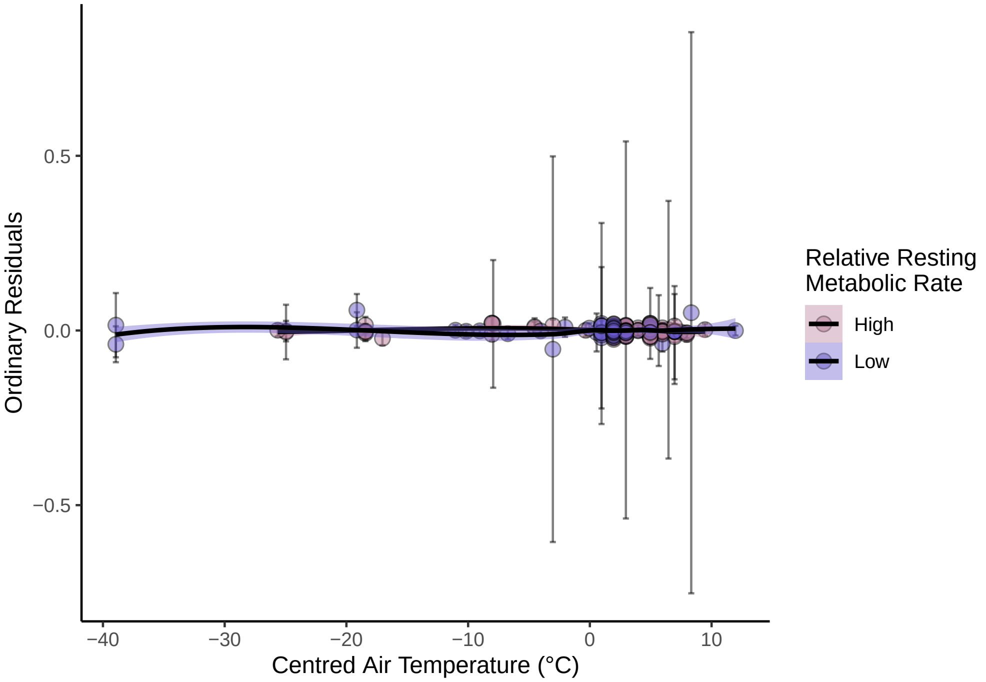


Figure 41: Ordinary residuals across both mean-centred ambient temperature measurements (°C; during stress-induced change in body temperature measurements) and relative resting metabolic expenditure (residuals). Ordinary residual values are means across posterior draws and are derived from a revised Bayesian, linear mixed effects model predicting stress-induced changes in body temperature (natural log-transformed response ratio, lRR'). Errorbars represent standard errors aroung mean residual values.

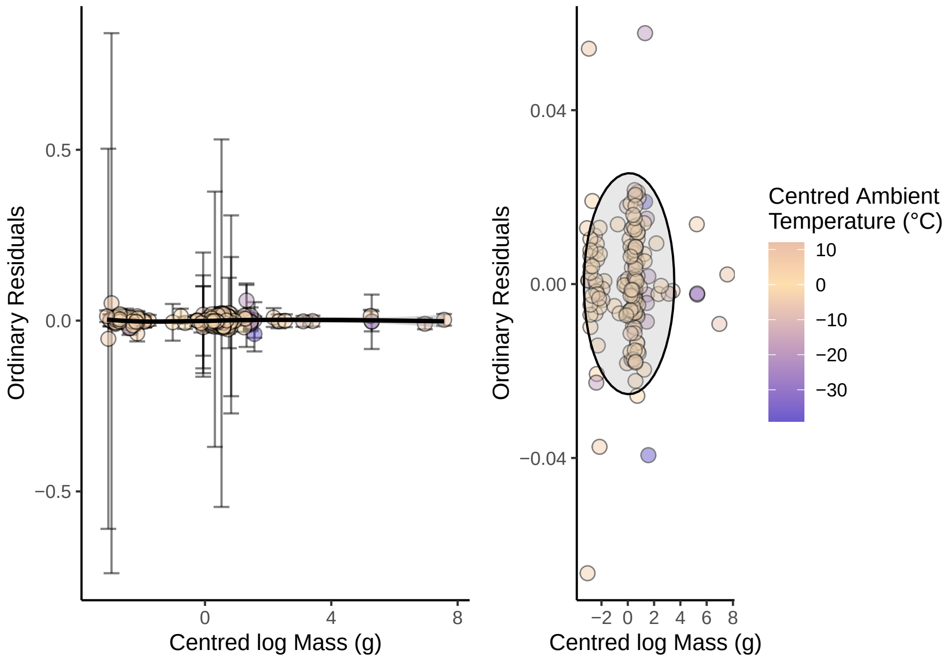


Figure 42: Ordinary residuals across mean-centred, natural log-transfomred body mass of sample populations (g; left panel), and both log-tranformed body mass and mean-centred ambient temperature values (°C; during stress-induced change in body temperature measurements; right panel). Ordinary residual values are means across posterior draws and are derived from a revised Bayesian, linear mixed effects model predicting stress-induced changes in body temperature (natural log-transformed response ratio, lRR'). Errorbars represent standard errors aroung mean residual values.

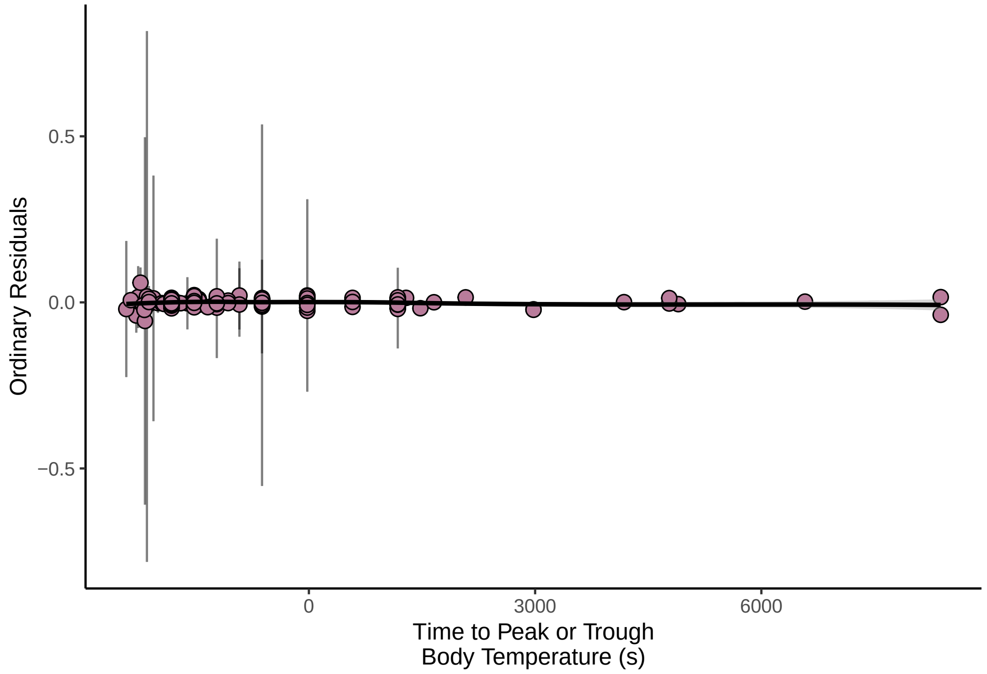


Figure 43: Ordinary residuals across latency between stress-exposure and maximum or minimum stress-induced change in body temperature measurements (s). Ordinary residual values are means across posterior draws and are derived from a revised Bayesian, linear mixed effects model predicting stress-induced changes in body temperature (natural log-transformed response ratio, lRR'). Errorbars represent standard errors aroung mean residual values.

```
        "Est.Error", "lRR"]) %>%
    ggplot(aes(x = Fit, y = lRR)) + geom_errorbarh(aes(y = lRR,
    xmin = Fit - Fit_SE, xmax = Fit + Fit_SE), height = 0.01,
    alpha = 0.5) + geom_point(size = 2, pch = 21, fill = nice_pink,
    alpha = 0.5) + geom_smooth(method = "lm", colour = "black",
    linetype = "dashed", se = FALSE) + theme_classic() + xlab("Fitted Value") +
    ylab("Log Response Ratio")
```

```
# Notably poorer. Calculating loo R2 from each approach.

loo_R2(Core_Model_New)
```

```
##                      Estimate   Est.Error        Q2.5      Q97.5
## R2CentredlogMass 0.0055499478 0.009285829 -0.02050895 0.01572797
## R2CentredAT      0.0001227698 0.018787794 -0.05280667 0.01838870
## R2lRR            0.5081212359 0.083036273  0.33895190 0.65899876
```

```
## And lastly, checking whether posteriors fit within a
## reasonable scope of our data.

post_samples <- Core_Model_New %>%
    spread_draws(b_lRR_Intercept, b_lRR_Age_Class, b_lRR_ResMR_Bin,
        b_lRR_Technique, b_lRR_polyCentred_Time21, b_lRR_polyCentred_Time22,
        bsp_lRR_miCentred_AT, `bsp_lRR_miCentred_AT:miCentred_logMass`,
        `bsp_lRR_miCentred_AT:ResMR_Bin`, bsp_lRR_miCentred_logMass) %>%
    rename(Intercept = b_lRR_Intercept, Age = b_lRR_Age_Class,
        Residual_MR = b_lRR_ResMR_Bin, Technique = b_lRR_Technique,
        Time_1 = b_lRR_polyCentred_Time21, Time_2 = b_lRR_polyCentred_Time22,
        Air_Temp = bsp_lRR_miCentred_AT, Mass = bsp_lRR_miCentred_logMass,
        AirTemp_by_Mass = `bsp_lRR_miCentred_AT:miCentred_logMass`,
        AirTemp_by_ResMR = `bsp_lRR_miCentred_AT:ResMR_Bin`) %>%
    select(-c(.chain, .iteration, .draw))
```

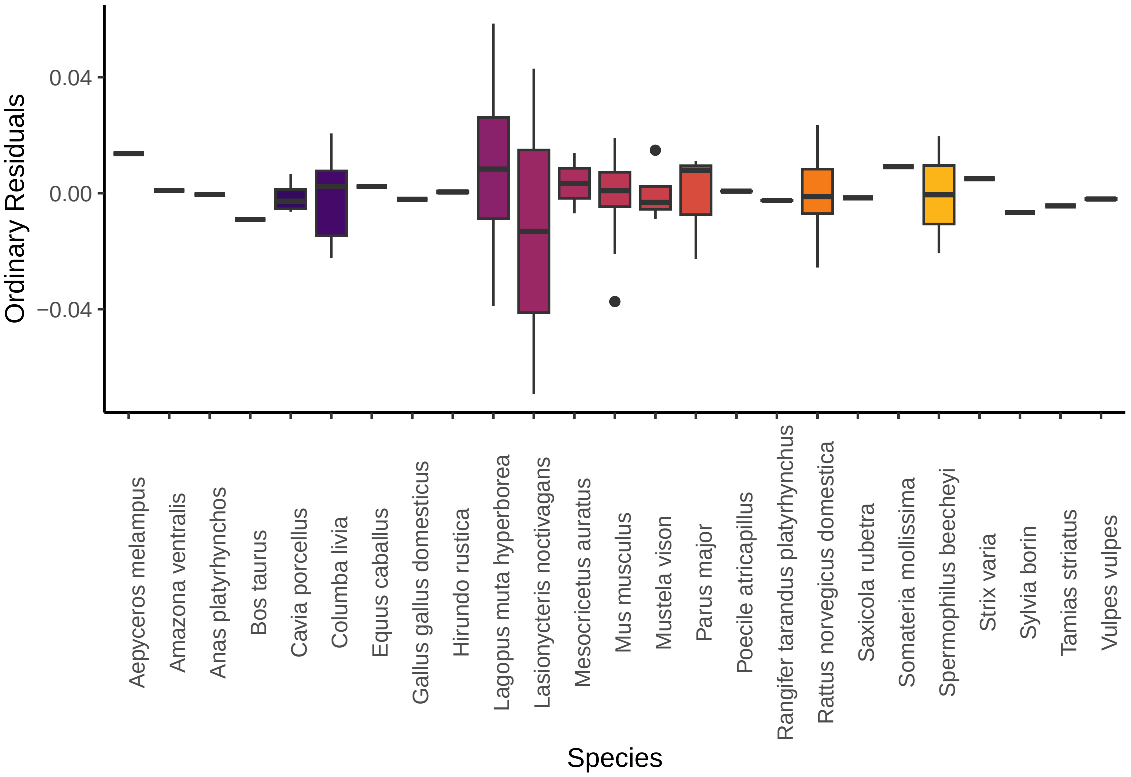


Figure 44: Distribution of ordinary residuals according to species. Ordinary residual values are means across posterior draws and are derived from a revised Bayesian, linear mixed effects model predicting stress-induced changes in body temperature (natural log-transformed response ratio, lRR’). Boxplots represent median values (central bars) with inter-quartiles ranges (box limits) and ranges exluding outliers (whiskers). Outliers (black dots) represent values beyond the median +/- 1.5 x the interquartile range.

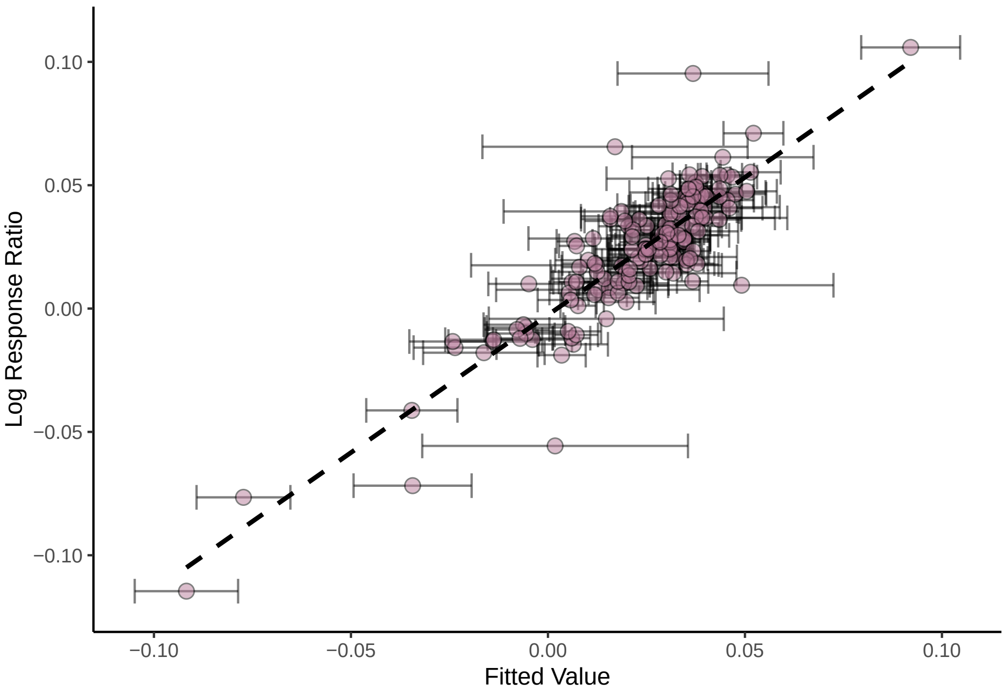


Figure 45: True stress-induced change in body temperature values (natural log-transformed response ratios) against fitted values from a revised Bayesian, linear mixed effects model predicting stress-induced changes in body temperature. Dots represent mean fitted values across model posterior draws and errorbards represent standard errors around means values. The dashed line represents a line of best fit (estimated for a linear correlation).

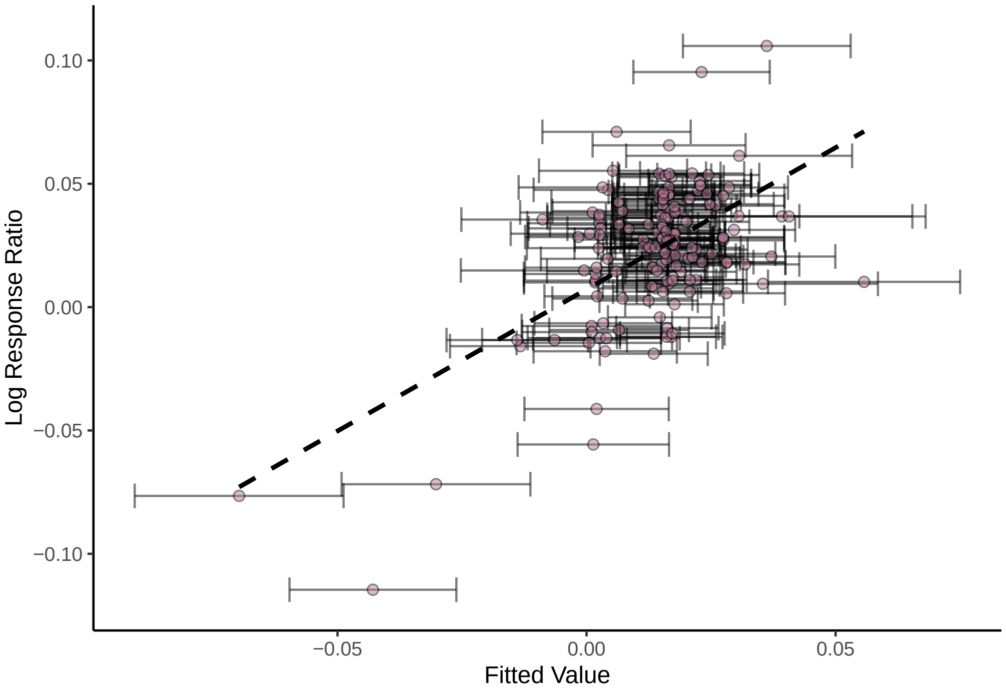


Figure 46: True stress-induced change in body temperature values (natural log-transformed response ratios) against marginal fitted values from a revised Bayesian, linear mixed effects model predicting stress-induced changes in body temperature. Dots represent mean fitted values across model posterior draws and errorbards represent standard errors around means values. The dashed line represents a line of best fit (estimated for a linear correlation). Note that fitted values here exclude the influence of group-level, model parameters (i.e. species, study identity, and phylogeny).

```
rbind(post_samples %>%
    mutate(Age = median(Intercept) + Age * range(Core_Data_MR_Shortened$Age_Class)[1],
        Residual_MR = median(Intercept) + Residual_MR * 0, Time_1 = median(Intercept) +
            Time_1 * range(poly(Core_Data_MR_Shortened$Centred_Time,
                2)[, 1])[1], Time_2 = median(Intercept) + Time_2 *
            range(poly(Core_Data_MR_Shortened$Centred_Time, 2)[,
                2])[1], Air_Temp_1 = median(Intercept) + Air_Temp *
            range(Core_Data_MR_Shortened$Centred_AT)[1], Mass_1 = median(Intercept) +
            Mass * range(Core_Data_MR_Shortened$Centred_logMass)[1],
        AirTemp_by_Mass = median(Intercept) + Air_Temp * range(Core_Data_MR_Shortened$Centred_AT)[1] +
            Mass * range(Core_Data_MR_Shortened$Centred_logMass)[1] +
            AirTemp_by_Mass * range(range(Core_Data_MR_Shortened$Centred_AT)[1]) *
                range(Core_Data_MR_Shortened$Centred_logMass)[1],
        AirTemp_by_ResMR = median(Intercept) + Air_Temp * range(Core_Data_MR_Shortened$Centred_AT)[1] +
            Residual_MR * 0 + AirTemp_by_ResMR * range(range(Core_Data_MR_Shortened$Centred_AT)[1]) *
            0) %>%
    select(-c("Air_Temp", "Mass")) %>%
    rename(Air_Temp = "Air_Temp_1", Mass = "Mass_1") %>%
    mutate(Range = "Low"), post_samples %>%
    mutate(Age = median(Intercept) + Age * range(Core_Data_MR_Shortened$Age_Class)[2],
        Residual_MR = median(Intercept) + Residual_MR * 1, Time_1 = median(Intercept) +
            Time_1 * range(poly(Core_Data_MR_Shortened$Centred_Time,
                2)[, 1])[2], Time_2 = median(Intercept) + Time_2 *
            range(poly(Core_Data_MR_Shortened$Centred_Time, 2)[,
                2])[2], Air_Temp_1 = median(Intercept) + Air_Temp *
            range(Core_Data_MR_Shortened$Centred_AT)[2], Mass_1 = median(Intercept) +
            Mass * range(Core_Data_MR_Shortened$Centred_logMass)[2],
        AirTemp_by_Mass = median(Intercept) + Air_Temp * range(Core_Data_MR_Shortened$Centred_AT)[2] +
            Mass * range(Core_Data_MR_Shortened$Centred_logMass)[2] +
            AirTemp_by_Mass * range(range(Core_Data_MR_Shortened$Centred_AT)[2]) *
                range(Core_Data_MR_Shortened$Centred_logMass)[2],
        AirTemp_by_ResMR = median(Intercept) + Air_Temp * range(Core_Data_MR_Shortened$Centred_AT)[1] +
            Residual_MR * 1 + AirTemp_by_ResMR * range(range(Core_Data_MR_Shortened$Centred_AT)[1]) *
            1) %>%
    select(-c("Air_Temp", "Mass")) %>%
    rename(Air_Temp = "Air_Temp_1", Mass = "Mass_1") %>%
    mutate(Range = "High")) %>%
    mutate(across(where(is.double), function(x) exp(x) * 38 -
        38)) %>%
    pivot_longer(-Range) %>%
    arrange(name) %>%
    ggplot(aes(x = value, fill = Range)) + geom_density(colour = "black",
    adjust = 1.5, alpha = 0.4) + scale_fill_manual(values = c(nice_pink,
    "royalblue2")) + theme_classic() + xlab("Change in Body Temperature (°C)") +
    ylab("Density") + facet_wrap(~name, scales = "free") + annotate("rect",
    xmin = min(Core_Data_MR_Shortened$Delta.Tb, na.rm = T), xmax = max(Core_Data_MR_Shortened$Delta.Tb,
        na.rm = T), ymin = -Inf, ymax = Inf, colour = "black",
    fill = "grey20", alpha = 0.3) + ggtitle("Grey box represents values observed in data set.")
```

```
rm(post_samples)
```

```
# Roughly comparing conditional and marginal fits by regressing fitted values (obtained with and without
# group-level effects respectively) against observed values. Note that we use cmdstan to parallel
# process HMC sampling within chains. Information on how to install cmdstan is available at:
# https://mc-stan.org/docs/2_25/cmdstan-guide/cmdstan-installation.html. Those who do not have
# cmdstan installed and do not wish to do so should remove 'backend = "cmdstanr"' and
# 'threads = threading(2)' from the below models before executing them.

CompDat <- Core_Model_New$data %>%
  mutate(
    "CFit" = fitted(Core_Model_New)[, "Estimate", "lRR"],
    "CFit_SE" = fitted(Core_Model_New)[, "Est.Error", "lRR"],
    "MFit" = fitted(Core_Model_New, re_formula = NA)[, "Estimate", "lRR"],
    "MFit_SE" = fitted(Core_Model_New, re_formula = NA)[, "Est.Error", "lRR"]
  )

# Assuming a slope of 1 with little error.
```

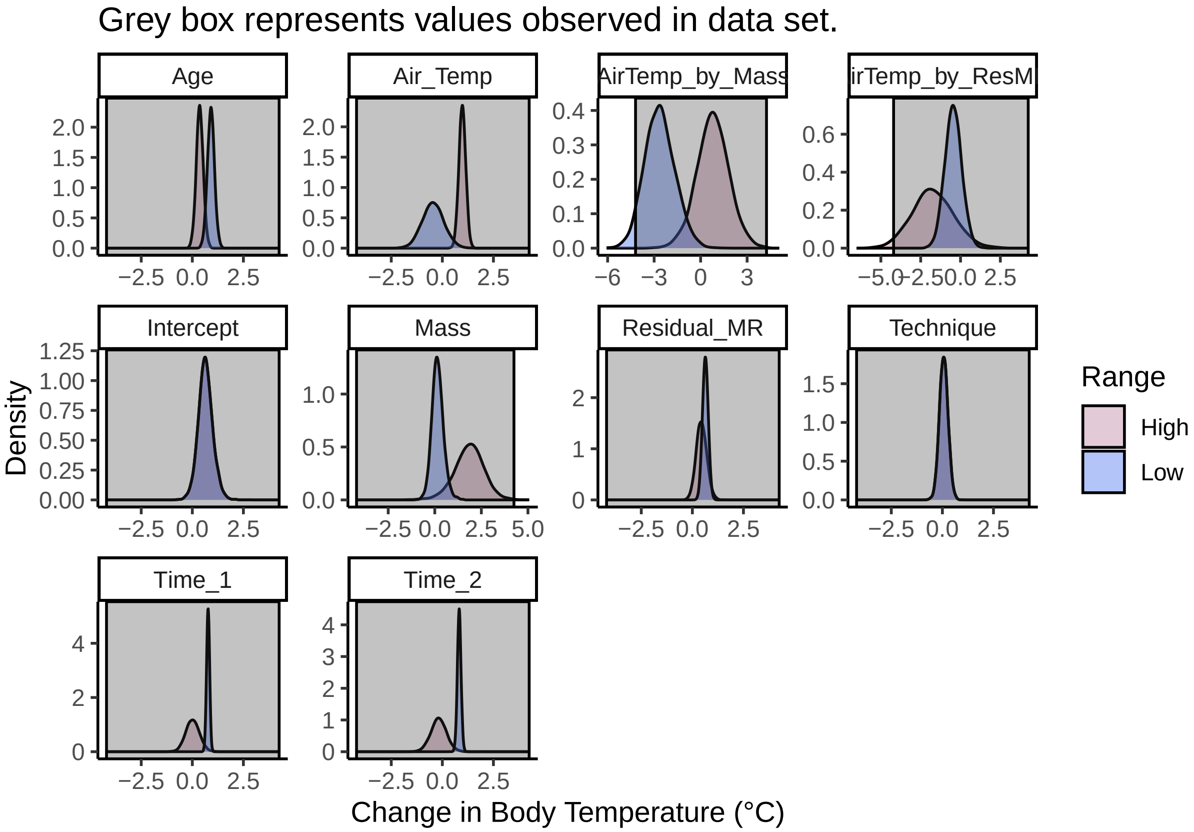


Figure 47: Density of coefficent estimates, per parameter, as derived from a revised Bayesian, linear mixed effects model predicting stress-induced changes in body temperature (natural log-transformed response ratios). Grey rectangles indicate ranges of true stress-induced change in body temperature values.

```r
prior_CFit <- c(
  set_prior("normal(1,1)", class = "b", coef = "CFit")
)

# Setting cmdstan path

#set_cmdstan_path("/home/joshk/git_repositories/cmdstan")

MO = "/Users/joshuatabh/Documents/researchProjects/trent/sihMetaregression/models/conditionalFitModel.Rds"

brm_fit1 <- brm(lRR ~ CFit,
  data = CompDat,
  cores = 1, chains = 2,
  #backend = "cmdstanr", threads = threading(2),
  seed = 200, silent = TRUE, refresh = 0,
  family = "gaussian",
  iter = 20000, warmup = 5000, thin = 10,
  control = list(adapt_delta = 0.95, max_treedepth = 13),
  prior = prior_CFit,
  file = MO
)

rm(MO)

prior_MFit <- c(
  set_prior("normal(1,1)", class = "b", coef = "MFit")
)

MO = "/Users/joshuatabh/Documents/researchProjects/trent/sihMetaregression/models/marginalFitModel.Rds"

brm_fit2 <- brm(lRR ~ MFit,
  data = CompDat,
  cores = 1, chains = 2,
  #backend = "cmdstanr", threads = threading(2),
  seed = 200, silent = TRUE, #refresh = 0,
  family = "gaussian",
  iter = 20000, warmup = 5000, thin = 10,
  control = list(adapt_delta = 0.95, max_treedepth = 13),
  prior = prior_MFit,
  file = MO
)

rm(MO)

# Comparing loo R2 values.

lapply(list(brm_fit1, brm_fit2), loo_R2)
```

```
## [[1]]
##    Estimate  Est.Error      Q2.5     Q97.5
## R2  0.75604 0.05576747 0.6375217 0.8529443
##
## [[2]]
##    Estimate Est.Error       Q2.5     Q97.5
## R2 0.281537 0.1051835 0.05646126 0.4704207
```

```r
# Conditional loo R2 = 0.76 and marginal loo R2 = 0.26.

rm(brm_fit1, brm_fit2)
```

Finally, we calculate the posterior mode of each model coefficient (here, rounded to 3 significant figures), alongside both 80% and 95% highest posterior density intervals.

```r
# Lastly, viewing and summarising model coefficients.

clean_dens(Core_Model_New, prs = c("b_lRR_Intercept", "b_lRR_Technique",
    "b_lRR_ResMR_Bin", "b_lRR_Age_Class", "b_lRR_polyCentred_Time21",
```

```
    "b_lRR_polyCentred_Time22"), names = c("Intercept", "Technique",
    "Residual MR", "Age Class", "Time 1", "Time 2")) + geom_vline(xintercept = 0,
    linetype = "dashed", colour = "black")
```

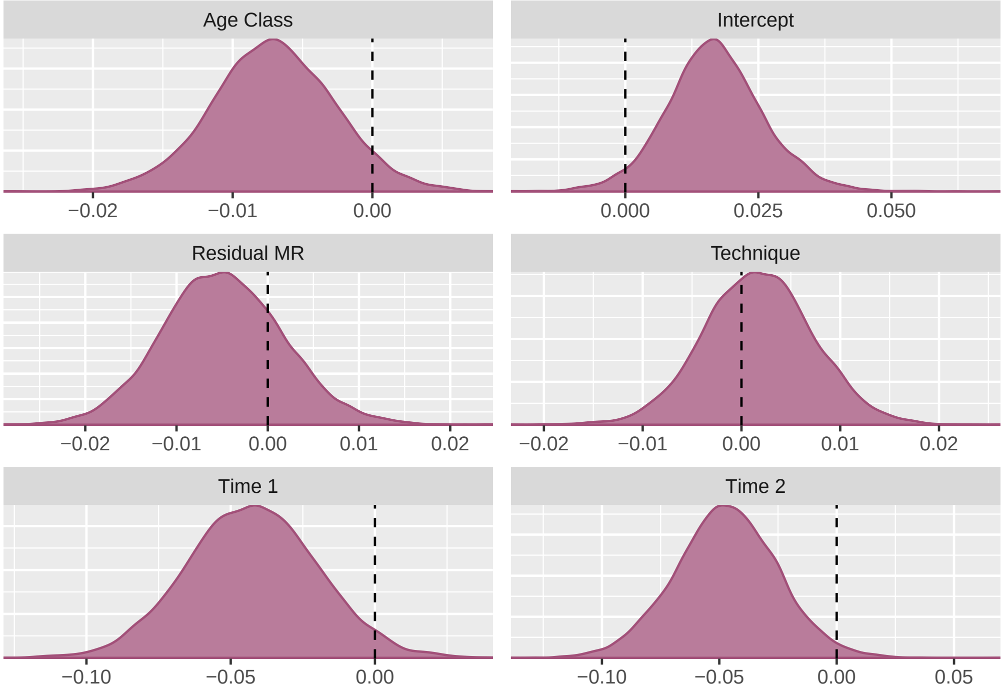


Figure 48: Density of coefficent estimates, per parameter, as derived from a revised Bayesian, linear mixed effects model predicting stress-induced changes in body temperature (natural log-transformed response ratios). Vertical dashed lines indicate 0 (no effect).

```
clean_dens(Core_Model_New, prs = c("bsp_lRR_miCentred_AT", "bsp_lRR_miCentred_AT:miCentred_logMass",
    "bsp_lRR_miCentred_AT:ResMR_Bin", "bsp_lRR_miCentred_logMass",
    "sigma_lRR"), names = c("Ambient °C", "log Mass by °C",
    "Residual MR by °C", "log Mass", "Sigma")) + geom_vline(xintercept = 0,
    linetype = "dashed", colour = "black")
```

```
# Very little kurtosis to posterior distributions, although
# those for phylogeny and species ID are a bit poorly
# defined.  Calculating modes and HPD intervals.

Core_Model_Table <- Core_Model_New %>%
    spread_draws(b_lRR_Intercept, bsp_lRR_miCentred_AT, b_lRR_ResMR_Bin,
        b_lRR_polyCentred_Time21, b_lRR_polyCentred_Time22, bsp_lRR_miCentred_logMass,
        b_lRR_Technique, b_lRR_Age_Class, `bsp_lRR_miCentred_AT:miCentred_logMass`,
        `bsp_lRR_miCentred_AT:ResMR_Bin`, sd_Phylo__lRR_Intercept,
        sd_StudyID__lRR_Intercept, sd_Species.Name__lRR_Intercept,
        sigma_lRR) %>%
    select(-c(.chain, .iteration, .draw)) %>%
    mutate_all(., .funs = round, 5) %>%
    summarise_all(., .funs = md) %>%
    pivot_longer(everything(), names_to = "Parameter", values_to = "Mode") %>%
    left_join(., simple_hdi(Core_Model_New, rnd = 5, cis = c(80,
        95), sci_note = FALSE), by = c("Parameter")) %>%
    right_join(tribble(~Par, ~Parameter, "Intercept", "b_lRR_Intercept",
        "Air Temperature", "bsp_lRR_miCentred_AT", "Residual RMR",
        "b_lRR_ResMR_Bin", "Time (1st order)", "b_lRR_polyCentred_Time21",
        "Time (2nd order)", "b_lRR_polyCentred_Time22", "Measurement Technique",
        "b_lRR_Technique", "Age Class", "b_lRR_Age_Class", "log Mass",
        "bsp_lRR_miCentred_logMass", "Air Temperature by log Mass",
        "bsp_lRR_miCentred_AT:miCentred_logMass", "Air Temperature by Residual RMR",
```

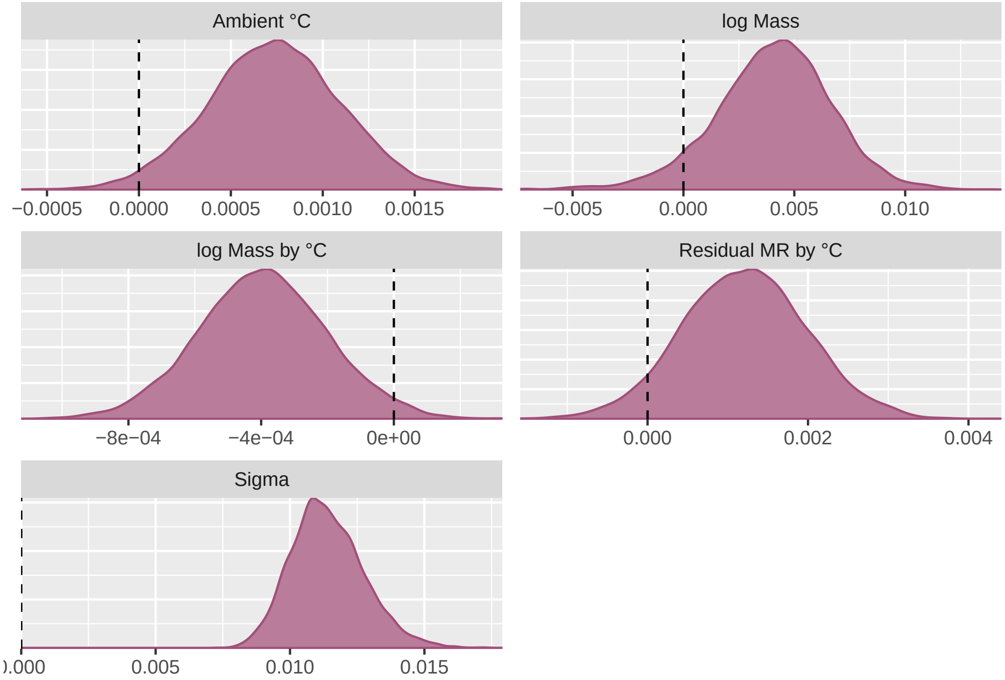


Figure 49: Density of coefficent estimates, per parameter, as derived from a revised Bayesian, linear mixed effects model predicting stress-induced changes in body temperature (natural log-transformed response ratios). Vertical dashed lines indicate 0 (no effect).

```
        "bsp_lRR_miCentred_AT:ResMR_Bin", "Phylogeny", "sd_Phylo__lRR_Intercept",
        "Species", "sd_Species.Name__lRR_Intercept", "Study ID",
        "sd_StudyID__lRR_Intercept", "Sigma", "sigma_lRR"), .,
        by = c("Parameter")) %>%
    select(-c(Parameter)) %>%
    mutate(` ` = ifelse((Low_HDI_80 < 0 & High_HDI_80 < 0) |
        (Low_HDI_80 > 0 & High_HDI_80 > 0), "*", ""), `  ` = ifelse((Low_HDI_95 <
        0 & High_HDI_95 < 0) | (Low_HDI_95 > 0 & High_HDI_95 >
        0), "*", "")) %>%
    select(Par, Mode, Low_HDI_80, High_HDI_80, " ", Low_HDI_95,
        High_HDI_95, "  ") %>%
    rename(Parameter = Par, `Low HDI (80%)` = Low_HDI_80, `High HDI (80%)` = High_HDI_80,
        `Low HDI (95%)` = Low_HDI_95, `High HDI (95%)` = High_HDI_95)

caption <- paste0("Paramaters predicting stress-induced changes in core body temperature",
    "across a sample of endotherms; results from a Bayesian mixed effects model.",
    " * indicates that highest density credible intervals do not cross 0.")

Core_Model_Table %>%
    rename(`Low 80%` = "Low HDI (80%)", `High 80%` = "High HDI (80%)",
        `Low 95%` = "Low HDI (95%)", `High 95%` = "High HDI (95%)",
        ) %>%
    mutate(Parameter = ifelse(Parameter == "Air Temperature by log Mass",
        "Air Temperature by\nlog Mass", Parameter)) %>%
    mutate(Parameter = ifelse(Parameter == "Air Temperature by Residual RMR",
        "Air Temperature by\nResidual RMR", Parameter)) %>%
    kbl(., longtable = T, booktabs = T, caption = caption) %>%
    kable_styling(latex_options = "striped")
```

Table 14: Paramaters predicting stress-induced changes in core body temperatureacross a sample of endotherms; results from a Bayesian mixed effects model. * indicates that highest density credible intervals do not cross 0.

| Parameter | Mode | Low 80% | High 80% | | Low 95% | High 95% | |
|---|---|---|---|---|---|---|---|
| Intercept | 0.01673 | 0.00441 | 0.02733 | * | -0.00178 | 0.03502 | |
| Air Temperature | 0.00077 | 0.00031 | 0.00123 | * | 0.00004 | 0.00144 | * |
| Residual RMR | -0.00398 | -0.01336 | 0.00343 | | -0.01764 | 0.00847 | |
| Time (1st order) | -0.03614 | -0.07137 | -0.01370 | * | -0.08533 | 0.00250 | |
| Time (2nd order) | -0.05039 | -0.07502 | -0.01957 | * | -0.08990 | -0.00334 | * |
| Measurement Technique | 0.00048 | -0.00545 | 0.00866 | | -0.00932 | 0.01238 | |
| Age Class | -0.00697 | -0.01253 | -0.00137 | * | -0.01619 | 0.00139 | |
| log Mass | 0.00425 | 0.00126 | 0.00768 | * | -0.00128 | 0.00917 | |
| Air Temperature by log Mass | -0.00040 | -0.00064 | -0.00015 | * | -0.00078 | -0.00001 | * |
| Air Temperature by Residual RMR | 0.00138 | 0.00026 | 0.00223 | * | -0.00027 | 0.00279 | |
| Phylogeny | 0.00150 | 0.00000 | 0.00223 | | 0.00000 | 0.00325 | |
| Species | 0.01492 | 0.00755 | 0.02941 | * | 0.00004 | 0.03219 | * |
| Study ID | 0.01455 | 0.01070 | 0.01807 | * | 0.00910 | 0.02066 | * |
| Sigma | 0.01075 | NA | NA | NA | NA | NA | NA |

```r
# Calculating descriptive statistics

## Assessing responses below 18˚C for various body masses
## and residual metabolic rates

Descriptive_Mass_Cold <- expand.grid(Centred_logMass = as.numeric(quantile(Core_Data_MR_Shortened$Centred_logMass)),
    logMass_SD = c(1e-04), Centred_AT = seq(-19.9, 17.9, length.out = 5) -
        mean(Core_Data_MR_Shortened$Ambient.Temp.Stressor), Ambient.Temp.Stressor.sd = c(1e-04),
    ResMR_Bin = 0, Technique = 0, Centred_Time = 0, Age_Class = 0,
    lRV = c(1e-04))

Descriptive_MR_Cold <- expand.grid(Centred_logMass = 0, logMass_SD = c(1e-04),
    Centred_AT = seq(-19.9, 17.9, length.out = 5) - mean(Core_Data_MR_Shortened$Ambient.Temp.Stressor),
    Ambient.Temp.Stressor.sd = c(1e-04), ResMR_Bin = c(-0.5,
        0.5), Technique = 0, Centred_Time = 0, Age_Class = 0,
    lRV = c(1e-04))

with(Core_Data_MR_Shortened, paste0("Mean time-to-stress-exposure (s): ",
    mean(Time.to.Peak.s.)))
```

```
## [1] "Mean time-to-stress-exposure (s): 1918.06842105263"
```

```r
caption <- paste0("Predicted changes in core body temperature after stress exposure",
    "at various air temperatures. Predictions are based upon body mass quantiles, ",
    "average time-post-stress-exposure (1683 s), mixed age groupings, and assume baseline ",
    "body temperature of 38°C.")

as.data.frame(predict(Core_Model_New, newdata = Descriptive_Mass_Cold,
    re_form = NA)[, , "lRR"]) %>%
    mutate(AT = rep(seq(-19.9, 17.9, length.out = 5), each = 5),
        CLM = rep(as.numeric(quantile(Core_Data_MR_Shortened$Centred_logMass)),
            5)) %>%
    group_by(AT, CLM) %>%
    summarise(Estimate = mean(Estimate), Error = mean(Est.Error)) %>%
    mutate(Delta_Tb = exp(Estimate) * 38 - 38, CLM = exp(CLM +
        mean(Core_Data_MR_Shortened$logMass))) %>%
    select(`Air Temperature (°C)` = AT, `Body Mass (g)` = CLM,
        Estimate, Error, `Delta Body Temperature (°C)` = Delta_Tb) %>%
    kbl(., longtable = T, booktabs = T, caption = caption) %>%
    kable_styling(latex_options = "striped")
```

Table 15: Predicted changes in core body temperature after stress exposureat various air temperatures. Predictions are based upon body mass quantiles, average time-post-stress-exposure (1683 s), mixed age groupings, and assume baseline body temperature of 38°C.

| Air Temperature (°C) | Body Mass (g) | Estimate | Error | Delta Body Temperature (°C) |
|---|---|---|---|---|
| -19.90 | 10.58817 | -0.0710539 | 0.0324956 | -2.6063547 |
| -19.90 | 121.12362 | -0.0228527 | 0.0206747 | -0.8585560 |

| | | | | |
|---|---|---|---|---|
| -19.90 | 384.88010 | -0.0004995 | 0.0204664 | -0.0189760 |
| -19.90 | 476.46780 | 0.0039828 | 0.0207200 | 0.1516490 |
| -19.90 | 457091.44411 | 0.1390308 | 0.0652751 | 5.6680616 |
| -10.45 | 10.58817 | -0.0522327 | 0.0277536 | -1.9338975 |
| -10.45 | 121.12362 | -0.0132810 | 0.0185084 | -0.5013396 |
| -10.45 | 384.88010 | 0.0050805 | 0.0180339 | 0.1935484 |
| -10.45 | 476.46780 | 0.0083938 | 0.0177916 | 0.3203070 |
| -10.45 | 457091.44411 | 0.1173178 | 0.0517461 | 4.7301163 |
| -1.00 | 10.58817 | -0.0335920 | 0.0232176 | -1.2552952 |
| -1.00 | 121.12362 | -0.0042229 | 0.0165571 | -0.1601301 |
| -1.00 | 384.88010 | 0.0098866 | 0.0161519 | 0.3775551 |
| -1.00 | 476.46780 | 0.0123906 | 0.0162739 | 0.4737735 |
| -1.00 | 457091.44411 | 0.0961725 | 0.0390414 | 3.8360611 |
| 8.45 | 10.58817 | -0.0152803 | 0.0198092 | -0.5762365 |
| 8.45 | 121.12362 | 0.0055897 | 0.0156054 | 0.2130048 |
| 8.45 | 384.88010 | 0.0151090 | 0.0148981 | 0.5785004 |
| 8.45 | 476.46780 | 0.0171081 | 0.0149814 | 0.6556999 |
| 8.45 | 457091.44411 | 0.0746479 | 0.0284644 | 2.9451779 |
| 17.90 | 10.58817 | 0.0033618 | 0.0181781 | 0.1279637 |
| 17.90 | 121.12362 | 0.0149017 | 0.0152801 | 0.5705052 |
| 17.90 | 384.88010 | 0.0202842 | 0.0146510 | 0.7786707 |
| 17.90 | 476.46780 | 0.0213703 | 0.0148297 | 0.8208090 |
| 17.90 | 457091.44411 | 0.0533866 | 0.0227154 | 2.0838213 |

```
caption <- paste0("Predicted changes in core body temperature after stress exposure",
    "according to relative resting energy expenditure and various air temperaturess. ",
    "Predictions are based upon average body mass, time-post-stress-exposure (1683 s), ",
    "mixed age groupings, and assume baseline body temperature of 38°C.")

as.data.frame(predict(Core_Model_New, newdata = Descriptive_MR_Cold,
    re_form = NA)[, , "lRR"]) %>%
    mutate(ResMR_Bin = c(rep("Low", 5), rep("High", 5)), AT = rep(seq(-19.9,
        17.9, length.out = 5), 2)) %>%
    group_by(AT, ResMR_Bin) %>%
    summarise(Estimate = mean(Estimate), Error = mean(Est.Error)) %>%
    arrange(ResMR_Bin) %>%
    mutate(Delta_Tb = exp(Estimate) * 38 - 38) %>%
    select(`Air Temperature (°C)` = AT, `Residual RMR Group` = ResMR_Bin,
        Estimate, Error, `Delta Body Temperature (°C)` = Delta_Tb) %>%
    kbl(., longtable = T, booktabs = T, caption = caption) %>%
    kable_styling(latex_options = "striped")
```

Table 16: Predicted changes in core body temperature after stress exposureaccording to relative resting energy expenditure and various air temperaturess. Predictions are based upon average body mass, time-post-stress-exposure (1683 s), mixed age groupings, and assume baseline body temperature of 38°C.

| Air Temperature (°C) | Residual RMR Group | Estimate | Error | Delta Body Temperature (°C) |
|---|---|---|---|---|
| -19.90 | High | -0.0360584 | 0.0253183 | -1.3458105 |
| -10.45 | High | -0.0233986 | 0.0217911 | -0.8788235 |
| -1.00 | High | -0.0110785 | 0.0184784 | -0.4186586 |
| 8.45 | High | 0.0019764 | 0.0161967 | 0.0751787 |
| 17.90 | High | 0.0143912 | 0.0153979 | 0.5508209 |
| -19.90 | Low | 0.0161197 | 0.0264142 | 0.6175123 |
| -10.45 | Low | 0.0176110 | 0.0224715 | 0.6751468 |
| -1.00 | Low | 0.0187922 | 0.0191405 | 0.7208555 |
| 8.45 | Low | 0.0200864 | 0.0166087 | 0.7709989 |
| 17.90 | Low | 0.0213774 | 0.0151535 | 0.8210867 |

```
## Assessing responses below 10˚C

Mass_Quant <- as.numeric(quantile(Core_Data_MR_Shortened$Centred_logMass))
```

```
Descriptive_Mass_VeryCold <- expand.grid(Centred_logMass = Mass_Quant,
    logMass_SD = c(1e-04), Centred_AT = seq(-19.9, 10, length.out = 5) -
        mean(Core_Data_MR_Shortened$Ambient.Temp.Stressor), Ambient.Temp.Stressor.sd = c(1e-04),
    ResMR_Bin = 0, Technique = 0, Centred_Time = 0, Age_Class = 0,
    lRV = c(1e-04))
rm(Mass_Quant)

Descriptive_MR_VeryCold <- expand.grid(Centred_logMass = 0, logMass_SD = c(1e-04),
    Centred_AT = seq(-19.9, 10, length.out = 5) - mean(Core_Data_MR_Shortened$Ambient.Temp.Stressor),
    Ambient.Temp.Stressor.sd = c(1e-04), ResMR_Bin = c(-0.5,
        0.5), Technique = 0, Centred_Time = 0, Age_Class = 0,
    lRV = c(1e-04))

caption <- paste0("Predicted changes in core body temperature after stress exposure at",
    "various air temperatures. Predictions are based upon body mass quantiles, ",
    "average time-post-stress-exposure (1683 s), mixed age groupings, and assume ",
    "baseline body temperature of 38°C.")

as.data.frame(predict(Core_Model_New, newdata = Descriptive_Mass_VeryCold,
    re_form = NA)[, , "lRR"]) %>%
    mutate(AT = rep(seq(-19.9, 10, length.out = 5), each = 5),
        CLM = rep(as.numeric(quantile(Core_Data_MR_Shortened$Centred_logMass)),
            5)) %>%
    group_by(AT, CLM) %>%
    summarise(Estimate = mean(Estimate), Error = mean(Est.Error)) %>%
    mutate(Delta_Tb = exp(Estimate) * 38 - 38, CLM = exp(CLM +
        mean(Core_Data_MR_Shortened$logMass))) %>%
    select(`Air Temperature (°C)` = AT, `Body Mass (g)` = CLM,
        Estimate, Error, `Delta Body Temperature (°C)` = Delta_Tb) %>%
    kbl(., longtable = T, booktabs = T, caption = caption) %>%
    kable_styling(latex_options = "striped")
```

Table 17: Predicted changes in core body temperature after stress exposure atvarious air temperatures. Predictions are based upon body mass quantiles, average time-post-stress-exposure (1683 s), mixed age groupings, and assume baseline body temperature of 38°C.

| Air Temperature (°C) | Body Mass (g) | Estimate | Error | Delta Body Temperature (°C) |
|---:|---:|---:|---:|---:|
| -19.900 | 10.58817 | -0.0708693 | 0.0324063 | -2.5998209 |
| -19.900 | 121.12362 | -0.0228690 | 0.0207303 | -0.8591619 |
| -19.900 | 384.88010 | -0.0002514 | 0.0202329 | -0.0095502 |
| -19.900 | 476.46780 | 0.0038560 | 0.0208747 | 0.1468124 |
| -19.900 | 457091.44411 | 0.1386906 | 0.0652075 | 5.6532095 |
| -12.425 | 10.58817 | -0.0561592 | 0.0285690 | -2.0752320 |
| -12.425 | 121.12362 | -0.0153142 | 0.0189139 | -0.5775076 |
| -12.425 | 384.88010 | 0.0037015 | 0.0183638 | 0.1409181 |
| -12.425 | 476.46780 | 0.0074049 | 0.0187306 | 0.2824293 |
| -12.425 | 457091.44411 | 0.1217884 | 0.0545863 | 4.9215709 |
| -4.950 | 10.58817 | -0.0414336 | 0.0246455 | -1.5423046 |
| -4.950 | 121.12362 | -0.0080537 | 0.0174437 | -0.3048117 |
| -4.950 | 384.88010 | 0.0077226 | 0.0168110 | 0.2945958 |
| -4.950 | 476.46780 | 0.0108860 | 0.0170530 | 0.4159293 |
| -4.950 | 457091.44411 | 0.1049670 | 0.0441025 | 4.2056123 |
| 2.525 | 10.58817 | -0.0267988 | 0.0218784 | -1.0048303 |
| 2.525 | 121.12362 | -0.0005707 | 0.0164113 | -0.0216816 |
| 2.525 | 384.88010 | 0.0121463 | 0.0158043 | 0.4643728 |
| 2.525 | 476.46780 | 0.0141879 | 0.0156136 | 0.5429813 |
| 2.525 | 457091.44411 | 0.0881036 | 0.0348112 | 3.4998473 |
| 10.000 | 10.58817 | -0.0121773 | 0.0194085 | -0.4599296 |
| 10.000 | 121.12362 | 0.0070486 | 0.0156782 | 0.2687934 |
| 10.000 | 384.88010 | 0.0161609 | 0.0149024 | 0.6191033 |
| 10.000 | 476.46780 | 0.0177790 | 0.0146856 | 0.6816421 |
| 10.000 | 457091.44411 | 0.0714270 | 0.0269704 | 2.8135086 |

```
caption <- paste0("Predicted changes in core body temperature after stress exposure",
    "according to relative resting energy expenditure and various air temperaturess. ",
    "Predictions are based upon average body mass, time-post-stress-exposure (1683 s), ",
    "mixed age groupings, and assume baseline body temperature of 38°C.")

as.data.frame(predict(Core_Model_New, newdata = Descriptive_MR_VeryCold,
    re_form = NA)[, , "lRR"]) %>%
    mutate(ResMR_Bin = rep(c("Low", "High"), each = 5), AT = rep(seq(-19.9,
        10, length.out = 5), 2)) %>%
    group_by(AT, ResMR_Bin) %>%
    summarise(Estimate = mean(Estimate), Error = mean(Est.Error)) %>%
    arrange(ResMR_Bin) %>%
    mutate(Delta_Tb = exp(Estimate) * 38 - 38) %>%
    select(`Air Temperature (°C)` = AT, `Residual RMR Group` = ResMR_Bin,
        Estimate, Error, `Delta Body Temperature (°C)` = Delta_Tb) %>%
    kbl(., longtable = T, booktabs = T, caption = caption) %>%
    kable_styling(latex_options = "striped")
```

Table 18: Predicted changes in core body temperature after stress exposureaccording to relative resting energy expenditure and various air temperaturess. Predictions are based upon average body mass, time-post-stress-exposure (1683 s), mixed age groupings, and assume baseline body temperature of 38°C.

| Air Temperature (°C) | Residual RMR Group | Estimate | Error | Delta Body Temperature (°C) |
|---:|---|---:|---:|---:|
| -19.900 | High | -0.0361905 | 0.0254786 | -1.3506501 |
| -12.425 | High | -0.0260172 | 0.0225166 | -0.9759050 |
| -4.950 | High | -0.0164043 | 0.0199465 | -0.6182772 |
| 2.525 | High | -0.0059780 | 0.0175850 | -0.2264856 |
| 10.000 | High | 0.0040354 | 0.0156889 | 0.1536569 |
| -19.900 | Low | 0.0167315 | 0.0265095 | 0.6411449 |
| -12.425 | Low | 0.0172953 | 0.0236002 | 0.6629374 |
| -4.950 | Low | 0.0184261 | 0.0202982 | 0.7066811 |
| 2.525 | Low | 0.0194765 | 0.0179930 | 0.7473632 |
| 10.000 | Low | 0.0203438 | 0.0162046 | 0.7809810 |

```
## Assessing trends at high ambient temperatures

masses = as.numeric(quantile(Core_Data_MR_Shortened$Centred_logMass))
Descriptive_Mass_Warm <- expand.grid(Centred_logMass = masses,
    logMass_SD = c(1e-04), Centred_AT = seq(20, 31, length.out = 5) -
        mean(Core_Data_MR_Shortened$Ambient.Temp.Stressor), Ambient.Temp.Stressor.sd = c(1e-04),
    ResMR_Bin = 0, Technique = 0, Centred_Time = 0, Age_Class = 0,
    lRV = c(1e-04))
rm(masses)

Descriptive_MR_Warm <- expand.grid(Centred_logMass = 0, logMass_SD = c(1e-04),
    Centred_AT = seq(20, 31, length.out = 5) - mean(Core_Data_MR_Shortened$Ambient.Temp.Stressor),
    Ambient.Temp.Stressor.sd = c(1e-04), ResMR_Bin = c(-0.5,
        0.5), Technique = 0, Centred_Time = 0, Age_Class = 0,
    lRV = c(1e-04))

caption <- paste0("Predicted changes in core body temperature after stress exposure at",
    "various air temperatures. Predictions are based upon body mass quantiles, ",
    "average time-post-stress-exposure (1683 s), mixed age groupings, and assume ",
    "baseline body temperature of 38°C.")

masses = rep(as.numeric(quantile(Core_Data_MR_Shortened$Centred_logMass)),
    5)
as.data.frame(predict(Core_Model_New, newdata = Descriptive_Mass_Warm,
    re_form = NA)[, , "lRR"]) %>%
    mutate(AT = rep(seq(20, 31, length.out = 5), each = 5), CLM = masses) %>%
    group_by(AT, CLM) %>%
    summarise(Estimate = mean(Estimate), Error = mean(Est.Error)) %>%
    mutate(Delta_Tb = exp(Estimate) * 38 - 38, CLM = exp(CLM +
        mean(Core_Data_MR_Shortened$logMass))) %>%
```

```
    select(`Air Temperature (°C)` = AT, `Body Mass (g)` = CLM,
        Estimate, Error, `Delta Body Temperature (°C)` = Delta_Tb) %>%
    kbl(., longtable = T, booktabs = T, caption = caption) %>%
    kable_styling(latex_options = "striped")
```

Table 19: Predicted changes in core body temperature after stress exposure atvarious air temperatures. Predictions are based upon body mass quantiles, average time-post-stress-exposure (1683 s), mixed age groupings, and assume baseline body temperature of 38°C.

| Air Temperature (°C) | Body Mass (g) | Estimate | Error | Delta Body Temperature (°C) |
|---:|---:|---:|---:|---:|
| 20.00 | 10.58817 | 0.0076724 | 0.0182738 | 0.2926714 |
| 20.00 | 121.12362 | 0.0169167 | 0.0154739 | 0.6483040 |
| 20.00 | 384.88010 | 0.0213679 | 0.0147650 | 0.8207178 |
| 20.00 | 476.46780 | 0.0223815 | 0.0145070 | 0.8600854 |
| 20.00 | 457091.44411 | 0.0486143 | 0.0226444 | 1.8929835 |
| 22.75 | 10.58817 | 0.0131304 | 0.0182278 | 0.5022462 |
| 22.75 | 121.12362 | 0.0198146 | 0.0153010 | 0.7604630 |
| 22.75 | 384.88010 | 0.0227121 | 0.0148960 | 0.8729354 |
| 22.75 | 476.46780 | 0.0235402 | 0.0149222 | 0.9051381 |
| 22.75 | 457091.44411 | 0.0425107 | 0.0232271 | 1.6502337 |
| 25.50 | 10.58817 | 0.0182922 | 0.0187626 | 0.7015019 |
| 25.50 | 121.12362 | 0.0223745 | 0.0154560 | 0.8598154 |
| 25.50 | 384.88010 | 0.0242662 | 0.0149648 | 0.9333930 |
| 25.50 | 476.46780 | 0.0245833 | 0.0149292 | 0.9457426 |
| 25.50 | 457091.44411 | 0.0362354 | 0.0246821 | 1.4021962 |
| 28.25 | 10.58817 | 0.0238716 | 0.0187816 | 0.9180331 |
| 28.25 | 121.12362 | 0.0252625 | 0.0157234 | 0.9722018 |
| 28.25 | 384.88010 | 0.0259560 | 0.0150750 | 0.9992394 |
| 28.25 | 476.46780 | 0.0259741 | 0.0152364 | 0.9999449 |
| 28.25 | 457091.44411 | 0.0298738 | 0.0264592 | 1.1523313 |
| 31.00 | 10.58817 | 0.0292583 | 0.0196921 | 1.1282409 |
| 31.00 | 121.12362 | 0.0280517 | 0.0160318 | 1.0810582 |
| 31.00 | 384.88010 | 0.0274366 | 0.0155899 | 1.0570265 |
| 31.00 | 476.46780 | 0.0274075 | 0.0155053 | 1.0558879 |
| 31.00 | 457091.44411 | 0.0236332 | 0.0291992 | 0.9087583 |

```
rm(masses)

caption <- paste0("Predicted changes in core body temperature after stress exposure",
    "according to relative resting energy expenditure and various air temperaturess. ",
    "Predictions are based upon average body mass, time-post-stress-exposure (1683 s), ",
    "mixed age groupings, and assume baseline body temperature of 38°C.")

as.data.frame(predict(Core_Model_New, newdata = Descriptive_MR_Warm,
    re_form = NA)[, , "lRR"]) %>%
    mutate(ResMR_Bin = rep(c("Low", "High"), each = 5), AT = rep(seq(20,
        31, length.out = 5), 2)) %>%
    group_by(AT, ResMR_Bin) %>%
    summarise(Estimate = mean(Estimate), Error = mean(Est.Error)) %>%
    arrange(ResMR_Bin) %>%
    mutate(Delta_Tb = exp(Estimate) * 38 - 38) %>%
    select(`Air Temperature (°C)` = AT, `Residual RMR Group` = ResMR_Bin,
        Estimate, Error, `Delta Body Temperature (°C)` = Delta_Tb) %>%
    kbl(., longtable = T, booktabs = T, caption = caption) %>%
    kable_styling(latex_options = "striped")
```

Table 20: Predicted changes in core body temperature after stress exposureaccording to relative resting energy expenditure and various air temperaturess. Predictions are based upon average body mass, time-post-stress-exposure (1683 s), mixed age groupings, and assume baseline body temperature of 38°C.

| Air Temperature (°C) | Residual RMR Group | Estimate | Error | Delta Body Temperature (°C) |
|---|---|---|---|---|

| | | | | |
|---|---|---|---|---|
| 20.00 | High | 0.0173282 | 0.0153456 | 0.6642079 |
| 22.75 | High | 0.0211567 | 0.0153392 | 0.8125204 |
| 25.50 | High | 0.0245838 | 0.0156722 | 0.9457626 |
| 28.25 | High | 0.0281301 | 0.0158322 | 1.0841190 |
| 31.00 | High | 0.0320872 | 0.0163694 | 1.2390849 |
| 20.00 | Low | 0.0215223 | 0.0151416 | 0.8267125 |
| 22.75 | Low | 0.0221029 | 0.0152161 | 0.8492617 |
| 25.50 | Low | 0.0223343 | 0.0154615 | 0.8582509 |
| 28.25 | Low | 0.0228554 | 0.0159430 | 0.8785062 |
| 31.00 | Low | 0.0232046 | 0.0162563 | 0.8920849 |

```
rm(Descriptive_Mass_Cold, Descriptive_MR_Cold, Descriptive_Mass_VeryCold,
    Descriptive_MR_VeryCold, Descriptive_Mass_Warm, Descriptive_MR_Warm)

# Checking mean baseline body temperatures between
# expenditure groups

Core_Data_MR_Shortened %>%
    mutate(ResMR_Bin = ifelse(ResMR_Bin == -0.5, "Low", "High")) %>%
    group_by(ResMR_Bin) %>%
    summarise(`Mean Baseline\nTb` = mean(Initial.Tb, na.rm = T),
        SD = sd(Initial.Tb, na.rm = T)) %>%
    rename(`Relative Resting\nMetabolic Rate` = ResMR_Bin)
```

```
## # A tibble: 2 x 3
##   `Relative Resting\nMetabolic Rate` `Mean Baseline\nTb`    SD
##   <chr>                                            <dbl> <dbl>
## 1 High                                              38.3  1.75
## 2 Low                                               37.5  1.70
```

```
# Calculating percentage of birds and mammals in high and
# low metabolic expenditure groupings.

Core_Data_MR_Shortened %>%
    mutate(ResMR_Bin = ifelse(ResMR_Bin == -0.5, "Low", "High")) %>%
    group_by(ResMR_Bin, Class, Species.Name) %>%
    count() %>%
    group_by(ResMR_Bin, Class) %>%
    count() %>%
    ungroup() %>%
    group_by(ResMR_Bin) %>%
    summarise(`%` = n/sum(n)) %>%
    ungroup() %>%
    mutate(Class = rep(unique(Core_Data_MR_Shortened$Class),
        2)) %>%
    select(ResMR_Bin, Class, `%`) %>%
    rename(`Relative Resting\nMetabolic Rate` = ResMR_Bin)
```

```
## # A tibble: 4 x 3
##   `Relative Resting\nMetabolic Rate` Class      `%`
##   <chr>                              <chr>    <dbl>
## 1 High                               Aves     0.571
## 2 High                               Mammalia 0.429
## 3 Low                                Aves     0.308
## 4 Low                                Mammalia 0.692
```

```
# Evaluating average change in body temperature by class

Core_Data_MR_Shortened %>%
    group_by(Class) %>%
    summarise(`Mean Change in\nBody Temperature (°C)` = mean(Delta.Tb,
        na.rm = T), SD = sd(Delta.Tb, na.rm = T))
```

```
## # A tibble: 2 x 3
##   Class    `Mean Change in\nBody Temperature (°C)`    SD
##   <chr>                                      <dbl> <dbl>
## 1 Aves                                       0.176 1.61
## 2 Mammalia                                   1.01  0.906
```

```
# And again, while setting body mass and ambient
# temperature at a mean

expand.grid(Centred_AT = 0, Ambient.Temp.Stressor.sd = 1e-04,
    Centred_logMass = 0, logMass_SD = 1e-04, Age_Class = 0, ResMR_Bin = 0,
    Centred_Time = 0, lRV = 1e-04, Technique = 0, Species.Name = unique(Core_Data_MR_Shortened$Species.Name)) %>%
    left_join(., Core_Data_MR_Shortened %>%
        select(Species.Name, Phylo), by = "Species.Name") %>%
    mutate(Fit = predict(Core_Model_New, newdata = ., newdata2 = list(A = Core_VCV_MR),
        re_formula = ~(1 | Species.Name) + (1 | gr(Phylo, cov = A)),
        allow_new_levels = TRUE, ndraws = 2500, )[, "Estimate",
        "lRR"]) %>%
    left_join(., Core_Data_MR_Shortened %>%
        select(Species.Name, Phylo, Class), by = c("Species.Name",
        "Phylo")) %>%
    group_by(Class) %>%
    summarise(`Mean Change in\nBody Temperature (°C)` = mean(Fit),
        SD = sd(Fit))
```

```
## # A tibble: 2 x 3
##   Class    `Mean Change in\nBody Temperature (°C)`      SD
##   <chr>                                       <dbl>   <dbl>
## 1 Aves                                       0.0129 0.00790
## 2 Mammalia                                   0.0329 0.00311
```

```
# Identifying maximum change points

Core_Data_MR_Shortened %>%
    filter(lRR %in% c(max(Core_Data_MR_Shortened$lRR), min(Core_Data_MR_Shortened$lRR))) %>%
    select(Species.Name, Class, `Body Mass (g)` = Mass, `Ta (°C)` = Ambient.Temp.Stressor,
        `Delta Tb (°C)` = Delta.Tb, lRR = lRR)
```

```
##          Species.Name    Class Body Mass (g) Ta (°C) Delta Tb (°C)        lRR
## 1 Aepyceros melampus Mammalia      42330.00      16          4.25  0.1058596
## 2         Parus major     Aves         20.25       2         -4.20 -0.1145666
```

```
# Plotting predicted trends across ambient temperature by
# class

predDataBase = expand.grid(Centred_AT = seq(min(Core_Data_MR_Shortened$Ambient.Temp.Stressor,
    na.rm = T), max(Core_Data_MR_Shortened$Ambient.Temp.Stressor,
    na.rm = T), by = 2.5) - mean(Core_Data_MR_Shortened$Ambient.Temp.Stressor),
    Ambient.Temp.Stressor.sd = 1e-04, Centred_logMass = 0, logMass_SD = 1e-04,
    Age_Class = 0, ResMR_Bin = 0, Centred_Time = 0, lRV = 1e-04,
    Technique = 0, Species.Name = unique(Core_Data_MR_Shortened$Species.Name)) %>%
    left_join(., Core_Data_MR_Shortened %>%
        select(Species.Name, Phylo), by = "Species.Name")
predData = cbind(predDataBase, predict(Core_Model_New, newdata = predDataBase,
    newdata2 = list(A = Core_VCV_MR), re_formula = ~(1 | Species.Name) +
        (1 | gr(Phylo, cov = A)), allow_new_levels = TRUE, ndraws = 5000)) %>%
    left_join(., Core_Data_MR_Shortened %>%
        select(Species.Name, Phylo, Class), by = c("Species.Name",
        "Phylo")) %>%
    mutate(Ambient = Centred_AT + mean(Core_Data_MR_Shortened$Ambient.Temp.Stressor))

ggplot(predData %>%
    group_by(Class, Ambient) %>%
    mutate(delta = exp(mean(Estimate.lRR)) * 38 - 38, UCL = exp(mean(Q2.5.lRR)) *
        38 - 38, LCL = exp(mean(Q97.5.lRR)) * 38 - 38), aes(x = Ambient,
    y = delta, linetype = Class)) + geom_ribbon(mapping = aes(x = Ambient,
    ymin = LCL, ymax = UCL, fill = Class), alpha = 0.7) + geom_smooth(colour = "black",
    se = FALSE) + theme_classic() + ylab("Change in Body Temperature (°C)") +
    xlab("Ambient Temperature (°C)") + scale_fill_manual(values = wes_palette("Rushmore",
    type = "continuous", 10)[c(2, 8)])
```

```
predData %>%
    mutate(delta = exp(Estimate.lRR) * 38 - 38) %>%
    ggplot(aes(x = delta, fill = Class)) + geom_density(adjust = 3) +
    geom_vline(xintercept = 0, colour = "black", linetype = "dashed") +
```

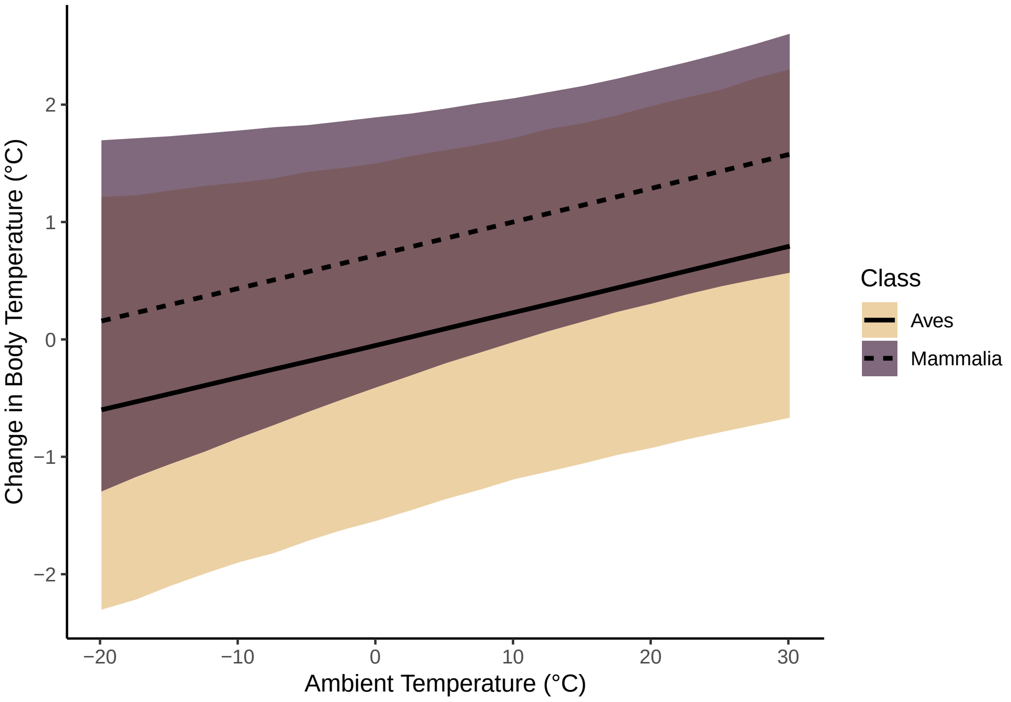


Figure 50: Predicted effect of ambient temperature (°C) on stress-induced changes in body temperature across birds and mammals. Predictions are derived from a revised Bayesian, linear mixed effects model predicting stress-induced changes in body temperature (natural log-transformed response ratios). Predicted trends (dashed and solid black lines) are marginalised (i.e. assume mean values for all other model parameters). Ribbons represent 95 percent credible intervals around trends.

```
    scale_fill_manual(values = wes_palette("Rushmore", type = "continuous",
        10)[c(2, 8)]) + theme_classic() + xlab("Change in Body Temperature (°C)")
```

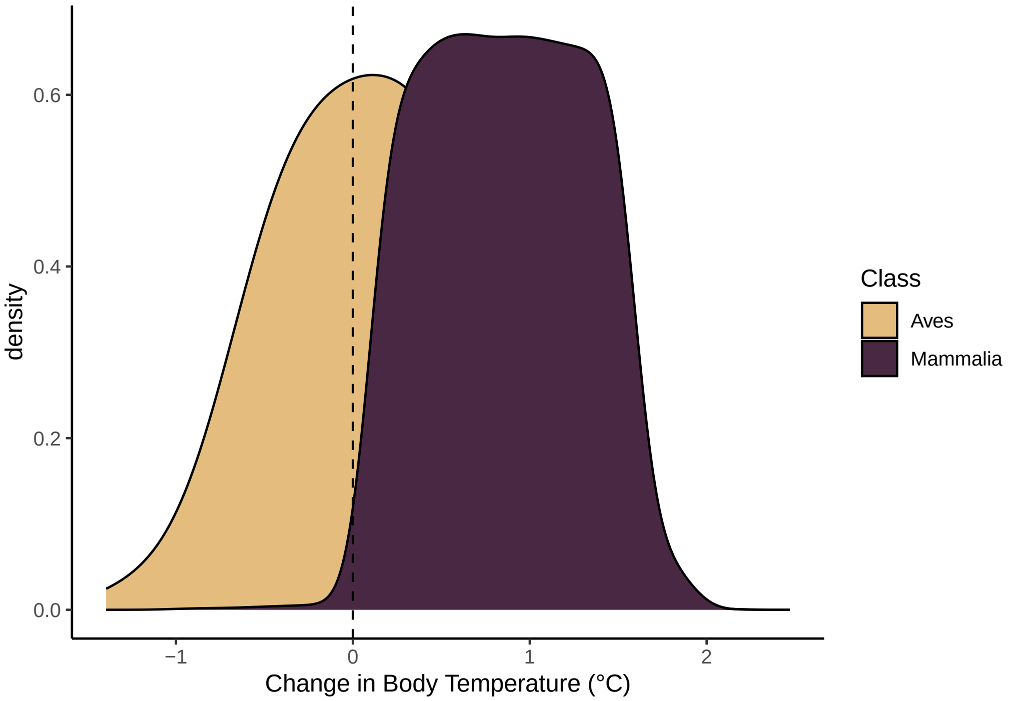


Figure 51: Density of predicted stress-induced changes in body temperature across birds and mammals. Predictions are derived from a revised Bayesian, linear mixed effects model predicting stress-induced changes in body temperature (natural log-transformed response ratios). The dashed vertical line indicates 0.

```
# Checking whether decrease in body temperature is more
# common among birds than mammals

posteriors = posterior_predict(Core_Model_New, newdata = predDataBase,
    newdata2 = list(A = Core_VCV_MR), re_formula = ~(1 | Species.Name) +
        (1 | gr(Phylo, cov = A)), allow_new_levels = TRUE, ndraws = 100,
    resp = "lRR")

predDataBase$declines = apply(posteriors, MARGIN = 2, FUN = function(x) {
    length(which(x < 0))
})
predDataBase$draws = 100

MO = "/Users/joshuatabh/Documents/researchProjects/trent/sihMetaregression/models/classModel.Rds"

classModel = brm(declines | trials(draws) ~ Class, data = predDataBase %>%
    left_join(., Core_Data_MR_Shortened %>%
        select(Species.Name, Phylo, Class) %>%
        distinct(), by = c("Species.Name", "Phylo")), cores = 1,
    chains = 4, seed = 200, silent = TRUE, refresh = 0, family = "binomial",
    iter = 15000, warmup = 2500, thin = 10, control = list(adapt_delta = 0.95,
        max_treedepth = 13), file = MO)

hypothesis(hypothesis = "ClassMammalia - Intercept < 0", class = "b",
    classModel)
```

```
## Hypothesis Tests for class b:
##                 Hypothesis Estimate Est.Error CI.Lower CI.Upper Evid.Ratio
## 1 (ClassMammalia-In... < 0    -1.16      0.02    -1.19    -1.13        Inf
##   Post.Prob Star
## 1         1    *
```

```
## ---
## 'CI': 90%-CI for one-sided and 95%-CI for two-sided hypotheses.
## '*': For one-sided hypotheses, the posterior probability exceeds 95%;
## for two-sided hypotheses, the value tested against lies outside the 95%-CI.
## Posterior probabilities of point hypotheses assume equal prior probabilities.
```

```
# Mammals tend to be less likely to decrease their body
# temperatures after stressors, after controlling for all
# other influential parameters, than birds.
```

### 7 | Repeating analyses with bias-corrected log response ratio as a measure of effect size

Although using the log-transformed response ratio as a measure of effect size is convenient for easily interpreting study outcomes, it does not account for biases imposed by small sample sizes (i.e. the over-representation of large effect sizes from studies with small sample size). Here, we sought to ensure that our findings were not influenced by small sample size biases. To do so, we recalculated our response ratio using bias correction methods reported by Lajeunesse (2015) (denoted as the "delta response ratio", or, "RR$^{\Delta}$"). For studies with a paired design (i.e. lacking a control group) our measure of RR$^{\Delta}$ was calculated as follows:

$$RR^{\Delta} = RR_i + \frac{1}{2} \cdot (\frac{\sigma^2_{1,i}}{n_{1,i} \cdot T_b 1^2_i} - \frac{\sigma^2_{0,i}}{n_{0,i} \cdot T_b 0^2_i})$$

where $RR_i$ represents the non-adjusted log response ratio for study $i$, $T_b 1_i$ represents an average minimal, or an average maximal body temperature of study population $i$ that was observed after the onset of a stress exposure treatment, $T_b 0_i$ represents the body temperature of study population $i$ prior to a stress exposure treatment, $n_{1,i}$ and $n_{0,i}$ represents the sample sizes under stress-exposure and baseline conditions respectively, and $\sigma_{n,i}$ represents the standard deviation around $T_b n_i$. Here, the variance around $RR^{\Delta}$ was estimated as follows:

$$var(RR^{\Delta}) = var(RR_i) + \frac{1}{2} \cdot (\frac{\sigma^4_{1,i}}{n^2_{1,i} \cdot T_b 1^4_i} - \frac{\sigma^4_{0,i}}{n^2_{0,i} \cdot T_b 0^4_i})$$

where all variables remain as previously described. If errors were only reported around a stress-induced change in body temperature and not around both baseline and stress-induced body temperature measurements, we assumed errors around baseline and stress-induced body temperatures to be equal to that around the stress-induced change in body temperature. For studies that did not use a paired experimental design and instead used a control group, the above equations were replaced with the following:

$$RR^{\Delta} = RR_i + \frac{1}{2} \cdot (\frac{\sigma^2_{1S,i}}{n_{S,i} \cdot T_b 1^2_{Si}} - \frac{\sigma^2_{1C,i}}{n_{C,i} \cdot T_b 1^2_{Ci}})$$

and:

$$var(RR^{\Delta}) = RR_i + \frac{1}{2} \cdot (\frac{\sigma^4_{1S,i}}{n^2_{S,i} \cdot T_b 1^4_{Si}} - \frac{\sigma^4_{1C,i}}{n^2_{C,i} \cdot T_b 1^4_{Ci}})$$

where $T_b 1_{Si}$ represents an average minimal, or an average maximal body temperature of the stress exposed population $i$, observed after the onset of a stress exposure treatment, $T_b 1_{Ci}$ represents the average body temperature of the control population at the equivalent time point, $\sigma_{1S,i}$ and $\sigma_{1C,i}$ represents the standard deviation around $T_b 1_{Si}$ and $T_b 1_{Ci}$ respectively, and $n_{S,i}$ and $n_{C,i}$ represents the sample sizes across observations $T_b 1_{Si}$ and $T_b 1_{Ci}$ respectively.

```
{
    Core_Data_AlRR <- Core_Data_MR_Shortened
    Core_Data_AlRR$Control.Sample.Size[c(which(Core_Data_AlRR$StudyID ==
        "F5"))] <- 9

    Core_Data_AlRR <- Core_Data_AlRR %>%
        mutate(SD1 = Peak.Tb.SE * sqrt(Exp.Sample.Size), SD2 = ifelse(Unmanipulated.Control.Y.N. ==
            "N", Initial.Tb.SE * sqrt(Exp.Sample.Size), Control.Tb.at.Stress.Peak.SE *
            sqrt(Control.Sample.Size))) %>%
        mutate(SD1 = ifelse(is.na(SD1) & !is.na(Delta.Tb.SE),
            Delta.Tb.SE, SD1)) %>%
        mutate(SD2 = ifelse(is.na(SD2) & !is.na(Delta.Tb.SE),
            Delta.Tb.SE, SD2)) %>%
        mutate(lRR_Delta = ifelse(Unmanipulated.Control.Y.N. ==
            "N", lRR + 0.5 * (SD1^2/(Exp.Sample.Size * Peak.Trough.Tb^2) -
            SD2^2/(Exp.Sample.Size * Initial.Tb^2)), ifelse(!is.na(Control.Tb.at.Stress.Peak),
            lRR + 0.5 * (SD1^2/(Exp.Sample.Size * Peak.Trough.Tb^2) -
                SD2^2/(Control.Sample.Size * Control.Tb.at.Stress.Peak^2)),
            lRR + 0.5 * (SD1^2/(Exp.Sample.Size * Peak.Trough.Tb^2) -
                SD2^2/(Exp.Sample.Size * Initial.Tb^2))))) %>%
        mutate(lRV_Delta = ifelse(Unmanipulated.Control.Y.N. ==
            "N", lRV + 0.5 * (SD1^4/(Exp.Sample.Size^2 * Peak.Trough.Tb^4) -
            SD2^4/(Exp.Sample.Size^2 * Initial.Tb^4)), ifelse(!is.na(Control.Tb.at.Stress.Peak),
            lRV + 0.5 * (SD1^4/(Exp.Sample.Size^2 * Peak.Trough.Tb^4) -
                SD2^4/(Control.Sample.Size^2 * Control.Tb.at.Stress.Peak^4)),
            lRV + 0.5 * (SD1^4/(Exp.Sample.Size^2 * Peak.Trough.Tb^4) -
                SD2^4/(Exp.Sample.Size^2 * Initial.Tb^4)))))

    # Plotting adjusted lRR against original lRR

    ggplot(Core_Data_AlRR, aes(x = lRR, y = lRR_Delta)) + geom_point(size = 3,
        pch = 21, colour = "black", fill = nice_pink, alpha = 0.7) +
        theme_classic() + xlab("log Response Ratio") + ylab("Adjusted log Response Ratio")

    # Adding errors

    ggplot(Core_Data_AlRR, aes(x = lRR, y = lRR_Delta)) + geom_errorbarh(aes(y = lRR_Delta,
        xmin = lRR - lRV, xmax = lRR + lRV), colour = "black") +
        geom_errorbar(aes(x = lRR, ymin = lRR_Delta - lRV_Delta,
            ymax = lRR_Delta + lRV_Delta), colour = "black") +
        geom_point(size = 3, pch = 21, colour = "black", fill = nice_pink,
            alpha = 0.7) + theme_classic() + xlab("log Response Ratio") +
        ylab("Adjusted log Response Ratio")

}
```

Following calculations, we then re-ran our previously described model but replacing the log response ratio with the bias-corrected log response ratio ($RR^{\Delta}$) as the response variable. After, we compared coefficients visually using paired forest plots (provided below).

Importantly, our model using $RR^{\Delta}$ as a response variable was not scrutinised to the same extent as that using the unadjusted log response ratio as a response variable. We chose to forgo this scrutinisation because we did not want our coefficient comparisons to be influenced by differences in model structure, or by sample inclusions, that may occur following after-the-fact model adjustments. Furthermore, we were primarily interested in broadly comparing coefficient estimates between models using each effect size metric as a response variable.

Priors, number of sample iterations, burn-in values, and thinning values for our model with $RR^{\Delta}$ as a response variable remained identical to those used for our model with the unadjusted log response ratio as our response variable.

```
# First, correcting model formula
```

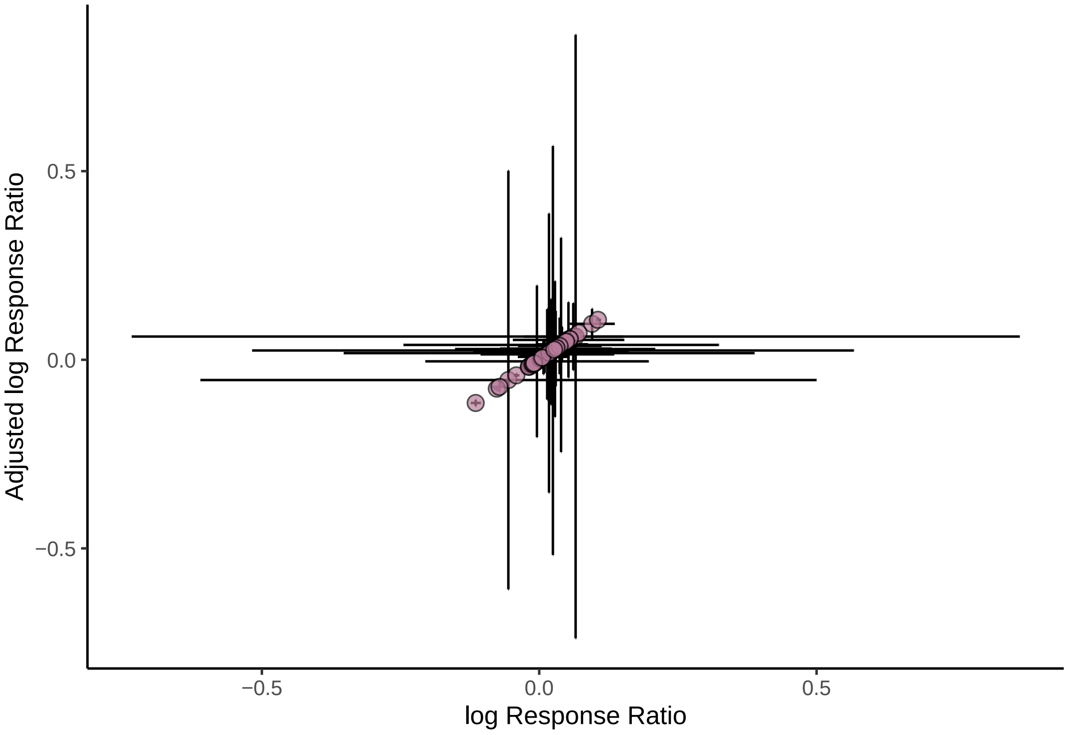


Figure 52: Stress-induced changes in body temperature represented as natural log-transformed response ratios against those represents as adjusted (delta; 'bias-corrected') log-transformed response ratios. Dots indicate mean values per observation and errorbars represent standard errors around means.

```
g(E1, E2, E3) %=% list(bf(paste(Core_Model_New$formula$forms$CentredlogMass)[1]),
    bf(paste(Core_Model_New$formula$forms$CentredAT)[1]), bf(gsub(pattern = ".*~",
        replacement = "lRR_Delta | se(lRV_Delta, sigma = TRUE) ~",
        paste(Core_Model_New$formula$forms$lRR)[1])))

form_lRRDelta <- E1 + E2 + E3

# Next, adjusting response variable names in our priors.

prior_Core_lRRDelta <- c(set_prior("gamma(1, 1)", class = "sd",
    resp = "CentredAT"), set_prior("gamma(1, 1)", class = "sd",
    resp = "CentredlogMass"), set_prior("normal(0, 0.5)", class = "Intercept",
    resp = "lRRDelta"), set_prior("normal(0, 0.1)", class = "b",
    coef = "miCentred_AT", resp = "lRRDelta"), set_prior("normal(0, 0.1)",
    class = "b", coef = "miCentred_logMass", resp = "lRRDelta"),
    set_prior("normal(0, 0.05)", class = "b", coef = "miCentred_AT:ResMR_Bin",
        resp = "lRRDelta"), set_prior("normal(0, 0.1)", class = "b",
        coef = "Age_Class", resp = "lRRDelta"), set_prior("skew_normal(0, 0.1, -2.1)",
        class = "b", coef = "Technique", resp = "lRRDelta"),
    set_prior("skew_normal(0, 0.1, -2.1)", class = "b", coef = "ResMR_Bin",
        resp = "lRRDelta"), set_prior("normal(0, 0.05)", class = "b",
        coef = "polyCentred_Time21", resp = "lRRDelta"), set_prior("normal(0, 0.1)",
        class = "b", coef = "polyCentred_Time22", resp = "lRRDelta"),
    set_prior("exponential(7.5)", class = "sd", resp = "lRRDelta"),
    set_prior("gamma(1,2)", class = "sigma", resp = "lRRDelta"))

# Executing model

Model_Output <- paste0("/Users/joshuatabh/Documents/researchProjects/trent/sihMetaregression/models/",
    "lRRDelta.Rds")
```

```
lRR_Delta_Core <- brm(formula = form_lRRDelta, prior = prior_Core_lRRDelta,
    data = Core_Data_AlRR, data2 = Core_Model_New$data2, cores = 1,
    chains = 2, seed = 100, family = "gaussian", iter = 15000,
    warmup = 2500, thin = 10, control = list(adapt_delta = 0.97,
        max_treedepth = 14), save_pars = save_pars(latent = TRUE),
    file = Model_Output)

rm(Model_Output)

# Printing posterior modes and HPD intervals (80% and 95%)
# alongside model using log response ratio as a response
# variable

lRR_Delta_Core_Table <- lRR_Delta_Core %>%
    spread_draws(b_lRRDelta_Intercept, bsp_lRRDelta_miCentred_AT,
        b_lRRDelta_ResMR_Bin, b_lRRDelta_polyCentred_Time21,
        b_lRRDelta_polyCentred_Time22, bsp_lRRDelta_miCentred_logMass,
        b_lRRDelta_Technique, b_lRRDelta_Age_Class, `bsp_lRRDelta_miCentred_AT:miCentred_logMass`,
        `bsp_lRRDelta_miCentred_AT:ResMR_Bin`, sd_Phylo__lRRDelta_Intercept,
        sd_StudyID__lRRDelta_Intercept, sd_Species.Name__lRRDelta_Intercept,
        sigma_lRRDelta) %>%
    select(-c(.chain, .iteration, .draw)) %>%
    mutate_all(., .funs = round, 5) %>%
    summarise_all(., .funs = md) %>%
    pivot_longer(everything(), names_to = "Parameter", values_to = "Mode") %>%
    left_join(., simple_hdi(Core_Model_New, rnd = 5, cis = c(80,
        95), sci_note = FALSE), by = c("Parameter")) %>%
    right_join(tribble(~Par, ~Parameter, "Intercept", "b_lRRDelta_Intercept",
        "Air Temperature", "bsp_lRRDelta_miCentred_AT", "Residual RMR",
        "b_lRRDelta_ResMR_Bin", "Time (1st order)", "b_lRRDelta_polyCentred_Time21",
        "Time (2nd order)", "b_lRRDelta_polyCentred_Time22",
        "Measurement Technique", "b_lRRDelta_Technique", "Age Class",
        "b_lRRDelta_Age_Class", "log Mass", "bsp_lRRDelta_miCentred_logMass",
        "Air Temperature by log Mass", "bsp_lRRDelta_miCentred_AT:miCentred_logMass",
        "Air Temperature by Residual RMR", "bsp_lRRDelta_miCentred_AT:ResMR_Bin",
        "Phylogeny", "sd_Phylo__lRRDelta_Intercept", "Species",
        "sd_Species.Name__lRRDelta_Intercept", "Study ID", "sd_StudyID__lRRDelta_Intercept",
        "Sigma", "sigma_lRRDelta"), ., by = c("Parameter")) %>%
    select(-c(Parameter)) %>%
    mutate(` ` = ifelse((Low_HDI_80 < 0 & High_HDI_80 < 0) |
        (Low_HDI_80 > 0 & High_HDI_80 > 0), "*", ""), `  ` = ifelse((Low_HDI_95 <
        0 & High_HDI_95 < 0) | (Low_HDI_95 > 0 & High_HDI_95 >
        0), "*", "")) %>%
    select(Par, Mode, Low_HDI_80, High_HDI_80, " ", Low_HDI_95,
        High_HDI_95, "  ") %>%
    rename(Parameter = Par, `Low HDI (80%)` = Low_HDI_80, `High HDI (80%)` = High_HDI_80,
        `Low HDI (95%)` = Low_HDI_95, `High HDI (95%)` = High_HDI_95)
```

```r
Table_Caption <- paste0("Effects of various parameters on stress-induced changes in core body",
    " temperature; results of a Bayesian linear mixed effects model with bias-corrected ",
    "log response ratio (RR Delta) as the response variable.")

lRR_Delta_Core_Table %>%
    kbl(caption = Table_Caption, longtable = T, booktabs = T) %>%
    kable_classic(full_width = TRUE) %>%
    kable_styling(latex_options = "striped")
```

Table 21: Effects of various parameters on stress-induced changes in core body temperature; results of a Bayesian linear mixed effects model with bias-corrected log response ratio (RR Delta) as the response variable.

| Parameter | Mode | Low HDI (80%) | High HDI (80%) | | Low HDI (95%) | High HDI (95%) | |
|---|---|---|---|---|---|---|---|
| Intercept | 0.01463 | NA | NA | NA | NA | NA | NA |
| Air Temperature | 0.00085 | NA | NA | NA | NA | NA | NA |
| Residual RMR | -0.00788 | NA | NA | NA | NA | NA | NA |
| Time (1st order) | -0.02203 | NA | NA | NA | NA | NA | NA |
| Time (2nd order) | -0.04613 | NA | NA | NA | NA | NA | NA |
| Measurement Technique | -0.00039 | NA | NA | NA | NA | NA | NA |
| Age Class | -0.00973 | NA | NA | NA | NA | NA | NA |
| log Mass | 0.00466 | NA | NA | NA | NA | NA | NA |
| Air Temperature by log Mass | -0.00049 | NA | NA | NA | NA | NA | NA |
| Air Temperature by Residual RMR | 0.00135 | NA | NA | NA | NA | NA | NA |
| Phylogeny | 0.00012 | NA | NA | NA | NA | NA | NA |
| Species | 0.02015 | NA | NA | NA | NA | NA | NA |
| Study ID | 0.01420 | NA | NA | NA | NA | NA | NA |
| Sigma | 0.01131 | NA | NA | NA | NA | NA | NA |

```r
Core_Comparison <- rbind(Core_Model_Table %>%
    mutate(`Response Variable` = "log Response Ratio"), lRR_Delta_Core_Table %>%
    mutate(`Response Variable` = "Bias Corrected\nlog Response Ratio")) %>%
    arrange(Parameter, `Response Variable`) %>%
    select(Parameter, `Response Variable`, Mode, `Low HDI (80%)`, `High HDI (80%)`,
        `Low HDI (95%)`, `High HDI (95%)`)

Table_Caption <- "Stress-induced core body temperature responses: Comparison of
coefficients derived from models using the log-response ratio and bias-corrected
log response ratio (RR Delta) as measures of effect size."

Core_Comparison %>%
    mutate(Parameter = ifelse(lead(Parameter, 1) == Parameter, Parameter, "")) %>%
    mutate(Parameter = ifelse(is.na(Parameter), "", Parameter)) %>%
    kbl(caption = Table_Caption, longtable = T, booktabs = T) %>%
    kable_classic(full_width = TRUE) %>%
    kable_styling(latex_options = "striped")
```

Table 22: Stress-induced core body temperature responses: Comparison of coefficients derived from models using the log-response ratio and bias-corrected log response ratio (RR Delta) as measures of effect size.

| Parameter | Response Variable | Mode | Low HDI (80%) | High HDI (80%) | Low HDI (95%) | High HDI (95%) |
|---|---|---|---|---|---|---|

| | | | | | | |
|---|---|---|---|---|---|---|
| Age Class | Bias Corrected log Response Ratio | -0.00973 | NA | NA | NA | NA |
| | log Response Ratio | -0.00697 | -0.01253 | -0.00137 | -0.01619 | 0.00139 |
| Air Temperature | Bias Corrected log Response Ratio | 0.00085 | NA | NA | NA | NA |
| | log Response Ratio | 0.00077 | 0.00031 | 0.00123 | 0.00004 | 0.00144 |
| Air Temperature by Residual RMR | Bias Corrected log Response Ratio | 0.00135 | NA | NA | NA | NA |
| | log Response Ratio | 0.00138 | 0.00026 | 0.00223 | -0.00027 | 0.00279 |
| Air Temperature by log Mass | Bias Corrected log Response Ratio | -0.00049 | NA | NA | NA | NA |
| | log Response Ratio | -0.00040 | -0.00064 | -0.00015 | -0.00078 | -0.00001 |
| Intercept | Bias Corrected log Response Ratio | 0.01463 | NA | NA | NA | NA |
| | log Response Ratio | 0.01673 | 0.00441 | 0.02733 | -0.00178 | 0.03502 |
| Measurement Technique | Bias Corrected log Response Ratio | -0.00039 | NA | NA | NA | NA |
| | log Response Ratio | 0.00048 | -0.00545 | 0.00866 | -0.00932 | 0.01238 |
| Phylogeny | Bias Corrected log Response Ratio | 0.00012 | NA | NA | NA | NA |
| | log Response Ratio | 0.00150 | 0.00000 | 0.00223 | 0.00000 | 0.00325 |
| Residual RMR | Bias Corrected log Response Ratio | -0.00788 | NA | NA | NA | NA |
| | log Response Ratio | -0.00398 | -0.01336 | 0.00343 | -0.01764 | 0.00847 |
| Sigma | Bias Corrected log Response Ratio | 0.01131 | NA | NA | NA | NA |
| | log Response Ratio | 0.01075 | NA | NA | NA | NA |
| Species | Bias Corrected log Response Ratio | 0.02015 | NA | NA | NA | NA |
| | log Response Ratio | 0.01492 | 0.00755 | 0.02941 | 0.00004 | 0.03219 |
| Study ID | Bias Corrected log Response Ratio | 0.01420 | NA | NA | NA | NA |
| | log Response Ratio | 0.01455 | 0.01070 | 0.01807 | 0.00910 | 0.02066 |
| Time (1st order) | Bias Corrected log Response Ratio | -0.02203 | NA | NA | NA | NA |
| | log Response Ratio | -0.03614 | -0.07137 | -0.01370 | -0.08533 | 0.00250 |
| Time (2nd order) | Bias Corrected log Response Ratio | -0.04613 | NA | NA | NA | NA |
| | log Response Ratio | -0.05039 | -0.07502 | -0.01957 | -0.08990 | -0.00334 |
| log Mass | Bias Corrected log Response Ratio | 0.00466 | NA | NA | NA | NA |
| | log Response Ratio | 0.00425 | 0.00126 | 0.00768 | -0.00128 | 0.00917 |

```
Core_Comparison %>%
    mutate(Mode = as.numeric(Mode), `Low HDI (80%)` = as.numeric(`Low HDI (80%)`),
        `High HDI (80%)` = as.numeric(`High HDI (80%)`), `Low HDI (95%)` = as.numeric(`Low HDI (95%)`),
        `High HDI (95%)` = as.numeric(`High HDI (95%)`)) %>%
    ggplot(aes(x = Mode, y = Parameter, fill = `Response Variable`)) +
    geom_errorbarh(aes(y = Parameter, xmin = `Low HDI (80%)`,
        xmax = `High HDI (80%)`), colour = "black", size = 0.75,
        height = 0.3) + geom_errorbarh(aes(y = Parameter, xmin = `Low HDI (95%)`,
    xmax = `High HDI (95%)`), colour = "grey60", size = 0.75,
    height = 0.3) + geom_point(size = 3, pch = 21, colour = "black") +
    geom_vline(xintercept = 0, size = 1, colour = "black", linetype = "dashed") +
    scale_fill_manual(values = c(nice_pink, "grey80")) + facet_grid(~`Response Variable`,
    scales = "free_x") + theme_classic() + xlab("Model Coefficient") +
    theme(axis.title.y = element_blank())
```

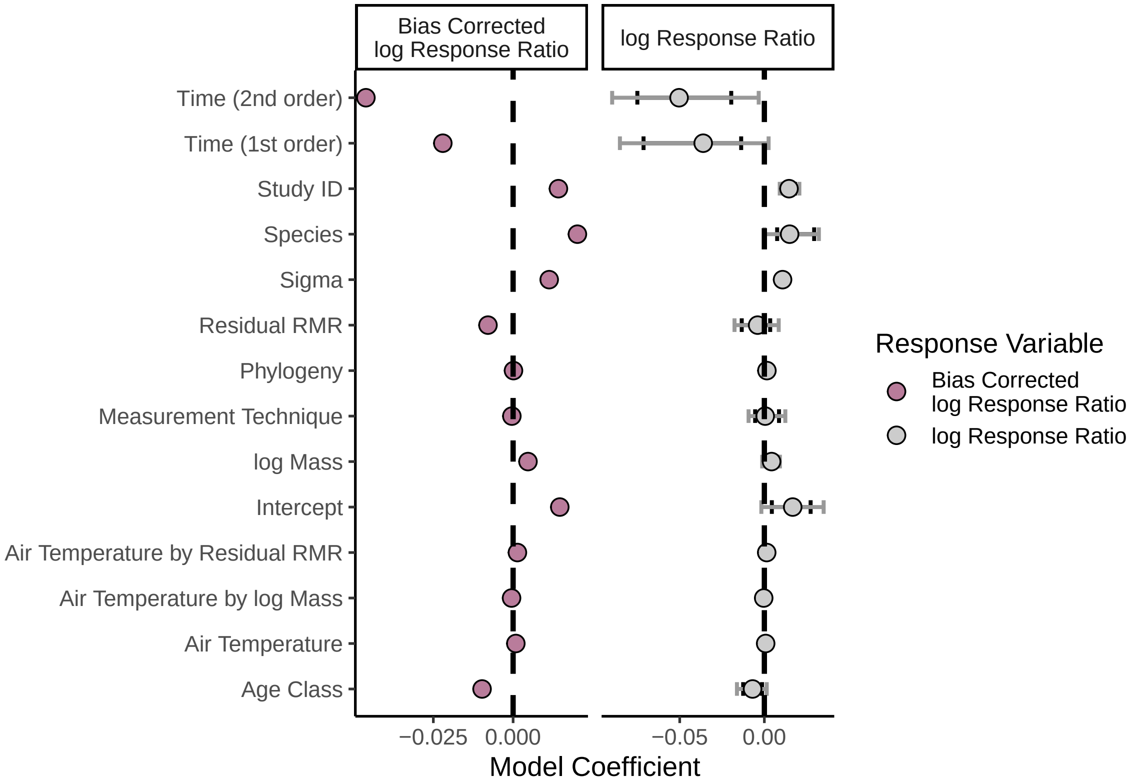


Figure 53: Predict effects of select parameters on stress-induced changes in body temperature, represented as either natural log-transformed response ratios or bias-corrected ('adjusted') log-transformed response ratios. Coefficients are derived from Bayesian linear mixed effects models predicting stress-induced changes in body temperature (either log response ratio or bias-corrected log response ratio) and represent modal values across model draws. Errorbars indicate 95 percent highest density credible intervals around modal coefficents.

```
Core_Comparison %>%
    mutate(Mode = as.numeric(Mode), `Low HDI (80%)` = as.numeric(`Low HDI (80%)`),
        `High HDI (80%)` = as.numeric(`High HDI (80%)`), `Low HDI (95%)` = as.numeric(`Low HDI (95%)`),
        `High HDI (95%)` = as.numeric(`High HDI (95%)`)) %>%
    filter(!(Parameter %in% c("Time (2nd order)", "Time (1st order)",
        "Intercept", "Species", "StudyID", "Sigma"))) %>%
    ggplot(aes(x = Mode, y = Parameter, fill = `Response Variable`)) +
    geom_errorbarh(aes(y = Parameter, xmin = `Low HDI (80%)`,
        xmax = `High HDI (80%)`), colour = "black", size = 0.75,
        height = 0.3) + geom_errorbarh(aes(y = Parameter, xmin = `Low HDI (95%)`,
    xmax = `High HDI (95%)`), colour = "grey60", size = 0.75,
    height = 0.3) + geom_point(size = 3, pch = 21, colour = "black") +
    geom_vline(xintercept = 0, size = 1, colour = "black", linetype = "dashed") +
```

```
    scale_fill_manual(values = c(nice_pink, "grey80")) + facet_grid(~`Response Variable`,
    scales = "free_x") + theme_classic() + xlab("Model Coefficient") +
    theme(axis.title.y = element_blank())
```

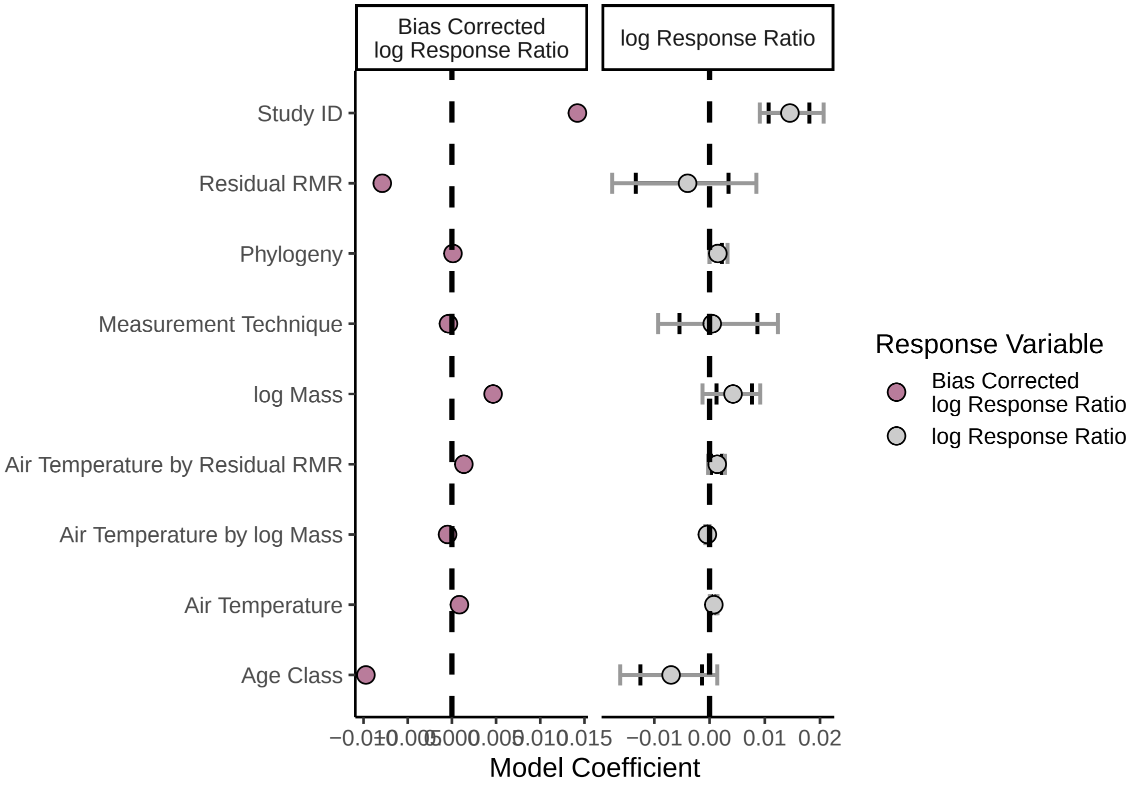


Figure 54: Predict effects of further select parameters on stress-induced changes in body temperature, represented as either natural log-transformed response ratios or bias-corrected ('adjusted') log-transformed response ratios. Coefficients are derived from Bayesian linear mixed effects models predicting stress-induced changes in body temperature (either log response ratio or bias-corrected log response ratio) and represent modal values across model draws. Errorbars indicate 95 percent highest density credible intervals around modal coefficents.

#### 8 | Plotting final results

Finally, below, we detail the methods by which we produced the final figures shown in our main text. Brief figure captions have been provided for simple interpretation, but are not equivalent to those reported in our main text. For this reason, we advise that readers seeking to understand our findings, and not simply reproduce our figures, should refer to figures and captions given in our main text.

```
# First, producing PRISMA flowchart using the R package
# 'PRISMAstatement'

found_lab <- paste0("Records identified through database searching\nusing keywords:",
    "\\'stress\\' AND\n\\'body temperature\\' NOT \\'heat stress\\'\n(n = 3022)")
screened_lab <- paste0("Abstracts screened\n(n = 3053)")
found_other_lab <- paste0("Additional records identified\nfrom",
    "reference lists\n(n = 31)")
screen_exclusions_lab <- paste0("Stress-induced changes in\ncore body temperature",
    "not\nmeasured, study species not\nendothermic, ", "pharmaceuticals\nemployed\n(n = 2917)")
fte_lab <- paste0("Full texts excluded:\ndata necessary for analysis\nnot available or\noutlying\n(n = ",
    136 - length(unique(Core_Model_New$data$StudyID)), ")")
```

```
qualitative_lab <- paste0("Studies included in\nquantitative synthesis\n(meta-regression)\n(n = ",
    length(unique(Core_Model_New$data$StudyID)), ")")

PChart <- prisma(found = 3022, found_other = 31, no_dupes = 3053,
    screened = 3053, screen_exclusions = 2917, full_text = 136,
    full_text_exclusions = 136 - length(unique(Core_Model_New$data$StudyID)),
    qualitative = length(unique(Core_Model_New$data$StudyID)),
    label = list(found = found_lab, screened = screened_lab,
        found_other = found_other_lab, screen_exclusions = screen_exclusions_lab,
        full_text_exclusions = fte_lab, qualitative = qualitative_lab))
Diagram_Mod <- gsub("qual -> quant\n    ", "", sub("(.*)quant .*",
    "\\1", PChart$x$diagram))
Diagram_Mod <- paste0(sub("(.*)\\n.*", "\\1", Diagram_Mod), "\n  }")
PChart$x$diagram <- Diagram_Mod

# Note that the packages 'rsvg' and 'DiagrammeRsvg' are
# required for the saving function used below.

require(DiagrammeRsvg)
require(rsvg)
FPath = "/Users/joshuatabh/Documents/researchProjects/trent/sihMetaregression/figures/final/Figure_2.pdf"
PRISMAstatement:::prisma_pdf(PChart, FPath)
knitr::include_graphics(path = FPath)
```

```
rm(Diagram_Mod, PChart)

# Next, producing organised forest plots for core body
# temperature responses to stress exposure

wrapper <- function(x, ...) {
    Wrap_Out <- paste(strwrap(x, width = 100, ...), collapse = "\n")
    return(Wrap_Out)
}
```

```
pal <- wes_palette("Rushmore1", n = 13, type = "continuous")
prs <- c("Sigma", "Time (1st order)", "Time (2nd order)", "Intercept", "Species", "Study ID")
xname <- "Effect on Body Temperature\nResponses to Stressors"

p1top <- Core_Model_Table %>%
  mutate(
    "Mode" = as.numeric(Mode),
    "Low HDI (80%)" = as.numeric(`Low HDI (80%)`),
    "High HDI (80%)" = as.numeric(`High HDI (80%)`),
    "Low HDI (95%)" = as.numeric(`Low HDI (95%)`),
    "High HDI (95%)" = as.numeric(`High HDI (95%)`)
  ) %>%
  filter(!Parameter %in% prs) %>%
  mutate(Parameter = ifelse(
    Parameter == "Air Temperature by log Mass",
      "Air Temperature\nby log Mass",
    ifelse(Parameter == "Residual RMR",
           "Relative Resting\nMetabolic Rate",
           ifelse(Parameter == "Air Temperature by Residual RMR",
                  "Air Temperature by\nRelative Resting\nMetabolic Rate",
                  ifelse(Parameter == "Measurement Technique",
                         "Measurement\nTechnique", Parameter))))) %>%
  mutate(Parameter = factor(Parameter,
    levels = c(
      "Age Class",
      "Air Temperature",
      "log Mass",
      "Relative Resting\nMetabolic Rate",
      "Air Temperature\nby log Mass",
      "Air Temperature by\nRelative Resting\nMetabolic Rate",
      "Measurement\nTechnique",
      "Phylogeny"
    )
  )) %>%
```

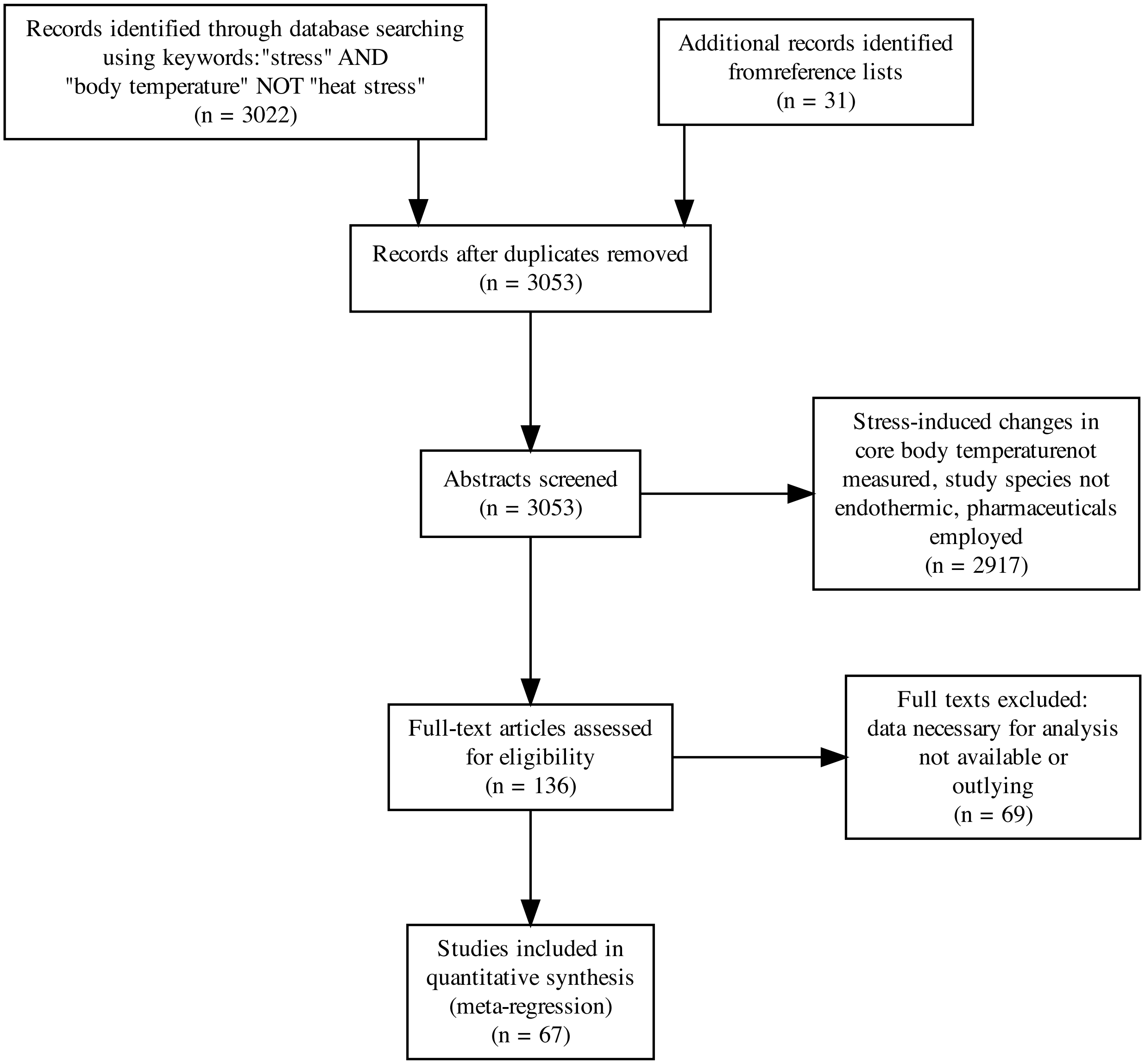


Figure 55: PRISMA flowchart detailing the process used to select studies for subsequent data collection.

```
  ggplot(aes(x = Mode, y = Parameter)) +
  geom_errorbarh(aes(
    y = Parameter, xmin = `Low HDI (95%)`,
    xmax = `High HDI (95%)`
  ),
  colour = "grey70", size = 0.75, height = 0.3, alpha = 0.7
  ) +
  geom_errorbarh(aes(
    y = Parameter, xmin = `Low HDI (80%)`,
    xmax = `High HDI (80%)`
  ),
  colour = "grey5", size = 0.75, height = 0.25, alpha = 0.7
  ) +
  geom_point(size = 3, pch = 21, colour = "black", fill = pal[7]) +
  geom_vline(xintercept = 0, size = 1, colour = "black", linetype = "dashed") +
  scale_y_discrete(limits = rev) +
  scale_x_continuous(sec.axis = sec_axis(~ ., breaks = c(-0.018, 0.013), labels = c("-", "+"),
  name = xname)) +
  theme_classic() +
  theme(
    axis.line.x.bottom = element_line(color="grey20"),
    axis.title.x.bottom = element_blank(),
    axis.title.y = element_blank(),
    legend.position = "none",
    axis.title = element_text(size = 14, family = "Noto Sans"), # Previously 12
    axis.text = element_text(size = 12, family = "Noto Sans"), # Previously 8
    axis.ticks.x.top = element_blank(),
    axis.text.x.top = element_text(size = 14, family = "Noto Sans") # Previously 12
  )

p1bottom <- Core_Model_Table %>%
  mutate(
    "Mode" = as.numeric(Mode),
    "Low HDI (80%)" = as.numeric(`Low HDI (80%)`),
    "High HDI (80%)" = as.numeric(`High HDI (80%)`),
    "Low HDI (95%)" = as.numeric(`Low HDI (95%)`),
    "High HDI (95%)" = as.numeric(`High HDI (95%)`)
  ) %>%
  filter(Parameter %in% c("Time (1st order)", "Time (2nd order)", "Intercept", "Species", "Study ID")) %>%
  mutate(Parameter = ifelse(Parameter == "Time (2nd order)", "   Time (2nd order)", Parameter)) %>%
  mutate(Parameter = factor(Parameter,
    levels = c(
      "Intercept",
      "Time (1st order)",
      "   Time (2nd order)",
      "Species",
      "Study ID"
    )
  )) %>%
  ggplot(aes(x = Mode, y = Parameter)) +
  geom_errorbarh(aes(
    y = Parameter, xmin = `Low HDI (95%)`,
    xmax = `High HDI (95%)`
  ),
  colour = "grey70", size = 0.75, height = 0.3, alpha = 0.7
  ) +
  geom_errorbarh(aes(
    y = Parameter, xmin = `Low HDI (80%)`,
    xmax = `High HDI (80%)`
  ),
  colour = "grey5", size = 0.75, height = 0.25, alpha = 0.7
  ) +
  geom_point(size = 3, pch = 21, colour = "black", fill = pal[7]) +
  geom_vline(xintercept = 0, size = 1, colour = "black", linetype = "dashed") +
  xlim(c(-0.11, 0.05)) +
  scale_y_discrete(limits = rev) +
  theme_classic() +
```

```
  xlab("Estimated Effect (Slope of lRR)") +
  theme(
    axis.title.y = element_blank(),
    legend.position = "none",
    axis.title = element_text(size = 16, family = "Noto Sans"),
    axis.text = element_text(size = 12, family = "Noto Sans")
  )

require('ggpubr')

p1 <- ggarrange(p1top, p1bottom,
          heights = c(3.5,1), ncol = 1)

dest <-"/Users/joshuatabh/Documents/researchProjects/trent/sihMetaregression/figures/final/Figure_3.jpg"
ggsave(dest, p1, width = 7.5, height = 7.5, dpi = 800)
rm(dest)

# Air temperature by body mass plot

Ambient_Mass <- expand.grid(
  "Centred_AT" = seq(min(Core_Data_MR_Shortened$Ambient.Temp.Stressor, na.rm = T),
                                max(Core_Data_MR_Shortened$Ambient.Temp.Stressor, na.rm = T),
                                by = 0.1) - mean(Core_Data_MR_Shortened$Ambient.Temp.Stressor),
  "Ambient.Temp.Stressor.sd" = 0.0001,
  "Centred_logMass" = log(c(
    10, 100, 1000, 10000
  )) - mean(Core_Data_MR_Shortened$logMass),
  "logMass_SD" = 0.0001,
  "Age_Class" = 0,
  "ResMR_Bin" = 0,
  "Centred_Time" = 0,
  "lRV" = 0.0001,
  "Technique" = 0
)

Ambient_Mass_Pred <- posterior_predict(Core_Model_New,
  resp = "lRR",
  newdata = Ambient_Mass,
  re_form = NA,
  nsamples = 2500
)

Ambient_Mass <- cbind(
  Ambient_Mass,
  as.data.frame(t(apply(Ambient_Mass_Pred,
    MARGIN = 2,
    HDInterval::hdi,
    credMass = c(0.8)
  ))) %>%
    rename("LCL_80" = "lower",
           "UCL_80" = "upper"),
  as.data.frame(t(apply(Ambient_Mass_Pred,
    MARGIN = 2,
    HDInterval::hdi,
    credMass = c(0.95)
  ))) %>%
    rename("LCL_95" = "lower",
           "UCL_95" = "upper")
)

Ambient_Mass <- cbind(Ambient_Mass,
  data.frame("Fit" = apply(Ambient_Mass_Pred,
    MARGIN = 2,
    function(x){md(round(x, digits = 5))}
  )))

rm(Ambient_Mass_Pred)
```

```
pal <- wes_palette("Rushmore1", n = 4, type = "continuous")

p2_data <- Ambient_Mass %>%
  mutate(Mass = exp(Centred_logMass + mean(Core_Data_MR_Shortened$logMass))) %>%
  mutate(Mass = as.character(Mass)) %>%
  mutate(Ambient.Temp = Centred_AT + mean(Core_Data_MR_Shortened$Ambient.Temp.Stressor)) %>%
  group_by(Mass, Ambient.Temp) %>%
  summarise("Fit" = mean(Fit),
            "LCL_80" = mean(LCL_80),
            "LCL_95" = mean(LCL_95),
            "UCL_80" = mean(UCL_80),
            "UCL_95" = mean(UCL_95))

p2 <- ggplot(p2_data, aes(
    x = Ambient.Temp, y = Fit,
    fill = Mass, linetype = Mass
  )) +
  geom_ribbon(
    data = p2_data %>%
      filter(Mass == 10000),
    aes(
    x = Ambient.Temp,
    ymin = LCL_95,
    ymax = UCL_95
  ), alpha = 0.5) +
  geom_ribbon(
    data = p2_data %>%
      filter(Mass == 10000),
    aes(
    x = Ambient.Temp,
    ymin = LCL_80,
    ymax = UCL_80
  ), alpha = 0.6) +
    geom_ribbon(
    data = p2_data %>%
      filter(Mass == 1000),
    aes(
    x = Ambient.Temp,
    ymin = LCL_95,
    ymax = UCL_95
  ), alpha = 0.5) +
  geom_ribbon(
    data = p2_data %>%
      filter(Mass == 1000),
    aes(
    x = Ambient.Temp,
    ymin = LCL_80,
    ymax = UCL_80
  ), alpha = 0.6) +
    geom_ribbon(
    data = p2_data %>%
      filter(Mass == 100),
    aes(
    x = Ambient.Temp,
    ymin = LCL_95,
    ymax = UCL_95
  ), alpha = 0.5) +
  geom_ribbon(
    data = p2_data %>%
      filter(Mass == 100),
    aes(
    x = Ambient.Temp,
    ymin = LCL_80,
    ymax = UCL_80
  ), alpha = 0.6) +
    geom_ribbon(
    data = p2_data %>%
```

```
      filter(Mass == 10),
    aes(
    x = Ambient.Temp,
    ymin = LCL_95,
    ymax = UCL_95
  ), alpha = 0.5) +
  geom_ribbon(
    data = p2_data %>%
      filter(Mass == 10),
    aes(
    x = Ambient.Temp,
    ymin = LCL_80,
    ymax = UCL_80
  ), alpha = 0.6) +
  geom_smooth(method = "lm", colour = "black", se = FALSE) +
  geom_segment(lineend = "round", linejoin = "round",
               size = 0.3, linetype = "solid", colour = "grey10",
    aes(x = 32, y = 0.01, xend = 32, yend = 0.13),
                  arrow = arrow(length = unit(0.2, "cm"))) +
  geom_segment(lineend = "round", linejoin = "round",
               size = 0.3, linetype = "solid", colour = "grey10",
    aes(x = 32, y = -0.01, xend = 32, yend = -0.14),
                  arrow = arrow(length = unit(0.2, "cm"))) +
  annotate(geom = "text", label = "Body Temperature\nIncreasing",
          x = 35, y = 0.07, size = 5, family = "Noto Sans", angle = 270) + # Size previously 4
  annotate(geom = "text", label = "Body Temperature\nDecreasing",
          x = 35, y = -0.075, size = 5, family = "Noto Sans", angle = 270) + # Size previously 4
  scale_fill_manual(values = pal, name = "Body Mass (g)") +
  scale_linetype_manual(values = c("solid", "dashed", "longdash", "twodash", "dotdash"), name = "Body Mass (g)") +
  scale_x_continuous(name = "Ambient Temperature (°C)",
                     breaks = c(-20, -10, 0, 10, 20, 30),
                     labels = c("-20", "-10", "0", "10", "20", "30")) +
  ylab("Stress-Induced Change in\nBody Temperature\n(log Response Ratio)") +
  theme_classic() +
  theme(axis.title = element_text(size = 16, family = "Noto Sans"), # Previously 14
        axis.text = element_text(size = 13), # Previously 12
        legend.title = element_text(size = 15, family = "Noto Sans"), # Previous 12
        legend.text = element_text(size = 13, family = "Noto Sans"),
        legend.position = "bottom")

dest <- "/Users/joshuatabh/Documents/researchProjects/trent/sihMetaregression/figures/final/Figure_4a.jpg"
ggsave(dest, p2, width = 9, height = 7, dpi = 800)
rm(dest)

## Descriptive panel

descData <- expand.grid("Centred_logMass" = seq(min(Core_Model_New$data$Centred_logMass),
                              max(Core_Model_New$data$Centred_logMass), by = 1),
           "Centred_AT" = seq(min(Core_Model_New$data$Centred_AT),
                              max(Core_Model_New$data$Centred_AT), by = 1),
           "Ambient.Temp.Stressor.sd" = 0.0001,
           "logMass_SD" = 0.0001,
           "Age_Class" = 0,
           "ResMR_Bin" = 0,
           "Centred_Time" = 0,
           "lRV" = 0.0001,
           "Technique" = 0
) %>% mutate("massGroup" = ifelse(Centred_logMass < mean(Core_Model_New$data$Centred_logMass), "Small", "Big"),
             "tempGroup" = ifelse(Centred_AT < 0, "Cold", "Warm")) %>%
  arrange(massGroup, tempGroup)

pDescRaw <- predict(Core_Model_New, newdata = descData, resp = "lRR", re_form = NA, summary = FALSE)

SC <- which(descData$massGroup == "Small" & descData$tempGroup == "Cold")
BC <- which(descData$massGroup == "Big" & descData$tempGroup == "Cold")
SW <- which(descData$massGroup == "Small" & descData$tempGroup == "Warm")
BW <- which(descData$massGroup == "Big" & descData$tempGroup == "Warm")
```

```
pDesc <- rbind(data.frame("Pred" = c(pDescRaw[, SC]),
                          "massGroup" = "Small", "tempGroup" = "Cold"),
               data.frame("Pred" = c(pDescRaw[, BC]),
                          "massGroup" = "Big", "tempGroup" = "Cold"),
               data.frame("Pred" = c(pDescRaw[, SW]),
                          "massGroup" = "Small", "tempGroup" = "Warm"),
               data.frame("Pred" = c(pDescRaw[, BW]),
                          "massGroup" = "Big", "tempGroup" = "Warm")
               ) %>%
  mutate("predRaw" = exp(Pred)*38 - 38) %>%
  group_by(massGroup, tempGroup) %>%
  summarise(meanPred = mean(predRaw),
            sdPred = sd(predRaw),
            LCL80 = as.numeric(bayestestR::hdi(predRaw, ci = 0.8)[2]),
            UCL80 = as.numeric(bayestestR::hdi(predRaw, ci = 0.8)[3]),
            LCL95 = as.numeric(bayestestR::hdi(predRaw)[2]),
            UCL95 = as.numeric(bayestestR::hdi(predRaw)[3])
            ) %>%
  left_join(., Core_Data_MR_Shortened %>%
              mutate("massGroup" = ifelse(Centred_logMass < 0, "Small", "Big"),
             "tempGroup" = ifelse(Centred_AT < 0, "Cold", "Warm")) %>%
              group_by(massGroup, tempGroup) %>%
              count())
rm(SC,BC,SW,BW)

pal <- wes_palette("Rushmore", n = 12, "continuous")[c(3,8)]
meanCLM <- mean(Core_Model_New$data$Centred_logMass)

p2b <- pDesc %>%
  as.data.frame() %>%
  mutate(tempGroup = ifelse(tempGroup == "Cold", "Cool\n(<20.1°C)", "Warm\n(>20.1°C)"),
         massGroup = ifelse(massGroup == "Big", "Large (>222 g)", "Small (<222 g)")) %>%
  mutate(massGroup = factor(massGroup, levels = c("Small (<222 g)", "Large (>222 g)"))) %>%
ggplot(aes(x = tempGroup, y = meanPred, fill = massGroup, colour = massGroup)) +
  geom_point(data = Core_Model_New$data %>%
               mutate("massGroup" = ifelse(Centred_logMass < meanCLM, "Small (<222 g)", "Large (>222 g)"),
             "tempGroup" = ifelse(Centred_AT < 0, "Cool\n(<20.1°C)", "Warm\n(>20.1°C)"),
             "meanPred" = exp(lRR)*38 - 38) %>%
               mutate(massGroup = factor(massGroup, levels = c("Small (<222 g)", "Large (>222 g)"))),
             size = 2, pch = 21, alpha = 0.6, position = position_jitterdodge(jitter.width = 0.2, dodge.width = 0.5)) +
  geom_errorbar(aes(x = tempGroup, ymin = LCL95, ymax = UCL95),
                colour = "grey70", width = 0.2, position = position_dodge(width = 0.5)) +
  geom_errorbar(aes(x = tempGroup, ymin = LCL80, ymax = UCL80),
                colour = "black", width = 0.1, position = position_dodge(width = 0.5)) +
  geom_point(size = 5, pch = 21, colour = "black", position = position_dodge(width = 0.5)) +
  theme_classic() +
  scale_fill_manual(values = pal, name = "Body Mass") +
  scale_colour_manual(values = pal, name = "Body Mass") +
  ylab("Change in\nBody Temperature (°C)") +
  xlab("Ambient Temperature") +
  theme(axis.title = element_text(size = 16, family = "Noto Sans"),
        axis.text = element_text(size = 13),
        legend.title = element_text(size = 15, family = "Noto Sans"),
        legend.text = element_text(size = 13, family = "Noto Sans"))

dest <- "/Users/joshuatabh/Documents/researchProjects/trent/sihMetaregression/figures/final/Figure_4b.jpg"
ggsave(dest, p2b, width = 8, height = 7, dpi = 800)
rm(dest)

# Panel for change in metabolic rate panel

energeticConsequences = c()
for (i in 1:nrow(Core_Data_MR_Shortened)){
  if (!is.na(Core_Data_MR_Shortened$Initial.Tb[i])){
    tBase = Core_Data_MR_Shortened$Initial.Tb[i]
  } else {
    tBase = Core_Data_MR_Shortened$Control.Tb.at.Stress.Peak[i]
```

```
  }
  tResponse = exp(Core_Data_MR_Shortened$lRR[i])*tBase

  if (Core_Data_MR_Shortened$Class[i] == "Aves"){
    shape = 1.25
    maxCore = 45
  } else if (Core_Data_MR_Shortened$Class[i] == "Mammalia"){
    shape = 1.25
    maxCore = 40
  }
  depth = log10(Core_Data_MR_Shortened$Mass[i])/100

  basalModel = endoR(TA = Core_Data_MR_Shortened$Ambient.Temp.Stressor[i],
    AMASS = Core_Data_MR_Shortened$Mass[i]/1000, TC= tBase, TC_MAX = maxCore,
    VEL = 0.1, RH = 50, SHAPE = 4, SHAPE_B = shape, FURTHRMK = 0.0272, SHADE = 50,
    ZFURD = depth, ZFURV = depth
  )
  basal = as.data.frame(basalModel$enbal)$QGEN

  responseModel = endoR(TA = Core_Data_MR_Shortened$Ambient.Temp.Stressor[i],
    AMASS = Core_Data_MR_Shortened$Mass[i]/1000, TC= tResponse, TC_MAX = maxCore,
    VEL = 0.1, RH = 50, SHAPE = 4, SHAPE_B = shape, FURTHRMK = 0.0272, SHADE = 50,
    ZFURD = depth, ZFURV = depth
  )
  response = as.data.frame(responseModel$enbal)$QGEN
  energeticConsequences[i] = (response/basal)*100 - 100
}

Core_Data_MR_Shortened$energeticConsequences = energeticConsequences

# Quickly summarising energetic consequences of stress-induced thermal responses

mean(Core_Data_MR_Shortened$energeticConsequences, na.rm = T)
```

```
## [1] -4.802855
```

```
sd(Core_Data_MR_Shortened$energeticConsequences, na.rm = T)
```

```
## [1] 7.129824
```

```
min(Core_Data_MR_Shortened$energeticConsequences, na.rm = T)
```

```
## [1] -24.44755
```

```
Core_Data_MR_Shortened %>%
  mutate("massGroup" = ifelse(Centred_logMass < mean(Core_Model_New$data$Centred_logMass),
                                      "Small (<222 g)",
                                      "Large (>222 g)"),
              "tempGroup" = ifelse(Centred_AT < 0, "Cool\n(<20.1°C)", "Warm\n(>20.1°C)")) %>%
  group_by(massGroup, tempGroup) %>%
  summarise("Mean" = mean(energeticConsequences, na.rm = T),
            "SD" = sd(energeticConsequences, na.rm = T))
```

```
## # A tibble: 4 x 4
## # Groups:   massGroup [2]
##   massGroup      tempGroup           Mean    SD
##   <chr>          <chr>              <dbl> <dbl>
## 1 Large (>222 g) "Cool\n(<20.1°C)" -6.65   8.38
## 2 Large (>222 g) "Warm\n(>20.1°C)" -6.44   4.15
## 3 Small (<222 g) "Cool\n(<20.1°C)" -5.21   4.83
## 4 Small (<222 g) "Warm\n(>20.1°C)" -0.326  9.61
```

```
pal <- wes_palette("Rushmore", n = 12, "continuous")[c(3,8)]

p2c = ggplot(Core_Data_MR_Shortened %>%
       mutate("massGroup" = ifelse(Centred_logMass < mean(Core_Model_New$data$Centred_logMass),
                                      "Small (<222 g)",
                                      "Large (>222 g)"),
              "tempGroup" = ifelse(Centred_AT < 0, "Cool\n(<20.1°C)", "Warm\n(>20.1°C)")) %>%
       mutate(massGroup = factor(massGroup, levels = c("Small (<222 g)", "Large (>222 g)"))),
```

```
      aes(x = tempGroup, y = energeticConsequences, fill = massGroup, colour = massGroup)) +
    geom_point(size = 2, colour = "black", pch = 21, alpha = 0.2,
               position = position_jitterdodge(jitter.width = 0.25, dodge.width = 0.25)) +
    stat_summary(geom = "errorbar",
                 fun.data = function(x){CIs = (sd(x)/sqrt(length(x)))*1.96;
                 list("ymin" = mean(x) - CIs, "ymax" = mean(x) + CIs)},
                 width = 0.2, colour = "grey70",
                 position = position_dodge(width = 0.25)) +
    stat_summary(geom = "errorbar",
                 fun.data = function(x){sem = sd(x)/sqrt(length(x));
                 list("ymin" = mean(x) - sem, "ymax" = mean(x) + sem)},
                 width = 0.1, colour = "black",
                 position = position_dodge(width = 0.25)) +
    stat_summary(geom = "point", colour = "black", fun = "mean", size = 4, pch = 21,
                 position = position_dodge(width = 0.25)) +
    geom_hline(yintercept = 0, linetype = "dashed", colour = "black") +
    scale_fill_manual(values = pal, name = "Body Mass") +
    scale_colour_manual(values = pal, name = "Body Mass") +
    xlab("Ambient Temperature") +
    ylab("Relative Change in\nEnergy Expenditure (%)") +
    theme_classic() +
    theme(axis.title = element_text(size = 16, family = "Noto Sans"),
        axis.text = element_text(size = 13),
        legend.title = element_text(size = 15, family = "Noto Sans"),
        legend.text = element_text(size = 13, family = "Noto Sans"))

# Identifying point with an ~30% increase in expenditure.

Core_Data_MR_Shortened[c(with(Core_Data_MR_Shortened, which(energeticConsequences > 28 & Ambient.Temp.Stressor > 20.1))), "Obser
```

```
## [1] "GG"
```

```
dest <- "/Users/joshuatabh/Documents/researchProjects/trent/sihMetaregression/figures/final/Figure_4c.jpg"
ggsave(dest, p2c, width = 6, height = 6, dpi = 800)
rm(dest)

p2full <- ggarrange(p2,
                    ggarrange(p2b, p2c, ncol = 2, labels = c("B", "C"), common.legend = TRUE,
                              legend = "right"),
                    ncol = 1, heights = c(1.7,1), labels = "A")

dest <- "/Users/joshuatabh/Documents/researchProjects/trent/sihMetaregression/figures/final/Figure_4.jpg"
ggsave(dest, p2full, height = 10, width = 10, dpi = 800)
rm(dest)

# Air temperature by relative resting metabolic rate

Ambient_MR <- expand.grid(
  "Centred_AT" = seq(min(Core_Data_MR_Shortened$Ambient.Temp.Stressor),
                                max(Core_Data_MR_Shortened$Ambient.Temp.Stressor),
                                by = 0.1) - mean(Core_Data_MR_Shortened$Ambient.Temp.Stressor),
  "Ambient.Temp.Stressor.sd" = 0.0001,
  "Centred_logMass" = 0,
  "logMass_SD" = 0.0001,
  "Age_Class" = 0,
  "ResMR_Bin" = c(-0.5, 0.5),
  "Centred_Time" = 0,
  "lRV" = 0.0001,
  "Technique" = 0
)

Ambient_MR_Pred <- posterior_predict(Core_Model_New,
  resp = "lRR",
  newdata = Ambient_MR,
  re_form = NA,
  nsamples = 2500
)
```

```
Ambient_MR <- cbind(
  Ambient_MR,
  as.data.frame(t(apply(Ambient_MR_Pred,
    MARGIN = 2,
    HDInterval::hdi,
    credMass = c(0.8)
  ))) %>%
    rename("LCL_80" = "lower",
           "UCL_80" = "upper"),
  as.data.frame(t(apply(Ambient_MR_Pred,
    MARGIN = 2,
    HDInterval::hdi,
    credMass = c(0.95)
  ))) %>%
    rename("LCL_95" = "lower",
           "UCL_95" = "upper")
)

Ambient_MR <- cbind(Ambient_MR,
  data.frame("Fit" = apply(Ambient_MR_Pred,
    MARGIN = 2,
    function(x){md(round(x, digits = 5))}
  )))

rm(Ambient_MR_Pred)

pal <- wes_palette("Rushmore1", n = 10, type = "continuous")

p3_Data <-Ambient_MR %>%
  mutate(Ambient.Temp = Centred_AT + mean(Core_Data_MR_Shortened$Ambient.Temp.Stressor)) %>%
  mutate(ResMR_Bin = ifelse(ResMR_Bin == -0.5, "Low", "High")) %>%
  mutate(ResMR_Bin = factor(ResMR_Bin, levels = c("Low", "High"))) %>%
  group_by(ResMR_Bin, Ambient.Temp) %>%
  summarise("Fit" = mean(Fit),
            "LCL_80" = mean(LCL_80),
            "LCL_95" = mean(LCL_95),
            "UCL_80" = mean(UCL_80),
            "UCL_95" = mean(UCL_95))

p3 <- ggplot(p3_Data %>%
               mutate(factor(ResMR_Bin, levels = "Low", "High")),
             aes(
    x = Ambient.Temp, y = Fit,
    fill = ResMR_Bin, linetype = ResMR_Bin
  )) +
  geom_ribbon(aes(
    x = Ambient.Temp,
    ymin = LCL_95,
    ymax = UCL_95
  ), alpha = 0.5) +
  geom_ribbon(aes(
    x = Ambient.Temp,
    ymin = LCL_80,
    ymax = UCL_80
  ), alpha = 0.6) +
  geom_smooth(method = "lm", colour = "black", se = FALSE) +
  geom_segment(lineend = "round", linejoin = "round",
               size = 0.3, linetype = "solid", colour = "grey10",
    aes(x = 32, y = 0.005, xend = 32, yend = 0.07),
                  arrow = arrow(length = unit(0.2, "cm"))) +
  geom_segment(lineend = "round", linejoin = "round",
               size = 0.3, linetype = "solid", colour = "grey10",
    aes(x = 32, y = -0.005, xend = 32, yend = -0.08),
                  arrow = arrow(length = unit(0.2, "cm"))) +
  annotate(geom = "text", label = "Body Temperature\nIncreasing",
           x = 35, y = 0.04, size = 5, family = "Noto Sans", angle = 270) + # Size previously 4
  annotate(geom = "text", label = "Body Temperature\nDecreasing",
```

```
           x = 35, y = -0.045, size = 5, family = "Noto Sans", angle = 270) +
  scale_x_continuous(name = "Ambient Temperature (°C)",
                     breaks = c(-20, -10, 0, 10, 20, 30),
                     labels = c("-20", "-10", "0", "10", "20", "30")) +
  scale_fill_manual(values = pal[c(7,9)], name = "Relative Energy\nExpenditure at Rest") +
  scale_linetype_manual(values = c("dashed", "solid"), name = "Relative Energy\nExpenditure at Rest") +
  xlab("Ambient Temperature (°C)") +
  ylab("Stress-Induced Change in\nBody Temperature\n(log Response Ratio)") +
  theme_classic() +
  theme(axis.title = element_text(size = 16, family = "Noto Sans"),
        axis.text = element_text(size = 13),
        legend.title = element_text(size = 15, family = "Noto Sans"),
        legend.text = element_text(size = 13, family = "Noto Sans"),
        legend.position = "bottom")

dest <- "/Users/joshuatabh/Documents/researchProjects/trent/sihMetaregression/figures/final/Figure_5a.jpg"
ggsave(dest, p3, width = 9, height = 7, dpi = 800)
rm(dest)

## Descriptive Panel

mrDescData <- expand.grid("Centred_logMass" = 0,
           "Centred_AT" = seq(min(Core_Model_New$data$Centred_AT),
                              max(Core_Model_New$data$Centred_AT), by = 1),
           "Ambient.Temp.Stressor.sd" = 0.0001,
           "logMass_SD" = 0.0001,
           "Age_Class" = 0,
           "ResMR_Bin" = c(-0.5, 0.5),
           "Centred_Time" = 0,
           "lRV" = 0.0001,
           "Technique" = 0
) %>% mutate("tempGroup" = ifelse(Centred_AT < 0, "Cold", "Warm")) %>%
  arrange(ResMR_Bin, tempGroup)

pmrDescRaw <- predict(Core_Model_New, newdata = mrDescData, resp = "lRR", re_form = NA, summary = FALSE)

LC <- which(mrDescData$ResMR_Bin == -0.5 & mrDescData$tempGroup == "Cold")
HC <- which(mrDescData$ResMR_Bin == 0.5 & mrDescData$tempGroup == "Cold")
LW <- which(mrDescData$ResMR_Bin == -0.5 & mrDescData$tempGroup == "Warm")
HW <- which(mrDescData$ResMR_Bin == 0.5 & mrDescData$tempGroup == "Warm")

pmrDesc <- rbind(data.frame("Pred" = c(pmrDescRaw[, LC]),
                            "ResMR_Bin" = "Low", "tempGroup" = "Cold"),
               data.frame("Pred" = c(pmrDescRaw[, HC]),
                          "ResMR_Bin" = "High", "tempGroup" = "Cold"),
               data.frame("Pred" = c(pmrDescRaw[, LW]),
                          "ResMR_Bin" = "Low", "tempGroup" = "Warm"),
               data.frame("Pred" = c(pmrDescRaw[, HW]),
                          "ResMR_Bin" = "High", "tempGroup" = "Warm")
               ) %>%
  mutate("predRaw" = exp(Pred)*38 - 38) %>%
  group_by(ResMR_Bin, tempGroup) %>%
  summarise(meanPred = mean(predRaw),
            sdPred = sd(predRaw),
            LCL80 = as.numeric(bayestestR::hdi(predRaw, ci = 0.8)[2]),
            UCL80 = as.numeric(bayestestR::hdi(predRaw, ci = 0.8)[3]),
            LCL95 = as.numeric(bayestestR::hdi(predRaw)[2]),
            UCL95 = as.numeric(bayestestR::hdi(predRaw)[3])
            ) %>%
  left_join(., Core_Data_MR_Shortened %>%
              mutate("tempGroup" = ifelse(Centred_AT < 0, "Cold", "Warm")) %>%
              mutate("ResMR_Bin" = ifelse(ResMR_Bin == -0.5, "Low", "High")) %>%
              group_by(ResMR_Bin, tempGroup) %>%
              count())

p3b <- pmrDesc %>%
  ungroup() %>%
```

```
  as.data.frame() %>%
  mutate(tempGroup = ifelse(tempGroup == "Cold", "Cool\n(<20.1°C)", "Warm\n(>20.1°C)"),
         ResMR_Bin = factor(ResMR_Bin, levels = c("Low", "High"))) %>%
ggplot(aes(x = tempGroup, y = meanPred, fill = ResMR_Bin, colour = ResMR_Bin)) +
  geom_point(data = Core_Model_New$data %>%
               mutate("tempGroup" = ifelse(Centred_AT < 0, "Cool\n(<20.1°C)", "Warm\n(>20.1°C)"),
             "meanPred" = exp(lRR)*38 - 38,
             "ResMR_Bin" = ifelse(ResMR_Bin == -0.5, "Low", ifelse(ResMR_Bin == 0.5, "High", NA))) %>%
               mutate(ResMR_Bin = factor(ResMR_Bin, levels = c("Low", "High"))),
             size = 2, pch = 21, alpha = 0.3, position = position_jitterdodge(jitter.width = 0.2, dodge.width = 0.5)) +
  geom_errorbar(aes(x = tempGroup, ymin = LCL95, ymax = UCL95),
                colour = "grey50", width = 0.2, position = position_dodge(width = 0.5)) +
  geom_errorbar(aes(x = tempGroup, ymin = LCL80, ymax = UCL80),
                colour = "black", width = 0.1, position = position_dodge(width = 0.5)) +
  geom_point(size = 5, pch = 21, colour = "black", position = position_dodge(width = 0.5)) +
  theme_classic() +
  scale_fill_manual(values = pal[c(7,9)], name = "Relative Energy\nExpenditure at Rest") +
  scale_colour_manual(values = pal[c(7,9)], name = "Relative Energy\nExpenditure at Rest") +
  ylab("Change in\nBody Temperature (°C)") +
  xlab("Ambient Temperature") +
  theme(axis.title = element_text(size = 16, family = "Noto Sans"),
        axis.text = element_text(size = 13),
        legend.title = element_text(size = 15, family = "Noto Sans"),
        legend.text = element_text(size = 13, family = "Noto Sans"))

dest <- "/Users/joshuatabh/Documents/researchProjects/trent/sihMetaregression/figures/final/Figure_5b.jpg"
ggsave(dest, p3b, width = 8, height = 7, dpi = 800)
rm(dest)

# Change in energy expenditure panel

pal <- wes_palette("Rushmore1", n = 10, type = "continuous")

p3c = ggplot(Core_Data_MR_Shortened %>%
       mutate("tempGroup" = ifelse(Centred_AT < 0, "Cool\n(<20.1°C)", "Warm\n(>20.1°C)")) %>%
       mutate(tempGroup = factor(tempGroup, levels = c("Cool\n(<20.1°C)", "Warm\n(>20.1°C)")),
              ResMR_Bin = ifelse(ResMR_Bin == -0.5, "Low", "High")) %>%
        mutate(ResMR_Bin = factor(ResMR_Bin, levels = c("Low", "High"))),
        aes(x = tempGroup, y = energeticConsequences, fill = ResMR_Bin, colour = ResMR_Bin)) +
    geom_point(size = 2, colour = "black", pch = 21, alpha = 0.2,
               position = position_jitterdodge(jitter.width = 0.25, dodge.width = 0.25)) +
    stat_summary(geom = "errorbar",
                 fun.data = function(x){CIs = (sd(x)/sqrt(length(x)))*1.96;
                 list("ymin" = mean(x) - CIs, "ymax" = mean(x) + CIs)},
                 width = 0.2, colour = "grey70",
                 position = position_dodge(width = 0.25)) +
    stat_summary(geom = "errorbar",
                 fun.data = function(x){sem = sd(x)/sqrt(length(x));
                 list("ymin" = mean(x) - sem, "ymax" = mean(x) + sem)},
                 width = 0.1, colour = "black",
                 position = position_dodge(width = 0.25)) +
    stat_summary(geom = "point", colour = "black", fun = "mean", size = 4, pch = 21,
                 position = position_dodge(width = 0.25)) +
    geom_hline(yintercept = 0, linetype = "dashed", colour = "black") +
    scale_fill_manual(values = pal[c(7,9)], name = "Relative Energy\nExpenditure at Rest") +
    scale_colour_manual(values = pal[c(7,9)], name = "Relative Energy\nExpenditure at Rest") +
    xlab("Ambient Temperature") +
    ylab("Relative Change in\nEnergy Expenditure (%)") +
    scale_y_continuous(limits = c(-25, 25)) +
    theme_classic() +
    theme(axis.title = element_text(size = 16, family = "Noto Sans"),
        axis.text = element_text(size = 13),
        legend.title = element_text(size = 15, family = "Noto Sans"),
        legend.text = element_text(size = 13, family = "Noto Sans"))

dest <- "/Users/joshuatabh/Documents/researchProjects/trent/sihMetaregression/figures/final/Figure_5c.jpg"
ggsave(dest, p3c, width = 6, height = 6, dpi = 800)
```

```
rm(dest)

p3full <- ggarrange(p3,
                    ggarrange(p3b, p3c, ncol = 2, labels = c("B", "C"), common.legend = TRUE,
                              legend = "right"),
                    ncol = 1, heights = c(1.7,1), labels = "A")

dest <- "/Users/joshuatabh/Documents/researchProjects/trent/sihMetaregression/figures/final/Figure_5.jpg"
ggsave(dest, p3full, height = 10, width = 10, dpi = 800)
rm(dest)

# Summarising energetic consequences again.

Core_Data_MR_Shortened %>%
       mutate("tempGroup" = ifelse(Centred_AT < 0, "Cool\n(<20.1°C)", "Warm\n(>20.1°C)")) %>%
       mutate(tempGroup = factor(tempGroup, levels = c("Cool\n(<20.1°C)", "Warm\n(>20.1°C)")),
              ResMR_Bin = ifelse(ResMR_Bin == -0.5, "Low", "High")) %>%
        mutate(ResMR_Bin = factor(ResMR_Bin, levels = c("Low", "High"))) %>%
  group_by(tempGroup, ResMR_Bin) %>%
  summarise("Mean" = mean(energeticConsequences, na.rm = T),
            "SD" = sd(energeticConsequences, na.rm = T))
```

```
## # A tibble: 4 x 4
## # Groups:   tempGroup [2]
##   tempGroup         ResMR_Bin  Mean    SD
##   <fct>             <fct>     <dbl> <dbl>
## 1 "Cool\n(<20.1°C)" Low       -7.74  8.10
## 2 "Cool\n(<20.1°C)" High      -5.20  7.26
## 3 "Warm\n(>20.1°C)" Low       -3.08  9.04
## 4 "Warm\n(>20.1°C)" High      -5.81  3.82
```

```
## Time plot

Time_Data <- expand.grid(
  "Centred_AT" = 0,
  "Ambient.Temp.Stressor.sd" = 0.0001,
  "Centred_logMass" = 0,
  "logMass_SD" = 0.0001,
  "Age_Class" = 0,
  "ResMR_Bin" = 0,
  "Centred_Time" = c(seq(min(Core_Data_MR_Shortened$Centred_Time),
                         max(Core_Data_MR_Shortened$Centred_Time),
                         by = 10)),
  "lRV" = 0.0001,
  "Technique" = 0
)

Time_Pred <- posterior_predict(Core_Model_New,
  resp = "lRR",
  newdata = Time_Data,
  re_form = NA,
  nsamples = 2500
)

Time_Data <- cbind(
  Time_Data,
  as.data.frame(t(apply(Time_Pred,
    MARGIN = 2,
    HDInterval::hdi,
    credMass = c(0.8)
  ))) %>%
    rename("LCL_80" = "lower",
           "UCL_80" = "upper"),
  as.data.frame(t(apply(Time_Pred,
    MARGIN = 2,
    HDInterval::hdi,
    credMass = c(0.95)
  ))) %>%
```

```
    rename("LCL_95" = "lower",
           "UCL_95" = "upper")
)

Time_Data <- cbind(Time_Data,
  data.frame("Fit" = apply(Time_Pred,
    MARGIN = 2,
    function(x){md(round(x, digits = 5))}
  )))

rm(Time_Pred)

p4 <- Time_Data %>%
  mutate(Time = Centred_Time + mean(Core_Data_MR_Shortened$Time.to.Peak.s.)) %>%
  ggplot(aes(
    x = Time, y = Fit
  )) +
  geom_ribbon(aes(
    x = Time,
    ymin = LCL_95,
    ymax = UCL_95
  ), alpha = 0.5,
  fill = "#E1BD6D") +
  geom_ribbon(aes(
    x = Time,
    ymin = LCL_80,
    ymax = UCL_80
  ), alpha = 0.6,
  fill = "#E1BD6D") +
  geom_smooth(method = "lm", formula = y ~ poly(x, 2), colour = "black", linetype = "dashed", se = FALSE) +
  geom_segment(lineend = "round", linejoin = "round",
               size = 0.3, linetype = "solid", colour = "grey10",
    aes(x = 11000, y = 0.005, xend = 11000, yend = 0.06),
                  arrow = arrow(length = unit(0.2, "cm"))) +
  geom_segment(lineend = "round", linejoin = "round",
               size = 0.3, linetype = "solid", colour = "grey10",
    aes(x = 11000, y = -0.005, xend = 11000, yend = -0.065),
                  arrow = arrow(length = unit(0.2, "cm"))) +
  annotate(geom = "text", label = "Body Temperature\nIncreasing",
           x = 11500, y = 0.028, size = 5, family = "Noto Sans", angle = 270) +
  annotate(geom = "text", label = "Body Temperature\nDecreasing",
           x = 11500, y = -0.03, size = 5, family = "Noto Sans", angle = 270) +
    scale_x_continuous(name = "Time Since Stress Exposure (s)",
                       breaks = c(0, 2500, 5000, 7500, 10000),
                       labels = c("0", "2500", "5000", "7500", "10000")) +
  ylab("Stress-Induced Change in\nBody Temperature\n(log Response Ratio)") +
  theme_classic() +
  theme(axis.title = element_text(size = 16, family = "Noto Sans"),
        axis.text = element_text(size = 13),
        legend.position = "none")

dest <- "/Users/joshuatabh/Documents/researchProjects/trent/sihMetaregression/figures/final/Figure_6.jpg"
p4
```

```
ggsave(dest, p4, width = 9, height = 7, dpi = 800)
```

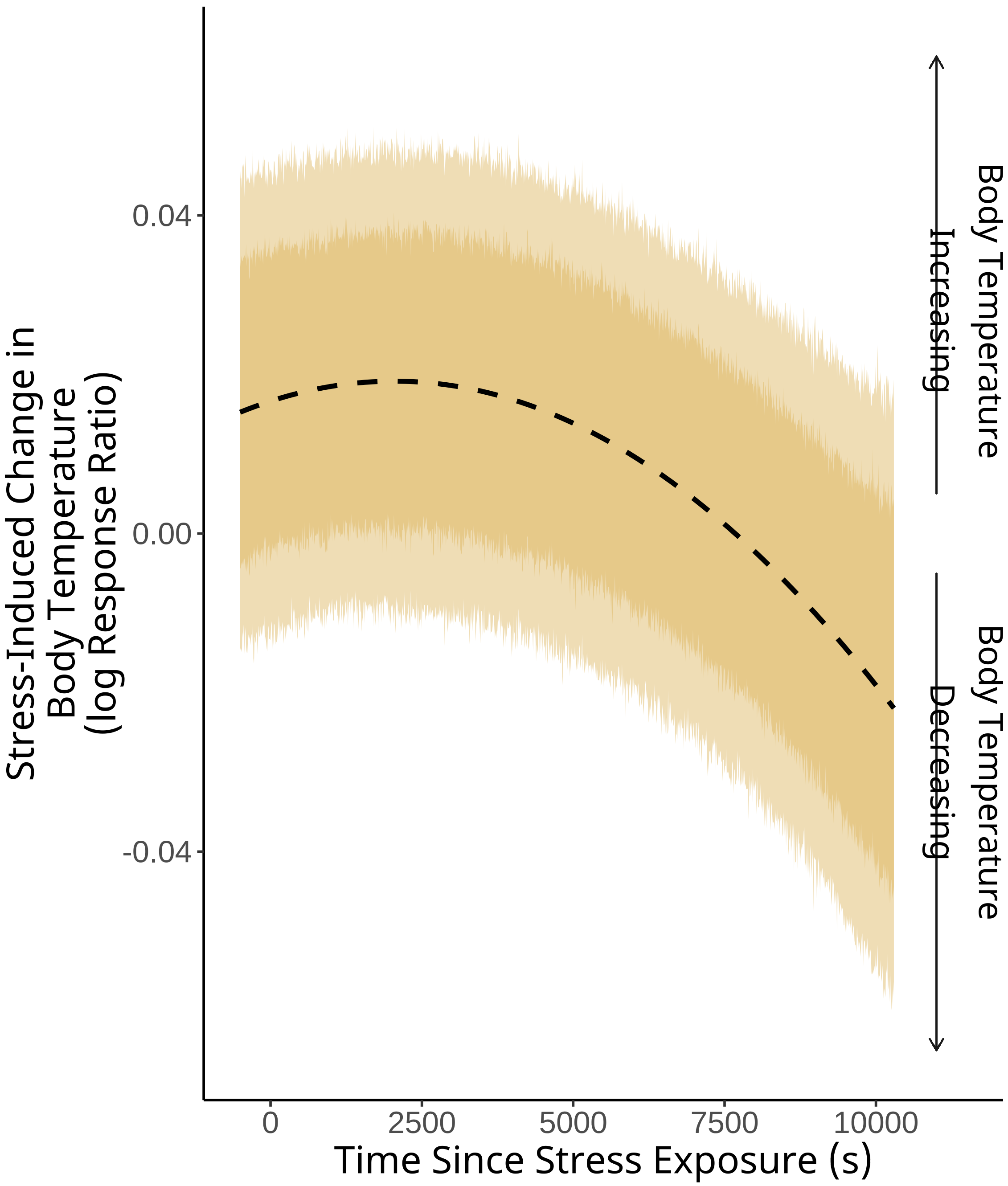


Figure 56: Effect of time post-stress exposure on stress-induced changes in body temperature. Time post stress-exposure is represented by the latency between stress exposure treatment and the measurement of a maximum or minimum body temperature measurement (in seconds; s). Negative log response ratios represent stress-induced declines in body temperature, while positive log response ratios represent stress-induced increases in body temperature. The dark ribbon represents 80 percent highest posterior density intervals (HPDIs) and the pale ribbons represent 95 percent HPDIs. The trend lines and credible intervals are marginalized across all other model predictors.

ranging leaf-eared mouse. *Evolutionary Ecology Research, 9(1)*: 547-554.

Bürkner, P.C. (2018). Advanced Bayesian multilevel modeling with the R package brms. *The R Journal, 10(1)*: 395-411.

Brase, C.H., Brase, C.P. (2017). Understandable statistics: concepts and methods (AP edition). *Cengage Learning*, Boston, United States of America.

Briese, E. (1991). Cold increases and warmth diminishes stress-induced rise of colonic temperature in rats. *Physiology and Behaviour, 51(1)*: 881-883.

Briese, E., Cabanac, M. (1991). Stress hyperthermia: physiological arguments that it is a fever. *Physiology and Behavior, 49(6)*: 1153-1157.

Buchanan, A.R., Hill, R.M. (1947).Temperature regulation in albino rats correlated with determinations of myelin density in the hypothalamus. *Proceedings of the Society for Experimental Biology and Medicine, 66(3)*: 602-608.

Careau, V. (2017). Energy intake, basal metabolic rate, and within-individual trade-offs in men and women training for a half marathon: a reanalysis. *Physiological and Biochemical Zoology 90(3)*:392-398.

Drugan, R.C., Eren, S., Hazi, A., Silva, J., Christianson, J.P., Kent, S. (2005). Impact of water temperature and stressor controllability on swim stress-induced changes in body temperature, serum corticosterone, and immobility in rats. *Pharmacology, Biochemistry and Behavior, 82(1)*: 397-403.

Ericson, P.G.P., Anderson, Britton, C.L., Eizanowski, T.A., Johansson, U.F., K{"a}llersj{"o}, M., Ohlson, J.I., Parsons, T.J., Zuccon, D., Mayr, G. (2006). Diversification of Neoaves: integration of molecular sequence data and fossils. *Biology Letters 2(1)*: 543-547.

Jerem, P., Jenni-Eiermann, S., Herborn, K., McKeegan, D., McCafferty, D.J., Nager, R.G. (2018). Eye region surface temperature reflects both energy reserves and circulating glucocorticoids in a wild bird. *Scientific Reports, 8(1)*: 1-10.

Jerem, P., Jenni-Eiermann, S., McKeegan, D., McCafferty, D.J., Nager, R.G. (2019). Eye region surface temperature dynamics during acute stress relate to baseline glucocorticoids independently of environmental conditions. *Physiology and Behavior, 210(1)*: 112627.

Jetz, W., Thomas, G.H., Joy, J.B., Hartmann, K., Mooers, A.O. (2012). The global diversity of birds in space and time. *Nature, 491(7424)*: 444-448.

Koteja, P. (1991). On the relation between basal and field metabolic rates in birds and mammals. *Functional Ecology, 5(1)*:56-64.

Lajeunesse, M.J. (2015). Bias and correction for the log response ratio in ecological meta-analysis. *Ecology, 96(8)*:2056-2063.

Lewden, A., Nord, A., Petit, M., Vezina, F. (2017). Body temperature responses to handling stress in wintering Black-capped Chickadees (*Poecile atricapillus L.*). *Physiology and Behavior, 179(1)*: 49-54.

Long, N.C., Vander, A.J., Kluger, M.J. (1990). Stress-induced rise of body temperature in rats is the same in warm and cool environments. *Physiology and Behavior, 47(4)*: 773-775.

Muise, K.A., Menzies, A.K., Willis, C.K. (2018). Stress-induced changes in body temperature of Silver-haired Bats (*Lasionycteris noctivagans*). *Physiology and Behavior, 194(1)*: 356-361.

Møller, A.P. (2010). Body temperature and fever in a free-living bird. *Comparative Biochemistry and Physiology Part B: Biochemistry and Molecular Biology, 156(1)*: 68-74.

Nagy, K.A. (2005). Field metabolic rate and body size. *Journal of Experimental Biology, 208(9)*: 1621-1625.

Nord, A., Folkow, L.P. (2019). Ambient temperature effects on stress-induced hyperthermia in Svalbard ptarmigan. *Biology Open, 8(6)*: bio043497.

Paradis E., Schliep K. (2019). ape 5.0: an environment for modern phylogenetics and evolutionary analyses in R. *Bioinformatics 35(1)*: 526-528.

Pough, F.H. (1980). The advantages of ectothermy for tetrapods. *The American Naturalist, 115(1)*: 92-112.

R Core Team. (2020). R: A language and environment for statistical computing. *R Foundation for Statistical Computing*, Vienna, Austria.

Robertson, J.K., Mastromonaco, G., Burness, G. (2020a). Evidence that stress-induced changes in surface temperature serve a thermoregulatory function. *Journal of Experimental Biology, 223(4)*: jeb213421.

Robertson, J.K., Mastromonaco, G.F., Burness, G. (2020b). Social hierarchy reveals thermoregulatory trade-offs in response to repeated stressors. *Journal of Experimental Biology, 223(21)*: jeb229047.

Stearns, S.C. (1992). *The evolution of life histories.* London, UK: Oxford University Press.

Szafranska, P.A., Andreasson, F., Nord, A., Nilsson, J.{AA}. (2020). Deep body and surface temperature responses to hot and cold environments in the zebra finch. *Journal of Thermal Biology, 94(1)*: 102776.

Upham, N.S., Esselstyn, J.A., Jetz, W. (2019). Inferring the mammal tree: species-level sets of phylogenies for questions in ecology, evolution, and conservation. *PLoS Biology, 17(12)*: e3000494.

White, C.R., Seymour, R.S. (2003). Mammalian basal metabolic rate is proportional to body mass 2/3. *Proceedings of the National Academy of Sciences, 100(7)*: 4046-4049.

Wilber, C.G., Robinson, P.F. (1958). Effect of restraint on body temperature in Guinea Pigs. *Journal of Applied Physiology, 12(2)*: 214-216.

Winder, L.A., White, S.A., Nord, A., Helm, B., McCafferty, D.J. (2020). Body surface temperature responses to food restriction in wild and captive great tits. *Journal of Experimental Biology, 223(8)*: jeb220046.

Yokoi, Y. (1966). Effect of ambient temperature upon emotional hyperthermia and hypothermia in rabbits. *Journal of Applied Physiology, 21(6)*: 1795-1798.
